# Generation of *C9orf72^h370^* mice, an intron 1 humanised *C9orf72* repeat-expansion knock-in model

**DOI:** 10.1101/2025.02.06.636691

**Authors:** Remya R Nair, Mireia Carcolé, David Thompson, Charlotte Tibbit, Ross McLeod, Alexander Cammack, Tatiana Jakubcova, Daniel Biggs, Matthew Wyles, Matthew Parker, Adam Caulder, Lydia Teboul, Chloe L Fisher-Ward, Ali R Awan, Michael Flower, Benjamin Davies, Adrian M Isaacs, Elizabeth MC Fisher, Thomas J Cunningham

**Affiliations:** MRC Harwell Institute, Harwell Campus, Oxfordshire, UK; Nucleic Acid Therapy Accelerator (NATA), Harwell Campus, Oxfordshire, UK; UK Dementia Research Institute at UCL and Department of Neurodegenerative Disease, UCL Queen Square Institute of Neurology, London, UK; Medical Sciences Division, Department of Pharmacology, University of Oxford, Oxford, UK; MRC Prion Unit at University College London, UCL Institute of Prion Diseases, London, UK; Wellcome Centre for Human Genetics, University of Oxford, Oxford, UK; School of Biosciences, University of Sheffield, Sheffield, UK; Oxford Nanopore Technologies, UK; The Mary Lyon Centre, MRC Harwell Institute, Harwell Campus, Oxfordshire, UK; Genomics Innovation Unit, Guy’s and St Thomas’ NHS Trust, London, UK; Comprehensive Cancer Centre, King’s College London, London, UK; Francis Crick Institute, London, UK; Department of Neuromuscular Diseases and Queen Square Motor Neuron Disease Centre, UCL Queen Square Institute of Neurology, London, UK

## Abstract

An autosomal dominant GGGGCC repeat expansion in intron 1 of the *C9orf72* gene is the most common genetic cause of both amyotrophic lateral sclerosis (ALS) and frontotemporal dementia (FTD). Here, we set out to engineer a gene targeted mouse model harbouring a pathogenic length humanised *C9orf72* repeat expansion allele, in order to model pathological mechanisms in a physiological context. In human disease, pathogenic repeats typically range from the hundreds to thousands of units in length, representing a considerable challenge for cellular and in vivo model generation given the instability of GC rich and repetitive DNA sequences during molecular cloning. To overcome this challenge, we developed new methodology to synthetically and iteratively build pure GGGGCC repeats within a linear vector system, which we then seamlessly and scarlessly embedded within the native human genomic sequence. This created a gene targeting DNA vector for homologous recombination of the human sequence in mouse embryonic stem cells. We used this novel targeting vector to generate a new gene targeted mouse allele, *C9orf72^h370^*, that for the first time has mouse *C9orf72* intron 1 scarlessly replaced with human intron 1 including a pure (GGGGCC)_370_ hexanucleotide repeat expansion. We confirm that the mouse model expresses human intron 1-derived RNA and produces dipeptide repeat proteins derived from the GGGGCC repeat expansion. We now provide this model as a new freely available resource for the field. In addition, we demonstrate the utility of our cloning method for engineering diverse repeat expansion sequences for modelling other disorders, such as Fragile X Syndrome.

## Introduction

ALS and FTD are related, incurable neurodegenerative disorders with shared genetic and pathological features. A GGGGCC (G4C2) repeat expansion in intron 1 of the *C9orf72* gene is the most frequent genetic cause of ALS, FTD, and ALS/FTD in European and North American populations (DeJesus-Hernandez et al. 2011; Renton et al. 2011). The presence of the repeat allele elicits a cascade of primary molecular events (Balendra and Isaacs 2018; Mayl, Shaw, and Lee 2021): (i) Repeat-containing sense and antisense RNA foci form in the nucleus and (less frequently) in the cytoplasm, (ii) Repeat-associated non-AUG (RAN) translation through all reading frames of sense- and antisense repeat RNA harbouring species leads to production and aggregation of five dipeptide repeat proteins (DPRs), poly-GA, GP, -GR, -PR, and -PA, in the nucleus and cytoplasm, and (iii) Repeat-mediated *C9orf72* promoter hypermethylation and transcriptional silencing results in *C9orf72* haploinsufficiency. An added layer of complexity and likely source of disease heterogeneity stems from somatic repeat instability, leading to variable repeat length between tissues and cells, and between generations (Nordin et al. 2015; van Blitterswijk et al. 2013).

RNA foci and DPRs are likely gain-of function species that are thought to disrupt fundamental cellular processes, whilst haploinsufficiency may provide additional triggers for disease or exacerbate toxic gain-of-function mechanisms (Balendra and Isaacs 2018; Mayl, Shaw, and Lee 2021; Braems, Swinnen, and Van Den Bosch 2020). However, there is no consensus as to which processes are predominately responsible for triggering neurodegeneration in patients. Of note, TDP-43 pathology (loss from the nucleus, aggregation in the cytoplasm) is a common feature in *C9orf72*-ALS/FTD and is broadly considered a central pathological event in the vast majority of all ALS cases and a significant subset of FTD cases. Understanding the physiological pathways that link *C9orf72* repeat expansion to TDP-43 pathology and neurodegeneration is critical for identifying primary therapeutic targets.

Here, we have created a strategy to generate a genetically humanised knock-in mouse harbouring the pathogenic *C9orf72* locus, to enable study of aberrant molecular events within a physiological setting, triggered by the repeat in its native genomic context. Humanising large genomic loci is challenging, requiring in the first instance engineering of bespoke DNA constructs for homologous recombination (Devoy et al. 2021). Including a large, unstable G4C2 repeat in this process presents an additional obstacle. Previously, we determined that a linear DNA vector system (pJazz) (Godiska et al. 2010) could successfully stabilise pathogenic-length G4C2 repeat sequences (Nair et al. 2021). Here, we build on this approach by devising a more tractable synthetic repeat-building methodology within this linear vector system, with a pre-designed mechanism for shuttling to gene-targeting constructs, which we demonstrate is applicable to diverse repeat sequences. Using this system, we generated a 1000x G4C2 repeat-harbouring targeting construct. We used this construct to target mouse embryonic stem cells, which successfully yielded clones for blastocyst injection and resulted in a new knock-in mouse model. This *C9orf72^h370^* mouse line, for the first time, comprises human intron 1 harbouring 370 G4C2 hexanucleotide repeats, seamlessly replacing mouse intron 1, and devoid of additional exogenous sequences, providing a refined physiological model to investigate *C9orf72*-ALS/FTD.

## Results

### De novo cloning of large repeat expansion sequences

We adapted the concept of recursive cloning in a circular vector (Meyer and Chilkoti 2002; Mizielinska et al. 2014) to the linear pJazz vector system, with refinements. We first cloned a small cassette into pJazz harbouring four G4C2 repeat units and flanked by XmaI and AarI restriction sites positioned to cleave the repeat ends to generate 5’-CCGG overhangs (thus enabling isolation of a pure G4C2 sequence) (**Figure 1A**). Repeat doubling consisted of cleaving the vector in separate digestion reactions with XmaI and AarI, respectively, and subsequently ligating together the repeat containing short and long arm fragments (**Figure 1B,C**). We successfully performed eight rounds of repeat doubling, synthesising an excisable, pure 1024-repeat sequence (pJazz-V1; **Figure 1D-E**).

**Figure 1.**
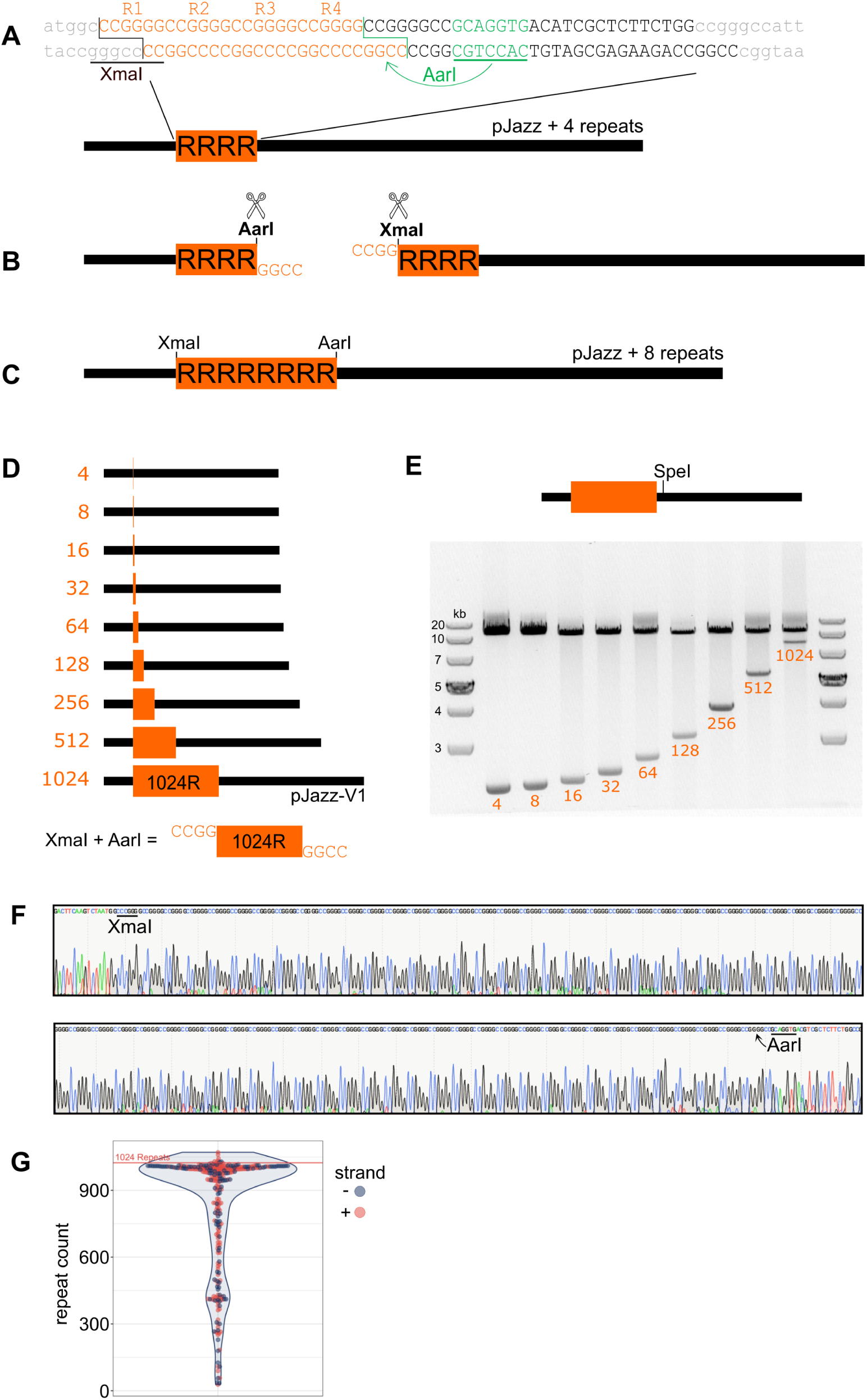
De novo generation of a large, pure, excisable repeat expansion in a linear vector. A. A pair of annealed DNA oligos (shown in uppercase) comprising four G4C2 repeats flanked by XmaI and AarI sites were designed and cloned into pJazz (flanking pJazz sequence in lowercase). B. This 4-repeat vector was digested with XmaI and AarI in two independent reactions, and each repeat harbouring fragment was isolated via gel electrophoresis (not to scale). C. Ligation of these isolated fragments doubles the repeat size of the parent vector. D. This process was repeated eight times to generate a 1024-repeat vector (pJazz-V1), which when digested with XmaI and AarI simultaneously, releases a pure G4C2 repeat with 5’ CCGG overhangs to enable subsequent cloning. E. Agarose gel electrophoresis of SpeI digestion fragments from pJazz repeat vectors at each cycle of cloning. The non-repeat harbouring 3’ fragment is 10.2 kb in each vector, while the repeat harbouring fragments rise from 2.3 kb in the 4-repeat vector, to 8.4 kb in the 1024 repeat vector. Note that DNA fragments with large repeat expansions bind with poor affinity to gel intercalating dyes. F. Sanger sequencing after 4 cycles of cloning, through the entire repeat, shows the expected uninterrupted 64 G4C2 repeat sequence. G. Summary ONT sequence data of the 1024-repeat vector showing presence of a repeat expansion in the vicinity of 1000 G4C2 repeats (median 984, 3^rd^ quartile mean 1006, maximum 1071).

We validated repeat sequence purity via Sanger sequencing in vectors up to the 64-repeat stage (**Figure 1F**), followed by Oxford Nanopore Technology (ONT) sequencing in the 1024-repeat pJazz-V1 vector (**Figure 1G**).

As proof of concept that this approach can be adapted to other repeat expansion sequences, we cloned a 10-repeat CGG cassette into pJazz, associated with Fragile X Syndrome. In this instance, we flanked repeats by AarI and BpiI, and expanded the construct up to 960 CGG repeats in length (**Supplementary Figure 1**). AarI and BpiI are both offset-cutting Type-IIS restriction enzymes, meaning different repeat units can be cloned within the cassette, expanded, and excised in a pure form.

### Engineering repeat harbouring targeting constructs

In parallel to synthetically engineering a large G4C2 repeat, we used standard recombineering technology (Copeland, Jenkins, and Court 2001; Devoy et al. 2021) to build a targeting construct for humanisation of the mouse *C9orf72* locus. Two bacterial artifical chromosome (BAC) constructs harbouring the mouse and human *C9orf72* loci (lacking a repeat), respectively, were recombined to include human *C9orf72* spanning from intron 1 to the translational stop codon; flanked by mouse homology arms at the orthologous locus (**Figure 2A**). We also included a kanamycin/neomycin selection cassette flanked by PiggyBac inverted terminal repeats, at an endogenous TTAA site within human intron 1 to enable scarless removal (Rad et al. 2010).

**Figure 2.**
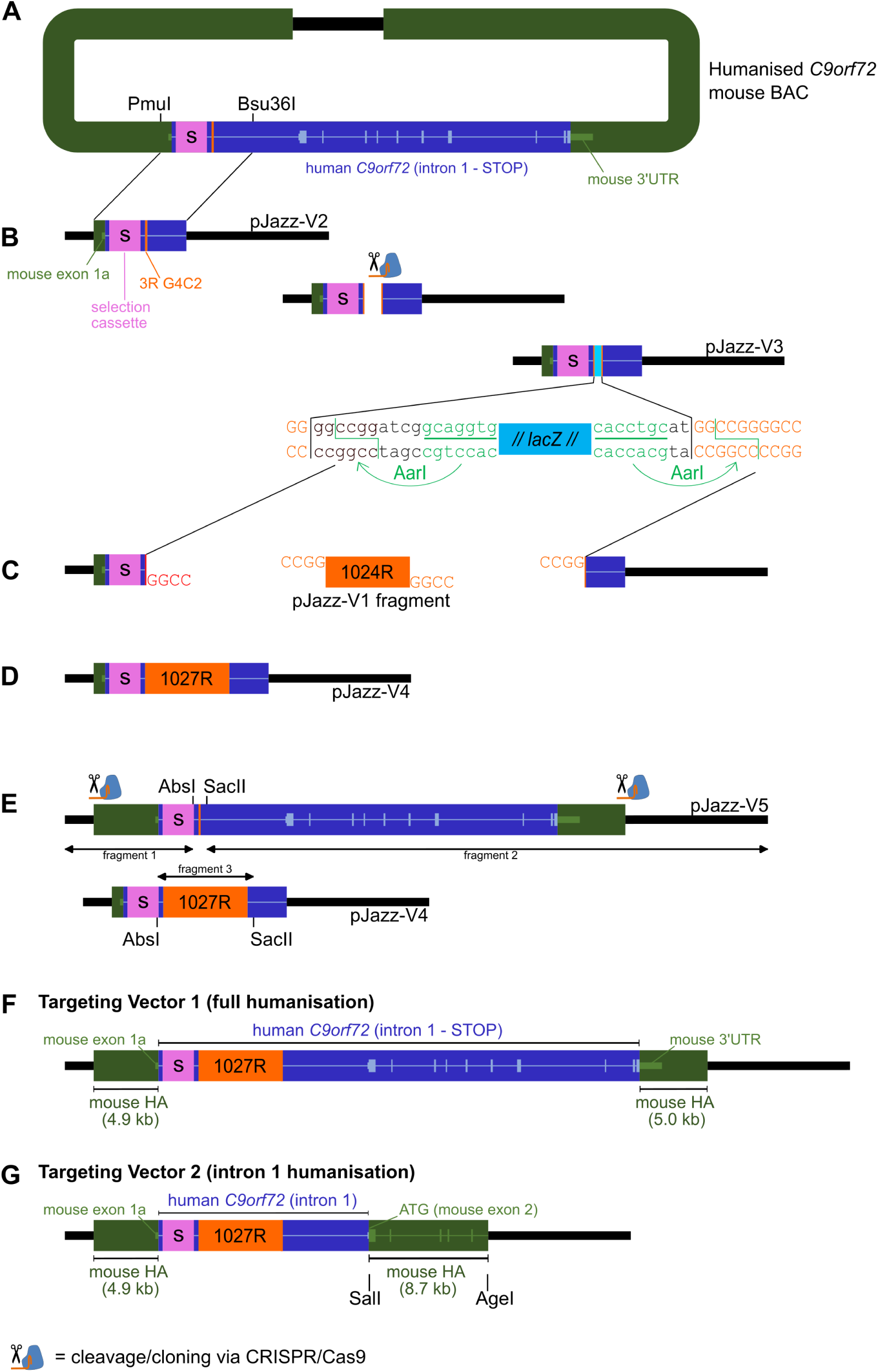
Seamless cloning of the de novo generated G4C2 repeat expansion into the human genomic sequence, within a targeting vector. A. The human *C9orf72* genomic sequence spanning from human intron 1 to the STOP codon, derived from a human BAC harbouring three G4C2 repeats, was recombineered into a mouse BAC, including recombineering of a PiggyBac-flanked neomycin selection cassette (s) 5’ to the repeat region. B. From this, a ∼6 kb fragment harbouring the 3-repeat region was cloned into pJazz using PmuI and Bsu36I restriction digestion (pJazz-V2); importantly, this fragment did not contain an AarI recognition sequence to allow the subsequent steps. pJazz-V2 was cleaved via CRISPR/Cas9 within the 3-repeat sequence and a *lacZ* landing pad was inserted via HiFi assembly to generate pJazz-V3. C. Digestion of pJazz-V3 with AarI released the landing pad, enabling the 5’ and 3’ pJazz-V3 fragments with 5’-CCGG sticky ends to be isolated and ligated together with the 1024-repeat sequence released from pJazz-V1 (Figure 1D) following AarI and XmaI digestion. D. The resulting vector, pJazz-V4, includes 1027 repeats (1024 + 3) within the native human genomic context, and flanked by proximal targeting vector elements, but lacking distal targeting vector elements. E. A pJazz targeting vector lacking the large repeat expansion was generated via CRISPR/Cas9 cleavage and blunt cloning of a fragment derived from the humanised *C9orf72* mouse BAC, including ∼5 kb mouse homology arms. An AbsI-SacII fragment from pJazz-V5 harbouring the 3-repeat region was replaced with an equivalent fragment from pJazz-V4, yielding F. Targeting vector 1, harbouring human *C9orf72* from intron 1 to the STOP codon, including 1027 repeats, and ∼5 kb mouse homology arms (HA). G. A second targeting vector was generated from targeting vector 1 designed to only humanise C9or72 intron 1 plus 1027 repeats. This was achieved by digesting targeting vector 1 with SalI, which cuts within the ATG sequence of both mouse and human *C9orf72*, and ligating to a SalI-AgeI fragment isolated from a mouse *C9orf72* BAC to generate a new 8.7 kb 3’ mouse homology arm, plus an isolated pJazz long arm backbone fragment (XmaI digested to create an AgeI compatible sticky end).

Given the stabilising attributes of the pJazz vector backbone versus a BAC vector (Nair et al. 2021), we decided to engineer the final targeting construct, including seamless integration of the large repeat, within pJazz – which required multiple steps. First, a ∼6 kb PmuI-Bsu36I restriction digest fragment (importantly lacking an AarI site) with the three-repeat G4C2 sequence at its centre, was isolated from the humanised *C9orf72* mouse BAC and cloned into a modified pJazz backbone with a mutated AarI site (GCAGGTG to GC<u>T</u>GGTG) (**Figure 2B**; pJazz-V2). pJazz-V2 was then cleaved using Cas9 and an sgRNA guide targeting the three-repeat G4C2, into which a *lacZ* ‘landing-pad’ was inserted, designed with flanking AarI sites that cut within the bisected 3-repeat G4C2 sequence (**Figure 2B**; pJazz-V3). Digestion of pJazz-V3 with AarI released the landing-pad, generating 5’ and 3’ fragments with 5’-CCGG sticky ends, which were then cloned together with the synthetic 1024-repeat XmaI-AarI fragment excised from pJazz-V1 (**Figure 2C-D**, **Figure 1D**; pJazz-V4). Next, CRISPR/Cas9 cleavage was used to isolate a larger fragment from the humanised BAC including ∼5 kb of each mouse homology arm, which was cloned into pJazz (pJazz-V5) (**Figure 2E**). An AbsI-SacII fragment from pJazz-V5 was then replaced with the equivalent AbsI-SacII fragment from pJazz-V4 harbouring the large G4C2 repeat; this constituted Targeting Vector 1 for full humanisation (**Figure 2F**). Given this construct was designed to execute a technically challenging targeting event (replacing ∼33 kb of mouse sequence with ∼25 kb of orthologous human sequence, plus a large repeat expansion), as a contingency we additionally designed a shorter construct to humanise intron 1 and 44 bp of exon 2 (up to the ATG start codon) plus the repeat expansion, which required a simple additional cloning step (**Figure 2G**; Targeting Vector 2).

Given that Targeting Construct 2 was derived from Targeting Construct 1 we performed PacBio sequencing on Targeting Construct 2, and found reads aligned correctly to the expected sequence, including through the repeat. Repeat-retracted reads were evident in a small minority, but it is likely their contribution was inflated given that coverage through the repeat region was significantly lower compared to flanking regions (**Supplementary Figure 2A**). Repeat length was independently validated by restriction digest using a panel of enzymes, yielding the expected banding patterns and a showing a repeat in the vicinity of 1000 G4C2 repeats, with no detectable evidence of retractions, again suggesting that the previously observed retracted species are in the minority (**Supplementary Figure 2B**).

### ES cell and mouse model allele QC

Both targeting constructs were used for CRISPR/Cas9 assisted gene targeting in JM8F6 C57BL/6N embryonic stem (ES) cells. DNA extracted from lipofected ES cell clones was screened by on-locus PCR, copy count qPCR of mouse and human *C9orf72* regions, and Southern blotting, to detect correctly targeted clones. Southern blotting on positive clones revealed a range of repeat sizes in individual clones up to ∼400 repeats in length, lower than the targeting vector repeat length, indicating retraction during homologous recombination (**Supplementary Figure 3**). Positive clones with the longest repeat lengths were selected for injection into mouse blastocysts.

In mice, one ES cell clone from each targeting strategy successfully passed through the germline following blastocyst injections, at which point further quality control was conducted. Targeted locus amplification (TLA; Cergentis) identified that the clone targeted for full humanisation + repeat did not fully incorporate the 3’ end of the human sequence from the targeting vector, and as such was discontinued (data not shown). TLA on the clone targeted for intron 1 humanisation + repeat revealed an intact locus as designed (**Supplementary File 1**). Primer sets specific to the targeting vector were used to capture genomic regions proximal to the insertion site in this ES cell clone. Alignment to the mouse genome revealed coverage only at the intended integration site on mouse chromosome 4 (at the *C9orf72* locus), extending beyond the mouse homology arms, and with an expected 7 kb coverage gap representing removal of mouse *C9orf72* intron 1. Alignment to the expected modified allele revealed continuous coverage indicating seamless integration of human intron 1 except for an expected coverage gap representing the G4C2 repeat sequence, which cannot be resolved with short read sequencing. No evidence for off target integrations, vector backbone integration, structural variants, or sequence variants deviating from the expected allele were found (**Supplementary File 1**).

Given that the short-read nature of the TLA method provided no information on repeat length, we next used Southern blotting. Southern blotting on XbaI digested genomic DNA from mouse spleen (from three F3 generation animals) using a (GGGGCC)_5_-DIG probe revealed a 4.5-5 kb band (**Figure 3**). Subtracting flanking non-repeat sequence (2.36 kb in length) from this figure revealed a repeat length approximately ∼2.1-2.6 kb or ∼350-440 G4C2 repeats – consistent with the repeat length observed in ES cell clone A8 from which the animals were derived (**Supplementary Figure 3**). DNA samples from animals 3.2 and 3.3 were ONT sequenced using adaptive sampling methodology. Repeat distribution analysis on reads spanning the entire repeat revealed a mean length of 373 repeats (**Figure 3C**); consistent with the repeat length estimation from the Southern blot data. Consensus sequences of repeat-flanking regions perfectly matched the expected reference sequence, and show these regions reading into the G4C2 repeat (**Figure 3D**). Heterozygous mice used in this study will be denoted as *C9orf72^h370/+^*.

**Figure 3.**
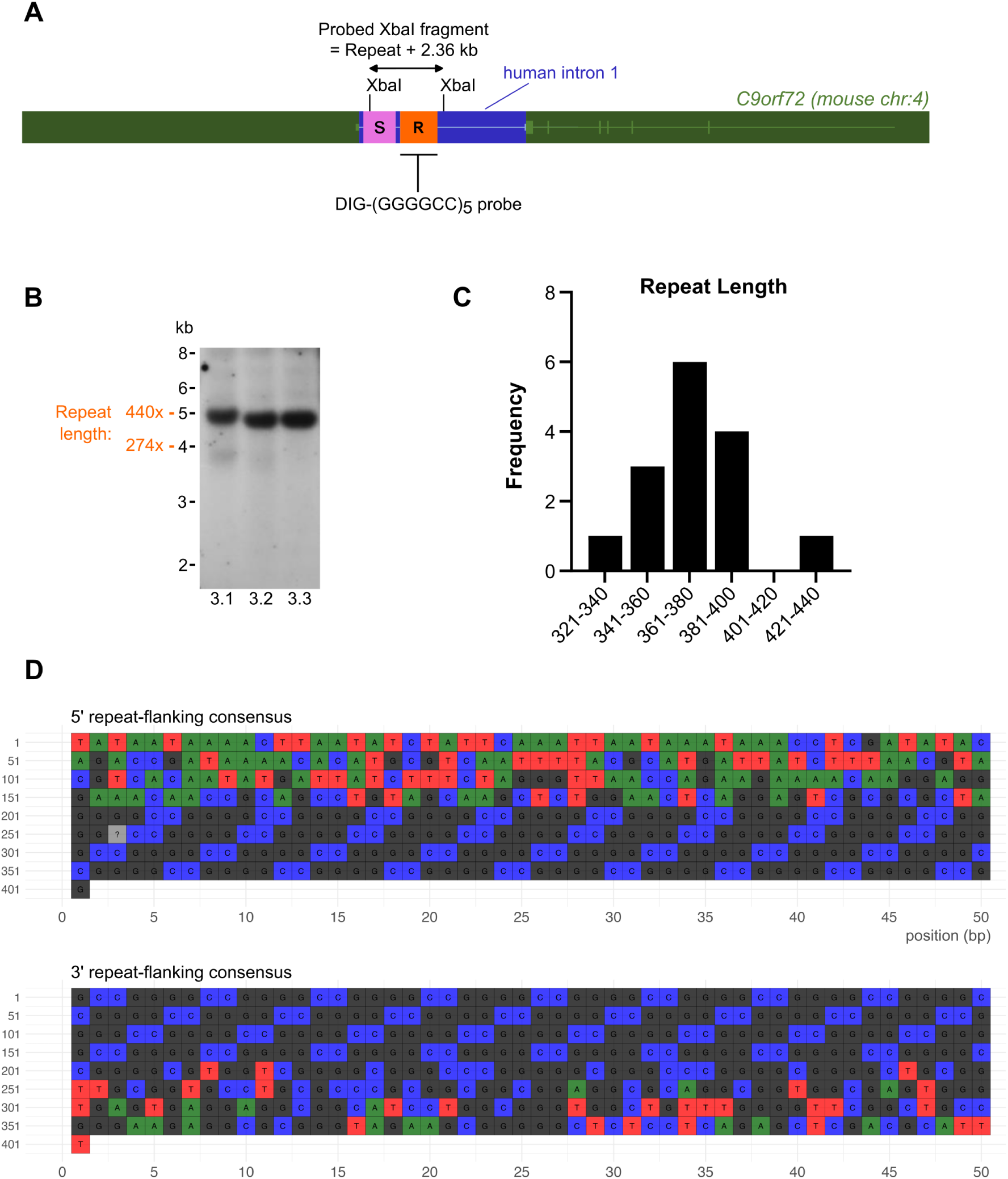
Southern blotting of the humanised intron 1 allele in *C9orf72^h370/+^* mice. A. Schematic of the gene targeted allele at the mouse *C9orf72* locus, whereby mouse intron 1 plus the 5’ UTR in exon 2 preceding the ATG start codon was replaced by orthologous human sequence, including a pathogenic G4C2 repeat expansion. Mouse sequences are in dark green, human sequences are in dark blue, while light green and light blue boxes and horizontal lines depict exons and introns of the *C9orf72* gene. Orange depicts the repeat expansion (R) and magenta depicts the selection cassette (S). B. Southern blotting on XbaI digested genomic DNA from three F3 generation mice revealed a 4.5-5kb band, suggesting the presence of ∼350-440 repeats. C. Consistent with this blot, repeat length distribution analysis from ONT sequencing of the same samples (combined sequencing data from animals 3.2 and 3.3) revealed a mean repeat length of 373 repeats. D. Consensus sequences, generated from ONT sequencing data, representing repeat flanking regions (200 bp, 5’ and 3’) show the correct endogenous human sequence reading into the G4C2 repeat. Note for the 5’ flanking region, position 1-127 represents the 3’ end of the selection cassette and position 128-200 represents human sequence reading into the G4C2 repeat.

To remove the selection cassette, we crossed the *C9orf72^h370^* line to mice expressing PiggyBac transposase (Rad et al. 2010). Sanger sequencing across the selection cassette region from DNA extracted from progeny of this cross confirmed a scarless excision event and thus a seamless replacement of mouse intron 1 with human intron 1 (**Figure 4**). Southern blotting in the same samples revealed the presence of an expected 3.1 kb band, representing stable inheritance of repeat length (370 repeats) from the previous generation (the expected lower size due to the removal of the selection cassette). In addition, a larger band equivalent to the repeat band prior to selection cassette removal was simultaneously present, indicating mosaicism in this generation with respect to selection cassette removal. This was consistent with qPCR copycount genotyping data from the same animals, whereby the internal selection cassette assay yielded reduced, but not zero, values. Genotyping of the subsequent (F5) generation confirmed germline transmission of the final selection-cassette-free allele (i.e. positive for the humanised allele and zero values for the internal selection cassette assay (**Supplementary Figure 4**).

**Figure 4.**
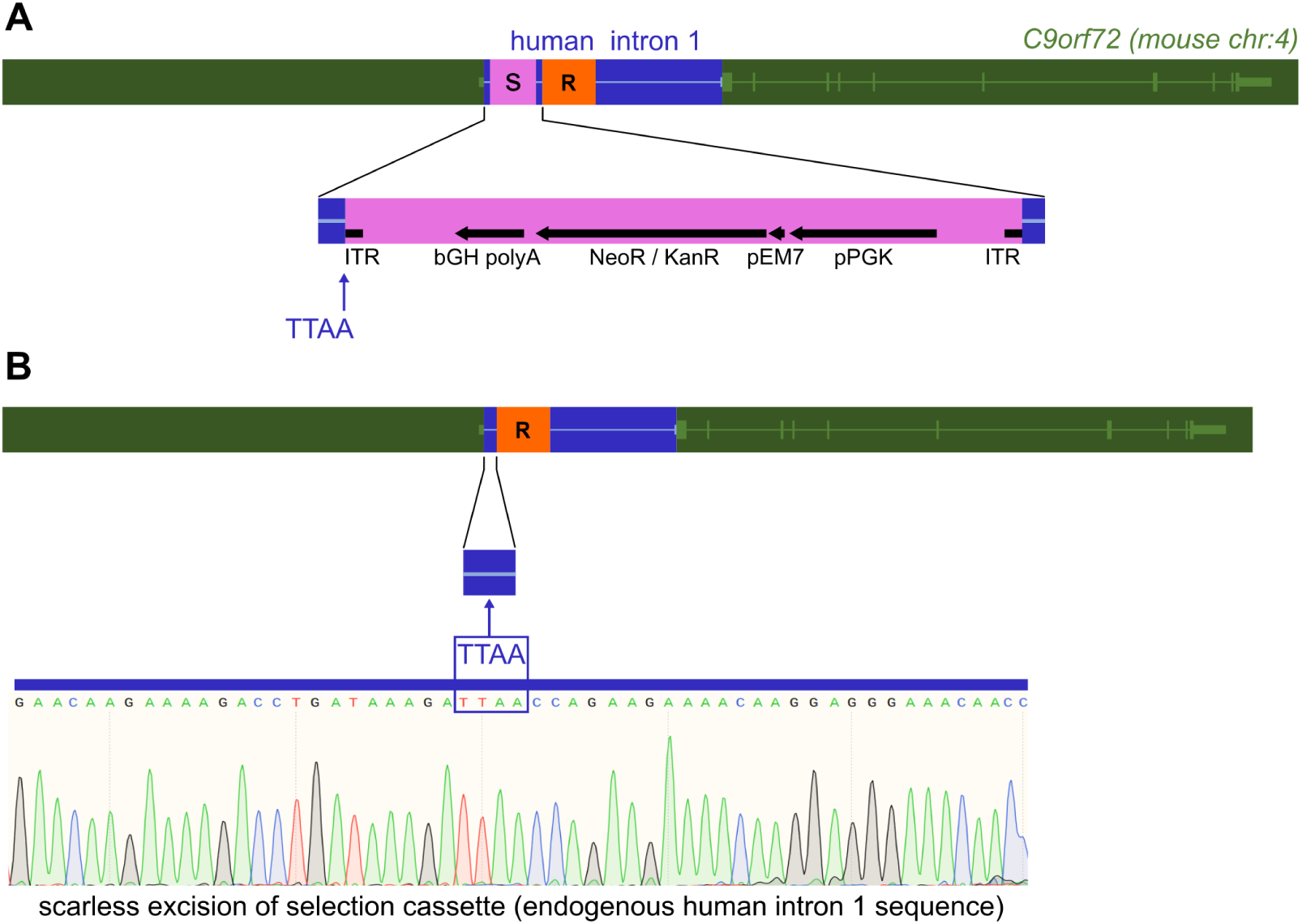
Scarless selection cassette removal from the *C9orf72^h370^* allele. A. Schematic of the gene targeted allele at the mouse *C9orf72* locus, with magnified detail of the selection cassette region (magenta); including pEM7 and pPGK promoters, dual neomycin/kanamycin antibiotic resistance gene, bGH polyA sequence, and PiggyBac inverted terminal repeats (ITR). Human sequence in blue, mouse sequence in green, G4C2 repeat in orange (R). The selection cassette was engineered at an endogenous TTAA site within human intron 1, 5’ to the repeat expansion (indicated with a blue arrow). B. Schematic of the allele following selection cassette removal. PiggyBac transposase excises regions flanked by PiggyBac ITRs, leaving behind the original TTAA insertion site. Thus, the excision was scarless, verified by Sanger sequencing on DNA samples from F4 generation mice (trace shown).

### Mutant allele expression analysis in *C9orf72^h370/+^* mice

Mouse brains were collected from F4 generation animals heterozygous for the inserted allele, for expression analyses. We first amplified a target spanning exons 5 and 6 (present in both wild type and mutant alleles), revealing that total *C9orf72* mRNA levels in heterozygous mutant mice were equivalent to wild type (**Figure 5A**). We next amplified a human *C9orf72* intron 1 target, revealing expression only in the mutant animals, as expected. At the protein level, we detected no significant difference in C9orf72 levels between genotypes (**Figure 5C, D**), consistent with total mRNA levels. Finally, to determine whether the inserted repeats underwent non-AUG RAN translation into DPRs, we utilised our established immunoassay for poly-GA (Quaegebeur et al. 2020). Poly-GA was significantly higher than controls in *C9orf72^h370/+^* mouse brains at 2.5-4 months of age, confirming that the repeats undergo RAN translation (**Figure 5E**).

**Figure 5.**
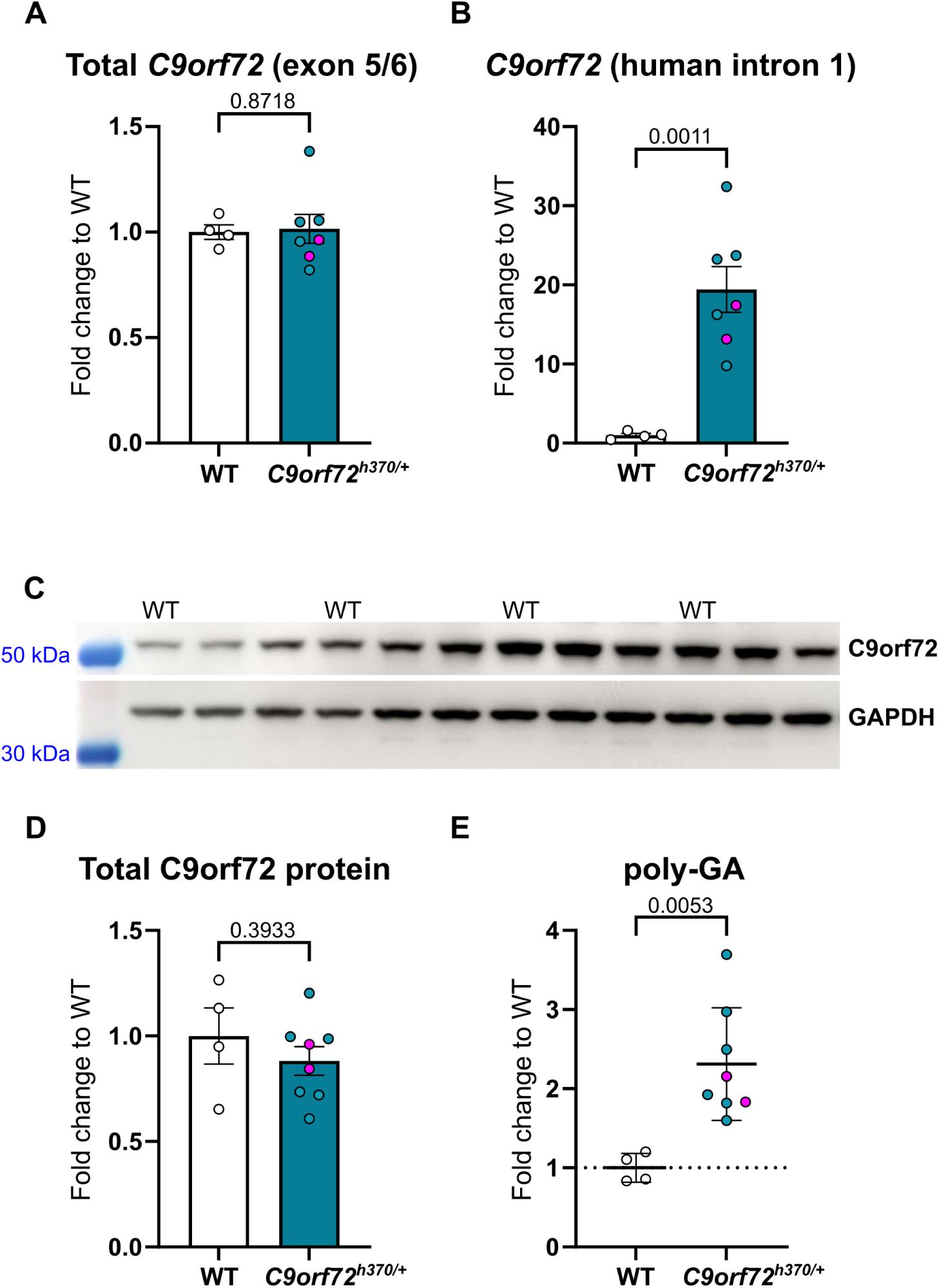
*C9orf72* locus expression analysis in mouse brain. A. qRT-PCR detection of total *C9orf72* mRNA, using primers that detect an amplicon spanning exon 5-6, present in both wild type and humanised transcripts; normalised to *Gapdh* (n=4 wild type, n=8 *C9orf72^h370/+^*). B. qRT-PCR detection of human *C9orf72* intron 1 retaining transcripts, normalised to *Gapdh* (n=4 wild type, n=7 *C9orf72^h370/+^*). C. Immunoblot detection of total C9orf72 protein normalised to GAPDH (n=4 wild type, n=8 *C9orf72^h370/+^*); wild type lanes are indicated, the remaining lanes are *C9orf72^h370^* samples. D. Quantification of C9orf72 immunoblot data, normalised to GAPDH. E. Immunoassay detection of poly-GA protein (n=4 wild type, n=8 *C9orf72^h370/+^*). Wild type control brains (WT) were from C57BL/6N mice aged 91-92 days. *C9orf72^h370^* mice were from the F4 generation, aged 77-121 days. Magenta coloured data points represent *C9orf72^h370/+^* samples from animals without any selection cassette removal. P values shown above each chart, unpaired *t*-test.

## Discussion

Here, we have devised novel methodology to synthetically engineer pathogenic repeat expansions within their endogenous genomic context for generating models of human disease. We successfully applied this method to generate the first mouse model with seamless humanisation of *C9orf72* intron 1 including ∼370 G4C2 hexanucleotide repeats, the *C9orf72^h370^* strain. Total C9orf72 protein levels were not depleted in young animals, suggesting relatively normal expression from the mutant allele. Future studies will determine whether repeat-allele mediated transcriptional silencing leading to haploinsufficiency is a function of age in this model. We were able to detect poly-GA in young *C9orf72^h370/+^* mouse brains, demonstrating RAN translation as occurs in the human condition, but again, progression with age is yet to be determined. Effects of homozygosity have not yet been tested.

Generating models of age-related neurodegeneration is challenging given that humans carrying pathogenic mutations, such as the *C9orf72* repeat expansion, typically develop symptoms after several decades of life. Overexpression models have been employed to accelerate pathological mechanisms within the short lifespan of small animal models. For example, overexpression of *C9orf72* constructs in Drosophila importantly demonstrated the neurotoxic effects of dipeptide repeat proteins (Mizielinska et al. 2014). In mice, AAV overexpression of G4C2 constructs leads to hallmark molecular changes including DPR production and TDP-43 proteinopathy alongside cognitive and motor deficits (Chew et al. 2015; Chew et al. 2019).

Nevertheless, to attempt to recreate more physiological expression and therefore understand early stage mechanisms, other strategies have been developed, such as the creation of bacterial artificial chromosome (BAC) transgenic mouse models that include proximal human gene regulatory sequences to control a pathogenic human *C9orf72* allele – although reported phenotypes from these models is variable, both between models and also between labs using the same model (O’Rourke et al. 2015; Peters et al. 2015; Jiang et al. 2016; Liu et al. 2016). This variability may in part arise from the effects of random genomic integration of the transgenic constructs including local mutation (Goodwin et al. 2019), the exact extent of human regulatory sequence included, and the mouse genetic background used (Mordes et al. 2020; Nguyen et al. 2020).

More precise approaches involve gene targeting, such as that of DPR-expressing constructs into the mouse *C9orf72* locus to enable specific investigation of pathology triggered by individual DPRs (Milioto et al. 2024). Such models are less likely to develop overt clinical disease (because they do not overexpress the sequence of interest), but are important for studying physiologically relevant early-stage processes.

A refinement of gene targeting is genomic humanisation, whereby mouse loci are replaced with the orthologous human genomic sequence, including pathogenic variants if desired (Devoy et al. 2021; Nair et al. 2019; Zhu et al. 2019; Benzow et al. 2024; Saito et al. 2019; Barendrecht et al. 2023; Saito et al. 2014; Knouff et al. 1999; Mann et al. 2004; Huynh et al. 2019; Foley et al. 2022). We have applied this latter approach here as part of wider efforts to generate humanised models for ALS/FTD research (Devoy et al. 2021).

We successfully applied novel molecular cloning methodology to construct a targeting vector enabling generation of a mouse model with seamless humanisation of *C9orf72* intron 1 including ∼370 G4C2 repeats. This represents a reduction from the ∼1000 G4C2 repeats present in the targeting construct, suggesting that recombination involving repeat sequences during gene targeting is vulnerable to retraction events, although we cannot rule out that recombination occurred with a minor retracted species within the purified targeting construct DNA preparation. A similar retraction phenomenon was reported during the creation of a gene targeted mouse model harbouring a CTG expansion allele in the *Dmpk* gene (Nutter et al. 2019). In our model, the repeat in the *C9orf72^h370^*allele appeared relatively stable in length following blastocyst injection of the ES cell clone and four generations of breeding from the F0 founder. This mouse provides an ideal platform for studying genetic or environmental factors that could influence repeat stability.

Of note, another group has in parallel independently engineered a similar *C9orf72* repeat knock-in mouse model (Kojak et al. 2024), but with some key differences: (1) The initial knock-in repeat length engineered in mice by Kojak et al was 96 G4C2 repeats, which was later found to be sensitive to expansion or retraction by inducing a double strand break in the repeat with CRISPR/Cas9, in zygotes, enabling mice with longer repeat lengths to be generated. Interestingly, repeat lengths over 400 were found to be more unstable, which is the length we are approaching in our model. (2) Our model has a seamless and scarless integration of the repeat sequence into the human genomic context, while the repeat in the Kojak et al model is flanked by exogenous sequences 5’ and 3’ to the repeat, and also includes a loxP site as a remnant of selection cassette removal. (3) Our model integrates all of human intron 1 plus 44 bp of exon 2 up to the ATG start codon; while Kojak et al introduced ∼1 kb of human intron 1 (5’ end), plus a 3’ portion of human exon 1a (**Supplementary Figure 5**). It remains to be determined if any of these differences impact repeat-length dynamics or RAN translation, but the generation of two independent *C9orf72* repeat knock-in lines and the allelic series of different repeat lengths generated by Kojak et al now provides a powerful resource for investigating these questions and downstream effects on pathological processes. Finally, at this time we have not yet generated a line of humanised mice with a non-pathogenic repeat, but endonuclease-mediated repeat-targeted cleavage, in zygotes, will provide an effective means to do so, and in turn provides a means for driving intergenerational repeat expansion.

This new gene targeted model will enable in vivo investigation of the consequences of a *C9orf72* repeat allele in its native genomic context providing a physiological platform to study, in combination, the spectrum of primary downstream molecular events, including: intergenerational and somatic instability, repeat-RNA species generation and dynamics, repeat-induced haploinsufficiency, and DPR production from sense and antisense strands. Furthermore, the *C9orf72^h370^* model provides a physiological platform for testing candidate therapies targeting the human repeat allele. While physiological neurodegeneration models are typically slower to produce clinical phenotypes, combination with additional humanised alleles, models expressing known genetic risk factors, incorporation of purported environmental risk factors, or combining with proteopathic seeding paradigms provides potential for additional model refinement. We will therefore make this model freely available to the research community to enable this work.

## Supplementary Figures

**Supplementary Figure 1.**
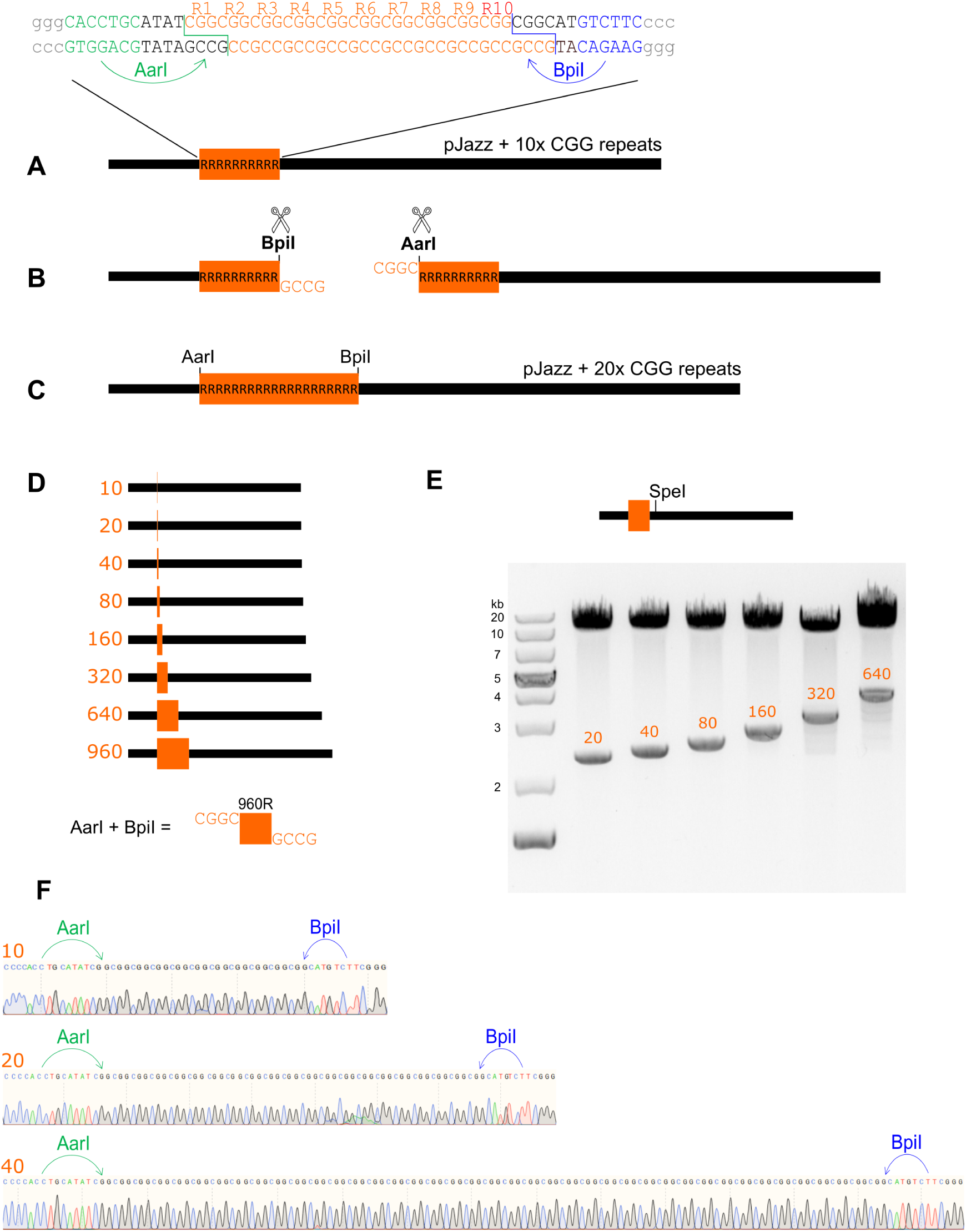
De novo generation of a large, pure, excisable CGG repeat expansion in a linear vector. A. A pair of annealed DNA oligos comprising 10 CGG repeats flanked by AarI and BpiI sites was designed and cloned into pJazz. B. This 10-repeat vector was digested with AarI and BpiI in two independent reactions, and each repeat harbouring fragment was isolated via gel electrophoresis. C. Ligation of these isolated fragments doubles the repeat size of the parent vector. D. This process was repeated six times to generate a 640-repeat CGG vector. Doubling the 640-repeat vector did not yield stable 1280-repeat clones, but combining 640- and 320-repeat constructs yielded stable 960-repeat clones. Digestion with AarI and BpiI simultaneously releases a pure CGG repeat with 5’-CGGC overhangs to enable subsequent cloning. E. Agarose gel electrophoresis of SpeI digestion fragments from pJazz repeat vectors at each cycle of cloning. The non-repeat harbouring 3’ fragment is 10.2 kb in each vector, while the repeat harbouring fragments rise from 2.3 kb in the 10-repeat vector, to 4.2 kb in the 640-repeat vector (960-repeat gel not shown). F. Sanger sequencing after 1, 2, and 3 cycles of cloning shows the expected number of uninterrupted CGG repeats.

**Supplementary Figure 2.**
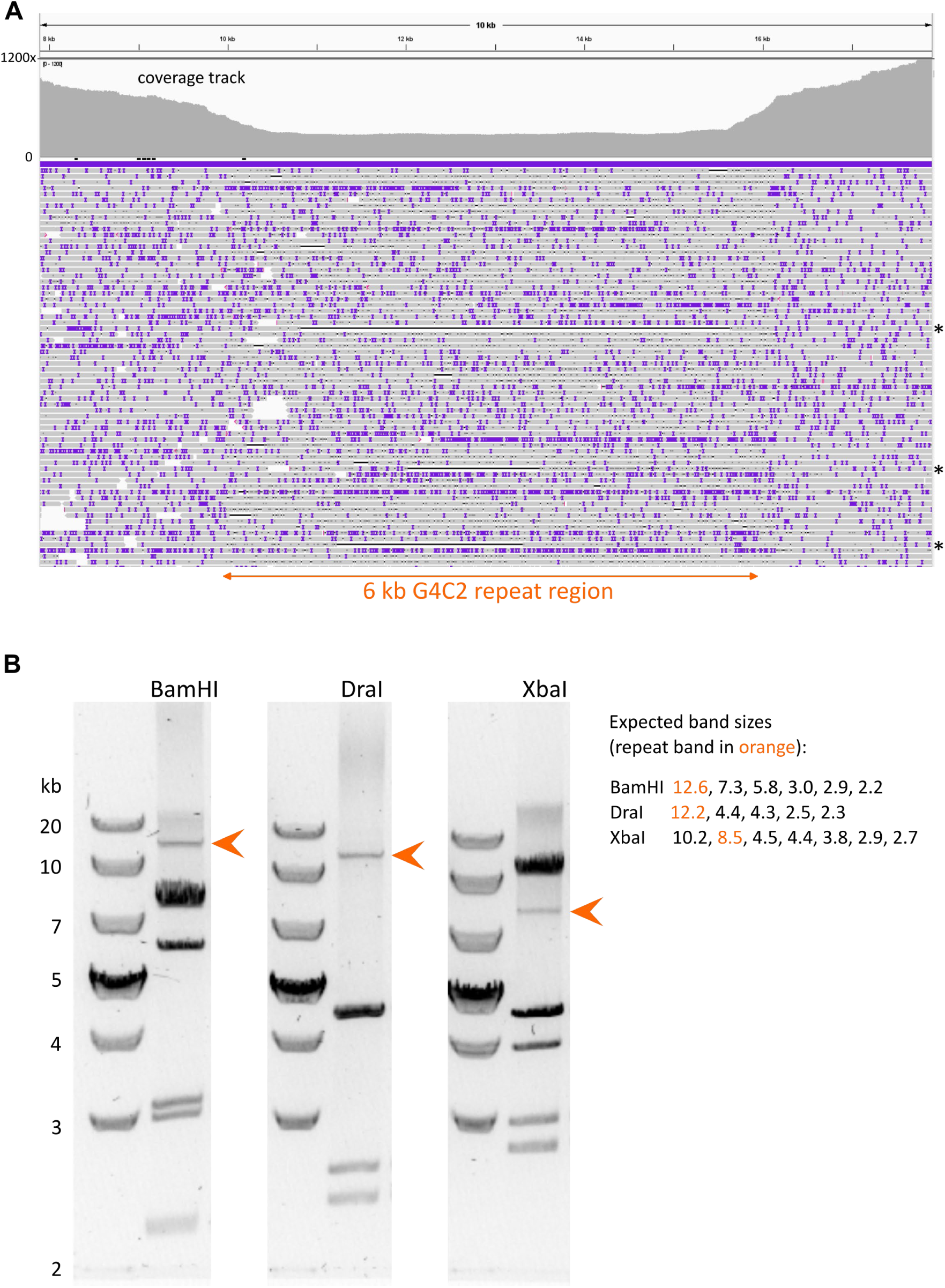
Sequence and repeat analysis of Targeting Construct 2. A. Targeting Construct 2 PacBio sequencing data. IGV visualization is shown for a 10 kb region including the 6 kb G4C2 repeat, aligned to the expected engineered sequence. The coverage profile at the top shows coverage dipping below 300x across the repeat, while climbing to over 1000x in flanking regions. Individual read tracks marked by asterisks (right hand side) highlight reads with a notably retracted G4C2 sequence (shown by horizontal black bars). B. Restriction digest fragment analysis of Targeting Construct 2 purified DNA. Left lane of each gel image was loaded with Generuler 1 kb plus DNA ladder; right lane represents restriction digest as indicated above gel. Banding patterns match those expected, including repeat harbouring bands (orange arrowhead and orange text) matching that expected for a repeat in the vicinity of 1000 G4C2 repeats. Of note, the G4C2 repeat binds with low affinity to the DNA intercalating dye (GelRed), resulting in weak fluorescence of repeat harbouring bands under UV light. No evidence of retraction was detected. Retractions would be represented by bands up to 6 kb lower in size – the DraI digest shows this most clearly as it shows no bands within 6 kb of the repeat band.

**Supplementary Figure 3.**
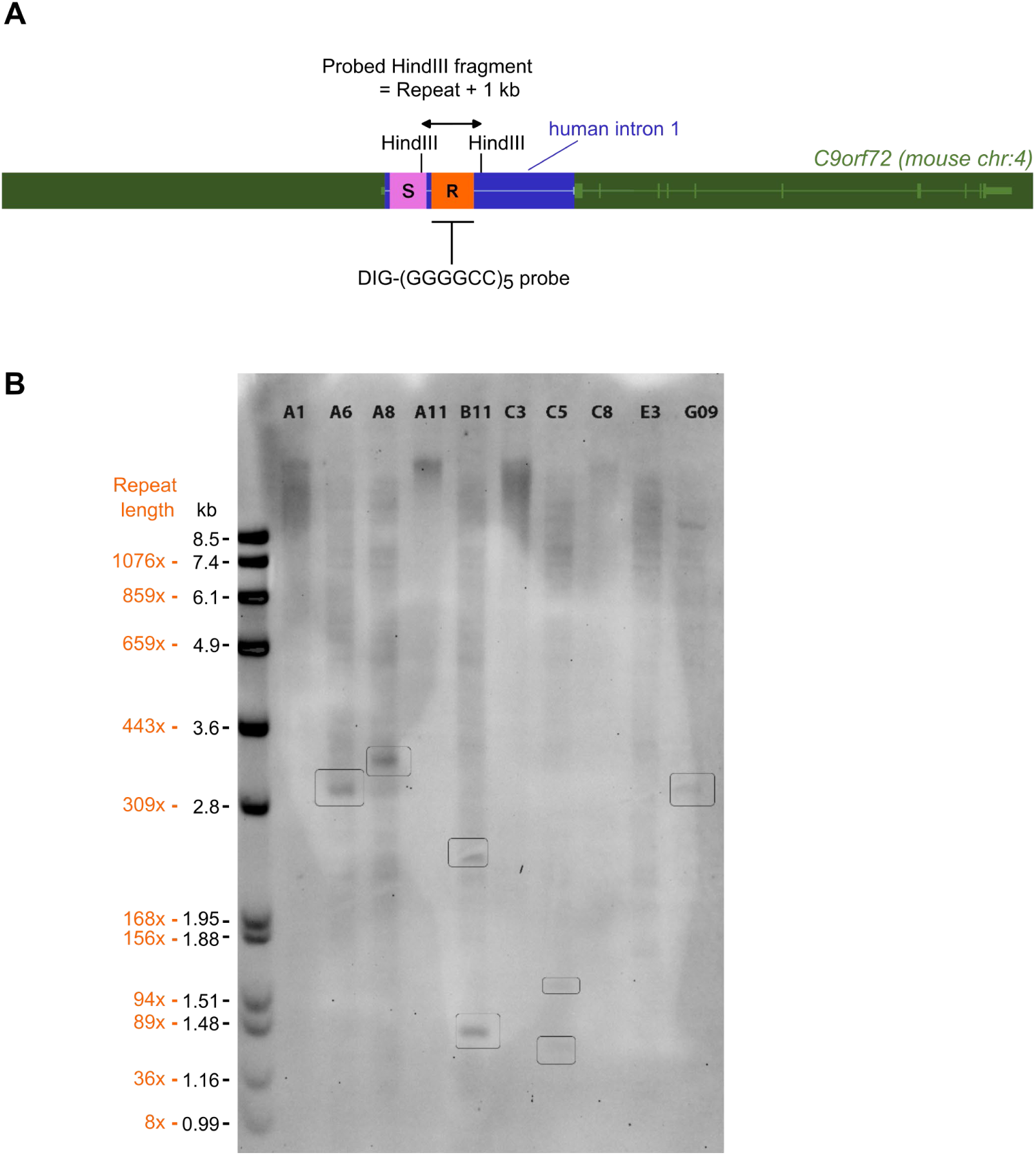
Southern blotting genomic DNA from candidate ES cell clones. A. Schematic of the gene targeted allele at the mouse *C9orf72* locus. Mouse sequences are in dark green, human sequences are in dark blue, while light green and light blue depict exons and introns of the *C9orf72* gene. Orange depicts the repeat expansion (R) and magenta depicts the selection cassette (S). B. Southern blotting on HindIII digested genomic DNA from targeted ES cell clones revealed a range of repeat lengths suggesting repeats are vulnerable to retraction events during gene targeting. The *C9orf72^h370^* line was derived from clone A8. Repeat length in clone A8 is consistent with the repeat length observed in *C9orf72^h370/+^* mice (370 repeats).

**Supplementary Figure 4.**
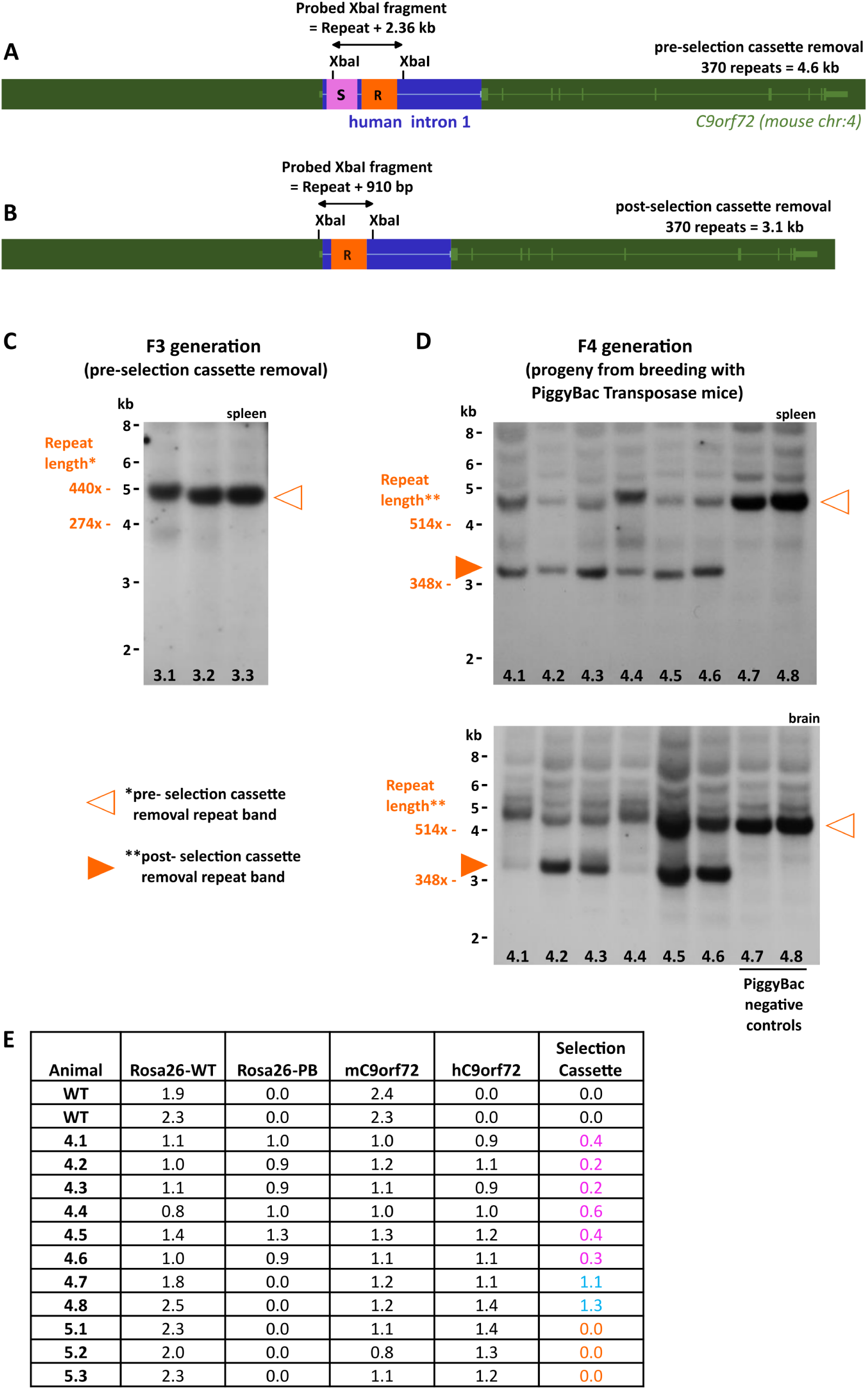
Repeat length was maintained in *C9orf72^h370/+^* mice following selection cassette removal. A. Schematic of the gene targeted *C9orf72^h370^* allele pre-selection cassette removal. B. Schematic of the gene targeted allele post-selection cassette removal. In A and B, the expected band size for 370 G4C2 repeats is indicated. Mouse sequences are in dark green, human sequences are in dark blue, while light green and light blue depict exons and introns of the *C9orf72* gene. Orange (R) depicts the repeat expansion and magenta depicts the selection cassette (S). C. Southern blotting on F3 generation mouse spleen DNA samples prior to selection cassette removal reveals a band consistent with a repeat length of 370 G4C2 repeats (orange outlined arrowhead). D. Southern blotting on F4 generation mouse DNA spleen and brain samples (from the progeny of breeding to PiggyBac transposase expressing mice) reveals an expected band at 3.1 kb, representing a 370 G4C2 repeat allele minus the selection cassette (orange arrowhead), together with the pre-selection cassette removal band (orange outlined arrowhead), indicating mosaicism. F4 generation controls that did not inherit the PiggyBac transposase allele only show the pre-selection cassette removal band (animals 4.7 and 4.8). E. qPCR genotyping results show copycount assay values for presence of the PiggyBAC transposase allele (Rosa26-WT), the wild type Rosa26 allele (Rosa26-PB), the wild type mouse *C9orf72* allele (mC9orf72), the humanised allele (hC9orf72), and for the presence of the selection cassette. The selection cassette assay does not drop to zero in F4 generation animals that inherited the PiggyBAC transposase allele (magenta text), consistent with mosaicism observed in Southern blot data. F4 generation animals that did not inherit the PiggyBAC transposase allele maintained a value of 1 for the selection cassette assay (blue). In the subsequent F5 generation, the humanised allele minus the selection cassette successfully transmitted through the germline as shown by selection cassette assay values of zero (orange) (data shown from males from which sperm was cryopreserved).

**Supplementary Figure 5.**
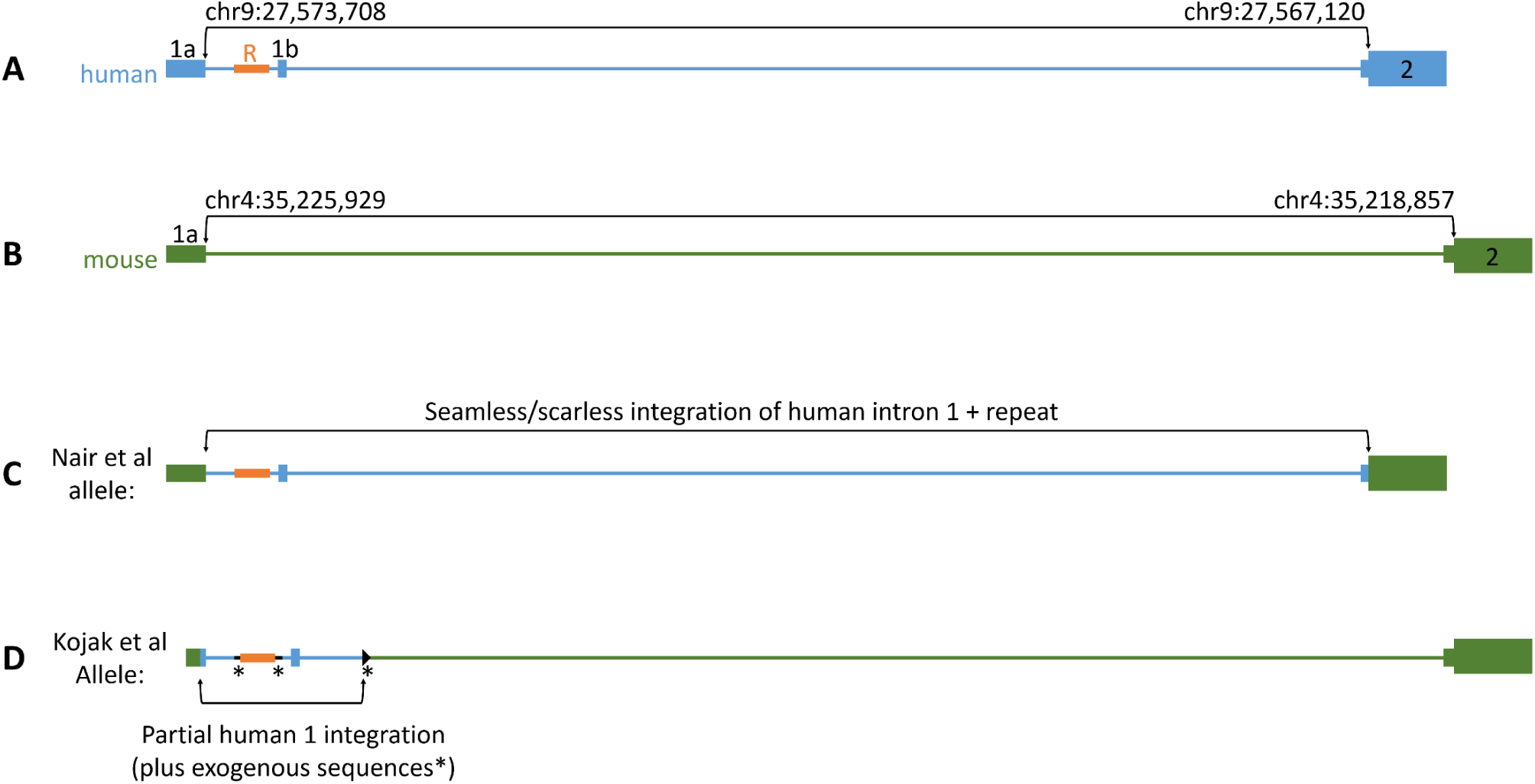
Schematic comparison of the *C9orf72^h370^* allele versus the Kojak et al *C9orf72* allele. A. Schematic of the human *C9orf72* locus. Exons 1a, 1b, and 2 are numbered, the repeat (R) is represented in orange. The black arrowed bracket (above) represents the human region we integrated into the mouse *C9orf72* locus on chromosome 4 to generate the *C9orf72^h370^* humanised allele. B. Schematic of the mouse *C9orf72* locus. The black arrowed bracket (above) represents the mouse region we replaced with the human region depicted in A. C. Schematic of the *C9orf72^h370^* humanised allele combining parts indicated in A and B. D. Schematic of the Kojak et al allele. Green represents mouse regions, blue represents human, orange represents the G4C2 repeat, boxes represent exons, lines represent introns, exogenous sequences are marked with *, the black triangle represents a remnant loxP site.

## Methods

### Molecular Cloning

pJazz vector sequences, commercially sourced inserts, vector derived fragments, and annealed oligo pairs used in pJazz cloning steps can be found in **Supplementary File 2.**

Two versions of pJazz (Lucigen) were used: pJazz-OK (kanamycin resistance) and pJazz-OC (chloramphenicol resistance). For repeat building synthetic G4C2 constructs (to create pJazz-V1), a pair of annealed DNA oligonucleotides (IDT) was designed with 5’-CCGG overhangs to clone into XmaI-predigested pJazz-OK. For repeat building synthetic CGG constructs, a pair of annealed DNA oligonucleotides (IDT) was designed to blunt clone into SmaI-predigested pJazz-OK-deltaAarI. pJazz-OK-deltaAarI was generated by replacing a PpuMI-SacI fragment from the long arm of the backbone with an equivalent fragment obtained commercially (IDT gBlock) harbouring an AarI mutation (GCAGGTG to GC<u>T</u>GGTG). pJazzV2-V5 and the final Targeting Constructs were cloned using pJazz-OC.

Vector fragments and inserts were isolated via gel extraction and purified, or synthesised commercially, prior to ligation steps. Following restriction digestion or Cas9 digestion, vector fragments were electrophoresed on agarose gels using GelGreen DNA dye (Biotium), cut from gels under blue light, and purified using NucleoSpin Gel and PCR Clean-up kit (Machery-Nagel) or Zymoclean Large Fragment DNA Recovery kit.

Ligation steps in pJazz were performed according to the BigEasy v2.0 Linear Cloning Kit instructions (Lucigen), in 10 μl reactions using CloneSmart DNA Ligase, and electroporating using BigEasy-TSA Electrocompetent Cells (provided in the kit) and E. coli Pulser Transformation Apparatus (Biorad) at 1.8 kV, using 0.1 cm gap Gene Pulser/MicroPulser Cuvettes (Biorad). Transformed cells were recovered in 975 μl recovery medium for 2 hours at room temperature (RT), 150 rpm, and plated on low salt LB-agar plates.

Antibiotic concentrations used were: 30 μg/ml kanamycin (pJazz-OK) or 12.5 μg/ml chloramphenicol (pJazz-OC). For blue/white screening in pJazz, 20 μg/ml X-gal and 1 mM IPTG were added to plates. CRISPR/Cas9 digestions were performed as described previously (Nair et al. 2021). sgRNA target sites for Cas9 digestion shown in **Figure 2** were AGGAGTCGCGCGCTAGGGGC for bisection of the non-pathogenic repeat (**Figure 2B**); GAGGCTTTGAAGATATAGCA and TGGTCCATAGGCCAGTCTAG for excising a Targeting Construct precursor sequence from the humanised mouse BAC (**Figure 2E**).

The cloning strategy for engineering the Targeting Constructs in pJazz was as described in the text, but for simplicity the following iterative steps were not mentioned in the main text. Generation of the pJazz-V2 PmlI-Bsu36I fragment (**Figure 2A**) required two steps. First, a 6.15 kb PmuI fragment was blunt cloned from the humanised *C9orf72* mouse BAC into pJazz-OC. Next, the pJazz long arm side of this vector was trimmed via Bsu36I digestion (to remove a distal AarI site), and a new pJazz long arm (deltaAarI) fragment (via XmaI digestion) was added via a pair of annealed oligos acting as sticky-ended bridge.

Adding the repeat from pJazz-V4 into pJazz-V5 (**Figure 2E**) also required several iterative steps due to the inefficiency in cloning very long fragments together with a long repeat expansion in a single step. Beginning with adding the full 3’ end of the targeting construct to pJazz-V4, pJazz-V5 was digested with SacII, and the long arm end (40.5 kb) was gel isolated. pJazz-V4 was then digested via CRISPR/Cas9 using an sgRNA guide targeting a site downstream of the SacII restriction site (CCAGGCTTTGTATTCAGAAA), and the previously used *lacZ* landing pad was blunt cloned into this site creating pJazz-V4(*lacZ*.1) (to avoid having to gel isolate the repeat expansion fragment, which proved prohibitive due to the similar size of the remaining SacII fragment). pJazz-V4(*lacZ*.1) was digested with SacII and the resulting fragments were ligated to the 40.5 kb fragment from pJazz-V5. Selecting white colonies using blue/white screening was successful in identifying a desired clone with the full 3’ end of the targeting construct together with the repeat expansion (pJazz-V5a). To add the 5’ end of the targeting construct a similar process was followed. First, the *lacZ* landing pad was cloned into the short arm backbone of pJazz-V4 via CRISPR/Cas9 digestion and blunt-end cloning, using a pair of sgRNA guide targets (AATGAAATAAGATCACTACCGGG and TACTGCGATGAGTGGCAGGGCGG) targeting the chloramphenicol cassette, creating pJazz-V4(*lacZ*.2), which was subsequently digested with AbsI and the short arm side (7.2 kb) was gel isolated and purified. In parallel, pJazz-V5a was also digested with AbsI and the resultant fragments were ligated to the 7.2 kb fragment from pJazz-V4(*lacZ*.2). Selecting blue colonies using blue/white screening was successful in identifying a desired clone of Targeting Vector 1 (antibiotic selection here relied on the dual neomycin/kanamycin selection cassette).

The recombineering workflow, including sequence detail, to produce the humanised *C9orf72* BAC is provided in **Supplementary File 3**. Standard recombineering protocols were followed using DH10B electrocompetent E. coli and the Red/ET plasmid system (Gene Bridges) (Hofemeister et al. 2011).

### ES cell targeting

CRISPR/Cas9 site-specific nucleases were designed against the extremities of exchange event at the *C9orf72* locus using the CRISPOR algorithm. These target sites were cloned as synthetic linkers into the BbsI restriction site of pX330 (Addgene Plasmid #42230), modified with the addition of a puromycin resistant cassette. 1x10^6^ mouse embryonic stem cells (JM8F6) were lipofected with 1.25 μg of Targeting Construct 1 or 2, together with 625 ng of the 5’ and the 3’ CRISPR/Cas9 plasmids using Lipofectamine LTX (Thermofisher). 600 ng/ml puromycin was applied 24 hours after lipofection for a period of 48 hours, followed by selection in 210 μg/ml G418. Individual G418 resistant clones were propagated on 96 well-plates and were genotyped by long-range PCR over the 5’ homology arm, using a forward primer binding upstream of the 5’ homology arm and a reverse primer within the humanized intron to detect homologously recombined ES cell clones. Additional on-locus PCR was performed to assess for the presence of human intron 1 sequence 3’ to the repeat, and the 3’ human-mouse junction. Additionally, probe-based qPCR assays were performed to assess copy number with respect to the inserted allele and the wild type mouse *C9orf72* allele. The sequences of the CRISPR/Cas9 target sites, the cloning oligonucleotides and the qPCR assays are shown in **Supplementary File 4.**

Correctly targeted ES cell clones were microinjected into albino C57BL/6J (Jax stock 000058) blastocysts and chimeras showing ES cell contribution by coat colour were generated. The chimeras were mated with albino C57BL/6J females and germline transmission of the humanized allele was confirmed in the resulting F1 generation mice by the 5’ screening PCR.

### Mice

Mice were maintained and studied according to UK Home Office legislation at MRC Harwell with local ethical approval (MRC Harwell ethical committee). Mice were fed ad libitum (Rat and Mouse Breeding 3 (RM3), Special Diet Services) with free access to water (chlorinated to 9-13 ppm). Mice were kept under constant conditions with a regular 12-hour light (19:00-07:00) and dark (07:00-19:00) cycle, temperature of 21±2°C and humidity 55±10%. Mice were housed in same sex cages of up to five mice. Cages contained Aspen Chips bedding (Datesand), shredded paper and a rodent tunnel for enrichment. F1 generation *C9orf72^h370/+^* animals were crossed for two generations to C57BL/6N wild type mice. F3 generation animals were used for initial Southern blotting and ONT sequencing. F3 generation animals were crossed to transgenic mice expressing PiggyBac transposase (gene targeted at the *Rosa26* locus) (Rad et al. 2010), bred on a C57BL6/N background, to remove the selection cassette. Samples from the resulting F4 generation were used for further Southern blotting. F4 generation animals were crossed one more generation with C57BL6/N wild type mice to breed out the PiggyBAC transposase allele, before cryopreservation of sperm. qPCR probe assays were used to genotype the presence or absence of the humanised allele or the wild type mouse *C9orf72* allele; plus assays were designed to assay for presence or absence of the PiggyBAC transposase allele, and for the presence or absence of the selection cassette sequence. The selection cassette region was also amplified and Sanger sequenced to confirm scarless excision in F4 generation mice. F4 generation mice were found to be mosaic for selection cassette removal. One further cross to C57BL/6N wild type mice enabled selection of animals positive for the humanised allele, negative for the selection cassette, and negative for the PiggyBAC expressing Rosa26 allele, for sperm cryopreservation. Assay and primer details can be found in **Supplementary File 4.**

### Southern Blotting

Southern blotting on DNA from mouse ES cell clones was performed as follows, adapted from a previously described protocol (DeJesus-Hernandez et al. 2011). 10 μg genomic DNA of each sample was digested with HindIII at 37°C overnight with 30 units of enzyme. After digestion, an aliquot of each digest was visualized on an agarose gel to ensure genomic DNAs were digested to completion. Digests were electrophoresed on a 0.8% agarose gel at 60V for 16 hours. Gels were then treated with denaturation solution (Merck, N1531-1L), and then a neutralisation solution (Merck, N6019-1L), followed by overnight transfer to a nylon membrane by capillary blotting (Roche 11417240001), and cross-linked by UV. The membrane was treated with prehybridization solution (Roche 11603558001) containing 100 μg/ml salmon sperm DNA (Thermo 15632011) at 48°C for 3 hours. Next, the membrane was hybridized with DIG Easy hybridization solution containing DIG-(GGGGCC)_5_ probe (IDT; 2 ng/ml) at 48°C for 16 hours. Membranes were washed in 2x SSC and 5% SDS for 15 minutes while the hybridization oven ramped from 48°C to 65°C. Next the membrane was further washed in 2x SSC and 0.5% SDS, 0.5x SSC and 0.5% SDS, and 0.15% SSC and 0.5% SDS, for 15 minutes each at 65°C. Antibody detection was performed according to the DIG Application Manual for non-radioactive *in situ* hybridisation (Roche), using the DIG Wash and Block Buffer Set (Roche 11585762001), Anti-DIG-AP Fab fragments (Roche 11093274910) at 1:10000 dilution, and CDP-star ready to use (Roche 12041677001). Signals were visualised using ChemiDoc XRS+ system (BioRad).

Southern blotting of mouse DNA samples was performed commercially (Celplor). Approximately 2 μg genomic DNA of each sample was digested with XbaI restriction enzyme at 37°C for 24 hours with 10 units of enzyme. After digestion, an aliquot of each digest was visualized on an agarose gel to ensure genomic DNAs were digested to completion. A DIG-(GGGGCC)_5_ probe was used for repeat fragment detection. Southern blotting and signal recording were performed following Celplor LLC standard operating procedures.

### Sequencing

#### ONT sequencing on pJazz-V1

Approximately 375ng of pJazz-V1 was digested using PacI and PmeI (NEB). A 1:1 Agencourt XP bead clean-up was performed to purify digested DNA. 200ng of DNA was recovered and used to construct a ONT sequencing library with the Ligation Sequencing kit (LSK109). Library was loaded onto a flongle (R9.4) and sequenced for 4.5 hours. Resulting data was base called with High Accuracy MinKNOW (v19.12.6) (guppy v3.2.10). Mapping to the vector reference was via minimap2 (v2.10-r761). Repeat counting was via STRique (flanking cut-off of 4).

#### PacBio sequencing on Targeting Construct 2

With CD Genomics. 5μg DNA was fragmented using Covaris g-TUBE devices and subsequently repaired by treating the sample with a DNA damage repair mix. Following DNA-damage repair, blunt ends are created on each end. Then, hairpin adapters incorporating a unique barcode were ligated to each blunt end. The SMRTbell DNA template libraries with a fragment size >10 kb were selected using a Bluepippin system. Library quality was analyzed by Qubit and real-time PCR System, and average fragment size was estimated using an Agilent 2100 Bioanalyzer. Sequencing was performed on the PacBio Sequel II platform. Alignment of reads to the Targeting Construct 2 reference sequence was performed using minimap2.

#### ONT adaptive sampling on mouse genomic DNA samples

High molecular weight DNA was extracted from tissues using a Monarch HMW DNA Extraction Kit for Tissue (NEB). A protocol for sample preparation and ONT processing was developed by the Genomics England Scientific R&D Team with some minor modifications. 6-6.4 µg of DNA was fragmented to 10-15 kb using Covaris g-TUBEs (520079, Covaris). Samples were centrifuged using an Eppendorf 5424 (022620401, Eppendorf) centrifuge for 1 minute at 4500-4800 rpm, rotated 180 degrees and centrifuged for a further minute at 4500-4800 rpm. Fragments under 5 kb were reduced using the SRE XS kit (SKU 102-208-20,0 PacBio). An equal volume of Buffer SRE-XS was added to the fragmented DNA sample and centrifuged at 10,000 *g* for 30 minutes before discarding the supernatant. The DNA pellet was washed with 200 µl of 70% ethanol for a total of two ethanol washes. The remaining ethanol was evaporated off at 37°C for 2-15 minutes as required. 50 µl of PacBio Buffer EB was used to resuspend the pellet at 37°C for 10-20 minutes then 4°C overnight. For library preparation 1-1.5 µg of sample in 48 µl of nuclease-free water (NFW) was mixed with 3.5 µl NEBNext FFPE DNA Repair Buffer, 3.5 µl Ultra II End-Prep Reaction Buffer, 2 µl NEBNext FFPE DNA Repair Mix (M6630, NEB) and 3 µl Ultra II End-Prep Enzyme Mix (E7546, NEB) and incubated at 20°C for 10 minutes followed by 65°C for 10 minutes. The reaction was then incubated at room temperature for 10 minutes on a hula mixer with 60 µl of AMPure XP Beads (A63881, Beckman Coulter). The beads were pelleted on a magnet and washed with 200 µl of 70% ethanol twice before eluting in 64 µl NFW at room temperature for 10-30 minutes. The library was processed using ONT Ligation Sequencing Kit V14 (SQK-LSK114). 62 µl of eluted DNA was added to a mix of 25 µL ONT Ligation Buffer, 8 µl NEBNext Quick T4 DNA Ligase (E6056, NEB) and 5 µl ONT Ligation Adapter and incubated at room temperature for 30 minutes. 40 µl of AMPure XP beads were added and incubated on a hula mixer at room temperature for 10 minutes then pelleted on a magnet. The beads were washed twice with 250 µl ONT Long Fragment Buffer then eluted in 34 µl ONT Elution Buffer at 37°C for 15 minutes. 10-15 fmol of library was loaded onto a single PromethION Flow Cell (R10.4.1) following manufacturer’s instructions. Adaptive sampling was carried out as described below. The run lasted 72 hours with two 1-hour nuclease flushes between 20-24 hours and 44-48 hours using the ONT Flow Cell Wash Kit (EXP-WSH004) following manufacturer’s instructions. The library was stored at 4°C during the run and 10-15 fmol of library was loaded after each nuclease flush. For adaptive sampling and analysis, all code can be found at: https://github.com/rainwala/GIU-rep_exp_blast_finder. Adaptive Sampling was carried out using the Adaptive Sampling option on MinKnow version 24.02.10. The reference genome used was GRCh38.p14 (https://www.ncbi.nlm.nih.gov/datasets/genome/GCF_000001405.40/) and the co-ordinates were specified in bed format as follows: chr9 27543537 27603536 C9orf72_expansion_region. This corresponds to the file C9orf72_adaptive.bed in the Github repository above.

A custom reference sequence was constructed based on the Ensembl human *C9orf72* reference (ENSG00000147894), incorporating the two regions flanking the hexanucleotide repeat. Reads were aligned to this custom reference using minimap2 with the ‘ont’ presets. Alignment SAM files were converted to sorted and indexed BAM files using samtools. Alignment quality was assessed in R by evaluating mapping quality, analysing the CIGAR strings, and calculating alignment length, mismatch, and indel counts. High-confidence alignments were selected based on mapping quality scores and consistent directionality. Antisense strands were reoriented to the sense strand to maintain consistency. Following alignment, custom R functions were used to detect hexanucleotide repeat motifs. Gaps between repeat units were allowed to account for sequence variants and sequencing artifacts. The repeat length was determined as the longest continuous stretch of repeat units, requiring at least a perfect repeat at both the start and end of the tract. This repeat length was then divided by the hexanucleotide motif length (6 bp) to calculate the total number of repeats.

The consensus sequence of aligned reads was generated by performing multiple sequence alignment (MSA) using a custom R function that incorporates the msa package with the ClustalW algorithm. Pairwise alignment scores were calculated for each sequence relative to the consensus sequence, and outliers were identified based on standard deviation from the mean alignment score. The final consensus sequences were generated for the full repeat region, as well as for 400 base pair windows centred on the left and right flanks of the repeat. Wrapped consensus sequence plots were created using ggplot2 in R.

The bioinformatics pipeline is available at https://github.com/mike-flower/othello.

### Immunoblotting and MSD immunoassays

Protein was extracted from mouse brain by lysing cortex tissue in 10% w/v lysis buffer (RIPA buffer + 2 % SDS + protease inhibitors) for 10 minutes on ice, before homogenising at 6,500 rpm for 45 sec with a Precellys Evolution tissue homogenizer (Bertin Instruments) with 1.4 mm zirconium beads. Samples were centrifuged at 17,000 *g* at 4°C for 20 minutes and the supernatant collected and stored at -80°C. Protein concentration was determined with a BCA protein assay kit (ThermoFisher) as per manufacturer’s instructions.

Protein extracts (10 µg) were heated at 95°C in 6X Laemmli SDS sample buffer (Alfa Aesar) and separated on NuPAGE™ 4 to 12% bis-tris gels (Invitrogen). Proteins were electrotransferred to nitrocellulose membranes (Bio-Rad Laboratories). Membranes were blocked in 10% milk in PBS-T (PBS, 0.1% Tween-20) for 1 hour at room temperature. The membranes were incubated overnight at 4 °C with anti-C9orf72 antibody (GTX634482; 1:1000 dilution). After 3 washes in PBS-T, membranes were incubated with secondary HRP-conjugated antibody (1:5000 dilution) for 1 hour at room temperature. Signals were visualized by chemiluminescence (Amersham imager 680, GE lifesciences) and quantifications were performed using ImageJ software. GAPDH (14C10; #2118, Cell Signaling; 1:5000 dilution) was used as a housekeeping control for normalisation.

Meso Scale Discovery (MSD) immunoassay was performed in singleplex using 96-well SECTOR plates (MSD, Rockville, Maryland) to quantify poly(GA) expression levels. The assay was performed as previously described (Simone et al. 2018). DPR protein levels were measured using lysates prepared in RIPA buffer containing 2% SDS and 180 µL of protein was loaded per well in duplicate. Unlabelled anti-poly(GA) antibody (Sigma Aldrich, #MABN889, 2 µg/ml) was used as capture, and a biotinylated anti-poly(GA) (GA 5F2, kindly gifted by Prof Dieter Edbauer, 2 µg/ml) was used as detector.

### qRT-PCR

Total RNA was extracted from frozen mouse brain tissue using the Direct-zol RNA Miniprep Plus Kit (Zymo). RNA samples underwent reverse transcription using SuperScriptTM IV VILOTM Master Mix (ThermoFisher) with random hexamers to produce cDNA according to manufacturer’s instructions. qPCR analysis was performed using SYBR Green Master Mix (ThermoFisher) (primer sequences provided in **Supplementary File 4**) and measured with a LightCyclerTM 480 (Roche). qPCR was performed in technical triplicate with data normalised to *Gapdh* expression levels.

## Acknowledgements

EMCF, TJC, RRN, CT, RM were funded by the UK Medical Research Council (MRC) (MC_EX_MR/N501931/1). DT was funded by an MRC PhD studentship. AMI, MC, AC were supported by UK Dementia Research Institute, through UK DRI Ltd, principally funded by the MRC, grant UKDRI-1203

We thank all staff involved at the Mary Lyon Centre, MRC Harwell Institute, for animal husbandry, colony management, tissue sampling, and cryopreservation activities.

## References

Balendra, R., and A. M. Isaacs. 2018. ‘C9orf72-mediated ALS and FTD: multiple pathways to disease’, Nat Rev Neurol, 14: 544–58.

Barendrecht, S., A. Schreurs, S. Geissler, V. Sabanov, V. Ilse, V. Rieckmann, R. Eichentopf, A. Kunemund, B. Hietel, S. Wussow, K. Hoffmann, K. Korber-Ferl, R. Pandey, G. W. Carter, H. U. Demuth, M. Holzer, S. Rossner, S. Schilling, C. Preuss, D. Balschun, and H. Cynis. 2023. ‘A novel human tau knock-in mouse model reveals interaction of Abeta and human tau under progressing cerebral amyloidosis in 5xFAD mice’, Alzheimers Res Ther, 15: 16.

Benzow, K., K. Karanjeet, A. L. Oblak, G. W. Carter, M. Sasner, and M. D. Koob. 2024. ’Gene replacement-Alzheimer’s disease (GR-AD): Modeling the genetics of human dementias in mice’, Alzheimers Dement, 20: 3080–87.

Braems, E., B. Swinnen, and L. Van Den Bosch. 2020. ‘C9orf72 loss-of-function: a trivial, stand-alone or additive mechanism in C9 ALS/FTD?’, Acta Neuropathol, 140: 625–43.

Chew, Jeannie, Casey Cook, Tania F. Gendron, Karen Jansen-West, Giulia Del Rosso, Lillian M. Daughrity, Monica Castanedes-Casey, Aishe Kurti, Jeannette N. Stankowski, Matthew D. Disney, Jeffrey D. Rothstein, Dennis W. Dickson, John D. Fryer, Yong Jie Zhang, and Leonard Petrucelli. 2019. ‘Aberrant deposition of stress granule-resident proteins linked to C9orf72-associated TDP-43 proteinopathy’, Molecular Neurodegeneration, 14: 1–15.

Chew, Jeannie, Tania F. Gendron, Mercedes Prudencio, Hiroki Sasaguri, Yong Jie Zhang, Monica Castanedes-Casey, Chris W. Lee, Karen Jansen-West, Aishe Kurti, Melissa E. Murray, Kevin F. Bieniek, Peter O. Bauer, Ena C. Whitelaw, Linda Rousseau, Jeannette N. Stankowski, Caroline Stetler, Lillian M. Daughrity, Emilie A. Perkerson, Pamela Desaro, Amelia Johnston, Karen Overstreet, Dieter Edbauer, Rosa Rademakers, Kevin B. Boylan, Dennis W. Dickson, John D. Fryer, and Leonard Petrucelli. 2015. ‘C9ORF72 repeat expansions in mice cause TDP-43 pathology, neuronal loss, and behavioral deficits’, Science, 348: 1151–54.

Copeland, N. G., N. A. Jenkins, and D. L. Court. 2001. ‘Recombineering: a powerful new tool for mouse functional genomics’, Nature Reviews: Genetics, 2: 769–79.

DeJesus-Hernandez, Mariely, Ian R. Mackenzie, Bradley F. Boeve, Adam L. Boxer, Matt Baker, Nicola J. Rutherford, Alexandra M. Nicholson, Ni Cole A. Finch, Heather Flynn, Jennifer Adamson, Naomi Kouri, Aleksandra Wojtas, Pheth Sengdy, Ging Yuek R. Hsiung, Anna Karydas, William W. Seeley, Keith A. Josephs, Giovanni Coppola, Daniel H. Geschwind, Zbigniew K. Wszolek, Howard Feldman, David S. Knopman, Ronald C. Petersen, Bruce L. Miller, Dennis W. Dickson, Kevin B. Boylan, Neill R. Graff-Radford, and Rosa Rademakers. 2011. ‘Expanded GGGGCC Hexanucleotide Repeat in Noncoding Region of C9ORF72 Causes Chromosome 9p-Linked FTD and ALS’, Neuron, 72: 245–56.

Devoy, Anny, Georgia Price, Francesca De Giorgio, Rosie Bunton-Stasyshyn, David Thompson, Samanta Gasco, Alasdair Allan, Gemma F. Codner, Remya R. Nair, Charlotte Tibbit, Ross McLeod, Zeinab Ali, Judith Noda, Alessandro Marrero-Gagliardi, José M. Brito-Armas, Chloe Williams, Muhammet M. Öztürk, Michelle Simon, Edward O’Neill, Sam Bryce-Smith, Jackie Harrison, Gemma Atkins, Silvia Corrochano, Michelle Stewart, Jonathan D. Gilthorpe, Lydia Teboul, Abraham Acevedo-Arozena, Elizabeth M. C. Fisher, and Thomas J. Cunningham. 2021. ‘Generation and analysis of innovative genomically humanized knockin SOD1, TARDBP (TDP-43), and FUS mouse models’, iScience, 24: 103463–63.

Foley, K. E., A. A. Hewes, D. T. Garceau, K. P. Kotredes, G. W. Carter, M. Sasner, and G. R. Howell. 2022. ‘The APOE (epsilon3/epsilon4) Genotype Drives Distinct Gene Signatures in the Cortex of Young Mice’, Front Aging Neurosci, 14: 838436.

Godiska, R., D. Mead, V. Dhodda, C. Wu, R. Hochstein, A. Karsi, K. Usdin, A. Entezam, and N. Ravin. 2010. ‘Linear plasmid vector for cloning of repetitive or unstable sequences in Escherichia coli’, Nucleic Acids Research, 38: e88.

Goodwin, L. O., E. Splinter, T. L. Davis, R. Urban, H. He, R. E. Braun, E. J. Chesler, V. Kumar, M. van Min, J. Ndukum, V. M. Philip, L. G. Reinholdt, K. Svenson, J. K. White, M. Sasner, C. Lutz, and S. A. Murray. 2019. ‘Large-scale discovery of mouse transgenic integration sites reveals frequent structural variation and insertional mutagenesis’, Genome Research, 29: 494–505.

Hofemeister, H., G. Ciotta, J. Fu, P. M. Seibert, A. Schulz, M. Maresca, M. Sarov, K. Anastassiadis, and A. F. Stewart. 2011. ‘Recombineering, transfection, Western, IP and ChIP methods for protein tagging via gene targeting or BAC transgenesis’, Methods, 53: 437–52.

Huynh, T. V., C. Wang, A. C. Tran, G. T. Tabor, T. E. Mahan, C. M. Francis, M. B. Finn, R. Spellman, M. Manis, R. E. Tanzi, J. D. Ulrich, and D. M. Holtzman. 2019. ‘Lack of hepatic apoE does not influence early Abeta deposition: observations from a new APOE knock-in model’, Mol Neurodegener, 14: 37.

Jiang, J., Q. Zhu, T. F. Gendron, S. Saberi, M. McAlonis-Downes, A. Seelman, J. E. Stauffer, P. Jafar-Nejad, K. Drenner, D. Schulte, S. Chun, S. Sun, S. C. Ling, B. Myers, J. Engelhardt, M. Katz, M. Baughn, O. Platoshyn, M. Marsala, A. Watt, C. J. Heyser, M. C. Ard, L. De Muynck, L. M. Daughrity, D. A. Swing, L. Tessarollo, C. J. Jung, A. Delpoux, D. T. Utzschneider, S. M. Hedrick, P. J. de Jong, D. Edbauer, P. Van Damme, L. Petrucelli, C. E. Shaw, C. F. Bennett, S. Da Cruz, J. Ravits, F. Rigo, D. W. Cleveland, and C. Lagier-Tourenne. 2016. ‘Gain of Toxicity from ALS/FTD-Linked Repeat Expansions in C9ORF72 Is Alleviated by Antisense Oligonucleotides Targeting GGGGCC-Containing RNAs’, Neuron, 90: 535–50.

Knouff, C., M. E. Hinsdale, H. Mezdour, M. K. Altenburg, M. Watanabe, S. H. Quarfordt, P. M. Sullivan, and N. Maeda. 1999. ‘Apo E structure determines VLDL clearance and atherosclerosis risk in mice’, J Clin Invest, 103: 1579–86.

Kojak, N., J. Kuno, K. E. Fittipaldi, A. Khan, D. Wenger, M. Glasser, R. A. Donnianni, Y. Tang, J. Zhang, K. Huling, R. Ally, A. O. Mujica, T. Turner, G. Magardino, P. Y. Huang, S. Y. Kerk, G. Droguett, M. Prissette, J. Rojas, T. Gomez, A. Gagliardi, C. Hunt, J. S. Rabinowitz, G. Gong, W. Poueymirou, E. Chiao, B. Zambrowicz, C. J. Siao, and D. Kajimura. 2024. ‘Somatic and intergenerational G4C2 hexanucleotide repeat instability in a human C9orf72 knock-in mouse model’, Nucleic Acids Research, 52: 5732–55.

Liu, Y., A. Pattamatta, T. Zu, T. Reid, O. Bardhi, D. R. Borchelt, A. T. Yachnis, and L. P. Ranum. 2016. ‘C9orf72 BAC Mouse Model with Motor Deficits and Neurodegenerative Features of ALS/FTD’, Neuron, 90: 521–34.

Mann, K. M., F. E. Thorngate, Y. Katoh-Fukui, H. Hamanaka, D. L. Williams, S. Fujita, and B. T. Lamb. 2004. ‘Independent effects of APOE on cholesterol metabolism and brain Abeta levels in an Alzheimer disease mouse model’, Hum Mol Genet, 13: 1959–68.

Mayl, K., C. E. Shaw, and Y. B. Lee. 2021. ‘Disease Mechanisms and Therapeutic Approaches in C9orf72 ALS-FTD’, Biomedicines, 9.

Meyer, D. E., and A. Chilkoti. 2002. ‘Genetically encoded synthesis of protein-based polymers with precisely specified molecular weight and sequence by recursive directional ligation: examples from the elastin-like polypeptide system’, Biomacromolecules, 3: 357–67.

Milioto, C., M. Carcole, A. Giblin, R. Coneys, O. Attrebi, M. Ahmed, S. S. Harris, B. I. Lee, M. Yang, R. A. Ellingford, R. S. Nirujogi, D. Biggs, S. Salomonsson, M. Zanovello, P. de Oliveira, E. Katona, I. Glaria, A. Mikheenko, B. Geary, E. Udine, D. Vaizoglu, S. Anoar, K. Jotangiya, G. Crowley, D. M. Smeeth, M. L. Adams, T. Niccoli, R. Rademakers, M. van Blitterswijk, A. Devoy, S. Hong, L. Partridge, A. N. Coyne, P. Fratta, D. R. Alessi, B. Davies, M. A. Busche, L. Greensmith, E. M. C. Fisher, and A. M. Isaacs. 2024. ‘PolyGR and polyPR knock-in mice reveal a conserved neuroprotective extracellular matrix signature in C9orf72 ALS/FTD neurons’, Nature Neuroscience, 27: 643–55.

Mizielinska, Sarah, S. Gronke, Teresa Niccoli, Charlotte E. Ridler, Emma L. Clayton, Anny Devoy, Thomas Moens, Frances E. Norona, I. O. C. Woollacott, Julian Pietrzyk, Karen Cleverley, Andrew J. Nicoll, Stuart Pickering-Brown, Jacqueline Dols, Melissa Cabecinha, Oliver Hendrich, Pietro Fratta, E. M. C. Fisher, Linda Partridge, and Adrian M. Isaacs. 2014. ‘C9orf72 repeat expansions cause neurodegeneration in Drosophila through arginine-rich proteins’, Science, 345: 1192–94.

Mordes, D. A., B. M. Morrison, X. H. Ament, C. Cantrell, J. Mok, P. Eggan, C. Xue, J. Y. Wang, K. Eggan, and J. D. Rothstein. 2020. ‘Absence of Survival and Motor Deficits in 500 Repeat C9ORF72 BAC Mice’, Neuron, 108: 775–83 e4.

Nair, Remya R., Silvia Corrochano, Samanta Gasco, Charlotte Tibbit, David Thompson, Cheryl Maduro, Zeinab Ali, Pietro Fratta, Abraham Acevedo Arozena, Thomas J. Cunningham, and Elizabeth M. C. Fisher. 2019. ‘Uses for humanised mouse models in precision medicine for neurodegenerative disease’, Mammalian Genome, 30: 173–91.

Nair, Remya R., Charlotte Tibbit, David Thompson, Ross McLeod, Asif Nakhuda, Michelle M. Simon, Robert H. Baloh, Elizabeth M. C. Fisher, Adrian M. Isaacs, and Thomas J. Cunningham. 2021. ‘Sizing, stabilising, and cloning repeat-expansions for gene targeting constructs’, Methods, 191: 15–22.

Nguyen, L., L. A. Laboissonniere, S. Guo, F. Pilotto, O. Scheidegger, A. Oestmann, J. W. Hammond, H. Li, A. Hyysalo, R. Peltola, A. Pattamatta, T. Zu, M. H. Voutilainen, H. A. Gelbard, S. Saxena, and L. P. W. Ranum. 2020. ‘Survival and Motor Phenotypes in FVB C9-500 ALS/FTD BAC Transgenic Mice Reproduced by Multiple Labs’, Neuron, 108: 784–96 e3.

Nordin, A., C. Akimoto, A. Wuolikainen, H. Alstermark, P. Jonsson, A. Birve, S. L. Marklund, K. S. Graffmo, K. Forsberg, T. Brannstrom, and P. M. Andersen. 2015. ‘Extensive size variability of the GGGGCC expansion in C9orf72 in both neuronal and non-neuronal tissues in 18 patients with ALS or FTD’, Human Molecular Genetics, 24: 3133–42.

Nutter, C. A., J. L. Bubenik, R. Oliveira, F. Ivankovic, L. J. Sznajder, B. M. Kidd, B. S. Pinto, B. A. Otero, H. A. Carter, E. A. Vitriol, E. T. Wang, and M. S. Swanson. 2019. ‘Cell-type-specific dysregulation of RNA alternative splicing in short tandem repeat mouse knockin models of myotonic dystrophy’, Genes & Development, 33: 1635–40.

O’Rourke, J. G., L. Bogdanik, Akmg Muhammad, T. F. Gendron, K. J. Kim, A. Austin, J. Cady, E. Y. Liu, J. Zarrow, S. Grant, R. Ho, S. Bell, S. Carmona, M. Simpkinson, D. Lall, K. Wu, L. Daughrity, D. W. Dickson, M. B. Harms, L. Petrucelli, E. B. Lee, C. M. Lutz, and R. H. Baloh. 2015. ‘C9orf72 BAC Transgenic Mice Display Typical Pathologic Features of ALS/FTD’, Neuron, 88: 892–901.

Peters, O. M., G. T. Cabrera, H. Tran, T. F. Gendron, J. E. McKeon, J. Metterville, A. Weiss, N. Wightman, J. Salameh, J. Kim, H. Sun, K. B. Boylan, D. Dickson, Z. Kennedy, Z. Lin, Y. J. Zhang, L. Daughrity, C. Jung, F. B. Gao, P. C. Sapp, H. R. Horvitz, D. A. Bosco, S. P. Brown, P. de Jong, L. Petrucelli, C. Mueller, and R. H. Brown, Jr. 2015. ‘Human C9ORF72 Hexanucleotide Expansion Reproduces RNA Foci and Dipeptide Repeat Proteins but Not Neurodegeneration in BAC Transgenic Mice’, Neuron, 88: 902–09.

Quaegebeur, A., I. Glaria, T. Lashley, and A. M. Isaacs. 2020. ‘Soluble and insoluble dipeptide repeat protein measurements in C9orf72-frontotemporal dementia brains show regional differential solubility and correlation of poly-GR with clinical severity’, Acta Neuropathol Commun, 8: 184.

Rad, R., L. Rad, W. Wang, J. Cadinanos, G. Vassiliou, S. Rice, L. S. Campos, K. Yusa, R. Banerjee, M. A. Li, J. de la Rosa, A. Strong, D. Lu, P. Ellis, N. Conte, F. T. Yang, P. Liu, and A. Bradley. 2010. ‘PiggyBac transposon mutagenesis: a tool for cancer gene discovery in mice’, Science, 330: 1104–7.

Renton, Alan E., Elisa Majounie, Adrian Waite, Javier Simón-Sánchez, Sara Rollinson, J. Raphael Gibbs, Jennifer C. Schymick, Hannu Laaksovirta, John C. van Swieten, Liisa Myllykangas, Hannu Kalimo, Anders Paetau, Yevgeniya Abramzon, Anne M. Remes, Alice Kaganovich, Sonja W. Scholz, Jamie Duckworth, Jinhui Ding, Daniel W. Harmer, Dena G. Hernandez, Janel O. Johnson, Kin Mok, Mina Ryten, Danyah Trabzuni, Rita J. Guerreiro, Richard W. Orrell, James Neal, Alex Murray, Justin Pearson, Iris E. Jansen, David Sondervan, Harro Seelaar, Derek Blake, Kate Young, Nicola Halliwell, Janis Bennion Callister, Greg Toulson, Anna Richardson, Alex Gerhard, Julie Snowden, David Mann, David Neary, Michael A. Nalls, Terhi Peuralinna, Lilja Jansson, Veli Matti Isoviita, Anna Lotta Kaivorinne, Maarit Hölttä-Vuori, Elina Ikonen, Raimo Sulkava, Michael Benatar, Joanne Wuu, Adriano Chiò, Gabriella Restagno, Giuseppe Borghero, Mario Sabatelli, David Heckerman, Ekaterina Rogaeva, Lorne Zinman, Jeffrey D. Rothstein, Michael Sendtner, Carsten Drepper, Evan E. Eichler, Can Alkan, Ziedulla Abdullaev, Svetlana D. Pack, Amalia Dutra, Evgenia Pak, John Hardy, Andrew Singleton, Nigel M. Williams, Peter Heutink, Stuart Pickering-Brown, Huw R. Morris, Pentti J. Tienari, and Bryan J. Traynor. 2011. ‘A hexanucleotide repeat expansion in C9ORF72 is the cause of chromosome 9p21-linked ALS-FTD’, Neuron, 72: 257–68.

Saito, T., Y. Matsuba, N. Mihira, J. Takano, P. Nilsson, S. Itohara, N. Iwata, and T. C. Saido. 2014. ’Single App knock-in mouse models of Alzheimer’s disease’, Nat Neurosci, 17: 661–3.

Saito, T., N. Mihira, Y. Matsuba, H. Sasaguri, S. Hashimoto, S. Narasimhan, B. Zhang, S. Murayama, M. Higuchi, V. M. Y. Lee, J. Q. Trojanowski, and T. C. Saido. 2019. ‘Humanization of the entire murine Mapt gene provides a murine model of pathological human tau propagation’, J Biol Chem, 294: 12754–65.

Simone, R., R. Balendra, T. G. Moens, E. Preza, K. M. Wilson, A. Heslegrave, N. S. Woodling, T. Niccoli, J. Gilbert-Jaramillo, S. Abdelkarim, E. L. Clayton, M. Clarke, M. T. Konrad, A. J. Nicoll, J. S. Mitchell, A. Calvo, A. Chio, H. Houlden, J. M. Polke, M. A. Ismail, C. E. Stephens, T. Vo, A. A. Farahat, W. D. Wilson, D. W. Boykin, H. Zetterberg, L. Partridge, S. Wray, G. Parkinson, S. Neidle, R. Patani, P. Fratta, and A. M. Isaacs. 2018. ‘G-quadruplex-binding small molecules ameliorate C9orf72 FTD/ALS pathology in vitro and in vivo’, EMBO Mol Med, 10: 22–31.

van Blitterswijk, M., M. DeJesus-Hernandez, E. Niemantsverdriet, M. E. Murray, M. G. Heckman, N. N. Diehl, P. H. Brown, M. C. Baker, N. A. Finch, P. O. Bauer, G. Serrano, T. G. Beach, K. A. Josephs, D. S. Knopman, R. C. Petersen, B. F. Boeve, N. R. Graff-Radford, K. B. Boylan, L. Petrucelli, D. W. Dickson, and R. Rademakers. 2013. ‘Association between repeat sizes and clinical and pathological characteristics in carriers of C9ORF72 repeat expansions (Xpansize-72): a cross-sectional cohort study’, Lancet Neurol, 12: 978–88.

Zhu, Fei, Remya R. Nair, Elizabeth M. C. Fisher, and Thomas J. Cunningham. 2019. ‘Humanising the mouse genome piece by piece’, Nature Communications, 10: 1845–45.

