## Supplemental files for "Generation of *C9orf72^h370^* mice, an intron 1 humanised *C9orf72* repeat-expansion knock-in model"

### Supplementary File 1

### Transgene analysis and integration site sequencing of 1 mouse spleen sample containing vector hC9orf72 short humanisation construct

| | |
|---|---|
| Prepared for: | MRC Harwell |
| | Mammalian Genetics Unit, Harwell Campus, Oxford, OX110RD, GB |
| Customer name: | David Thompson |
| | Scientist - Mouse Models of Neurodegeneration group |
| | |
| Internal project number: | 2888-B |
| Quote number: | 2023 - 3935 |
| Version: | 1 |
| Date: | 14-Aug-2023 |

### Goal

In this study, 1 transgenic mouse sample with the hC9orf72 short humanisation construct vector sequence was analyzed.

The aim of this analysis was to:

1. Study the vector integrity:
    - Determine the presence of sequence variants and their allele frequency.
    - Determine the presence of vector-vector breakpoints that represent concatemerization of multiple copies of the vector and/or structural rearrangements in a single vector sequence.
2. Identify vector integration site(s) and breakpoint sequences between the vector and genome.
3. Assess the presence of structural variants surrounding the vector integration site(s).

An overview of the TLA technology and technical details of the performed analyses is provided in the manual “Introduction to the terminology and methods used in TLA analyses_v2”.

### Summary

| Sample | Vector Integrity | Integration site(s) | Structural variants at the integration site | Notes |
|---|---|---|---|---|
| **hC9orf72 short humanisation** | 0 sequence variants, 0 structural variants | chr4:35,218,855-35,225,929 | 7 kb deletion | Correct targeting |

### Conclusion

The generated data shows that the vector integrated correctly at the targeted location. A 7 kb genomic deletion was observed at the integration site. Additionally, no sequence and structural variants were identified within the integrated vector.

### TLA, sequencing and data mapping

Viable frozen mouse splenocytes were used and processed according to Cergentis' TLA protocol (de Vree et al. Nat Biotechnol. Oct 2014). An overview of the TLA technology and technical details of the performed analyses is provided in the manual "Introduction to the terminology and methods used in TLA analyses_v2".

TLA was performed with 2 independent primer sets specific for the vector sequence (Table 1).

**Table 1**: Primers used in TLA analysis

| Primer set | Name/VP | Direction | Binding position<br>*hC9orf72 short humanisation construct* | Sequence |
|---|---|---|---|---|
| **1** | NEO | RV | 8,517 | TGTTCGCCAGGCTCAAGG |
| | | FW | 8,654 | GTAGCCGGATCAAGCGTATG |
| **2** | hC9orf72 intron 1 | RV | 22,023 | TCAATTCCTAACCCTTGGTG |
| | | FW | 22,143 | GCCAAATCTCCAGTCATACT |

The NGS reads were aligned to the vector sequence, human hg38 and host genome. The mouse mm10 genome was used as host reference genome sequence.

### Results hC9orf72 short humanisation

#### Vector integrity

Figure 1 depicts the NGS coverage across the vector sequence using primer sets 1 and 2.

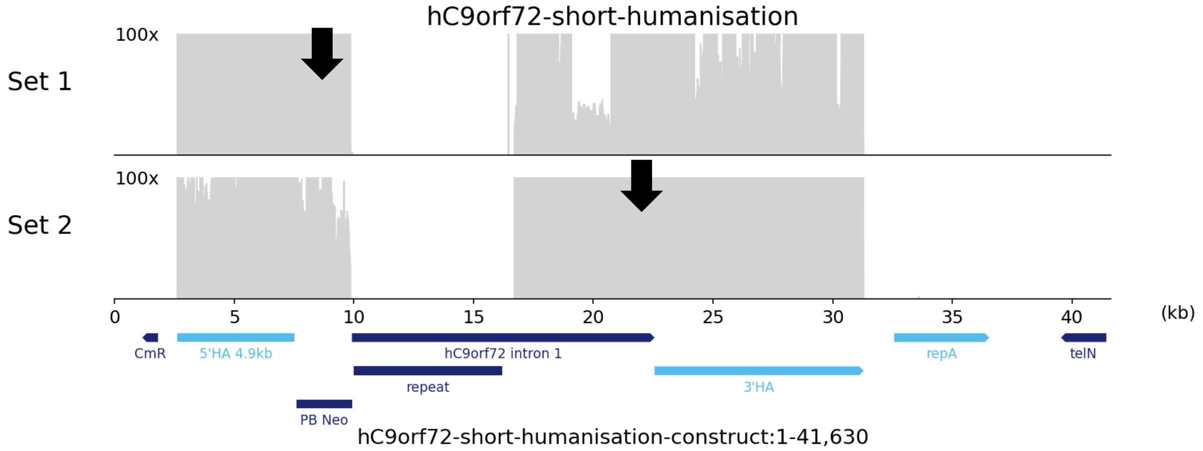

**Figure 1a:** NGS sequencing coverage (in grey) across the vector. Black arrows indicate the primer location. The vector map is shown on the bottom. Y-axes are limited to 100x.

Coverage is observed across the vector sequence Vector: 2,625-9,858 and Vector: 16,707-31,299, within the homology arms. No coverage is observed across the Vector: 1-2,624 (CmR) and Vector: 31,300-41,630 (repA and telN), indicating that the backbone has not integrated in this sample. Moreover, no coverage is observed across the vector sequence Vector: 9,859-16,706 (repeat and part of hC9orf72 intron 1) due to the nature of the sequence. Local dips in coverage are due to GC rich regions that are less efficiently sequenced and due to the nature of the sequence.

Sequence variants and structural variants were called in the covered regions.

##### Sequence variants

No sequence variants were detected in this sample

##### Vector concatemerization and structural variants

No vector concatemerization and structural variants were detected in this sample.

### Integration sites

#### Whole genome coverage plot

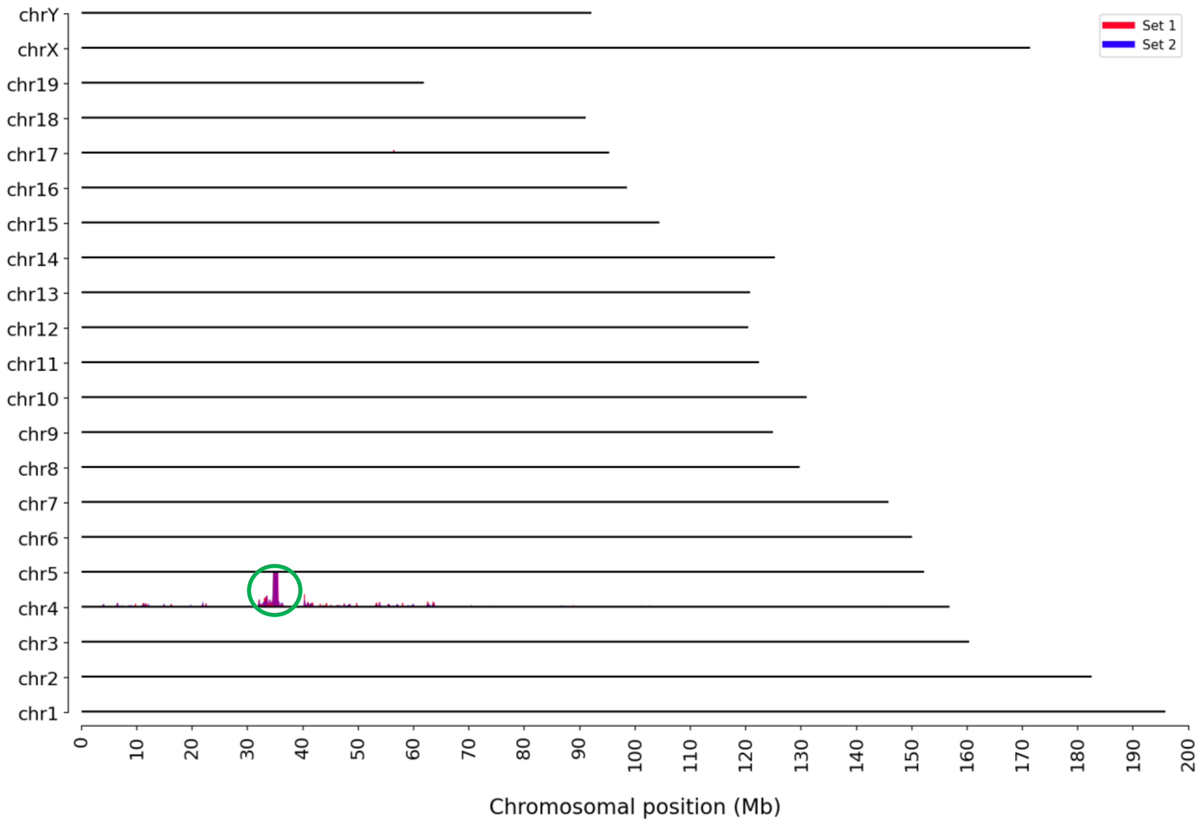

**Figure 2:** TLA sequence coverage across the mouse genome using primer set 1 (red) and set 2 (blue). The chromosomes are indicated on the y-axis, the chromosomal position on the x-axis. Identified integration site is encircled in green.

As shown in Figure 2, the vector has integrated on chromosome 4 at the intended location.

#### Locus-wide coverage

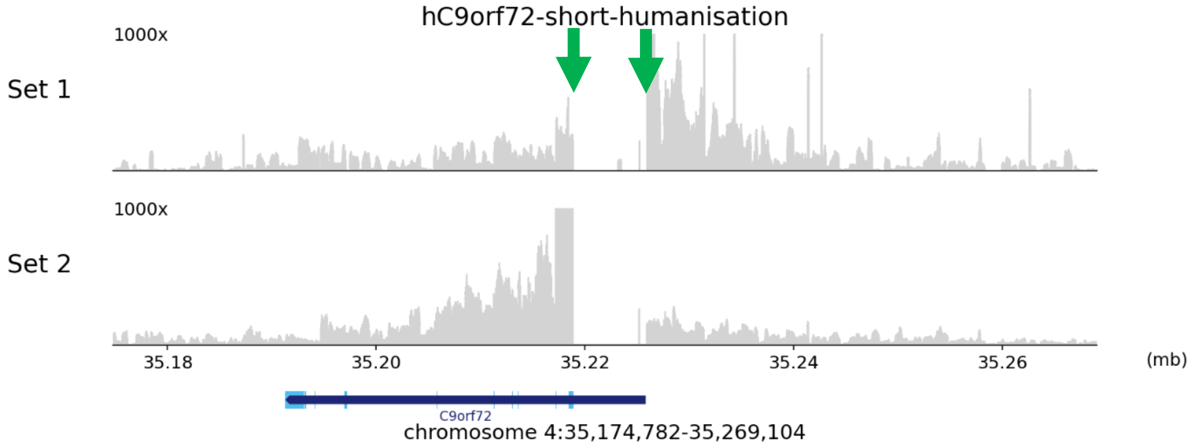

**Figure 3:** TLA sequence coverage (in grey) across the vector integration locus, mouse chr4:35,174,782-35,269,104. The green arrows indicate the location of the breakpoint sequences. Y-axes are limited to 1000x.

Coverage is observed across the vector integration site as shown in Figure 3.

##### Breakpoint sequences

When the data is aligned to the genome, the following reads at the inner edges of the homology arms indicate the vector integration at the intended location:

5' integration site:
chr4:35,218,855 (tail) fused to Vector:22,562 (tail)
AGCAAAGGTAGCCGCCAACAAGGGTGATTCACCACTTAAAGCAATCTCTGTCTTGGCAACAGCAGGA
GATGGTGGGGGGCAGATAGTCGACATCACTGCATTCCAACTGTCACATTATCCAAATGCTCCGGAGAT
ATCTAAACAATGACATATGAAACCAATGATTAGGTTCAGCAATTTAAAGATATCCATCAAAACCCCAAA
TGATTTAGACATATTTGGTT

3' integration site:
Vector:7,524 (head) fused to chr4:35,225,929 (head)
CATGCGTCAATTTTACGCAGACTATCTTTCTAGGGTTAATCTTTATCAGGTCTTTTCTTGTTCACCCTCA
GCGAGTACTGTGAGAGCAAGTAGTGGGGAGAGAGGGTGGGAAAAACAAAAACACACACCCAGAATC
TGCAGGGCTCCCCGGTGCGGATCTCTGCAGTCTTGTTTCCTTCTTTTGTCCTTGGTCACCGTTGTGGA
GTTAAGTTGCGAGCTGAGGCAACCAAAACGCACCTCCCTCGCACACC

Please note, due to homology between the vector and the genome sequence the identified breakpoint reads completely align to the vector sequence as well, which is as expected in a correct targeting event.

The coverage profile in figure 3 shows that a genomic deletion has occurred in the region of the integration site. The 7 kb genomic sequence in between the two identified breakpoints is deleted.

Figure 4 depicts the NGS coverage across the expected modified allele using primer sets 1 and 2.

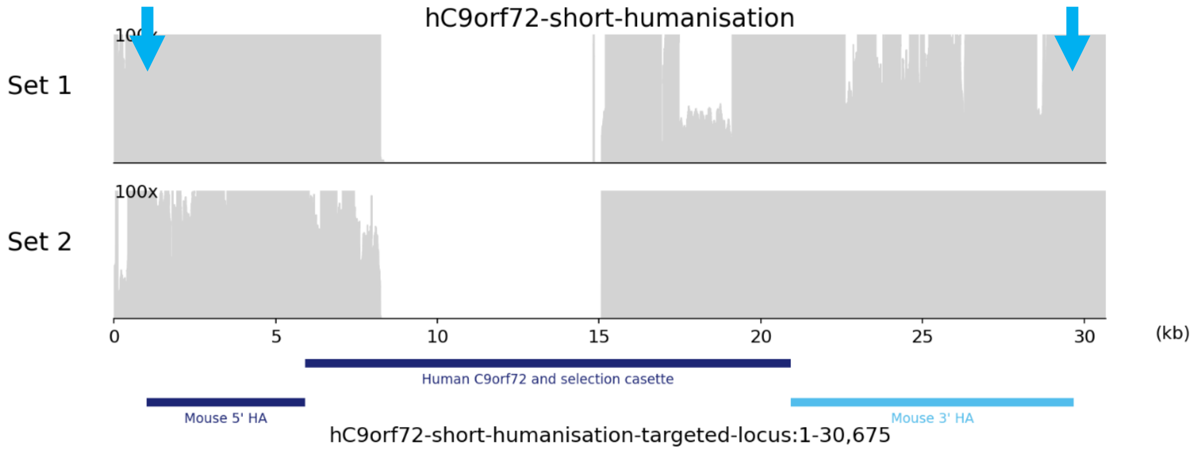

**Figure 4**: TLA sequence coverage (in grey) across the expected modified allele. The blue arrows indicate the integration sites. The integrated features are shown on the bottom. Y-axes are limited to 100x.

Continuous coverage is observed across the modified allele (except the gap due to the nature of the sequence), indicating vector integration. At the outer edge of homology arm (blue arrows) the wild type sequence was identified as expected.

#### Targeting accuracy

The accuracy of a targeting event is checked on the basis of the following criteria:

- Evidence for any structural variation within the vector: No
- Off-target integration site: No
- Sequence coverage on the backbone sequences of the vector: No
- Indication of any structural variation surrounding the integration site (e.g. genomic duplication): No

In this sample, there is no evidence of an incorrect targeting, indicating that the targeting has occurred as intended.

From this data it is concluded that the vector has integrated at mouse chr4:35,218,855-35,225,929 as shown in Figure 5. According to the RefSeq, there is an intron of *C9orf72* at the integration site.

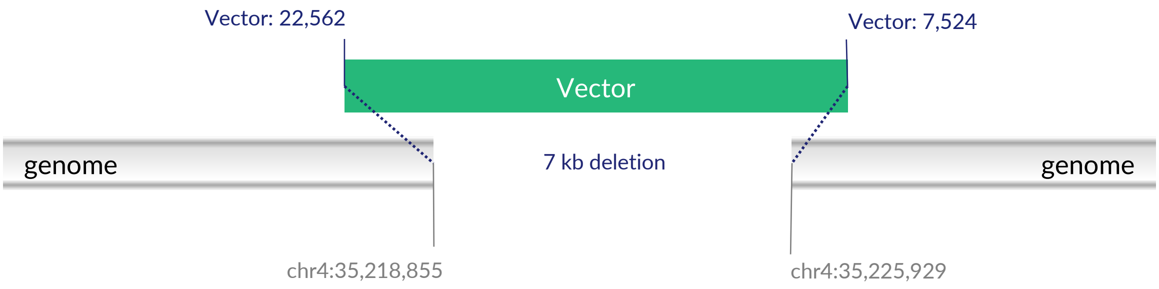

**Figure 5:** Schematic representation of the integration site.

### QC information

#### Sample and Study details

| | |
|---|---|
| Sample receipt date | 07-Jun-2023 |
| Condition of sample at receipt | Frozen |
| Start date in the lab | 12-Jun-2023 |
| Sequencing run | 23-027 |
| Date data analysis | 03-Aug-2023 |
| Deviations from the protocol | No |
| TLApp version: | 1.6.0 |

#### Study Personnel

| | |
|---|---|
| Lab technician | Beatriz Almeida, MSc |
| Data Analyst | Andrea Oneglia, PhD |
| QC Analysis and Report | Irina Sergeeva, PhD |

TESTING
RvA L 671

#### Quality control

The results are independently verified and reviewed and are an accurate and complete representation of the study. The scope of accreditation for ISO/IEC 17025:2017, accredited by the Dutch Accreditation Council RvA, Registration number L671, entails all analytical services including, determination of the integrity of the transgene vector sequence; determination of the vector integration site(s) and breakpoint sequences between the vector and genome, determination of the presence of structural variants surrounding the vector integration site(s), next generation sequencing (NGS) and bio-informatic data analysis.

| | |
|---|---|
| Scientific approval | Cheryl Dambrot, PhD - Scientific Account Manager |
| Date | 14-Aug-2023 |
| Signature | |

### Supplementary File 2

#### 4x G4C2 repeat building cassette

See Figure 1A. Annealed pair of DNA oligos; cloned into XmaI digested pJAZZ-OK-delta-AarI
5’-CCGGGGCCGGGGCCGGGGCCGGGGCCGGGGCCGCAGGTGACATCGCTCTTCTGG-3’
3’-CCGGCCCCGGCCCCGGCCCCGGCCCCGGCGTCCACTGTAGCGAGAAGACCGGCC-5’

#### pJazz-V1

See figure 1. Excisable repeat in **red**. XmaI and AarI sites underlined. pJazz backbone in lowercase.

gcgtataatgggcaattgtgtgctgatatgttccgtaattgagatatgatatagtgcatatgttccgaattattgatacattaatgttgcattcctatgttgtgttctg
ttatggtgttcttagatagtggggatgtcacatggattgatgttctttgattgtgtttcaatatagagacggaagagatagagatgcctcaaagtgtataaacagcgt
gaaaatcagcatgttcgcttaacttttgatagcaattggtatgatttattatcatttaaaatcattaagttaggttttaaatataccaacgtaagtaataccgttgcgc
ttagatgaagctaaatagcttctgtagcgcatcaaaaaattgatttgaaccttaactttgcgttttctaagtcactttgctttttttatgttgctttgctttaaaaacg
taaagtatgatgaaaaacacaaagataagggcatgtttgagaggaaacaggatgtcgttaattaatttgctgaaagactgtatcaatcgcggtcaggaaatgacgcgcg
ctattgccatcgcacagttcggcgatgacagcccggaagcgcgtcgtattactcgccgctggggtataacagaagttgcagacctgatcggcgtaacaccgcaggctat
cagagacgcggaaaaagctggtcgtctaccggctcctgattttgagatgaggggccgagtagaacgtcgcgctggctacacaattgaccaaattagccatatgcggagt
gtgtttggtaacccaaaccaaagacctgacgataaaaacccggttgtactttccgttatgtcacacaagggaggggtttataaacctcatctgcagtacaccaggcgc
aatggttagccttgcaagggcaccgcgtcatactcgttgaagggaacgacccgcaagggactgcatctatgtaccacggttacgttcctgatttgcacattcacgcaga
cgatacactgcttccatttacctgggtaaacgcgacaacgctgaatatgcgataaaaccaacatgctggccgggtctcgacataatccccagctgcctggcccttcac
cgcattgaaacagatctagcagaaagtcaaaagcctccgaccggaggcttttgacttctgtcacctaggttagaaaaactcatcgagcatcaagtgaaactgcaattta
ttcatatcaggattatcaataccatatttttgaaaaagccgtttctgtaatgaaggagaaaactcaccgaggcagttccataggatggcaagatcctggtatcggtctg
cgattccgactcgtccaacatcaatacaacctattaatttcccctcgtcaaaaataaggttatcaagtgagaaatcaccatgagtgacgactgaatccggtgagaatgg
caaaagtttatgcatttctttccagacttgttcaacaggccagccattacgctcgtcatcaaaatcactcgcaccaaccaaaccgttattcattcgtgattgcgcctga
gcgagacgaaatacgcgatcgccgttaaaaggacaattacaaacaggaatcgaatgcaaccggcgcaggaacactgccagcgcatcaacaatattttcacctgaatcag
gatattcttctaatacctggaatgctgttttccctgggatcgcagtggtgagtaaccatgcatcatcaggagtacggataaaatgcttgatggtcggaagaggcataaa
ttccgtcagccagtttagcctgaccatctcatctgtaacatcattggcaacgctacctttgccatgtttcagaaacaactctggcgcatcgggcttcccatacaatcga
tagattgtcgcacctgattgcccgacattatcgcgagcccatttatacccatataaatcagcatccatgttggaatttaatcgcggcctcgagcaagacgtttcccgtt
gaatatggctcatttagcttccttagctcctgaaaatctcgataactcaaaaatacgcccggtagtgatcttatttcattatggtgaaagttggaacctcttacgtgc
cgatcagattaaaacgaaaggcccagtctttcgactgagcctttcgttttatttgaccatgttggtatgatttaaatggtcagttaatcagttctatgtaccagcaagg
tccagttgtaagcggccgcttgacttcaagtctaatggcCCGGGGCCGGGGCCGGGGCCGGGGCCGGGGCCGGGGCCGGGGCCGGGGCCGGGGCCGGGGCCGGGGCCGG
GGCCGGGGCCGGGGCCGGGGCCGGGGCCGGGGCCGGGGCCGGGGCCGGGGCCGGGGCCGGGGCCGGGGCCGGGGCCGGGGCCGGGGCCGGGGCCGGGGCCGGGGCCGGG
GCCGGGGCCGGGGCCGGGGCCGGGGCCGGGGCCGGGGCCGGGGCCGGGGCCGGGGCCGGGGCCGGGGCCGGGGCCGGGGCCGGGGCCGGGGCCGGGGCCGGGGCCGGGG
CCGGGGCCGGGGCCGGGGCCGGGGCCGGGGCCGGGGCCGGGGCCGGGGCCGGGGCCGGGGCCGGGGCCGGGGCCGGGGCCGGGGCCGGGGCCGGGGCCGGGGCCGGGGC
CGGGGCCGGGGCCGGGGCCGGGGCCGGGGCCGGGGCCGGGGCCGGGGCCGGGGCCGGGGCCGGGGCCGGGGCCGGGGCCGGGGCCGGGGCCGGGGCCGGGGCCGGGGCC
GGGGCCGGGGCCGGGGCCGGGGCCGGGGCCGGGGCCGGGGCCGGGGCCGGGGCCGGGGCCGGGGCCGGGGCCGGGGCCGGGGCCGGGGCCGGGGCCGGGGCCGGGGCCG
GGGCCGGGGCCGGGGCCGGGGCCGGGGCCGGGGCCGGGGCCGGGGCCGGGGCCGGGGCCGGGGCCGGGGCCGGGGCCGGGGCCGGGGCCGGGGCCGGGGCCGGGGCCGG
GGCCGGGGCCGGGGCCGGGGCCGGGGCCGGGGCCGGGGCCGGGGCCGGGGCCGGGGCCGGGGCCGGGGCCGGGGCCGGGGCCGGGGCCGGGGCCGGGGCCGGGGCCGGG
GCCGGGGCCGGGGCCGGGGCCGGGGCCGGGGCCGGGGCCGGGGCCGGGGCCGGGGCCGGGGCCGGGGCCGGGGCCGGGGCCGGGGCCGGGGCCGGGGCCGGGGCCGGGG
CCGGGGCCGGGGCCGGGGCCGGGGCCGGGGCCGGGGCCGGGGCCGGGGCCGGGGCCGGGGCCGGGGCCGGGGCCGGGGCCGGGGCCGGGGCCGGGGCCGGGGCCGGGGC
CGGGGCCGGGGCCGGGGCCGGGGCCGGGGCCGGGGCCGGGGCCGGGGCCGGGGCCGGGGCCGGGGCCGGGGCCGGGGCCGGGGCCGGGGCCGGGGCCGGGGCCGGGGCC
GGGGCCGGGGCCGGGGCCGGGGCCGGGGCCGGGGCCGGGGCCGGGGCCGGGGCCGGGGCCGGGGCCGGGGCCGGGGCCGGGGCCGGGGCCGGGGCCGGGGCCGGGGCCG
GGGCCGGGGCCGGGGCCGGGGCCGGGGCCGGGGCCGGGGCCGGGGCCGGGGCCGGGGCCGGGGCCGGGGCCGGGGCCGGGGCCGGGGCCGGGGCCGGGGCCGGGGCCGG
GGCCGGGGCCGGGGCCGGGGCCGGGGCCGGGGCCGGGGCCGGGGCCGGGGCCGGGGCCGGGGCCGGGGCCGGGGCCGGGGCCGGGGCCGGGGCCGGGGCCGGGGCCGGG
GCCGGGGCCGGGGCCGGGGCCGGGGCCGGGGCCGGGGCCGGGGCCGGGGCCGGGGCCGGGGCCGGGGCCGGGGCCGGGGCCGGGGCCGGGGCCGGGGCCGGGGCCGGGG
CCGGGGCCGGGGCCGGGGCCGGGGCCGGGGCCGGGGCCGGGGCCGGGGCCGGGGCCGGGGCCGGGGCCGGGGCCGGGGCCGGGGCCGGGGCCGGGGCCGGGGCCGGGGC
CGGGGCCGGGGCCGGGGCCGGGGCCGGGGCCGGGGCCGGGGCCGGGGCCGGGGCCGGGGCCGGGGCCGGGGCCGGGGCCGGGGCCGGGGCCGGGGCCGGGGCCGGGGCC
GGGGCCGGGGCCGGGGCCGGGGCCGGGGCCGGGGCCGGGGCCGGGGCCGGGGCCGGGGCCGGGGCCGGGGCCGGGGCCGGGGCCGGGGCCGGGGCCGGGGCCGGGGCCG
GGGCCGGGGCCGGGGCCGGGGCCGGGGCCGGGGCCGGGGCCGGGGCCGGGGCCGGGGCCGGGGCCGGGGCCGGGGCCGGGGCCGGGGCCGGGGCCGGGGCCGGGGCCGG
GGCCGGGGCCGGGGCCGGGGCCGGGGCCGGGGCCGGGGCCGGGGCCGGGGCCGGGGCCGGGGCCGGGGCCGGGGCCGGGGCCGGGGCCGGGGCCGGGGCCGGGGCCGGG
GCCGGGGCCGGGGCCGGGGCCGGGGCCGGGGCCGGGGCCGGGGCCGGGGCCGGGGCCGGGGCCGGGGCCGGGGCCGGGGCCGGGGCCGGGGCCGGGGCCGGGGCCGGGG
CCGGGGCCGGGGCCGGGGCCGGGGCCGGGGCCGGGGCCGGGGCCGGGGCCGGGGCCGGGGCCGGGGCCGGGGCCGGGGCCGGGGCCGGGGCCGGGGCCGGGGCCGGGGC
CGGGGCCGGGGCCGGGGCCGGGGCCGGGGCCGGGGCCGGGGCCGGGGCCGGGGCCGGGGCCGGGGCCGGGGCCGGGGCCGGGGCCGGGGCCGGGGCCGGGGCCGGGGCC
GGGGCCGGGGCCGGGGCCGGGGCCGGGGCCGGGGCCGGGGCCGGGGCCGGGGCCGGGGCCGGGGCCGGGGCCGGGGCCGGGGCCGGGGCCGGGGCCGGGGCCGGGGCCG
GGGCCGGGGCCGGGGCCGGGGCCGGGGCCGGGGCCGGGGCCGGGGCCGGGGCCGGGGCCGGGGCCGGGGCCGGGGCCGGGGCCGGGGCCGGGGCCGGGGCCGGGGCCGG
GGCCGGGGCCGGGGCCGGGGCCGGGGCCGGGGCCGGGGCCGGGGCCGGGGCCGGGGCCGGGGCCGGGGCCGGGGCCGGGGCCGGGGCCGGGGCCGGGGCCGGGGCCGGG
GCCGGGGCCGGGGCCGGGGCCGGGGCCGGGGCCGGGGCCGGGGCCGGGGCCGGGGCCGGGGCCGGGGCCGGGGCCGGGGCCGGGGCCGGGGCCGGGGCCGGGGCCGGGG
CCGGGGCCGGGGCCGGGGCCGGGGCCGGGGCCGGGGCCGGGGCCGGGGCCGGGGCCGGGGCCGGGGCCGGGGCCGGGGCCGGGGCCGGGGCCGGGGCCGGGGCCGGGGC
CGGGGCCGGGGCCGGGGCCGGGGCCGGGGCCGGGGCCGGGGCCGGGGCCGGGGCCGGGGCCGGGGCCGGGGCCGGGGCCGGGGCCGGGGCCGGGGCCGGGGCCGGGGCC
GGGGCCGGGGCCGGGGCCGGGGCCGGGGCCGGGGCCGGGGCCGGGGCCGGGGCCGGGGCCGGGGCCGGGGCCGGGGCCGGGGCCGGGGCCGGGGCCGGGGCCGGGGCCG
GGGCCGGGGCCGGGGCCGGGGCCGGGGCCGGGGCCGGGGCCGGGGCCGGGGCCGGGGCCGGGGCCGGGGCCGGGGCCGGGGCCGGGGCCGGGGCCGGGGCCGGGGCCGG
GGCCGGGGCCGGGGCCGGGGCCGGGGCCGGGGCCGGGGCCGGGGCCGGGGCCGGGGCCGGGGCCGGGGCCGGGGCCGGGGCCGGGGCCGGGGCCGGGGCCGGGGCCGGG
GCCGGGGCCGGGGCCGGGGCCGGGGCCGGGGCCGGGGCCGGGGCCGGGGCCGGGGCCGGGGCCGGGGCCGGGGCCGGGGCCGGGGCCGGGGCCGGGGCCGGGGCCGGGG
CCGGGGCCGGGGCCGGGGCCGGGGCCGGGGCCGGGGCCGGGGCCGGGGCCGGGGCCGGGGCCGGGGCCGGGGCCGGGGCCGGGGCCGGGGCCGGGGCCGGGGCCGGGGC
CGGGGCCGGGGCCGGGGCCGGGGCCGGGGCCGGGGCCGGGGCCGGGGCCGGGGCCGGGGCCGGGGCCGGGGCCGGGGCCGGGGCCGGGGCCGGGGCCGGGGCCGGGGCC
GGGGCCGGGGCCGGGGCCGGGGCCGGGGCCGGGGCCGGGGCCGGGGCCGGGGCCGGGGCCGGGGCCGGGGCCGGGGCCGGGGCCGGGGCCGGGGCCGGGGCCGGGGCCG
GGGCCGGGGCCGGGGCCGGGGCCGGGGCCGGGGCCGGGGCCGGGGCCGGGGCCGGGGCCGGGGCCGGGGCCGGGGCCGGGGCCGGGGCCGGGGCCGGGGCCGGGGCCGG
GGCCGGGGCCGGGGCCGGGGCCGGGGCCGGGGCCGGGGCCGGGGCCGGGGCCGGGGCCGGGGCCGGGGCCGGGGCCGGGGCCGGGGCCGGGGCCGGGGCCGGGGCCGGG
GCCGGGGCCGGGGCCGGGGCCGGGGCCGGGGCCGGGGCCGGGGCCGGGGCCGGGGCCGGGGCCGGGGCCGGGGCCGGGGCCGGGGCCGGGGCCGGGGCCGGGGCCGGGG
CCGGGGCCGGGGCCGGGGCCGGGGCCGGGGCCGGGGCCGGGGCCGGGGCCGGGGCCGGGGCCGGGGCCGGGGCCGGGGCCGGGGCCGGGGCCGGGGCCGGGGCCGGGGC
CGGGGCCGGGGCCGGGGCCGGGGCCGGGGCCGGGGCCGGGGCCGGGGCCGGGGCCGGGGCCGGGGCCGGGGCCGGGGCCGGGGCCGGGGCCGGGGCCGGGGCCGGGGCC
GGGGCCGGGGCCGGGGCCGGGGCCGGGGCCGGGGCCGGGGCCGGGGCCGGGGCCGGGGCCGGGGCCGGGGCCGGGGCCGGGGCCGGGGCCGGGGCCGGGGCCGGGGCCG
GGGCCGGGGCCGGGGCCGGGGCCGGGGCCGGGGCCGGGGCCGGGGCCGGGGCCGGGGCCGGGGCCGGGGCCGGGGCCGGGGCCGGGGCCGGGGCCGGGGCCGGGGCCGG
GGCCGGGGCCGGGGCCGGGGCCGGGGCCGGGGCCGGGGCCGGGGCCGGGGCCGGGGCCGGGGCCGGGGCCGGGGCCGGGGCCGGGGCCGGGGCCGGGGCCGGGGCCGGG

GCCGGGGCCGGGGCCGGGGCCGGGGCCGGGGCCGGGGCCGGGGCCGGGGCCGGGGCCGGGGCCGGGGCCGGGGCCGGGGCCGGGGCCGGGGCCGGGGCCGGGGCCGGGGCCGGGGCCGGGG
CCGGGGCCGGGGCCGGGGCCGGGGCCGGGGCCGGGGCCGGGGCCGGGGCCGGGGCCGGGGCCGGGGCCGGGGCCGGGGCCGGGGCCGGGGCCGGGGCCGGGGCCGGGGCCGGGGCCGGGGC
CGGGGCCGGGGCCGGGGCCGGGGCCGGGGCCGGGGCCGGGGCCGGGGCCGGGGCCGGGGCCGGGGCCGGGGCCGGGGCCGGGGCCGGGGCCGGGGCCGGGGCCGGGGCCGGGGCCGGGGCC
GGGGCCGGGGCCGGGGCCGGGGCCGGGGCCGGGGCCGGGGCCGGGGCCGGGGCCGGGGCCGGGGCCGGGGCCGGGGCCGGGGCCGGGGCCGGGGCCGGGGCCGGGGCCGGGGCCGGGGCCG
GGGCCGGGGCCGGGGCCGGGGCCGGGGCCGGGGCCGGGGCCGGGGCCGGGGCCGGGGCCGGGGCCGGGGCCGGGGCCGGGGCCGGGGCCGGGGCCGGGGCCGGGGCCGGGGCCGGGGCCGG
GGCCGGGGCCGGGGCCGGGGCCGGGGCCGGGGCCGGGGCCGGGGCCGGGGCCGGGGCCGGGGCCGGGGCCGGGGCCGGGGCCGGGGCCGGGGCCGGGGCCGGGGCCGGGGCCGGGGCCGGG
GCCGGGGCCGGGGCCGGGGCCGGGGCCGGGGCCGGGGCCGGGGCCGGGGCCGGGGCCGGGGCCGGGGCCGGGGCCGGGGCCGGGGCCGGGGCCGGGGCCGCAGGTGACATCGCTCTTCTGG
ccgggccattgggccctagacaagttgtcgcggccgcactagtgactccagcgtaactggactggccacagttaggccgcaaatgtaatcacactggctcaccttcggg
tgggcctttctgcgtctagaggatcccctgttatgcccttccttacccatgcgctgagaggtgaaatcactctttaatttgtacgcggtacggacctctatatcgcttt
gtcgtagaggtggaatctccccagcttgcagccattgatagactgcgccgggtgtcacgcccacgccaaccgctgctttttccacatcgccgaaatggcgcacaagttc
ttcgggtttcatggggattattataatcaataagtgaaaattaaagctaggtataatttataaaaattatagccagctataaagagatcatttatgattaagggtatga
aaacacgaggcgaacgactgaaagcacgccgtttagagctgaaactgacgctgaagcaagtggcagaagctgtaggaatctctcttccgggcgtccaaaacttagaacg
tggcgacgtgatgccgtcgctggagatcggctatcgctggcaaaatgcctgcgtaagcccgtgcaatggatactgtatggcactgaatctgatccagaccgcgttcct
gttattggtacgacagagaccggaccggatagagactggcagcctggagaacctgccaacacagagcgattcctgccgttcgtgagtcaacggaacaccgtttacgcac
tgacggtcgggaaccagattcagcgaaactaccagccgggagacgttatcctggtcgattcttcacttacgcttgtaccgggtgaggatgtactggtttgtgataataa
cggcgagatcacaattcaacgattagcccgttatgacgagcagcattactaccttgatagtgctaactctcaacgggttatccatgataaaagtgatcttcaatttgtg
caccaagtagtcggtacgatcaaatcgttcatggttgagggtagatgatacaataacagggtttattgctgactataattctggtttaatccgatctatactttgtcgg
ggttgaaccagaccgtagcagccgaaaaagacgaaaaaaacccgagtcggcaaactcggcctttttcaggaagtagccacgataacgcagatacgtccttcagaaa
gatagtgcgtttattgtggctgctcctgcggatttttcaacccgaaaaaatgctaattcgcatggaaaggctaaaaaatgaccttacaagaattctacgcggagcgct
ttggcagcgatccgttttcactgcttgacgcggcacgtgatgagctgactgagctggcgaagatggccggtatcaactggcctgcgtgtgctgataagattcagctcaa
tccgcgcggtggtgtagagcgctataccacctacaataattcctccccggaagctctgcagaagagccttaaaggccgtgtggaaatctattcacgccttgagcagagc
aaggatggcatcagttatcctttcgttaacttcgttgaaaaagcgcatgacgctggctcctggagtggcttttcattcctgttttcggagtatcgccgtgaacagcaac
ggaatggtgcgaccgtggtcgctcagccggaagaagaacgtgcacgcatggagcgccaggctgaagcacgccgcctccgagctgaacaacaacgcgttaatgatttaaa
aaacaatcagatggagcaggaacgcctgcttggatggttggctttccatcgcgcctgggaacattcgccagctgaagatggctcctggccttatgcggtaaaaaggga
attcgtgacgtatttggcgcttgtgatatacgtcgcgtgaccagtcacgacagtgcgaaatggagccgtggaccaacaacttacatggcgatacctctgtcccacctgg
acggaagaaaagacggcggattgttggctggcaacgtatcgatctgaatggtggcaagttccagaccagcgccatcactaacggtgatttcgtcggtgcatgctttgt
tattggcgacctgcaggggcgcaaaaaattgcgactgcagagggtttcgccaccggggcatccatctggctggcgaccagaaacgacccgaaaaaacgctttgacgcg
gtagtcattgcggtttccgctaacaacatgattcacgttgtcgagcagctggtgaacatgtacccggctgcgcaaatcacttgcgctctcgataacgaccgcaaatcct
cagctgaaggaaaaggcaacacaggcctccgcaccggattcgacatcatggagaagttttccggcgtcaaatgtgtttacccaacttttgaggatgaccctgagctgga
atgcagcgattttaacgacctgcacagcctgagagggctgaaggaagtcgctcgccagttaacgagaaatcatctgagccgtgcaaccgatctgttgtcgatcacgctg
aataagctgcgtactctcccgcgtctgaacagacgcacttttgccaaagaactgcttcgcgccgttgatattggcatgctgacatgcccggtaccaaacagcccgaaag
aacttatgcgcctttttcagcagcacgctgcgggatatgggtatcgcagaaatttataacggtaccgttaaagatcacattacgcgccggttgaatcgcaaatgccgtgc
cgcgcaaacatcacgttcgttcagtgaacgcatcaccaacccgaaccttcgcccgtcacacatcacttacaaacggtttgaaacctccaggatgaccgatgaggtgatg
acatacgccgcacagctgcagggcatcgttattgtccgcgccgggatggggtcgggtaaatcgacaggcctcctgcgtccactgatgctgcagtccacgcgtggcgttt
ccgtcgcgcaccgcgtatccctgataggcggcctgcatgaaatgatgaccgaagggaaaggcgcgaaagccgacattctgcattatcaggatcccggctatcaggaaat
ggcgccatatgccaacaagctgactatttgcatcaactccatacttaaaggctgctggcaaccgctgatgcgccagcatgacttcttcggcttcgatgaagcaacacag
ggcctgcgtgccattctggccgggcgtgcgatggaaaacccggtaggcgtattcaacacgcttatcgacgcgctggcgcgtactgaagagcatgccattatggtggatg
ccgatgccaacgatctgcttgttgacctggctgaactggcgatgaagcgacgcgaggagctgggcctacctgcctggctgcaaattcacgtgattgaactcccggtcga
cgttcgcaaccgcgaaaccaacaagcctatccgcgtattttataccgagaaaaatcggatcatgaccgaggttatcgctgcagtgcaacgcggcgaacgaatcatgctg
gccaccgacagttcgacgttcgccgaagacgttaccatgcagctgagactacagttccctgacaaaaagtttctctgcgttaaccagaaaaacaaacaggagaaggaag
tcgacgatttcaccaaccagcctaaagtgatggtgaaaaaatatgacggcctcatctacagcccgtcgatatcttcaggtgtatcgattgaggagaaacacttccaccg
ccatttcggcatgttctgcggcgaagtggtccccagcgacgccatccagatgctgcgccgcgaccgtacagctcaggaatacattatcggtttcgacaagcttcgcggt
aaacgtgaaaccgatccggaaaaaatcaaacgcgcctacgcgcaggccttgctcgaaacggctggccactccgggctgctgacagacgttgtctttgacggtgaccgaa
tttcactcggcgtggctaactcttcattcatgcagctcaaaattaaagccgccgcgctcgaggcatcggccagaaatgattatgccagcaacatgatttgcatcatgca
tgatgatggctatcaggtcgcgcctatggccaccgacgcgcttgcaaacagcatcggtaaggatttacgtaaagaagcccgtgaactggtctttgagcagcttatggag
cgccacctgagtgtcgatactcctgaccaggccgaacacgatgagttgataaaaaaacgcaccctgtccctggatgagcaggcccagctggtccgctgggacatcgaaa
aggagctgcagctggatgtcgacgaggtggcgttaaaattctatttcgacggcggcctgaaaaaagtgcgcctgtttgaaaccatgcagctggatgagataaccgcccg
tcgccttgaccgtgaagaagcgttgattcactttacctacgcttatcgcgttgccggaagatggcagcagttcgtcactacggccatgacgcgggaacaggccgacgca
gagttccaggctaaattcccggccataaccgattaccgagtgaaatcgacccctgctgttgagataggcatgcgtggtttctacaccctcaaatcggcgacgctgcagc
agtacttccgcgactgcggtatcgatccaaaaaccctggaaggtgaagcagacatggacgccctcaaacgcgccagagataacctgctcacgccggaacggcgcgatct
gctaaacaacgttttacgcatcggcggcttcaacacagagaaaggcaagaagaaagcgcctgcagacctctgcataggcatcctggagtctatggggctatccagtaag
accagacgcgccagagacggcgatgcacgccccacgatgcgttccattgacccggactcggtcgagttcctcatgaatatcgtggagaagcgccgcgaggccggtttat
caattcacgcacgtaaggtggaaaaaaccaccatcgaagtggatcgcgatttggatctaaatatagatatacatggtaaccctcgatccaaaacagagcacgttccgga
cgccccacaatcagtaattatccaggcactggaggctatcccggtggcggtaccggaggcgtgggcggagaacgcgttgccagcaaccgaaatggaggcggtacgcctg
tggccagtggccagcatcgcgagaacgttcgcctcgctgtacatgaccgaatttatggacctgctatcagtacgcgaaataaggctgctgaaagcgtttctaagccagc
ggcaagccgtggcgatataaccggaggggtaatgggaaaacaacgggaatgccttgtcatcgtggagggctacgacccttatgtggtcaaggtcagccacgacgcgaca
agctatagcgattataaaactggccgggaacttccattcacgatcattcggatgaacgggttctcagatagtgagggagtttattgctgcattgccagcgataagagca
tggaagaccacacagacgaagccagggcgcttctaaaagcctataaggtgaagccactatgacgtcgacaccagagttcctgaagcaactggattacgaacagctgaag
tactgtcgcgagctgtgcgataaccgcatccgggctattgaggccgaagagaaaaagatggcctgggcagttactgatggtggcattaactacggctggttccgtacag
aagactacatgaaggcggtagaatgcctggttgctacagctgcagaacggtgggaagaatcggataaggctgacccatctggacgtggttggcttgacctttcaatccg
tggaagccgcctgccggtgtccgagtatgaagctttatttgctgatggccagtgggggtgaatagtggcgcatgatctacaaaaaatgatggatggtgagctggtccgc
ttacaacttgagattaagaaagaacaggagaggcgtttagccgaaaagatgattcctgtttttggtgtgtggcaatcacctattaatcaatatgaatttgctgatccag
agcaggcaatttcttgtgccgcagggctattggataaggttcttcgtgaggtcaaggacgatctttccaaagaaggtggattacagggatggaacggcaatttgctacg
catctacgtacagcatgtgaacgagagcgattttgcagttctacagccttatttaaaggacaaataatcatgatttatcgcagaggatgggtgcctgttttgtggagga
atgagctggaaaagcggctcaaagaagaagggttcgaaaactggaagcagataagtaactttctttgtggtgaagaggattgctactcggaagcagcaaaaacgccagg
acaatacgcttaccaggtcgtcgacaataccgaatggatggaacgtcgtgatgcgacgttctggcagcgtctgaaccgtttgtggttcgttccgctgtacctgctgact
attccgttccagtggctcattcgtggccgtatggggtttgagacaacatctaaggtggctgcggtcatcaataagattaccggactccaataactgatcaggttctgat
cagtcagtaaagcaaactataaaaacacctgtgcacgtgactggtcacggggtaaaatatccacctaaattattgacgtgcgttctgctttggcggtagagttaccccg
ctgcagcaaaatctgcagccgggcctctcaaccccgaatgacaagaagcgcacaacacgcgccagcgtgtttttttgtgtgttaaatctgcgcatacctgaattatggt
ggctcagatggggccaacttcggttgggccggtttcttcttgtaccggtgttgagaaccctgtctgggctaccacccctatagagattctcaactctggtggtagcacc
ctatacaagataggaatgcataccatgttcaaattcaaatttgcagctgtcgtccgcacggacaaaaaatcccatatccaccgcctttccaccattgcatcatccgagc
gagaagctcgtcgccagttcgccagccgttttgtcctcgttctgtcagcccgtatcccggtcagcgaggtgatcgcatgaaccagattcagttaaacgcccagggcctg
ctggagacgattgaggaacgcctgctgcaggttgaagcgcttgtgggctctgctcaccgcactatctccagttatgaggccacgctgtatttgcaggaagctgcggaac
ttctgcaggtcgcgcgcgaactcacgcaggaagctcgcggctgctccctgtccctttctgagcagctgaaatccggggaggcgcaatgaaggctctctccgtcttttca
tttcaggaaagccaccctatacgggtggttcttgtcggtggtgacccgtggtttgtggcgctggatatctgtgctgcattgaatatagccaacccttctgatgcattgc
gtaagctggatcatgatgaaaaattgaccctcggtttaaccgaggcacaaaaacttgatcggatggccagggaagtaaatgtggtatcggaatctggtctctacaccat
aatcctccgctgccgcgacgcggtgaaacaggggacgacagcctggcggttccgcaagtgggtgacgaacgaggttctgccagctatccggaaaaacggcgagtatgct
tttgtcgagcctgagcctaagaacgccggtgaaccactggactggcggcagaaagaagaacttcggggcctgataaacgacatagcacaaagttttcagtaccgtaacg

catgggttagcggcgtctggatggctctccgccgcgcctgcaggaacccatcaccgaacccaataaccgtggatgatctcccggcaatcatcgccgagctgcgccggat
attaacggctgcagaaacggctctgggcaacatgcgtgtctacgaacgggagctgctgcgcgatgtagttcgtggcggtcgccggagtatgtcttgtggagagctgccg
atcaccgatattgatacggaactggagaaggtgctgccagcacattttgagctggccattgaaaagctggaaacgctatccacgaaactaacatcgccagcgattgcag
aaaaatagggttggccagcccgaatttaagccccgctttacgcggggaagtccctgcagtcgggacggcgatgtccgtagttgcagctgcagtttccgcctaatccttc
cgtcccttgcaggggatgggtagggacaggcctcaataaaaaagccaccgattaacaccggtggcttttcttttagctgtagtacgtttcccatgcggatcgcattgcg
gccatagggctatcggcggggccggaccaggcataatgctttccatcgtattcaaactctatcttgtacgttccgtccccgttatttttgcaggcttgaagacgggtt
ttagagcagttggctgatgttcttctggcccttcctcttcggttggttcatcgccgccaccctcgtcgagctcaatttcatcttcatccagctcgtcgtcttgggactc
atcatccggttcgtcgatggtctcaacggattcttcatcaggcaggacgattgcaggtgtctctatcttcagctgccactgcccgttctcgccaacgaactgccccaat
gcatcagcggcaaactccaggtaccggctaatcatcgtcgggctaaatttaaaggcccggagagtgctgttggttatttttgctgatgggtcctgctccaccagctgct
taacggtttcatggagacggacgccagcgtcacctctggcaaagcctggcatttcatcgtccagtttctgcagagccaccagcctggtgttttcatccccaacttcagg
tcgccaggttctggagaagttggccagcttgaactgcttatagtgcagctgggtgttctcatcgtcgtgtccgagaatctccatgaagaacacatcctcgtcgacgttt
ttccaccgtggatcgacgcggaagaacatctcataagcgatgcgagcgtaaatagcgcggctatctttataaacacgacggtcatcgccgaaaaatgatttaacccaag
ggttaaatgcttttgctaaaatagcatttatcctgccgttctcagaccttgtatcatcctttccatatcctttaacaacctcatcgaaatcagatgcagcagagcaaga
acgcaattctgttaataattcaacgaataatttgcttcgcataaagtataaatcgttctggttacgcttttatcttcagagcgtttttagcttgccctgagaaatta
accgtatactttcctgaaacggcaaattcaccctgaaacattatctcaatcattcttcgccctgataccgcagccagagcaaaggccaaaggtgccattccagaacgag
tgtttaaactaaataaagtcgcaggattattcaaaatatcatagatagactgcatgtatgttgggtagtcaataaccacaacattacgcttcttctcgcgcagaacatc
ggcccatcgttgctgtatagatgtacgctccgcagggcttagctgcagatggtacagaacctcatggttgaccttgagctggtgtagttcttctaacaacgcagagcct
tgttggaataacttataaagatagtcacggcgctccttccaatcatcactgtttaaatcactaagagcaaaactccaatctggatatttttttattagtcttgctattt
ttgcatcactgcctttagagcctattcttacatttgataactcttcggcaagcggcattatttctttcaatttagattgtaacgatgacatgtgctggcgaatattagc
cgtaggcatagaaagccatgaagataattcttcgctgtaaagaggatacttttccgataatttattaatatttttatcaaagctatgatgtaatttatcatcaaaccgc
tttcttgccctgctcatataggcgttaaaagtattcgcggttattcttttctgcaatcctttcccacggaactttcttttatcattaaataacgcgttcttataccgtg
cggctgcggctttaattctcttcgttttgtcgccttgtgggcggtctgaggcatcaattgcctctacctcattcacaagcgtgttgatcaactcaccgattttacctt
gctcatcccggttcccccgtttctaagtctctggatatcgtaacacaatcaaagaacaagtgacgattaattatgctgctttcgtcacactataagagaacataatagc
agaataacaaaggaacacaacaccggaatagatacatattgttctgcgcttatgttctccttatcagcacacaatagtccattatacgc

#### Annealed-oligo bridge needed to create pJazz-V2.

5’-GGTGATCCACCTGGCTAGCTAAGCCGGCTTACTTAAGTAGGCCTCCCGG-3’
3’-AGTCCACTAGGTGGACCGATCGATTCGGCCGAATGAATTCATCCGGAG-5’

#### pJazz-V2

See figure 2B. PmuI-Bsu36 fragment from humanised C9orf72 mouse BAC cloned into pJazz-OC (delta-AarI) pre-digested with XmaI, via an annealed-oligo bridge (above). pJazz backbone in lowercase. Mouse sequence in **green**. Human sequence in **blue**, selection cassette in **magenta**, non-pathogenic repeat in **red**, repeat targeting sgRNA underlined, Cas9 cleavage site indicated by **|**, AarI site mutation indicated by **gcTggtg**.

gcgtataatgggcaattgtgtgctgatatgttccgtaattgagatatgatatagtgcatatgttccgaattattgatacattaatgttgcattcctatgttgtgttctg
ttatggtgttcttagatagtgggggatgtcacatggattgatgttctttgattgtgtttcaatatagagacggaagagatagagatgcctcaaagtgtataaacagcgt
gaaaatcagcatgttcgcttaacttttgatagcaattggtatgatttattatcatttaaaatcattaagttaggttttaaatataccaacgtaagtaataccgttgcgc
ttagatgaagctaaatagcttctgtagcgcatcaaaaaattgatttgaaccttaactttgcgttttctaagtcactttgctttttttatgttgctttgctttaaaaacg
taaagtatgatgaaaaacacaaagataagggcatgtttgagaggaaacaggatgtcgttaattaatttgctgaaagactgtatcaatcgcggtcaggaaatgacgcgcg
ctattgccatcgcacagttcggcgatgacagcccggaagcgcgtcgtattactcgccgctggggtataacagaagttgcagacctgatcggcgtaacaccgcaggctat
cagagacgcggaaaaagctggtcgtctaccggctcctgattttgagatgaggggccgagtagaacgtcgcgctggctacacaattgaccaaattagccatatgcggagt
gtgtttggtaacccaaaccaaagacctgacgataaaaacccggttgtactttccgttatgtcacacaagggaggggtttataaacctcatctgcagtacaccaggcgc
aatggttagccttgcaagggcaccgcgtcctactcgttgaagggaacgacccgcaagggactgcatctatgtaccacggttacgttcctgatttgcacattcacgcaga
cgatacactgcttccatttacctgggtaaacgcgacaacgctgaatatgcgataaaaccaacatgctggccgggtctcgacataatccccagctgcctggcccttcac
cgcattgaaacagatctagcagaaagtcaaaagcctccgaccggaggcttttgacttctgtcacctaggttacgccccgccctgccactcatcgcagtactgttgtaat
tcattaagcattctgccgacatggaagccatcacaaacggcatgatgaacctgaatcgccagcggcatcagcaccttgtcgccttgcgtataatatttgcccatggtga
aaacgggggcgaagaagttgtccatattggccacgtttaaatcaaaactggtgaaactcacccagggattggctgagacgaaaaacatattctcaataaaccctttagg
gaaataggccaggttttcaccgtaacacgccacatcttgcgaatatatgtgtagaaactgccggaaatcgtcgtggtattcactccagagcgatgaaaacgtttcagtt
tgctcatggaaaacggtgtaacaagggtgaacactatcccatatcaccagctcaccgtctttcattgccatacgaaattccggatgagcattcatcaggcgggcaagaa
tgtgaataaaggccggataaacttgtgcttatttttctttacggtctttaaaaaggccgtaatatccagctgaacggtctggttataggtacattgagcaactgactg
aaatgcctcaaaatgttctttacgatgccattgggatatatcaacggtggtatatccagtgatttttttctccattttagcttccttagctcctgaaaatctcgataac
tcaaaaaatacgcccggtagtgatcttatttcattatggtgaaagttggaacctcttacgtgccgatcagattaaaacgaaaggcccagtctttcgactgagcctttcg
ttttatttgaccatgttggtatgatttaaatggtcagttaatcagttctatgtaccagcaaggtccagttgtaagcggccgcttgacttcaagtctaatggcccGTGAA
AGATGGCGTTTGTAGTGACAGCCATCCCAATTGCCCTTTCCTTCTAGGTGGAAAGTGGTGTCTAGACAGTCCAGGGAGGGTGTGCGAGGGAGGTGCGTTTTGGTTGCCT
CAGCTCGCAACTTAACTCCACAACGGTGACCAAGGACAAAAGAAGGAAACAAGACTGCAGAGATCCGCACCGGGGAGCCCTGCAGATTCTGGGTGTGTGTTTTTGTTTT
TCCCACCCTCTCTCCCCACTACTTGCTCTCACAGTACTCGCTGAGGGTGAACAAGAAAAGACCTGATAAAGATTAACCCTAGAAAGATAGTCTGCGTAAAATTGACGCA
TGCATTCTTGAAATATTGCTCTCTCTTTCTAAATAGCGCGAATCCGTCGCTGTGCATTTAGGACATCTCAGTCGCCGCTTGGAGCTCCCGTGAGGCGTGCTTGTCAATG
CGGTAAGTGTCACTGATTTTGAACTATAACGACCGCGTGAGTCAAAATGACGCATGATTATCTTTTACGTGACTTTTAAGATTTAACTCATACGATAATTATATTGTTA
TTTCATGTTCTACTTACGTGATAACTTATTATATATATATTTTCTTGTTATAGATATCGCGGCCGCTCTAGCCTCGAGGCTAGAACTAGTGGATCTCGAGCCCCAGCTG
GTTCTTTCCGCCTCAGAAGCCATAGAGCCCACCGCATCCCCAGCATGCCTGCTATTGTCTTCCCAATCCTCCCCCTTGCTGTCCTGCCCCACCCCACCCCCCAGAATAG
AATGACACCTACTCAGACAATGCGATGCAATTTCCTCATTTTATTAGGAAAGGACAGTGGGAGTGGCACCTTCCAGGGTCAAGGAAGGCACGGGGGAGGGGCAAACAAC
AGATGGCTGGCAACTAGAAGGCACAGTCGAGGCTGATCAGCGAGCTCTAGAGCTCAGAAGAACTCGTCAAGAAGGCGATAGAAGGCGATGCGCTGCGAATCGGGAGCGG
CGATACCGTAAAGCACGAGGAAGCGGTCAGCCCATTCGCCGCCAAGCTCTTCAGCAATATCACGGGTAGCCAACGCTATGTCCTGATAGCGGTCCGCCACACCCAGCCG
GCCACAGTCGATGAATCCAGAAAAGCGGCCATTTTCCACCATGATATTCGGCAAGCAGGCATCGCCATGGGTCACGACGAGATCCTCGCCGTCGGGCATGCGCGCCTTG
AGCCTGGCGAACAGTTCGGCTGGCGCGAGCCCCTGATGCTCTTCGTCCAGATCATCCTGATCGACAAGACCGGCTTCCATCCGAGTACGTGCTCGCTCGATGCGATGTT
TCGCTTGGTGGTCGAATGGGCAGGTAGCCGGATCAAGCGTATGCAGCCGCCGCATTGCATCAGCCATGATGGATACTTTCTCGGCAGGAGCAAGGTGAGATGACAGGAG
ATCCTGCCCCGGCACTTCGCCCAATAGCAGCCAGTCCCTTCCCGCTTCAGTGACAACGTCGAGCACAGCTGCGCAAGGAACGCCCGTCGTGGCCAGCCACGATAGCCGC
GCTGCCTCGTCCTGCAGTTCATTCAGGGCACCGGACAGGTCGGTCTTGACAAAAAGAACCGGGCGCCCCTGCGCTGACAGCCGGAACACGGCGGCATCAGAGCAGCCGA
TCGTCTGTTGTGCCCAGTCATAGCCGAATAGCCTCTCCACCCAAGCGGCCGGAGAACCTGCGTGCAATCCATCTTGTTCAATGGCCGATCCCATGGTTTAGTTCCTCAC
CTTGTCGTATTATACTATGCCGATATACTATGCCGATGATTAATTGTCAACAGGCTGCAGGTCGAAAGGCCCGGAGATGAGGAAGAGGAGAACAGCGCGGCAGACGTGC
GCTTTTGAAGCGTGCAGAATGCCGGGCCTCCGGAGGACCTTCGGGCGCCCGCCCCGCCCCTGAGCCCGCCCCTGAGCCCGCCCCCGGACCCACCCCTTCCCAGCCTCTG
AGCCCAGAAAGCGAAGGAGCAAAGCTGCTATTGGCCGCTGCCCCAAAGGCCTACCCGCTTCCATTGCTCAGCGGTGCTGTCCATCTGCACGAGACTAGTGAGACGTGCT
ACTTCCATTTGTCACGTCCTGCACGACGCGAGCTGCGGGGCGGGGGGGAACTTCCTGACTAGGGGAGGAGTAGAAGGTGGCGCGAAGGGGCCACCAAAGAACGGAGCCG
GTTGGCGCCTACCGGTGGATGTGGAATGTGTGCGAGGCCAGAGGCCACTTGTGTAGCGCCAAGTGCCCAGCGGGGCTGCTAAAGCGCATGCTCCAGACTGCCTTGGGAA
AAGCGCCTCCCCTACCCGGTAGAATTAATTCGATATCAAGCTTGCTCGGATGATAGGGCCCTAACGTGTTTTTGCTTTGTTACTTTATAGAAGAAATTTTGAGTTTTTG

TTTTTTTTTAATAAATAAATAAACATAAATAAATTGTTTGTTGAATTTATTATTAGTATGTAAGTGTAAATATAATAAAACTTAATATCTATTCAAATTAATAAATAAA
CCTCGATATACAGACCGATAAAACACATGCGTCAATTTTACGCATGATTATCTTTAACGTACGTCACAATATGATTATCTTTCTAGGGTTAACCAGAAGAAAACAAGGA
GGGAAACAACCGCAGCCTGTAGCAAGCTCTGGAACTCAGGAGTCGCGCGCTAGG | GGCCGGGGCCGGGGCCGGGGCGTGGTCGGGGCGGGCCCGGGGGCGGGCCCGGGG
CGGGGCTGCGGTTGCGGTGCCTGCGCCCGCGGCGGCGGAGGCGCAGGCGGTGGCGAGTGGGTGAGTGAGGAGGCGGCATCCTGGCGGGTGGCTGTTTGGGGTTCGGCTG
CCGGGAAGAGGCGCGGGTAGAAGCGGGGGCTCTCCTCAGAGCTCGACGCATTTTTACTTTCCCTCTCATTTCTCTGACCGAAGCTGGGTGTCGGGCTTTCGCCTCTAGC
GACTGGTGGAATTGCCTGCATCCGGGCCCCGGGCTTCCCGGCGGCGGCGGCGGCGGCGGCGCAGGGACAAGGGATGGGGATCTGGCCTCTTCCTTGCTTTCCCGCC
CTCAGTACCCGAGCTGTCTCCTTCCCGGGGACCCGCTGGGAGCGCTGCCGCTGCGGGCTCGAGAAAAGGGAGCCTCGGGTACTGAGAGGCCTCGCCTGGGGGAAGGCCG
GAGGGTGGGCGGCGCGCGGCTTCTGCGGACCAAGTCGGGGTTCGCTAGGAACCCGAGACGGTCCCTGCCGGCGAGGAGATCATGCGGGATGAGATGGGGGTGTGGAGAC
GCCTGCACAATTTCAGCCCAAGCTTCTAGAGAGTGGTGATGACTTGCATATGAGGGCAGCAATGCAAGTCGGTGTGCTCCCCATTCTGTGGGACATGACCTGGTTGCTT
CACAGCTCCGAGATGACACAGACTTGCTTAAAGGAAGTGACTATTGTGACTTGGGCATCACTTGACTGATGGTAATCAGTTGTCTAAAGAAGTGCACAGATTACATGTC
CGTGTGCTCATTGGGTCTATCTGGCCGCGTTGAACACCACCAGGCTTTGTATTCAGAAACAGGAGGGAGGTCCTGCACTTTCCCAGGAGGGGTGGCCCTTTCAGATGCA
ATCGAGATTGTTAGGCTCTGGGAGAGTAGTTGCCTGGTTGTGGCAGTTGGTAAATTTCTATTCAAACAGTTGCCATGCACCAGTTGTTCACAACAAGGGTACGTAATCT
GTCTGGCATTACTTCTACTTTTGTACAAAGGATCAAAAAAAAAAAAAGATACTGTTAAGATATGATTTTTCTCAGACTTTGGGAAACTTTTAACATAATCTGTGAATATC
ACAGAAACAAGACTATCATATAGGGGATATTAATAACCTGGAGTCAGAATACTTGAAATACGGTGTCATTTGACACGGGCATTGTTGTCACCACCTCTGCCAAGGCCTG
CCACTTTAGGAAAACCCTGAATCAGTTGGAAACTGCTACATGCTGATAGTACATCTGAAACAAGAACGAGAGTAATTACCACATTCCAGATTGTTCACTAAGCCAGCAT
TTACCTGCTCCAGGAAAAAATTACAAGCACCTTATGAAGTTGATAAAATATTTTGTTTGGCTATGTTGGCACTCCACAATTTGCTTTCAGAGAAACAAAGTAAACCAAG
GAGGACTTCTGTTTTTCAAGTCTGCCCTCGGGTTCTATTCTACGTTAATTAGATAGTTCCCAGGAGGACTAGGTTAGCCTACCTATTGTCTGAGAAACTTGGAACTGTG
AGAAATGGCCAGATAGTGATATGAACTTCACCTTCCAGTCTTCCCTGATGTTGAAGATTGAGAAAGTGTTGTGAACTTTCTGGTACTGTAAACAGTTCACTGTCCTTGA
AGTGGTCCTGGGCAGCTCCTGTTGTGGAAAGTGGACGGTTTAGGATCCTGCTTCTCTTTGGGCTGGGAGAAAATAAACAGCATGGTTACAAGTATTGAGAGCCAGGTTG
GAGAAGGTGGCTTACACCTGTAATGCCAGAGCTTTGGGAGGCGGAGGCAAGAGGATCACTTGAAGCCAGGAGTTCAAGCTCAACCTGGGCAACGTAGACCCTGTCTCTA
CAAAAAATTAAAAACTTAGCCGGGCGTGGTGATGTGCACCTGTAGTCCTAGCTACTTGGGAGGCTGAGGCAGGAGGGTCATTTGAGCCCAAGAGTTTGAAGTTACCGAG
AGCTATGATCCTGCCAGTGCATTCCAGCCTGGATGACAAAACGAGACCCTGTCTCTAAAAAACAAGAAGTGAGGGCTTTATGATTGTAGAATTTTCACTACAATAGCAG
TGGACCAACCACCTTTCTAAATACCAATCAGGGAAGAGATGGTTGATTTTTTAACAGACGTTTAAAGAAAAAGCAAAACCTCAAACTTAGCACTCTACTAACAGTTTTA
GCAGATGTTAATTAATGTAATCATGTCTGCATGTATGGGATTATTTCCAGAAAGTGTATTGGGAAACCTCTCATGAACCCTGTGAGCAAGCCACCGTCTCACTCAATTT
GAATCTTGGCTTCCCTCAAAAGACTGGCTAATGTTTGGTAACTCTCTGGAGTAGACAGCACTACATGTACGTAAGATAGGTACATAAACAACTATTGGTTTTGAGCTGA
TTTTTTTCAGCTGCATTTGCATGTATGGATTTTTCTCACCAAAGACGATGACTTCAAGTATTAGTAAAATAATTGTACAGCTCTCCTGATTATACTTCTCTGTGACATT
TCATTTCCCAGGCTATTTCTTTTGGTAGGATTTAAAACTAAGCAATTCAGTATGATCTTTGTCCTTCATTTTCTTTCTTATTCTTTTTGTTTGTTTGTTTGTTTGTTTT
TTTCTTGAGGCAGAGTCTCTCTCTGTCGCCCAGGCTGGAGTGCAGTGGCGCCATCTCAGCTCATTGCAACCTCTGCCACCTCCGGGTTCAAGAGATTCTCCTGCCTCAG
CCTCCCGAGTAGCTGGGATTACAGGTGTCCACCACCACACCCGGCTAATTTTTTGTATTTTTAGTAGAGGTGGGGTTTCACCATGTTGGCCAGGCTGGTCTTGAGCTCC
TGACCTCAGGTGATCCACCTGGCTAGCTAAGCCGGCTTACTTAAGTAGGCCTCccgggccattgggccctagacaagttgtcgcggccgcactagtgactccagcgtaa
ctggactggccacagttaggccgcaaatgtaatcacactggctcaccttcgggtgggcctttctgcgtctagaggatcccctgttatgcccttccttacccatgcgctg
agaggtgaaatcactctttaatttgtacgcggtacggacctctatatcgctttgtcgtagaggtggaatctccccagcttgcagccattgatagactgcgccgggtgtc
acgcccacgccaaccgctgctttttccacatcgccgaaatggcgcacaagttcttcgggtttcatggggattattataatcaataagtgaaaattaaagctaggtataa
tttataaaaattatagccagctataaagagatcatttatgattaagggtatgaaaacacgaggcgaacgactgaaagcacgccgtttagagctgaaactgacgctgaag
caagtggcagaagctgtaggaatctctcttccgggcgtccaaaacttagaacgtggcgacgtgatgccgtcgctggagatcgggctatcgctggcaaaatgcctgcgta
agcccgtgcaatggatactgtatggcactgaatctgatccagaccgcgttcctgttattggtacgacagagaccggaccggatagagactggcagcctggagaacctgc
caacacagagcgattcctgccgttcgtgagtcaacggaacaccgtttacgcactgacggtcgggaaccagattcagcgaaactaccagccgggagacgttatcctggtc
gattcttcacttacgcttgtaccgggtgaggatgtactggtttgtgataataacggcgagatcacaattcaacgattagcccgttatgacgagcagcattactaccttg
atagtgctaactctcaacgggttatccatgataaaagtgatcttcaatttgtgcaccaagtagtcggtacgatcaaatcgttcatggttgagggtagatgatacaataa
cagggtttattgctgactataattctggtttaatccgatctatactttgtcggggttgaaccagaccgtagcagccgaaaaaagacgaaaaaaacccgagtcggcaaa
ctcgggcctttttcaggaagtagccacgataacgcagatacgtccttcagaaagatagtgcgtttattgtggctgctcctgcggatttttcaacccgaaaaaatgcta
attcgcatggaaaggctaaaaatgaccttacaagaattctacgcggagcgctttggcagcgatccgttttcactgcttgacgcggcacgtgatgagctgactgagctg
gcgaagatggccggtatcaactggcctgcgtgtgctgataagattcagctcaatccgcgcggtggtgtagagcgctataccacctacaataattcctccccggaagctc
tgcagaagagccttaaaggccgtgtggaaatctattcacgccttgagcagagcaaggatggcatcagttatcctttcgttaacttcgttgaaaaagcgcatgacgctgg
ctcctggagtggcttttcattcctgttttcggagtatcgccgtgaacagcaacggaatggtgcgaccgtggtcgctcagccggaagaagaacgtgcacgcatggagcgc
caggctgaagcacgccgcctccgagctgaacaacaacgcgttaatgatttaaaaaacaatcagatggagcaggaacgcctgcttggatggttggctttccatcgcgcct
gggaacattcgccagctgaagatggctcctggccttatgcggtaaaaaagggaattcgtgacgtatttggcgcttgtgatatacgtcgcgtgaccagtcacgacagtgc
gaaatggagccgtggaccaacaacttacatggcgatacctctgtcccacctggacggaagaaaagacgggcggattgttggctggcaacgtatcgatctgaatggtggc
aagttccagaccagcgccatcactaacggtgatttcgtcggtgcatgctttgttattggcgacctgcaggggggcgcaaaaaattgcgactgcagagggtttcgccaccg
gggcatccatctggctggcgaccagaaacgacccgaaaaaacgctttgacgcggtagtcattgcggtttccgctaacaacatgattcacgttgtcgagcagctggtgaa
catgtacccggctgcgcaaatcacttgcgctctcgataacgaccgcaaatcctcagctgaaggaaaaggcaacacaggcctccgcaccggattcgacatcatggagaag
ttttccggcgtcaaatgtgtttacccaacttttgaggatgaccctgagctggaatgcagcgattttaacgacctgcacagcctgagagggctgaaggaagtcgctcgcc
agttaacgagaaatcatctgagccgtgcaaccgatctgttgtcgatcacgctgaataagctgcgtactctcccgcgtctgaacagacgcactttgccaaagaactgct
tcgcgccgttgatattggcatgctgacatgcccggtaccaaacagcccgaaagaacttatgcgccttttcagcagcacgctgcgggatatgggtatcgcagaaatttat
aacggtaccgttaaagatcacattacgcgccggttgaatcgcaaatgccgtgccgcgcaaacatcacgttcgttcagtgaacgcatcaccaacccgaaccttcgcccgt
cacacatcacttacaaacggtttgaaacctccaggatgaccgatgaggtgatgacatacgccgcacagctgcagggcatcgttattgtccgcgccgggatggggtcggg
taaatcgacaggcctcctgcgtccactgatgctgcagtccacgcgtggcgtttccgtcgcgcaccgcgtatccctgataggcggcctgcatgaaatgatgaccgaaggg
aaaggcgcgaaagccgacattctgcattatcaggatcccggctatcaggaaatggcgccatatgccaacaagctgactatttgcatcaactccatacttaaaggctgct
ggcaaccgctgatgcgccagcatgacttcttcggcttcgatgaagcaacacagggcctgcgtgccattctggccgggcgtgcgatggaaaacccggtaggcgtattcaa
cacgcttatcgacgcgctggcgcgtactgaagagcatgccattatggtggatgccgatgccaacgatctgcttgttgacctggctgaactggcgatgaagcgacgcgag
gagctgggcctacctgcctggctgcaaattcacgtgattgaactcccggtcgacgttcgcaaccgcgaaaccaacaagcctatccgcgtattttataccgagaaaaatc
ggatcatgaccgaggttatcgctgcagtgcaacgcggcgaacgaatcatgctggccaccgacagttcgacgttcgccgaagacgttaccatgcagctgagactacagtt
ccctgacaaaaagtttctctgcgttaaccagaaaaacaaacaggagaaggaagtcgacgatttcaccaaccagcctaaagtgatggtgaaaaaatatgacggcctcatc
tacagcccgtcgatatcttcaggtgtatcgattgaggagaaacacttccaccgccatttcggcatgttctgcggcgaagtggtccccagcgacgccatccagatgctgc
gccgcgaccgtacagctcaggaatacattatcggtttcgacaagcttcgcggtaaacgtgaaaccgatccggaaaaaatcaaacgcgcctacgcgcaggccttgctcga
aacggctggccactccgggctgctgacagacgttgtctttgacggtgaccgaatttcactcggcgtggctaactcttcattcatgcagctcaaaattaaagccgccgcg
ctcgaggcatcggccagaaatgattatgccagcaacatgatttgcatcatgcatgatgatggctatcaggtcgcgcctatggccaccgacgcgcttgcaaacagcatcg
gtaaggatttacgtaaagaagcccgtgaactggtctttgagcagcttatggagcgccacctgagtgtcgatactcctgaccaggccgaacacgatgagttgataaaaaa
acgcaccctgtccctggatgagcaggcccagctggtccgctgggacatcgaaaaggagctgcagctggatgtcgacgaggtggcgttaaaattctatttcgacggcggc
ctgaaaaaagtgcgcctgtttgaaaccatgcagctggatgagataaccgcccgtcgccttgaccgtgaagaagcgttgattcactttacctacgcttatcgcgttgccg
gaagatggcagcagttcgtcactacggccatgacgcgggaacaggccgacgcagagttccaggctaaattcccggccataaccgattaccgagtgaaatcgacccctgc
tgttgagataggcatgcgtggtttctacaccctcaaatcggcgacgctgcagcagtacttccgcgactgcggtatcgatccaaaaaccctggaaggtgaagcagacatg
gacgccctcaaacgcgccagagataacctgctcacgccggaacggcgcgatctgctaaacaacgttttacgcatcggcggcttcaacacagagaaaggcaagaagaaag
cgcctgcagacctctgcataggcatcctggagtctatggggctatccagtaagaccagacgcgccagagacggcgatgcacgccccacgatgcgttccattgacccgga
ctcggtcgagttcctcatgaatatcgtggagaagcgccgcgaggccggtttatcaattcacgcacgtaaggtggaaaaaaccaccatcgaagtggatcgcgatttggat
ctaaatatagatatacatggtaaccctcgatccaaaacagagcacgttccggacgccccacaatcagtaattatccaggcactggaggctatcccggtggcggtaccgg
aggcgtgggcggagaacgcgttgccagcaaccgaaatggaggcggtacgcctgtggccagtggccagcatcgcgagaacgttcgcctcgctgtacatgaccgaatttat
ggacctgctatcagtacgcgaaataaggctgctgaaagcgtttctaagccagcggcaagccgtggcgatataaccggaggggtaatgggaaaacaacggaatgccttg

tcatcgtggagggctacgacccttatgtggtcaaggtcagccacgacgcgacaagctatagcgattataaaactggccgggaacttccattcacgatcattcggatgaa
cggcttctcagatagtgagggagtttattgctgcattgccagcgataagagcatggaagaccacacagacgaagccagggcgcttctaaaagcctataggtgaagcca
ctatgacgtcgacaccagagttcctgaagcaactggattacgaacagctgaagtactgtcgcgagctgtgcgataaccgcatccgggctattgaggccgaagagaaaaa
gatggcctgggcagttactgatggtggcattaactacggctggttccgtacagaagactacatgaaggcggtagaatgcctggttgctacagctgcagaacggtgggaa
gaatcggataaggctgacccatctggacgtggttggcttgacctttcaatccgtggaagccgcctgccggtgtccgagtatgaagctttatttgctgatggccagtggg
ggtgaatagtggcgcatgatctacaaaaaatgatggatggtgagctggtccgcttacaacttgagattaagaaagaacaggagaggcgtttagccgaaaagatgattcc
tgtttttggtgtgtggcaatcacctattaatcaatatgaatttgctgatccagagcaggcaatttcttgtgccgcagggctattggataaggttcttcgtgaggtcaag
gacgatctttccaaagaaggtggattacagggatggaacggcaatttgctacgcatctacgtacagcatgtgaacgagagcgattttgcagttctacagccttatttaa
aggacaaataatcatgatttatcgcagaggatgggtgcctgttttgtggaggaatgagctggaaaagcggctcaaagaagaagggttcgaaaactggaagcagataagt
aactttctttgtggtgaagaggattgctactcggaagcagcaaaaacgccaggacaatacgcttaccaggtcgtcgacaataccgaatggatggaacgtcgtgatgcga
cgttctggcagcgtctgaaccgtttgtggttcgttccgctgtacctgctgactattccgttccagtggctcattcgtggccgtatggggtttgagacaacatctaaggt
ggctgcggtcatcaataagattaccggactccaataactgatcaggttctgatcagtcagtaaagcaaactataaaaacacctgtgcacgtgactggtcacggggtaaa
atatccacctaaattattgacgtgcgttctgctttggcggtagagttaccccgctgcagcaaaatctgcagccgggcctctcaaccccgaatgacaagaagcgcacaac
acgcgccagcgtgttttttttgtgtgttaaatctgcgcatacctgaattatggtggctcagatggggccaacttcggttgggccggtttcttcttgtaccggtgttgaga
accctgtctgggctaccaccccctatagagattctcaactctggtggtagcaccctatacaagataggaatgcataccatgttcaaattcaaatttgcagctgtcgtccg
cacggacaaaaaaatcccatatccaccgcctttccaccattgcatcatccgagcgagaagctcgtcgccagttcgccagccgttttgtcctcgttctgtcagcccgtatc
ccggtcagcgaggtgatcgcatgaaccagattcagttaaacgcccagggcctgctggagacgattgaggaacgcctgctgcaggttgaagcgcttgtgggctctgctca
ccgcactatctccagttatgaggccacgctgtatttgcaggaagctgcggaacttctgcaggtcgcgcgcgaactcacgcaggaagctcgcggctgctccctgtccctt
tctgagcagctgaaatcggggaggcgcaatgaaggctctctccgtctttcatttcaggaaagccaccctatacgggtggttcttgtcggtggtgacccgtggtttgt
ggcgctggatatctgtgctgcattgaatatagccaaccccttctgatgcattgcgtaagctggatcatgatgaaaaattgaccctcggtttaaccgaggcacaaaaactt
gatcggatggccagggaagtaaatgtggtatcggaatctggtctctacaccataatcctccgctgccgcgacgcggtgaaacaggggacgacagcctggcggttccgca
agtgggtgacgaacgaggttctgccagctatccggaaaaacggcgagtatgcttttgtcgagcctgagcctaagaacgccggtgaaccactggactggcggcagaaaga
agaacttcggggcctgataaacgacatagcacaaagttttcagtaccgtaacgcatgggttagcggcgtctggatggctctccgccgcgcctgcaggaacccatcaccg
aacccaataaccgtggatgatctcccggcaatcatcgccgagctgcgccggatattaacggctgcagaaacggctctgggcaacatgcgtgtctacgaacgggagctgc
tgcgcgatgtagttcgtggcggtcgccggagtatgtcttgtggagagctgccgatcaccgatattgatacggaactggagaaggtgctgccagcacattttgagctggc
cattgaaaagctggaaacgctatccacgaactaacatcgccagcgattgcagaaaaatagggttggccagcccgaatttaagccccgctttacgcggggaagtccctg
cagtcgggacggcgatgtccgtagttgcagctgcagtttccgcctaatccttccgtcccttgcaggggatgggtagggacaggcctcaataaaaagccaccgattaac
accggtggcttttctttagctgtagtacgtttcccatgcggatcgcattgcggccatagggctatcggcggggccggaccaggcataatgctttccatcgtattcaaa
ctctatcttgtacgttccgtccccgttatttttgcaggcttgaagacgggttttagagcagttggctgatgttcttctggcccttcctcttcggttggttcatcgccg
ccaccctcgtcgagctcaatttcatcttcatccagctcgtcgtcttgggactcatcatccggttcgtcgatggtctcaacggattcttcatcaggcaggacgatt**gcTg**
**gtg**tctctatcttcagctgccactgcccgttctcgccaacgaactgccccaatgcatcagcggcaaactccaggtaccggctaatcatcgtcgggctaaatttaaaggc
ccggagagtgctgttggttatttttgctgatggtcctgctccaccagctgcttaacggtttcatggagacggacgccagcgtcacctctggcaaagcctggcatttca
tcgtccagtttctgcagagccaccagcctggtgttttcatccccaacttcaggtcgccaggttctggagaagttggccagcttgaactgcttatagtgcagctgggtgt
tctcatcgtcgtgtccgagaatctccatgaagaacacatcctcgtcgacgtttttccaccgtggatcgacgcggaagaacatctcataagcgatgcgagcgtaaatagc
gcggctatctttataaacacgacggtcatcgccgaaaaatgatttaacccaagggttaaatgcttttgctaaaatagcatttatcctgccgttctcagaccttgtatca
tcctttccatatcctttaacaacctcatcgaaatcagatgcagcagagcaagaacgcaattctgttaataattcaacgaataattttgcttcgcataaagtataaatcg
ttctggttacgcttttatcttcagagcgtttttttagcttgccctgagaaattaaccgtatactttcctgaaacggcaaattcaccctgaaacattatctcaatcattct
tcgccctgataccgcagccagagcaaaggccaaaggtgccattccagaacgagtgtttaaactaaataaagtcgcaggattattcaaaatatcatagatagactgcatg
tatgttgggtagtcaataaccacaacattacgcttcttctcgcgcagaacatcggcccatcgttgctgtatagatgtacgctccgcagggcttagctgcagatggtaca
gaacctcatggttgaccttgagctggtgtagttcttctaacaacgcagagccttgttggaataacttataaagatagtcacggcgctccttccaatcatcactgtttaa
atcactaagagcaaaactccaatctggatatttttttattagtcttgctatttttgcatcactgccttttagagcctattcttacatttgataactcttcggcaagcggc
attatttctttcaatttagattgtaacgatgacatgtgctggcgaatattagccgtaggcatagaaagccatgaagataattcttcgctgtaaagaggatacttttccg
ataatttattaatatttttatcaaagctatgatgtaatttatcatcaaaccgctttcttgccctgctcatataggcgttaaaagtattcgcggttattcttttctgcaa
tcctttcccacggaactttcttttatcattaaataacgcgttcttataccgtgcggctgcggctttaattctcttcgttttgtcgccttgtgggcggtctgaggcatca
attgcctctacctcattcacaagcgtgttgatcaactcaccgattttaccttgctcatcccggttcccccgtttctaagtctctggatatcgtaacacaatcaaagaa
caagtgacgattaattatgctgctttcgtcacactataagagaacataatagcagaataacaaaggaacacaacaccggaatagatacatattgttctgcgcttatgtt
ctcccttatcagcacacaatagtccattatacgc

#### lacZ landing pad

See Figure 2B.

ggccggatcggcaggtgatacttaagtaagccggcttagctagcgggacaggtttcccgactggaaagcgggcagtgagcgcaacgcaattaatgtgagttagctcact
cattaggcaccccaggctttacactttatgcttccggctcgtatgttgtgtggaattgtgagcggacaacaatttcacacaggaaacagctatgaccatgattacgcca
agctatttaggtgagactatagaatactcaagcttgcatgcgatacgtatcgttaacgatggatccgacgcacgtgcgaattcgccctatagtgagtcgtattacaatt
cactggccgtcgttttacaacgtcgtgactgggaaaaccctggcgtcacccaacttaatcgccttgcagcacatccccctttcgccagctggcgtaatagcgaagaggc
ccgcaccgatcgcccttcccaacagttgcgcagcctgaatggcgaatcttaagtaggcctcccccacctgcat

#### pJazz-V3

See Figure 2B. Insertion of landing pad into pJazz-V2 at Cas9 digestion site. pJazz backbone in lowercase. Mouse sequence in **green**. Human sequence in **blue**, selection cassette in **magenta**, non-pathogenic repeat in **red**, lacZ landing pad highlighted in grey, AarI sites underlined, AarI cut sites indicated by |. Backbone AarI site mutation indicated by **gcTggtg**.

gcgtataatgggcaattgtgtgctgatatgttccgtaattgagatatgatatagtgcatatgttccgaattattgatacattaatgttgcattcctatgttgtgttctg
ttatggtgttcttagatagtggggatgtcacatggattgatgttctttgattgtgtttcaatatagagacggaagagatagagatgcctcaaagtgtataaacagcgt
gaaaatcagcatgttcgcttaacttttgatagcaattggtatgatttattatcatttaaaatcattaagttaggttttaaatataccaacgtaagtaataccgttgcgc
ttagatgaagctaaatagcttctgtagcgcatcaaaaaaattgatttgaaccttaactttgcgttttctaagtcactttgcttttttttatgttgctttgctttaaaaacg
taaagtatgatgaaaaacacaaagataagggcatgtttgagaggaaacaggatgtcgttaattaatttgctgaaagactgtatcaatcgcggtcaggaaatgacgcgcg
ctattgccatcgcacagttcggcgatgacagcccggaagcgcgtcgtattactcgccgctggggtataacagaagttgcagacctgatcggcgtaacaccgcaggctat
cagagacgcggaaaaagctggtcgtctaccggctcctgattttgagatgaggggccgagtagaacgtcgcgctggctacacaattgaccaaattagccatatgcggagt
gtgtttggtaacccaaaccaaagacctgacgataaaaacccggttgtactttccgttatgtcacacaagggaggggtttataaaacctcatctgcagtacaccaggcgc
aatggttagccttgcaagggcaccgcgtcctactcgttgaagggaacgacccgcaagggactgcatctatgtaccacggttacgttcctgatttgcacattcacgcaga
cgatacactgcttccatttacctgggtaaacgcgacaacgctgaatatgcgataaaaccaacatgctggccgggtctcgacataatccccagctgcctggcccttcac
cgcattgaaacagatctagcagaaagtcaaaagcctccgaccggaggcttttgacttctgtcacctaggttacgccccgccctgccactcatcgcagtactgttgtaat
tcattaagcattctgccgacatggaagccatcacaaacggcatgatgaacctgaatcgccagcggcatcagcaccttgtcgccttgcgtataatatttgcccatggtga
aaacgggggcgaagaagttgtccatattggccacgtttaaatcaaaactggtgaaactcacccagggattggctgagacgaaaaacatattctcaataaaccctttagg
gaaataggccaggttttcaccgtaacacgccacatcttgcgaatatatgtgtagaaactgccggaaatcgtcgtggtattcactccagagcgatgaaaacgtttcagtt
tgctcatggaaaacggtgtaacaagggtgaacactatcccatatcaccagctcaccgtctttcattgccatacgaaattccggatgagcattcatcaggcgggcaagaa

tgtgaataaaggccggataaaacttgtgcttatttttctttacggtctttaaaaaggccgtaatatccagctgaacggtctggttataggtacattgagcaactgactg
aaatgcctcaaaatgttctttacgatgccattgggatatatcaacggtggtatatccagtgatttttttctccattttagcttccttagctcctgaaaatctcgataac
tcaaaaaatacgcccggtagtgatcttatttcattatggtgaaagttggaacctcttacgtgccgatcagattaaaacgaaaggcccagtctttcgactgagcctttcg
ttttatttgaccatgttggtatgatttaaatggtcagttaatcagttctatgtaccagcaaggtccagttgtaagcggccgcttgacttcaagtctaatggcccGTGAA
AGATGGCGTTTGTAGTGACAGCCATCCCAATTGCCCTTTCCTTCTAGGTGGAAAGTGGTGTCTAGACAGTCCAGGGAGGGTGTGCGAGGGAGGTGCGTTTTGGTTGCCT
CAGCTCGCAACTTAACTCCACAACGGTGACCAAGGACAAAAGAAGGAAACAAGACTGCAGAGATCCGCACCGGGGAGCCCTGCAGATTCTGGGTGTGTGTTTTTGTTTT
TCCCACCCTCTCTCCCCACTACTTGCTCTCACAGTACTCGCTGAGGGTGAACAAGAAAAGACCTGATAAAGATTAACCCTAGAAAGATAGTCTGCGTAAAATTGACGCA
TGCATTCTTGAAATATTGCTCTCTCTTTCTAAATAGCGCGAATCCGTCGCTGTGCATTTAGGACATCTCAGTCGCCGCTTGGAGCTCCCGTGAGGCGTGCTTGTCAATG
CGGTAAGTGTCACTGATTTTGAACTATAACGACCGCGTGAGTCAAAATGACGCATGATTATCTTTTACGTGACTTTTAAGATTTAACTCATACGATAATTATATTGTTA
TTTCATGTTCTACTTACGTGATAACTTATTATATATATATATTTTCTTGTTATAGATATCGCGGCCGCTCTAGCCTCGAGGCTAGAACTAGTGGATCTCGAGCCCCAGCTG
GTTCTTTCCGCCTCAGAAGCCATAGAGCCCACCGCATCCCCAGCATGCCTGCTATTGTCTTCCCAATCCTCCCCCTTGCTGTCCTGCCCCACCCCACCCCCCAGAATAG
AATGACACCTACTCAGACAATGCGATGCAATTTCCTCATTTTATTAGGAAAGGACAGTGGGAGTGGCACCTTCCAGGGTCAAGGAAGGCACGGGGGAGGGGCAAACAAC
AGATGGCTGGCAACTAGAAGGCACAGTCGAGGCTGATCAGCGAGCTCTAGAGCTCAGAAGAACTCGTCAAGAAGGCGATAGAAGGCGATGCGCTGCGAATCGGGAGCGG
CGATACCGTAAAGCACGAGGAAGCGGTCAGCCCATTCGCCGCCAAGCTCTTCAGCAATATCACGGGTAGCCAACGCTATGTCCTGATAGCGGTCCGCCACACCCAGCCG
GCCACAGTCGATGAATCCAGAAAAGCGGCCATTTTCCACCATGATATTCGGCAAGCAGGCATCGCCATGGGTCACGACGAGATCCTCGCCGTCGGGCATGCGCGCCTTG
AGCCTGGCGAACAGTTCGGCTGGCGCGAGCCCCTGATGCTCTTCGTCCAGATCATCCTGATCGACAAGACCGGCTTCCATCCGAGTACGTGCTCGCTCGATGCGATGTT
TCGCTTGGTGGTCGAATGGGCAGGTAGCCGGATCAAGCGTATGCAGCCGCCGCATTGCATCAGCCATGATGGATACTTTCTCGGCAGGAGCAAGGTGAGATGACAGGAG
ATCCTGCCCCGGCACTTCGCCCAATAGCAGCCAGTCCCTTCCCGCTTCAGTGACAACGTCGAGCACAGCTGCGCAAGGAACGCCCGTCGTGGCCAGCCACGATAGCCGC
GCTGCCTCGTCCTGCAGTTCATTCAGGGCACCGGACAGGTCGGTCTTGACAAAAAGAACCGGGCGCCCCTGCGCTGACAGCCGGAACACGGCGGCATCAGAGCAGCCGA
TCGTCTGTTGTGCCCAGTCATAGCCGAATAGCCTCTCCACCCAAGCGGCCGGAGAACCTGCGTGCAATCCATCTTGTTCAATGGCCGATCCCATGGTTTAGTTCCTCAC
CTTGTCGTATTATACTATGCCGATATACTATGCCGATGATTAATTGTCAACAGGCTGCAGGTCGAAAGGCCCGGAGATGAGGAAGAGGAGAACAGCGCGGCAGACGTGC
GCTTTTGAAGCGTGCAGAATGCCGGGCCTCCGGAGGACCTTCGGGCGCCCGCCCCGCCCCTGAGCCCGCCCCTGAGCCCGCCCCCGGACCCACCCCTTCCCAGCCTCTG
AGCCCAGAAAGCGAAGGAGCAAAGCTGCTATTGGCCGCTGCCCCAAAGGCCTACCCGCTTCCATTGCTCAGCGGTGCTGTCCATCTGCACGAGACTAGTGAGACGTGCT
ACTTCCATTTGTCACGTCCTGCACGACGCGAGCTGCGGGGCGGGGGGAACTTCCTGACTAGGGGAGGAGTAGAAGGTGGCGCGAAGGGGCCACCAAAGAACGGAGCCG
GTTGGCGCCTACCGGTGGATGTGGAATGTGTGCGAGGCCAGAGGCCACTTGTGTAGCGCCAAGTGCCCAGCGGGGCTGCTAAAGCGCATGCTCCAGACTGCCTTGGGAA
AAGCGCCTCCCCTACCCGGTAGAATTAATTCGATATCAAGCTTGCTCGGATGATAGGGCCCTAACGTGTTTTTGCTTTGTTACTTTATAGAAGAAATTTTGAGTTTTTG
TTTTTTTTTAATAAATAAATAAACATAAATAAATTGTTTGTTGAATTTATTATTAGTATGTAAGTGTAAATATAATAAAACTTAATATCTATTCAAATTAATAAATAAA
CCTCGATATACAGACCGATAAAACACATGCGTCAATTTTACGCATGATTATCTTTAACGTACGTCACAATATGATTATCTTTCTAGGGTTAACCAGAAGAAAACAAGGA
GGGAAACAACCGCAGCCTGTAGCAAGCTCTGGAACTCAGGAGTCGCGCGCTAGGgg|ccggatcggcaggtgatacttaagtaagccggcttagctagcgggacaggtt
tcccgactggaaagcgggcagtgagcgcaacgcaattaatgtgagttagctcactcattaggcaccccaggctttacactttatgcttccggctcgtatgttgtgtgga
attgtgagcggacaacaatttcacacaggaaacagctatgaccatgattacgccaagctatttaggtgagactatagaatactcaagcttgcatgcgatacgtatcgtt
aacgatggatccgacgcacgtgcgaattcgccctatagtgagtcgtattacaattcactggccgtcgttttacaacgtcgtgactgggaaaaccctggcgtcacccaac
ttaatcgccttgcagcacatccccctttcgccagctggcgtaatagcgaagaggcccgcaccgatcgcccttcccaacagttgcgcagctgaatggcgaatcttaagta
ggcctccccacctgcatgg|CCGGGGCCGGGGCCGGGGCGTGGTCGGGGCGGCCCGGGGGCGGCCCGGGGCGGGGCTGCGGTTGCGGTGCCTGCGCCCGCGGCGGCG
GAGGCGCAGGCGGTGGCGAGTGGGTGAGTGAGGAGGCGGCATCCTGGCGGGTGGCTGTTTGGGGTTCGGCTGCCGGGAAGAGGCGCGGGTAGAAGCGGGGGCTCTCCTC
AGAGCTCGACGCATTTTTACTTTCCCTCTCATTTCTCTGACCGAAGCTGGGTGTCGGGCTTTCGCCTCTAGCGACTGGTGGAATTGCCTGCATCCGGGCCCCGGGCTTC
CCGGCGGCGGCGGCGGCGGCGGCGGCGCAGGGACAAGGGATGGGGATCTGGCCTCTTCCTTGCTTTCCCGCCCTCAGTACCCGAGCTGTCTCCTTCCCGGGGACCCGCT
GGGAGCGCTGCCGCTGCGGGCTCGAGAAAAGGGAGCCTCGGGTACTGAGAGGCCTCGCCTGGGGGAAGGCCGGAGGGTGGGCGGCGCGCGGCTTCTGCGGACCAAGTCG
GGGTTCGCTAGGAACCCGAGACGGTCCCTGCCGGCGAGGAGATCATGCGGGATGAGATGGGGGTGTGGAGACGCCTGCACAATTTCAGCCCAAGCTTCTAGAGAGTGGT
GATGACTTGCATATGAGGGCAGCAATGCAAGTCGGTGTGCTCCCCATTCTGTGGGACATGACCTGGTTGCTTCACAGCTCCGAGATGACACAGACTTGCTTAAAGGAAG
TGACTATTGTGACTTGGGCATCACTTGACTGATGGTAATCAGTTGTCTAAAGAAGTGCACAGATTACATGTCCGTGTGCTCATTGGGTCTATCTGGCCGCGTTGAACAC
CACCAGGCTTTGTATTCAGAAACAGGAGGGAGGTCCTGCACTTTCCCAGGAGGGGTGGCCCTTTCAGATGCAATCGAGATTGTTAGGCTCTGGGAGAGTAGTTGCCTGG
TTGTGGCAGTTGGTAAATTTCTATTCAAACAGTTGCCATGCACCAGTTGTTCACAACAAGGGTACGTAATCTGTCTGGCATTACTTCTACTTTTGTACAAAGGATCAAA
AAAAAAAAAGATACTGTTAAGATATGATTTTTCTCAGACTTTGGGAAACTTTTAACATAATCTGTGAATATCACAGAAACAAGACTATCATATAGGGGATATTAATAAC
CTGGAGTCAGAATACTTGAAATACGGTGTCATTTGACACGGGCATTGTTGTCACCACCTCTGCCAAGGCCTGCCACTTTAGGAAAACCCTGAATCAGTTGGAAACTGCT
ACATGCTGATAGTACATCTGAAACAAGAACGAGAGTAATTACCACATTCCAGATTGTTCACTAAGCCAGCATTTACCTGCTCCAGGAAAAAATTACAAGCACCTTATGA
AGTTGATAAAATATTTTGTTTGGCTATGTTGGCACTCCACAATTTGCTTTCAGAGAAACAAAGTAAACCAAGGAGGACTTCTGTTTTTCAAGTCTGCCCTCGGGTTCTA
TTCTACGTTAATTAGATAGTTCCCAGGAGGACTAGGTTAGCCTACCTATTGTCTGAGAAACTTGGAACTGTGAGAAATGGCCAGATAGTGATATGAACTTCACCTTCCA
GTCTTCCCTGATGTTGAAGATTGAGAAAGTGTTGTGAACTTTCTGGTACTGTAAACAGTTCACTGTCCTTGAAGTGGTCCTGGGCAGCTCCTGTTGTGGAAAGTGGACG
GTTTAGGATCCTGCTTCTCTTTGGGCTGGGAGAAAATAAACAGCATGGTTACAAGTATTGAGAGCCAGGTTGGAGAAGGTGGCTTACACCTGTAATGCCAGAGCTTTGG
GAGGCGGAGGCAAGAGGATCACTTGAAGCCAGGAGTTCAAGCTCAACCTGGGCAACGTAGACCCTGTCTCTACAAAAAATTAAAAACTTAGCCGGGCGTGGTGATGTGC
ACCTGTAGTCCTAGCTACTTGGGAGGCTGAGGCAGGAGGGTCATTTGAGCCCAAGAGTTTGAAGTTACCGAGAGCTATGATCCTGCCAGTGCATTCCAGCCTGGATGAC
AAAACGAGACCCTGTCTCTAAAAAACAAGAAGTGAGGGCTTTATGATTGTAGAATTTTCACTACAATAGCAGTGGACCAACCACCTTTCTAAATACCAATCAGGGAAGA
GATGGTTGATTTTTTAACAGACGTTTAAAGAAAAAGCAAAACCTCAAACTTAGCACTCTACTAACAGTTTAGCAGATGTTAATTAATGTAATCATGTCTGCATGTATG
GGATTATTTCCAGAAAGTGTATTGGGAAACCTCTCATGAACCCTGTGAGCAAGCCACCGTCTCACTCAATTTGAATCTTGGCTTCCCTCAAAAGACTGGCTAATGTTTG
GTAACTCTCTGGAGTAGACAGCACTACATGTACGTAAGATAGGTACATAAACAACTATTGGTTTTGAGCTGATTTTTTTCAGCTGCATTTGCATGTATGGATTTTTCTC
ACCAAAGACGATGACTTCAAGTATTAGTAAAATAATTGTACAGCTCTCCTGATTATACTTCTCTGTGACATTTCATTTCCCAGGCTATTTCTTTTGGTAGGATTTAAAA
CTAAGCAATTCAGTATGATCTTTGTCCTTCATTTTCTTTCTTATTCTTTTTGTTTGTTTGTTTGTTTTTTTCTTGAGGCAGAGTCTCTCTCTGTCGCCCAGGCTG
GAGTGCAGTGGCGCCATCTCAGCTCATTGCAACCTCTGCCACCTCCGGGTTCAAGAGATTCTCCTGCCTCAGCCTCCCGAGTAGCTGGGATTACAGGTGTCCACCACCA
CACCCGGCTAATTTTTTGTATTTTTAGTAGAGGTGGGGTTTCACCATGTTGGCCAGGCTGGTCTTGAGCTCCTGACCTCAGGTGATCCACCTGGCTAGCTAAGCCGGCT
TACTTAAGTAGGCCTCccgggccattgggccctagacaagttgtcgcggccgcactagtgactccagcgtaactggactggccacagttaggccgcaaatgtaatcaca
ctggctcaccttcgggtgggcctttctgcgtctagaggatcccctgttatgcccttccttacccatgcgctgagaggtgaaatcactctttaatttgtacgcggtacgg
acctctatatcgctttgtcgtagaggtggaatctccccagcttgcagccattgatagactgcgccgggtgtcacgcccacgccaaccgctgctttttccacatcgccga
aatggcgcacaagttcttcgggtttcatggggattattataatcaataagtgaaaattaaagctaggtataatttataaaaattatagccagctataaagagatcattt
atgattaagggtatgaaaacacgaggcgaacgactgaaagcacgccgtttagagctgaaactgacgctgaagcaagtggcagaagctgtaggaatctctcttccgggcg
tccaaaacttagaacgtggcgacgtgatgccgtcgctggagatcgggctatcgctggcaaaatgcctgcgtaagcccgtgcaatggatactgtatggcactgaatctga
tccagaccgcgttcctgttattggtacgacagagaccggaccggatagagactggcagcctggagaacctgccaacacagagcgattcctgccgttcgtgagtcaacgg
aacaccgtttacgcactgacggtcgggaaccagattcagcgaaactaccagccgggagacgttatcctggtcgattcttcacttacgcttgtaccgggtgaggatgtac
tggtttgtgataataacggcgagatcacaattcaacgattagcccgttatgacgagcagcattactaccttgatagtgctaactctcaacgggttatccatgataaaag
tgatcttcaatttgtgcaccaagtagtcggtacgatcaaatcgttcatggttgagggtagatgatacaataacagggtttattgctgactataattctggtttaatccg
atctatactttgtcggggttgaaccagaccgtagcagccgaaaaaagacgaaaaaaaacccgagtcggcaaactcgggcctttttcaggaagtagccacgataacgcag
atacgtccttcagaaagatagtgcgtttattgtggctgctcctgcggatttttttcaacccgaaaaaatgctaattcgcatggaaaggctaaaaaatgaccttacaagaa
ttctacgcggagcgctttggcagcgatccgttttcactgcttgacgcggcacgtgatgagctgactgagctggcgaagatggccggtatcaactggcctgcgtgtgctg
ataagattcagctcaatccgcgcggtggtgtagagcgctataccacctacaataattcctccccggaagctctgcagaagagccttaaaggccgtgtggaaatctattc
acgccttgagcagagcaaggatggcatcagttatcctttcgttaacttcgttgaaaaagcgcatgacgctggctcctggagtggcttttcattcctgttttcggagtat
cgccgtgaacagcaacggaatggtgcgaccgtggtcgctcagccggaagaagaacgtgcacgcatggagcgccaggctgaagcacgccgcctccgagctgaacaacaac
gcgttaatgatttaaaaaacaatcagatggagcaggaacgcctgcttggatggttggctttccatcgcgcctgggaacattcgccagctgaagatggctcctggcctta
tgcggtaaaaaagggaattcgtgacgtatttggcgcttgtgatatacgtcgcgtgaccagtcacgacagtgcgaaatggagccgtggaccaacaacttacatggcgata

cctctgtcccacctggacggaagaaaagacgggcggattgttggctggcaacgtatcgatctgaatggtggcaagttccagaccagcgccatcactaacggtgatttcg
tcggtgcatgctttgttattggcgacctgcaggggcgcaaaaaattgcgactgcagagggtttcgccaccggggcatccatctggctggcgaccagaaacgacccgaa
aaaacgctttgacgcggtagtcattgcggtttccgctaacaacatgattcacgttgtcgagcagctggtgaacatgtacccggctgcgcaaatcacttgcgctctcgat
aacgaccgcaaatcctcagctgaaggaaaaggcaacacaggcctccgcaccggattcgacatcatggagaagttttccggcgtcaaatgtgtttacccaacttttgagg
atgaccctgagctggaatgcagcgattttaacgacctgcacagcctgagagggctgaaggaagtcgctcgccagttaacgagaaatcatctgagccgtgcaaccgatct
gttgtcgatcacgctgaataagctgcgtactctcccgcgtctgaacagacgcacttttgccaaagaactgcttcgcgccgttgatattggcatgctgacatgcccggta
ccaaacagcccgaaagaacttatgcgccttttcagcagcacgctgcgggatatgggtatcgcagaaatttataacggtaccgttaaagatcacattacgcgccggttga
atcgcaaatgccgtgccgcgcaaacatcacgttcgttcagtgaacgcatcaccaacccgaaccttcgcccgtcacacatcacttacaaacggtttgaaacctccaggat
gaccgatgaggtgatgacatacgccgcacagctgcagggcatcgttattgtccgcgccgggatggggtcgggtaaatcgacaggcctcctgcgtccactgatgctgcag
tccacgcgtggcgtttccgtcgcgcaccgcgtatccctgataggcggcctgcatgaaatgatgaccgaagggaaaggcgcgaaagccgacattctgcattatcaggatc
ccggctatcaggaaatggcgccatatgccaacaagctgactatttgcatcaactccatacttaaaggctgctggcaaccgctgatgcgccagcatgacttcttcggctt
cgatgaagcaacacagggcctgcgtgccattctggccgggcgtgcgatggaaaacccggtaggcgtattcaacacgcttatcgacgcgctggcgcgtactgaagagcat
gccattatggtggatgccgatgccaacgatctgcttgttgacctggctgaactggcgatgaagcgacgcgaggagctgggcctacctgcctggctgcaaattcacgtga
ttgaactcccggtcgacgttcgcaaccgcgaaaccaacaagcctatccgcgtattttataccgagaaaaatcggatcatgaccgaggttatcgctgcagtgcaacgcgg
cgaacgaatcatgctggccaccgacagttcgacgttcgccgaagacgttaccatgcagctgagactacagttccctgacaaaaagtttctctgcgttaaccagaaaaac
aaacaggagaaggaagtcgacgatttcaccaaccagcctaaagtgatggtgaaaaaatatgacggcctcatctacagcccgtcgatatcttcaggtgtatcgattgagg
agaaacacttccaccgccatttcggcatgttctgcggcgaagtggtccccagcgacgccatccagatgctgcgccgcgaccgtacagctcaggaatacattatcggttt
cgacaagcttcgcggtaaacgtgaaaccgatccggaaaaaatcaaacgcgcctacgcgcaggccttgctcgaaacggctggccactccgggctgctgacagacgttgtc
tttgacggtgaccgaatttcactcggcgtggctaactcttcattcatgcagctcaaaattaaagccgccgcgctcgaggcatcggccagaaatgattatgccagcaaca
tgatttgcatcatgcatgatgatggctatcaggtcgcgcctatggccaccgacgcgcttgcaaacagcatcggtaaggatttacgtaaagaagcccgtgaactggtctt
tgagcagcttatggagcgccacctgagtgtcgatactcctgaccaggccgaacacgatgagttgataaaaaaacgcaccctgtccctggatgagcaggcccagctggtc
cgctgggacatcgaaaaggagctgcagctggatgtcgacgaggtggcgttaaaattctatttcgacggcggcctgaaaaaagtgcgcctgtttgaaaccatgcagctgg
atgagataaccgcccgtcgccttgaccgtgaagaagcgttgattcactttacctacgcttatcgcgttgccggaagatggcagcagttcgtcactacggccatgacgcg
ggaacaggccgacgcagagttccaggctaaattcccggccataaccgattaccgagtgaaatcgacccctgctgttgagataggcatgcgtggtttctacaccctcaaa
tcggcgacgctgcagcagtacttccgcgactgcggtatcgatccaaaaaccctggaaggtgaagcagacatggacgccctcaaacgcgccagagataacctgctcacgc
cggaacggcgcgatctgctaaacaacgttttacgcatcggcggcttcaacacagagaaaggcaagaagaaagcgcctgcagacctctgcataggcatcctggagtctat
ggggctatccagtaagaccagacgcgccagagacggcgatgcacgccccacgatgcgttccattgacccggactcggtcgagttcctcatgaatatcgtggagaagcgc
cgcgaggccggtttatcaattcacgcacgtaaggtggaaaaaaccaccatcgaagtggatcgcgatttggatctaaatatagatatacatggtaaccctcgatccaaaa
cagagcacgttccggacgccccacaatcagtaattatccaggcactggaggctatcccggtggcggtaccggaggcgtgggcggagaacgcgttgccagcaaccgaaat
ggaggcggtacgcctgtggccagtggccagcatcgcgagaacgttcgcctcgctgtacatgaccgaatttatggacctgctatcagtacgcgaaataaggctgctgaaa
gcgtttctaagccagcggcaagccgtggcgatataaccggaggggtaatgggaaaacaacgggaatgccttgtcatcgtggagggctacgacccttatgtggtcaaggt
cagccacgacgcgacaagctatagcgattataaaactggccgggaacttccattcacgatcattcggatgaacgggttctcagatagtgagggagtttattgctgcatt
gccagcgataagagcatggaagaccacacagacgaagccagggcgcttctaaaagcctataaggtgaagccactatgacgtcgacaccagagttcctgaagcaactgga
ttacgaacagctgaagtactgtcgcgagctgtgcgataaccgcatccgggctattgaggccgaagagaaaaagatggcctgggcagttactgatggtggcattaactac
ggctggttccgtacagaagactacatgaaggcggtagaatgcctggttgctacagctgcagaacggtgggaagaatcggataaggctgacccatctggacgtggttggc
ttgacctttcaatccgtggaagccgcctgccggtgtccgagtatgaagctttatttgctgatggccagtgggggtgaatagtggcgcatgatctacaaaaaatgatgga
tggtgagctggtccgcttacaacttgagattaagaaagaacaggagaggcgtttagccgaaaagatgattcctgtttttggtgtgtggcaatcacctattaatcaatat
gaatttgctgatccagagcaggcaatttcttgtgccgcagggctattggataaggttcttcgtgaggtcaaggacgatctttccaaagaaggtggattacagggatgga
acggcaatttgctacgcatctacgtacagcatgtgaacgagagcgattttgcagttctacagccttatttaaaggacaaataatcatgatttatcgcagaggatgggtg
cctgttttgtggaggaatgagctggaaaagcggctcaaagaagaagggttcgaaaactggaagcagataagtaactttctttgtggtgaagaggattgctactcggaag
cagcaaaaacgccaggacaatacgcttaccaggtcgtcgacaataccgaatggatggaacgtcgtgatgcgacgttctggcagcgtctgaaccgtttgtggttcgttcc
gctgtacctgctgactattccgttccagtggctcattcgtggccgtatggggtttgagacaacatctaaggtggctgcggtcatcaataagattaccggactccaataa
ctgatcaggttctgatcagtcagtaaagcaaactataaaaacacctgtgcacgtgactggtcacggggtaaaatatccacctaaattattgacgtgcgttctgctttgg
cggtagagttaccccgctgcagcaaaatctgcagccgggcctctcaaccccgaatgacaagaagcgcacaacacgcgccagcgtgtttttttgtgtgttaaatctgcgc
atacctgaattatggtggctcagatggggccaacttcggttgggccggtttcttcttgtaccggtgttgagaaccctgtctgggctaccacccctatagagattctcaa
ctctggtggtagcaccctatacaagataggaatgcataccatgttcaaattcaaatttgcagctgtcgtccgcacggacaaaaaatcccatatccaccgcctttccacc
attgcatcatccgagcgagaagctcgtcgccagttcgccagccgttttgtcctcgttctgtcagcccgtatcccggtcagcgaggtgatcgcatgaaccagattcagtt
aaacgcccagggcctgctggagacgattgaggaacgcctgctgcaggttgaagcgcttgtgggctctgctcaccgcactatctccagttatgaggccacgctgtatttg
caggaagctgcggaacttctgcaggtcgcgcgcgaactcacgcaggaagctcgcggctgctccctgtccctttctgagcagctgaaatccgggaggcgcaatgaaggc
tctctccgtcttttcatttcaggaaagccaccctatacgggtggttcttgtcggtggtgacccgtggtttgtggcgctggatatctgtgctgcattgaatatagccaac
ccttctgatgcattgcgtaagctggatcatgatgaaaaattgaccctcggtttaaccgaggcacaaaaacttgatcggatggccagggaagtaaatgtggtatcggaat
ctggtctctacaccataatcctccgctgccgcgacgcggtgaaacaggggacgacagcctggcggttccgcaagtgggtgacgaacgaggttctgccagctatccggaa
aaacggcgagtatgcttttgtcgagcctgagcctaagaacgccggtgaaccactggactggcggcagaaagaagaacttcggggcctgataaacgacatagcacaaagt
tttcagtaccgtaacgcatgggttagcggcgtctggatggctctccgccgcgcctgcaggaacccatcaccgaacccaataaccgtggatgatctcccggcaatcatcg
ccgagctgcgccggatattaacggctgcagaaacggctctgggcaacatgcgtgtctacgaacgggagctgctgcgcgatgtagttcgtggcggtcgccggagtatgtc
ttgtggagagctgccgatcaccgatattgatacggaactggagaaggtgctgccagcacattttgagctggccattgaaaagctggaaacgctatccacgaaactaaca
tcgccagcgattgcagaaaaataggggttggccagcccgaatttaagccccgctttacgcggggaagtccctgcagtcgggacggcgatgtccgtagttgcagctgcagt
ttccgcctaatccttccgtcccttgcaggggatgggtagggacaggcctcaataaaaaagccaccgattaacaccggtggcttttcttttagctgtagtacgtttccca
tgcggatcgcattgcggccatagggctatcggcggggccggaccaggcataatgctttccatcgtattcaaactctatcttgtacgttccgtccccgttatttttgca
ggcttgaagacgggttttagagcagttggctgatgttcttctggcccttcctcttcggttggttcatcgccgccaccctcgtcgagctcaatttcatcttcatccagct
cgtcgtcttgggactcatcatccggttcgtcgatggtctcaacggattcttcatcaggcaggacgatt**gcTggtg**tctctatcttcagctgccactgcccgttctcgcc
aacgaactgccccaatgcatcagcggcaaactccaggtaccggctaatcatcgtcgggctaaatttaaaggcccggagagtgctgttggttattttttgctgatgggtcc
tgctccaccagctgcttaacggtttcatggagacggacgccagcgtcacctctggcaaagcctggcatttcatcgtccagtttctgcagagccaccagcctggtgtttt
catccccaacttcaggtcgccaggttctggagaagttggccagcttgaactgcttatagtgcagctgggtgttctcatcgtcgtgtccgagaatctccatgaagaacac
atcctcgtcgacgtttttccaccgtggatcgacgcggaagaacatctcataagcgatgcgagcgtaaatagcgcggctatctttataaacacgacggtcatcgccgaaa
aatgatttaacccaagggttaaatgcttttgctaaaatagcatttatcctgccgttctcagaccttgtatcatcctttccatatcctttaacaacctcatcgaaatcag
atgcagcagagcaagaacgcaattctgttaataattcaacgaataattttgcttcgcataaagtataaatcgttctggttacgcttttatcttcagagcgtttttagc
ttgccctgagaaattaaccgtatactttcctgaaacggcaaattcaccctgaaacattatctcaatcattcttcgccctgataccgcagccagagcaaaggccaaaggt
gccattccagaacgagtgtttaaactaaataaagtcgcaggattattcaaaatatcatagatagactgcatgtatgttgggtagtcaataaccacaacattacgcttct
tctcgcgcagaacatcggcccatcgttgctgtatagatgtacgctccgcagggcttagctgcagatggtacagaacctcatggttgaccttgagctggtgtagttcttc
taacaacgcagagccttgttggaataacttataaagatagtcacggcgctccttccaatcatcactgtttaaatcactaagagcaaaactccaatctggatatttttt
attagtcttgctattttgcatcactgcctttagagcctattcttacatttgataactcttcggcaagcggcattatttctttcaatttagattgtaacgatgacatgt
gctggcgaatattagccgtaggcatagaaagccatgaagataattcttcgctgtaaagaggatacttttccgataatttattaatatttttatcaaagctatgatgtaa
tttatcatcaaaccgctttcttgccctgctcatataggcgttaaaagtattcgcggttattcttttctgcaatcctttcccacggaactttcttttatcattaaataac
gcgttcttataccgtgcggctgcggctttaattctcttcgttttgtcgccttgtgggcggtctgaggcatcaattgcctctacctcattcacaagcgtgttgatcaact
caccgatttttaccttgctcatcccggttccccgtttctaagtctctggatatcgtaacacaatcaaagaacaagtgacgattaattatgctgctttcgtcacactat
aagagaacataatagcagaataacaaaggaacacaacaccggaatagatacatattgttctgcgcttatgttctccttatcagcacacaatagtccattatacgc

### pJazz-V4

See Figure 2D. Replacement of landing pad (AarI fragment removal) with pure repeat (derived from pJazz-V1). pJazz backbone in lowercase. Mouse sequence in **green**. Human sequence in **blue**, selection cassette in **magenta**, pathogenic repeat in **red**, AarI site mutation indicated by **gcTggtg**.

gcgtataatgggcaattgtgtgctgatatgttccgtaattgagatatgatatagtgcatatgttccgaattattgatacattaatgttgcattcctatgttgtgttctg
ttatggtgttcttagatagtggggatgtcacatggattgatgttctttgattgtgtttcaatatagagacggaagagatagagatgcctcaaagtgtataaacagcgt
gaaaatcagcatgttcgcttaacttttgatagcaattggtatgatttattatcatttaaaatcattaagttaggttttaaatataccaacgtaagtaataccgttgcgc
ttagatgaagctaaatagcttctgtagcgcatcaaaaaattgatttgaaccttaactttgcgttttctaagtcactttgctttttttatgttgctttgctttaaaaacg
taaagtatgatgaaaaacacaaagataagggcatgtttgagaggaaacaggatgtcgttaattaatttgctgaaagactgtatcaatcgcggtcaggaaatgacgcgcg
ctattgccatcgcacagttcggcgatgacagcccggaagcgcgtcgtattactcgccgctggggtataacagaagttgcagacctgatcggcgtaacaccgcaggctat
cagagacgcggaaaaagctggtcgtctaccggctcctgattttgagatgaggggccgagtagaacgtcgcgctggctacacaattgaccaaattagccatatgcggagt
gtgtttggtaacccaaaccaaagacctgacgataaaaacccggttgtactttccgttatgtcacacaagggaggggtttataaaacctcatctgcagtacaccaggcgc
aatggttagccttgcaagggcaccgcgtcctactcgttgaaggaacgacccgcaagggactgcatctatgtaccacggttacgttcctgatttgcacattcacgcaga
cgatacactgcttccatttacctgggtaaacgcgacaacgctgaatatgcgataaaaccaacatgctggccgggtctcgacataatccccagctgcctggcccttcac
cgcattgaaacagatctagcagaaagtcaaaagcctccgaccggaggcttttgacttctgtcacctaggttacgccccgccctgccactcatcgcagtactgttgtaat
tcattaagcattctgccgacatggaagccatcacaaacggcatgatgaacctgaatcgccagcggcatcagcaccttgtcgccttgcgtataatatttgcccatggtga
aaacgggggcgaagaagttgtccatattggccacgtttaaatcaaaactggtgaaactcacccagggattggctgagacgaaaaacatattctcaataaaccctttagg
gaaataggccaggttttcaccgtaacacgccacatcttgcgaatatatgtgtagaaactgccggaaatcgtcgtggtattcactccagagcgatgaaaacgtttcagtt
tgctcatggaaaacggtgtaacaagggtgaacactatcccatatcaccagctcaccgtctttcattgccatacgaaattccggatgagcattcatcaggcgggcaagaa
tgtgaataaaggccggataaacttgtgcttatttttctttacggtctttaaaaaggccgtaatatccagctgaacggtctggttataggtacattgagcaactgactg
aaatgcctcaaaatgttctttacgatgccattgggatatatcaacggtggtatatccagtgatttttttctccatttagcttccttagctcctgaaaatctcgataac
tcaaaaaatacgcccggtagtgatcttatttcattatggtgaaagttggaacctcttacgtgccgatcagattaaaacgaaaggcccagtctttcgactgagcctttcg
ttttatttgaccatgttggtatgatttaaatggtcagttaatcagttctatgtaccagcaaggtccagttgtaagcggccgcttgacttcaagtctaatggcccGTGAA
AGATGGCGTTTGTAGTGACAGCCATCCCAATTGCCCTTTCCTTCTAGGTGGAAAGTGGTGTCTAGACAGTCCAGGGAGGGTGTGCGAGGGAGGTGCGTTTTGGTTGCCT
CAGCTCGCAACTTAACTCCACAACGGTGACCAAGGACAAAAGAAGGAAACAAGACTGCAGAGATCCGCACCGGGGAGCCCTGCAGATTCTGGGTGTGTGTTTTTGTTTT
TCCCACCCTCTCTCCCCACTACTTGCTCTCACAGTACTCGCTGAGGGTGAACAAGAAAAGACCTGATAAAGATTAACCCTAGAAAGATAGTCTGCGTAAAATTGACGCA
TGCATTCTTGAAATATTGCTCTCTCTTTCTAAATAGCGCGAATCCGTCGCTGTGCATTTAGGACATCTCAGTCGCCGCTTGGAGCTCCCGTGAGGCGTGCTTGTCAATG
CGGTAAGTGTCACTGATTTTGAACTATAACGACCGCGTGAGTCAAAATGACGCATGATTATCTTTTACGTGACTTTTAAGATTTAACTCATACGATAATTATATTGTTA
TTTCATGTTCTACTTACGTGATAACTTATTATATATATATTTTCTTGTTATAGATATCGCGGCCGCTCTAGCCTCGAGGCTAGAACTAGTGGATCTCGAGCCCCAGCTG
GTTCTTTCCGCCTCAGAAGCCATAGAGCCCACCGCATCCCCAGCATGCCTGCTATTGTCTTCCCAATCCTCCCCCTTGCTGTCCTGCCCCACCCCACCCCCCAGAATAG
AATGACACCTACTCAGACAATGCGATGCAATTTCCTCATTTTATTAGGAAAGGACAGTGGGAGTGGCACCTTCCAGGGTCAAGGAAGGCACGGGGGAGGGGCAAACAAC
AGATGGCTGGCAACTAGAAGGCACAGTCGAGGCTGATCAGCGAGCTCTAGAGCTCAGAAGAACTCGTCAAGAAGGCGATAGAAGGCGATGCGCTGCGAATCGGGAGCGG
CGATACCGTAAAGCACGAGGAAGCGGTCAGCCCATTCGCCGCCAAGCTCTTCAGCAATATCACGGGTAGCCAACGCTATGTCCTGATAGCGGTCCGCCACACCCAGCCG
GCCACAGTCGATGAATCCAGAAAAGCGGCCATTTTCCACCATGATATTCGGCAAGCAGGCATCGCCATGGGTCACGACGAGATCCTCGCCGTCGGGCATGCGCGCCTTG
AGCCTGGCGAACAGTTCGGCTGGCGCGAGCCCCTGATGCTCTTCGTCCAGATCATCCTGATCGACAAGACCGGCTTCCATCCGAGTACGTGCTCGCTCGATGCGATGTT
TCGCTTGGTGGTCGAATGGGCAGGTAGCCGGATCAAGCGTATGCAGCCGCCGCATTGCATCAGCCATGATGGATACTTTCTCGGCAGGAGCAAGGTGAGATGACAGGAG
ATCCTGCCCCGGCACTTCGCCCAATAGCAGCCAGTCCCTTCCCGCTTCAGTGACAACGTCGAGCACAGCTGCGCAAGGAACGCCCGTCGTGGCCAGCCACGATAGCCGC
GCTGCCTCGTCCTGCAGTTCATTCAGGGCACCGGACAGGTCGGTCTTGACAAAAAGAACCGGGCGCCCCTGCGCTGACAGCCGGAACACGGCGGCATCAGAGCAGCCGA
TCGTCTGTTGTGCCCAGTCATAGCCGAATAGCCTCTCCACCCAAGCGGCCGGAGAACCTGCGTGCAATCCATCTTGTTCAATGGCCGATCCCATGGTTTAGTTCCTCAC
CTTGTCGTATTATACTATGCCGATATACTATGCCGATGATTAATTGTCAACAGGCTGCAGGTCGAAAGGCCCGGAGATGAGGAAGAGGAGAACAGCGCGGCAGACGTGC
GCTTTTGAAGCGTGCAGAATGCCGGGCCTCCGGAGGACCTTCGGGCGCCCGCCCCGCCCCTGAGCCCGCCCCTGAGCCCGCCCCCGGACCCACCCCTTCCCAGCCTCTG
AGCCCAGAAAGCGAAGGAGCAAAGCTGCTATTGGCCGCTGCCCCAAAGGCCTACCCGCTTCCATTGCTCAGCGGTGCTGTCCATCTGCACGAGACTAGTGAGACGTGCT
ACTTCCATTTGTCACGTCCTGCACGACGCGAGCTGCGGGGCGGGGGGGAACTTCCTGACTAGGGGAGGAGTAGAAGGTGGCGCGAAGGGGCCACCAAAGAACGGAGCCG
GTTGGCGCCTACCGGTGGATGTGGAATGTGTGCGAGGCCAGAGGCCACTTGTGTAGCGCCAAGTGCCCAGCGGGGCTGCTAAAGCGCATGCTCCAGACTGCCTTGGGAA
AAGCGCCTCCCCTACCCGGTAGAATTAATTCGATATCAAGCTTGCTCGGATGATAGGGCCCTAACGTGTTTTTGCTTTGTTACTTTATAGAAGAAATTTTGAGTTTTTG
TTTTTTTTTAATAAATAAATAAACATAAATAAATTGTTTGTTGAATTTATTATTAGTATGTAAGTGTAAATATAATAAAACTTAATATCTATTCAAATTAATAAATAAA
CCTCGATATACAGACCGATAAAACACATGCGTCAATTTTACGCATGATTATCTTTAACGTACGTCACAATATGATTATCTTTCTAGGGTTAACCAGAAGAAAACAAGGA
GGGAAACAACCGCAGCCTGTAGCAAGCTCTGGAACTCAGGAGTCGCGCGCTAGGGGCCGGGGCCGGGGCCGGGGCCGGGGCCGGGGCCGGGGCCGGGGCCGGGGCCGGG
GCCGGGGCCGGGGCCGGGGCCGGGGCCGGGGCCGGGGCCGGGGCCGGGGCCGGGGCCGGGGCCGGGGCCGGGGCCGGGGCCGGGGCCGGGGCCGGGGCCGGGGCCGGGG
CCGGGGCCGGGGCCGGGGCCGGGGCCGGGGCCGGGGCCGGGGCCGGGGCCGGGGCCGGGGCCGGGGCCGGGGCCGGGGCCGGGGCCGGGGCCGGGGCCGGGGCCGGGGC
CGGGGCCGGGGCCGGGGCCGGGGCCGGGGCCGGGGCCGGGGCCGGGGCCGGGGCCGGGGCCGGGGCCGGGGCCGGGGCCGGGGCCGGGGCCGGGGCCGGGGCCGGGGCC
GGGGCCGGGGCCGGGGCCGGGGCCGGGGCCGGGGCCGGGGCCGGGGCCGGGGCCGGGGCCGGGGCCGGGGCCGGGGCCGGGGCCGGGGCCGGGGCCGGGGCCGGGGCCG
GGGCCGGGGCCGGGGCCGGGGCCGGGGCCGGGGCCGGGGCCGGGGCCGGGGCCGGGGCCGGGGCCGGGGCCGGGGCCGGGGCCGGGGCCGGGGCCGGGGCCGGGGCCGG
GGCCGGGGCCGGGGCCGGGGCCGGGGCCGGGGCCGGGGCCGGGGCCGGGGCCGGGGCCGGGGCCGGGGCCGGGGCCGGGGCCGGGGCCGGGGCCGGGGCCGGGGCCGGG
GCCGGGGCCGGGGCCGGGGCCGGGGCCGGGGCCGGGGCCGGGGCCGGGGCCGGGGCCGGGGCCGGGGCCGGGGCCGGGGCCGGGGCCGGGGCCGGGGCCGGGGCCGGGG
CCGGGGCCGGGGCCGGGGCCGGGGCCGGGGCCGGGGCCGGGGCCGGGGCCGGGGCCGGGGCCGGGGCCGGGGCCGGGGCCGGGGCCGGGGCCGGGGCCGGGGCCGGGGC
CGGGGCCGGGGCCGGGGCCGGGGCCGGGGCCGGGGCCGGGGCCGGGGCCGGGGCCGGGGCCGGGGCCGGGGCCGGGGCCGGGGCCGGGGCCGGGGCCGGGGCCGGGGCC
GGGGCCGGGGCCGGGGCCGGGGCCGGGGCCGGGGCCGGGGCCGGGGCCGGGGCCGGGGCCGGGGCCGGGGCCGGGGCCGGGGCCGGGGCCGGGGCCGGGGCCGGGGCCG
GGGCCGGGGCCGGGGCCGGGGCCGGGGCCGGGGCCGGGGCCGGGGCCGGGGCCGGGGCCGGGGCCGGGGCCGGGGCCGGGGCCGGGGCCGGGGCCGGGGCCGGGGCCGG
GGCCGGGGCCGGGGCCGGGGCCGGGGCCGGGGCCGGGGCCGGGGCCGGGGCCGGGGCCGGGGCCGGGGCCGGGGCCGGGGCCGGGGCCGGGGCCGGGGCCGGGGCCGGG
GCCGGGGCCGGGGCCGGGGCCGGGGCCGGGGCCGGGGCCGGGGCCGGGGCCGGGGCCGGGGCCGGGGCCGGGGCCGGGGCCGGGGCCGGGGCCGGGGCCGGGGCCGGGG
CCGGGGCCGGGGCCGGGGCCGGGGCCGGGGCCGGGGCCGGGGCCGGGGCCGGGGCCGGGGCCGGGGCCGGGGCCGGGGCCGGGGCCGGGGCCGGGGCCGGGGCCGGGGC
CGGGGCCGGGGCCGGGGCCGGGGCCGGGGCCGGGGCCGGGGCCGGGGCCGGGGCCGGGGCCGGGGCCGGGGCCGGGGCCGGGGCCGGGGCCGGGGCCGGGGCCGGGGCC
GGGGCCGGGGCCGGGGCCGGGGCCGGGGCCGGGGCCGGGGCCGGGGCCGGGGCCGGGGCCGGGGCCGGGGCCGGGGCCGGGGCCGGGGCCGGGGCCGGGGCCGGGGCCG
GGGCCGGGGCCGGGGCCGGGGCCGGGGCCGGGGCCGGGGCCGGGGCCGGGGCCGGGGCCGGGGCCGGGGCCGGGGCCGGGGCCGGGGCCGGGGCCGGGGCCGGGGCCGG
GGCCGGGGCCGGGGCCGGGGCCGGGGCCGGGGCCGGGGCCGGGGCCGGGGCCGGGGCCGGGGCCGGGGCCGGGGCCGGGGCCGGGGCCGGGGCCGGGGCCGGGGCCGGG
GCCGGGGCCGGGGCCGGGGCCGGGGCCGGGGCCGGGGCCGGGGCCGGGGCCGGGGCCGGGGCCGGGGCCGGGGCCGGGGCCGGGGCCGGGGCCGGGGCCGGGGCCGGGG
CCGGGGCCGGGGCCGGGGCCGGGGCCGGGGCCGGGGCCGGGGCCGGGGCCGGGGCCGGGGCCGGGGCCGGGGCCGGGGCCGGGGCCGGGGCCGGGGCCGGGGCCGGGGC
CGGGGCCGGGGCCGGGGCCGGGGCCGGGGCCGGGGCCGGGGCCGGGGCCGGGGCCGGGGCCGGGGCCGGGGCCGGGGCCGGGGCCGGGGCCGGGGCCGGGGCCGGGGCC
GGGGCCGGGGCCGGGGCCGGGGCCGGGGCCGGGGCCGGGGCCGGGGCCGGGGCCGGGGCCGGGGCCGGGGCCGGGGCCGGGGCCGGGGCCGGGGCCGGGGCCGGGGCCG
GGGCCGGGGCCGGGGCCGGGGCCGGGGCCGGGGCCGGGGCCGGGGCCGGGGCCGGGGCCGGGGCCGGGGCCGGGGCCGGGGCCGGGGCCGGGGCCGGGGCCGGGGCCGG
GGCCGGGGCCGGGGCCGGGGCCGGGGCCGGGGCCGGGGCCGGGGCCGGGGCCGGGGCCGGGGCCGGGGCCGGGGCCGGGGCCGGGGCCGGGGCCGGGGCCGGGGCCGGG
GCCGGGGCCGGGGCCGGGGCCGGGGCCGGGGCCGGGGCCGGGGCCGGGGCCGGGGCCGGGGCCGGGGCCGGGGCCGGGGCCGGGGCCGGGGCCGGGGCCGGGGCCGGGG
CCGGGGCCGGGGCCGGGGCCGGGGCCGGGGCCGGGGCCGGGGCCGGGGCCGGGGCCGGGGCCGGGGCCGGGGCCGGGGCCGGGGCCGGGGCCGGGGCCGGGGCCGGGGC
CGGGGCCGGGGCCGGGGCCGGGGCCGGGGCCGGGGCCGGGGCCGGGGCCGGGGCCGGGGCCGGGGCCGGGGCCGGGGCCGGGGCCGGGGCCGGGGCCGGGGCCGGGGCC
GGGGCCGGGGCCGGGGCCGGGGCCGGGGCCGGGGCCGGGGCCGGGGCCGGGGCCGGGGCCGGGGCCGGGGCCGGGGCCGGGGCCGGGGCCGGGGCCGGGGCCGGGGCCG
GGGCCGGGGCCGGGGCCGGGGCCGGGGCCGGGGCCGGGGCCGGGGCCGGGGCCGGGGCCGGGGCCGGGGCCGGGGCCGGGGCCGGGGCCGGGGCCGGGGCCGGGGCCGG
GGCCGGGGCCGGGGCCGGGGCCGGGGCCGGGGCCGGGGCCGGGGCCGGGGCCGGGGCCGGGGCCGGGGCCGGGGCCGGGGCCGGGGCCGGGGCCGGGGCCGGGGCCGGG

GCCGGGGCCGGGGCCGGGGCCGGGGCCGGGGCCGGGGCCGGGGCCGGGGCCGGGGCCGGGGCCGGGGCCGGGGCCGGGGCCGGGGCCGGGGCCGGGGCCGGGGCCGGGG
CCGGGGCCGGGGCCGGGGCCGGGGCCGGGGCCGGGGCCGGGGCCGGGGCCGGGGCCGGGGCCGGGGCCGGGGCCGGGGCCGGGGCCGGGGCCGGGGCCGGGGCCGGGGC
CGGGGCCGGGGCCGGGGCCGGGGCCGGGGCCGGGGCCGGGGCCGGGGCCGGGGCCGGGGCCGGGGCCGGGGCCGGGGCCGGGGCCGGGGCCGGGGCCGGGGCCGGGGCC
GGGGCCGGGGCCGGGGCCGGGGCCGGGGCCGGGGCCGGGGCCGGGGCCGGGGCCGGGGCCGGGGCCGGGGCCGGGGCCGGGGCCGGGGCCGGGGCCGGGGCCGGGGCCG
GGGCCGGGGCCGGGGCCGGGGCCGGGGCCGGGGCCGGGGCCGGGGCCGGGGCCGGGGCCGGGGCCGGGGCCGGGGCCGGGGCCGGGGCCGGGGCCGGGGCCGGGGCCGG
GGCCGGGGCCGGGGCCGGGGCCGGGGCCGGGGCCGGGGCCGGGGCCGGGGCCGGGGCCGGGGCCGGGGCCGGGGCCGGGGCCGGGGCCGGGGCCGGGGCCGGGGCCGGG
GCCGGGGCCGGGGCCGGGGCCGGGGCCGGGGCCGGGGCCGGGGCCGGGGCCGGGGCCGGGGCCGGGGCCGGGGCCGGGGCCGGGGCCGGGGCCGGGGCCGGGGCCGGGG
CCGGGGCCGGGGCCGGGGCCGGGGCCGGGGCCGGGGCCGGGGCCGGGGCCGGGGCCGGGGCCGGGGCCGGGGCCGGGGCCGGGGCCGGGGCCGGGGCCGGGGCCGGGGC
CGGGGCCGGGGCCGGGGCCGGGGCCGGGGCCGGGGCCGGGGCCGGGGCCGGGGCCGGGGCCGGGGCCGGGGCCGGGGCCGGGGCCGGGGCCGGGGCCGGGGCCGGGGCC
GGGGCCGGGGCCGGGGCCGGGGCCGGGGCCGGGGCCGGGGCCGGGGCCGGGGCCGGGGCCGGGGCCGGGGCCGGGGCCGGGGCCGGGGCCGGGGCCGGGGCCGGGGCCG
GGGCCGGGGCCGGGGCCGGGGCCGGGGCCGGGGCCGGGGCCGGGGCCGGGGCCGGGGCCGGGGCCGGGGCCGGGGCCGGGGCCGGGGCCGGGGCCGGGGCCGGGGCCGG
GGCCGGGGCCGGGGCCGGGGCCGGGGCCGGGGCCGGGGCCGGGGCCGGGGCCGGGGCCGGGGCCGGGGCCGGGGCCGGGGCCGGGGCCGGGGCCGGGGCCGGGGCCGGG
GCCGGGGCCGGGGCCGGGGCCGGGGCCGGGGCCGGGGCCGGGGCCGGGGCCGGGGCCGGGGCCGGGGCCGGGGCCGGGGCCGGGGCCGGGGCCGGGGCCGGGGCCGGGG
CCGGGGCCGGGGCCGGGGCCGGGGCCGGGGCCGGGGCCGGGGCCGGGGCCGGGGCCGGGGCCGGGGCCGGGGCCGGGGCCGGGGCCGGGGCCGGGGCCGGGGCCGGGGC
CGGGGCCGGGGCCGGGGCCGGGGCCGGGGCCGGGGCCGGGGCCGGGGCCGGGGCCGGGGCCGGGGCCGGGGCCGGGGCCGGGGCCGGGGCCGGGGCCGGGGCCGGGGCC
GGGGCCGGGGCCGGGGCCGGGGCCGGGGCCGGGGCCGGGGCCGGGGCCGGGGCCGGGGCCGGGGCCGGGGCCGGGGCCGGGGCCGGGGCCGGGGCCGGGGCCGGGGCCG
GGGCCGGGGCCGGGGCCGGGGCCGGGGCCGGGGCCGGGGCCGGGGCCGGGGCCGGGGCCGGGGCCGGGGCCGGGGCCGGGGCCGGGGCCGGGGCCGGGGCCGGGGCCGG
GGCCGGGGCCGGGGCCGGGGCCGGGGCCGGGGCCGGGGCCGGGGCCGGGGCCGGGGCCGGGGCCGGGGCCGGGGCCGGGGCCGGGGCCGGGGCCGGGGCCGGGGCCGGG
GCCGGGGCCGGGGCCGGGGCCGGGGCCGGGGCCGGGGCCGGGGCCGGGGCCGGGGCCGGGGCCGGGGCCGGGGCCGGGGCCGGGGCCGGGGCCGGGGCCGGGGCCGGGG
CCGGGGCCGGGGCCGGGGCCGGGGCCGGGGCCGGGGCCGGGGCCGGGGCCGGGGCCGGGGCCGGGGCCGGGGCCGGGGCCGGGGCCGGGGCCGGGGCCGGGGCCGGGGC
CGGGGCCGGGGCCGGGGCCGGGGCCGGGGCCGGGGCCGGGGCCGGGGCCGGGGCCGGGGCCGGGGCCGGGGCCGGGGCCGGGGCCGGGGCCGGGGCCGGGGCCGGGGCC
GGGGCCGGGGCCGGGGCCGGGGCCGGGGCCGGGGCCGGGGCCGGGGCCGGGGCCGGGGCCGGGGCCGGGGCCGGGGCCGGGGCCGGGGCCGGGGCCGGGGCCGGGGCCG
GGGCCGGGGCCGGGGCCGGGGCCGGGGCCGGGGCCGGGGCCGGGGCCGGGGCCGGGGCCGGGGCCGGGGCCGGGGCCGGGGCCGGGGCCGGGGCCGGGGCCGGGGCCGG
GGCCGGGGCCGGGGCCGGGGCCGGGGCCGGGGCCGGGGCCGGGGCCGGGGCCGGGGCCGGGGCCGGGGCCGGGGCCGGGGCCGGGGCCGGGGCCGGGGCCGGGGCCGGG
GCCGGGGCCGGGGCCGGGGCCGGGGCCGGGGCCGGGGCCGGGGCCGGGGCCGGGGCCGGGGCCGGGGCCGGGGCCGGGGCCGGGGCCGGGGCCGGGGCCGGGGCCGGGG
CCGGGGCCGGGGCCGGGGCCGGGGCCGGGGCCGGGGCCGGGGCCGGGGCCGGGGCCGGGGCCGGGGCCGGGGCCGGGGCCGGGGCCGGGGCCGGGGCCGGGGCCGGGGC
CGGGGCGTGGTCGGGGCGGCCCGGGGGCGGCCCGGGGCGGGGCTGCGGTTGCGGTGCCTGCGCCCGCGGCGGCGGAGGCGCAGGCGGTGGCGAGTGGGTGAGTGAGG
AGGCGGCATCCTGGCGGGTGGCTGTTTGGGGTTCGGCTGCCGGGAAGAGGCGCGGGTAGAAGCGGGGGCTCTCCTCAGAGCTCGACGCATTTTTACTTTCCCTCTCATT
TCTCTGACCGAAGCTGGGTGTCGGGCTTTCGCCTCTAGCGACTGGTGGAATTGCCTGCATCCGGGCCCCGGGCTTCCCGGCGGCGGCGGCGGCGGCGGCGCAGGGA
CAAGGGATGGGGATCTGGCCTCTTCCTTGCTTTCCCGCCCTCAGTACCCGAGCTGTCTCCTTCCCGGGGACCCGCTGGGAGCGCTGCCGCTGCGGGCTCGAGAAAAGGG
AGCCTCGGGTACTGAGAGGCCTCGCCTGGGGGAAGGCCGGAGGGTGGGCGGCGCGCGGCTTCTGCGGACCAAGTCGGGGTTCGCTAGGAACCCGAGACGGTCCCTGCCG
GCGAGGAGATCATGCGGGATGAGATGGGGGTGTGGAGACGCCTGCACAATTTCAGCCCAAGCTTCTAGAGAGTGGTGATGACTTGCATATGAGGGCAGCAATGCAAGTC
GGTGTGCTCCCCATTCTGTGGGACATGACCTGGTTGCTTCACAGCTCCGAGATGACACAGACTTGCTTAAAGGAAGTGACTATTGTGACTTGGGCATCACTTGACTGAT
GGTAATCAGTTGTCTAAAGAAGTGCACAGATTACATGTCCGTGTGCTCATTGGGTCTATCTGGCCGCGTTGAACACCACCAGGCTTTGTATTCAGAAACAGGAGGGAGG
TCCTGCACTTTCCCAGGAGGGGTGGCCCTTTCAGATGCAATCGAGATTGTTAGGCTCTGGGAGAGTAGTTGCCTGGTTGTGGCAGTTGGTAAATTTCTATTCAAACAGT
TGCCATGCACCAGTTGTTCACAACAAGGGTACGTAATCTGTCTGGCATTACTTCTACTTTTGTACAAAGGATCAAAAAAAAAAAAGATACTGTTAAGATATGATTTTTC
TCAGACTTTGGGAAACTTTTAACATAATCTGTGAATATCACAGAAACAAGACTATCATATAGGGGATATTAATAACCTGGAGTCAGAATACTTGAAATACGGTGTCATT
TGACACGGGCATTGTTGTCACCACCTCTGCCAAGGCCTGCCACTTTAGGAAAACCCTGAATCAGTTGGAAACTGCTACATGCTGATAGTACATCTGAAACAAGAACGAG
AGTAATTACCACATTCCAGATTGTTCACTAAGCCAGCATTTACCTGCTCCAGGAAAAAATTACAAGCACCTTATGAAGTTGATAAAATATTTTGTTTGGCTATGTTGGC
ACTCCACAATTTGCTTTCAGAGAAACAAAGTAAACCAAGGAGGACTTCTGTTTTTCAAGTCTGCCCTCGGGTTCTATTCTACGTTAATTAGATAGTTCCCAGGAGGACT
AGGTTAGCCTACCTATTGTCTGAGAAACTTGGAACTGTGAGAAATGGCCAGATAGTGATATGAACTTCACCTTCCAGTCTTCCCTGATGTTGAAGATTGAGAAAGTGTT
GTGAACTTTCTGGTACTGTAAACAGTTCACTGTCCTTGAAGTGGTCCTGGGCAGCTCCTGTTGTGGAAAGTGGACGGTTTAGGATCCTGCTTCTCTTTGGGCTGGGAGA
AAATAAACAGCATGGTTACAAGTATTGAGAGCCAGGTTGGAGAAGGTGGCTTACACCTGTAATGCCAGAGCTTTGGGAGGCGGAGGCAAGAGGATCACTTGAAGCCAGG
AGTTCAAGCTCAACCTGGGCAACGTAGACCCTGTCTCTACAAAAAATTAAAAACTTAGCCGGGCGTGGTGATGTGCACCTGTAGTCCTAGCTACTTGGGAGGCTGAGGC
AGGAGGGTCATTTGAGCCCAAGAGTTTGAAGTTACCGAGAGCTATGATCCTGCCAGTGCATTCCAGCCTGGATGACAAAACGAGACCCTGTCTCTAAAAAACAAGAAGT
GAGGGCTTTATGATTGTAGAATTTTCACTACAATAGCAGTGGACCAACCACCTTTCTAAATACCAATCAGGGAAGAGATGGTTGATTTTTAACAGACGTTTAAAGAAA
AAGCAAAACCTCAAACTTAGCACTCTACTAACAGTTTTAGCAGATGTTAATTAATGTAATCATGTCTGCATGTATGGGATTATTTCCAGAAAGTGTATTGGGAAACCTC
TCATGAACCCTGTGAGCAAGCCACCGTCTCACTCAATTTGAATCTTGGCTTCCCTCAAAAGACTGGCTAATGTTTGGTAACTCTCTGGAGTAGACAGCACTACATGTAC
GTAAGATAGGTACATAAACAACTATTGGTTTTGAGCTGATTTTTTTCAGCTGCATTTGCATGTATGGATTTTTCTCACCAAAGACGATGACTTCAAGTATTAGTAAAAT
AATTGTACAGCTCTCCTGATTATACTTCTCTGTGACATTTCATTTCCCAGGCTATTTCTTTTGGTAGGATTTAAAACTAAGCAATTCAGTATGATCTTTGTCCTTCATT
TTCTTTCTTATTCTTTTTGTTTGTTTGTTTGTTTGTTTTTTTCTTGAGGCAGAGTCTCTCTCTGTCGCCCAGGCTGGAGTGCAGTGGCGCCATCTCAGCTCATTGCAAC
CTCTGCCACCTCCGGGTTCAAGAGATTCTCCTGCCTCAGCCTCCCGAGTAGCTGGGATTACAGGTGTCCACCACCACACCCGGCTAATTTTTTGTATTTTTAGTAGAGG
TGGGGTTTCACCATGTTGGCCAGGCTGGTCTTGAGCTCCTGACCTCAGGTGATCCACCTGGCTAGCTAAGCCGGCTTACTTAAGTAGGCCTCccgggccattgggccct
agacaagttgtcgcggccgcactagtgactccagcgtaactggactggccacagttaggccgcaaatgtaatcacactggctcaccttcgggtgggcctttctgcgtct
agaggatcccctgttatgcccttccttacccatgcgctgagaggtgaaatcactctttaatttgtacgcggtacggacctctatatcgctttgtcgtagaggtggaatc
tccccagcttgcagccattgatagactgcgccgggtgtcacgcccacgccaaccgctgctttttccacatcgccgaaatggcgcacaagttcttcgggtttcatgggga
ttattataatcaataagtgaaaattaaagctaggtataatttataaaaattatagccagctataaagagatcatttatgattaagggtatgaaaacacgaggcgaacga
ctgaaagcacgccgtttagagctgaaactgacgctgaagcaagtggcagaagctgtaggaatctctcttccgggcgtccaaaacttagaacgtggcgacgtgatgccgt
cgctggagatcgggctatcgctggcaaaatgcctgcgtaagcccgtgcaatggatactgtatggcactgaatctgatccagaccgcgttcctgttattggtacgacaga
gaccggaccggatagagactggcagcctggagaacctgccaacacagagcgattcctgccgttcgtgagtcaacggaacaccgtttacgcactgacggtcgggaaccag
attcagcgaaactaccagccgggagacgttatcctggtcgattcttcacttacgcttgtaccgggtgaggatgtactggtttgtgataataacggcgagatcacaattc
aacgattagcccgttatgacgagcagcattactaccttgatagtgctaactctcaacgggttatccatgataaaagtgatcttcaatttgtgcaccaagtagtcggtac
gatcaaatcgttcatggttgagggtagatgatacaataacagggtttattgctgactataattctggtttaatccgatctatactttgtcggggttgaaccagaccgta
gcagccgaaaaaagacgaaaaaaaacccgagtcggcaaactcgggccttttttcaggaagtagccacgataacgcagatacgtccttcagaaagatagtgcgtttattgt
ggctgctcctgcggatttttttcaacccgaaaaaaatgctaattcgcatggaaaggctaaaaaatgaccttacaagaattctacgcggagcgctttggcagcgatccgttt
tcactgcttgacgcggcacgtgatgagctgactgagctggcgaagatggccggtatcaactggcctgcgtgtgctgataagattcagctcaatccgcgcggtggtgtag
agcgctataccacctacaataattcctccccggaagctctgcagaagagccttaaaggccgtgtggaaatctattcacgccttgagcagagcaaggatggcatcagtta
tcctttcgttaacttcgttgaaaaagcgcatgacgctggctcctggagtggcttttcattcctgttttcggagtatcgccgtgaacagcaacggaatggtgcgaccgtg
gtcgctcagccggaagaagaacgtgcacgcatggagcgccaggctgaagcacgccgcctccgagctgaacaacaacgcgttaatgatttaaaaaacaatcagatggagc
aggaacgcctgcttggatggttggctttccatcgcgcctgggaacattcgccagctgaagatggctcctggccttatgcggtaaaaagggaattcgtgacgtatttgg
cgcttgtgatatacgtcgcgtgaccagtcacgacagtgcgaaatggagccgtggaccaacaacttacatggcgatacctctgtcccacctggacggaagaaaagacggg
cggattgttggctggcaacgtatcgatctgaatggtggcaagttccagaccagcgccatcactaacggtgatttcgtcggtgcatgctttgttattggcgacctgcagg
gggcgcaaaaaattgcgactgcagagggtttcgccaccggggcatccatctggctggcgaccagaaacgacccgaaaaaacgctttgacgcggtagtcattgcggtttc
cgctaacaacatgattcacgttgtcgagcagctggtgaacatgtacccggctgcgcaaatcacttgcgctctcgataacgaccgcaaatcctcagctgaaggaaaaggc
aacacaggcctccgcaccggattcgacatcatggagaagttttccggcgtcaaatgtgtttacccaacttttgaggatgaccctgagctggaatgcagcgattttaacg
acctgcacagcctgagagggctgaaggaagtcgctcgccagttaacgagaaatcatctgagccgtgcaaccgatctgttgtcgatcacgctgaataagctgcgtactct
cccgcgtctgaacagacgcacttttgccaaagaactgcttcgcgccgttgatattggcatgctgacatgcccggtaccaaacagcccgaaagaacttatgcgcctttttc
agcagcacgctgcgggatatgggtatcgcagaaatttataacggtaccgttaaagatcacattacgcgccggttgaatcgcaaatgccgtgccgcgcaaacatcacgtt

cgttcagtgaacgcatcaccaacccgaaccttcgcccgtcacacatcacttacaaacggtttgaaacctccaggatgaccgatgaggtgatgacatacgccgcacagct
gcagggcatcgttattgtccgcgccgggatggggtcgggtaaatcgacaggcctcctgcgtccactgatgctgcagtccacgcgtggcgtttccgtcgcgcaccgcgta
tccctgataggcggcctgcatgaaatgatgaccgaagggaaaggcgcgaaagccgacattctgcattatcaggatcccggctatcaggaaatggcgccatatgccaaca
agctgactatttgcatcaactccatacttaaaggctgctggcaaccgctgatgcgccagcatgacttcttcggcttcgatgaagcaacacagggcctgcgtgccattct
ggccgggcgtgcgatggaaaacccggtaggcgtattcaacacgcttatcgacgcgctggcgcgtactgaagagcatgccattatggtggatgccgatgccaacgatctg
cttgttgacctggctgaactggcgatgaagcgacgcgaggagctgggcctacctgcctggctgcaaattcacgtgattgaactcccggtcgacgttcgcaaccgcgaaa
ccaacaagcctatccgcgtattttataccgagaaaaatcggatcatgaccgaggttatcgctgcagtgcaacgcggcgaacgaatcatgctggccaccgacagttcgac
gttcgccgaagacgttaccatgcagctgagactacagttccctgacaaaaagtttctctgcgttaaccagaaaaacaaacaggagaaggaagtcgacgatttcaccaac
cagcctaaagtgatggtgaaaaaatatgacggcctcatctacagcccgtcgatatcttcaggtgtatcgattgaggagaaacacttccaccgccatttcggcatgttct
gcggcgaagtggtccccagcgacgccatccagatgctgcgccgcgaccgtacagctcaggaatacattatcggtttcgacaagcttcgcggtaaacgtgaaaccgatcc
ggaaaaaatcaaacgcgcctacgcgcaggccttgctcgaaacggctggccactccgggctgctgacagacgttgtctttgacggtgaccgaatttcactcggcgtggct
aactcttcattcatgcagctcaaaattaaagccgccgcgctcgaggcatcggccagaaatgattatgccagcaacatgatttgcatcatgcatgatgatggctatcagg
tcgcgcctatggccaccgacgcgcttgcaaacagcatcggtaaggatttacgtaaagaagcccgtgaactggtctttgagcagcttatggagcgccacctgagtgtcga
tactcctgaccaggccgaacacgatgagttgataaaaaaacgcaccctgtccctggatgagcaggcccagctggtccgctgggacatcgaaaaggagctgcagctggat
gtcgacgaggtggcgttaaaattctatttcgacggcggcctgaaaaaagtgcgcctgtttgaaaccatgcagctggatgagataaccgcccgtcgccttgaccgtgaag
aagcgttgattcactttacctacgcttatcgcgttgccggaagatggcagcagttcgtcactacggccatgacgcgggaacaggccgacgcagagttccaggctaaatt
cccggccataaccgattaccgagtgaaatcgaccccctgctgttgagataggcatgcgtggtttctacaccctcaaatcggcgacgctgcagcagtacttccgcgactgc
ggtatcgatccaaaaaccctggaaggtgaagcagacatggacgccctcaaacgcgccagagataacctgctcacgccggaacggcgcgatctgctaaacaacgttttac
gcatcggcggcttcaacacagagaaaggcaagaagaaagcgcctgcagacctctgcataggcatcctggagtctatggggctatccagtaagaccagacgcgccagaga
cggcgatgcacgccccacgatgcgttccattgacccggactcggtcgagttcctcatgaatatcgtggagaagcgccgcgaggccggtttatcaattcacgcacgtaag
gtggaaaaaaccaccatcgaagtggatcgcgatttggatctaaatatagatatacatggtaaccctcgatccaaaacagagcacgttccggacgccccacaatcagtaa
ttatccaggcactggaggctatcccggtggcggtaccggaggcgtgggcggagaacgcgttgccagcaaccgaaatggaggcggtacgcctgtggccagtggccagcat
cgcgagaacgttcgcctcgctgtacatgaccgaatttatggacctgctatcagtacgcgaaataaggctgctgaaagcgtttctaagccagcggcaagccgtggcgata
taaccggaggggtaatgggaaaacaacgggaatgccttgtcatcgtggagggctacgacccttatgtggtcaaggtcagccacgacgcgacaagctatagcgattataa
aactggccgggaacttccattcacgatcattcggatgaacgggttctcagatagtgagggagtttattgctgcattgccagcgataagagcatggaagaccacacagac
gaagccagggcgcttctaaaagcctataaggtgaagccactatgacgtcgacaccagagttcctgaagcaactggattacgaacagctgaagtactgtcgcgagctgtg
cgataaccgcatccggctattgaggccgaagagaaaaagatggcctgggcagttactgatggtggcattaactacggctggttccgtacagaagactacatgaaggcg
gtagaatgcctggttgctacagctgcagaacggtgggaagaatcggataaggctgacccatctggacgtggttggcttgacctttcaatccgtggaagccgcctgccgg
tgtccgagtatgaagctttatttgctgatggccagtggggggtgaatagtggcgcatgatctacaaaaaatgatggatggtgagctggtccgcttacaacttgagattaa
gaaagaacaggagaggcgtttagccgaaaagatgattcctgtttttggtgtgtggcaatcacctattaatcaatatgaatttgctgatccagagcaggcaatttcttgt
gccgcagggctattggataaggttcttcgtgaggtcaaggacgatctttccaaagaaggtggattacagggatggaacggcaatttgctacgcatctacgtacagcatg
tgaacgagagcgattttgcagttctacagccttatttaaaggacaaataatcatgatttatcgcagaggatgggtgcctgttttgtggaggaatgagctggaaaagcgg
ctcaaagaagaagggttcgaaaactggaagcagataagtaactttctttgtggtgaagaggattgctactcggaagcagcaaaaacgccaggacaatacgcttaccagg
tcgtcgacaataccgaatggatggaacgtcgtgatgcgacgttctggcagcgtctgaaccgtttgtggttcgttccgctgtacctgctgactattccgttccagtggct
cattcgtggccgtatggggtttgagacaacatctaaggtggctgcggtcatcaataagattaccggactccaataactgatcaggttctgatcagtcagtaaagcaaac
tataaaaacacctgtgcacgtgactggtcacggggtaaaatatccacctaaattattgacgtgcgttctgctttggcggtagagttaccccgctgcagcaaaatctgca
gccgggcctctcaaccccgaatgacaagaagcgcacaacacgcgccagcgtgtttttttgtgtgttaaatctgcgcatacctgaattatggtggctcagatggggccaa
cttcggttgggccggtttcttcttgtaccggtgttgagaaccctgtctgggctaccacccctatagagattctcaactctggtggtagcaccctatacaagataggaat
gcataccatgttcaaattcaaatttgcagctgtcgtccgcacggacaaaaaatcccatatccaccgcctttccaccattgcatcatccgagcgagaagctcgtcgccag
ttcgccagccgttttgtcctcgttctgtcagcccgtatcccggtcagcgaggtgatcgcatgaaccagattcagttaaacgcccagggcctgctggagacgattgagga
acgcctgctgcaggttgaagcgcttgtgggctctgctcaccgcactatctccagttatgaggccacgctgtatttgcaggaagctgcggaacttctgcaggtcgcgcgc
gaactcacgcaggaagctcgcggctgctccctgtccctttctgagcagctgaaatccggggaggcgcaatgaaggctctctccgtcttttcatttcaggaaagccaccc
tatacgggtggttcttgtcggtggtgacccgtggtttgtggcgctggatatctgtgctgcattgaatatagccaacccttctgatgcattgcgtaagctggatcatgat
gaaaaattgaccctcggtttaaccgaggcacaaaaacttgatcggatggccagggaagtaaatgtggtatcggaatctggtctctacaccataatcctccgctgccgcg
acgcggtgaaacaggggacgacagcctggcggttccgcaagtgggtgacgaacgaggttctgccagctatccggaaaaacggcgagtatgcttttgtcgagcctgagcc
taagaacgccggtgaaccactggactggcggcagaaagaagaacttcggggcctgataaacgacatagcacaaagttttcagtaccgtaacgcatgggttagcggcgtc
tggatggctctccgccgcgcctgcaggaacccatcaccgaacccaataaccgtggatgatctcccggcaatcatcgccgagctgcgccggatattaacggctgcagaaa
cggctctgggcaacatgcgtgtctacgaacgggagctgctgcgcgatgtagttcgtggcggtcgccggagtatgtcttgtggagagctgccgatcaccgatattgatac
ggaactggagaaggtgctgccagcacattttgagctggccattgaaaagctggaaacgctatccacgaaactaacatcgccagcgattgcagaaaaatagggttggcca
gcccgaatttaagccccgctttacgcggggaagtccctgcagtcggacggcgatgtccgtagttgcagctgcagtttccgcctaatccttccgtcccttgcaggggat
gggtagggacaggcctcaataaaaaagccaccgattaacaccggtggcttttcttttagctgtagtacgtttcccatgcggatcgcattgcggccatagggctatcggc
ggggccggaccaggcataatgctttccatcgtattcaaactctatcttgtacgttccgtccccgttatttttttgcaggcttgaagacgggttttagagcagttggctga
tgttcttctggcccttcctcttcggttggttcatcgccgccaccctcgtcgagctcaatttcatcttcatccagctcgtcgtcttgggactcatcatccggttcgtcga
tggtctcaacggattcttcatcaggcaggacgatt**gcTggtg**tctctatcttcagctgccactgcccgttctcgccaacgaactgccccaatgcatcagcggcaaactc
caggtaccggctaatcatcgtcgggctaaatttaaaggcccggagagtgctgttggttatttttgctgatgggtcctgctccaccagctgcttaacggtttcatggaga
cggacgccagcgtcacctctggcaaagcctggcatttcatcgtccagtttctgcagagccaccagcctggtgttttcatccccaacttcaggtcgccaggttctggaga
agttggccagcttgaactgcttatagtgcagctgggtgttctcatcgtcgtgtccgagaatctccatgaagaacacatcctcgtcgacgtttttccaccgtggatcgac
gcggaagaacatctcataagcgatgcgagcgtaaatagcgcggctatctttataaacacgacggtcatcgccgaaaaatgatttaacccaagggttaaatgcttttgct
aaaatagcatttatcctgccgttctcagaccttgtatcatcctttccatatcctttaacaacctcatcgaaatcagatgcagcagagcaagaacgcaattctgttaata
attcaacgaataatttgcttcgcataaagtataaatcgttctggttacgcttttatcttcagagcgtttttagcttgccctgagaaattaaccgtatactttcctga
aacggcaaattcaccctgaaacattatctcaatcattcttcgccctgataccgcagccagagcaaaggccaaaggtgccattccagaacgagtgtttaaactaaataaa
gtcgcaggattattcaaaatatcatagatagactgcatgtatgttgggtagtcaataaccacaacattacgcttcttctcgcgcagaacatcggcccatcgttgctgta
tagatgtacgctccgcagggcttagctgcagatggtacagaacctcatggttgaccttgagctggtgtagttcttctaacaacgcagagccttgttggaataacttata
aagatagtcacggcgctccttccaatcatcactgtttaaatcactaagagcaaaactccaatctggatatttttttattagtcttgctatttttgcatcactgccttta
gagcctattcttacatttgataactcttcggcaagcggcattatttctttcaatttagattgtaacgatgacatgtgctggcgaatattagccgtaggcatagaaagcc
atgaagataattcttcgctgtaaagaggatacttttccgataatttattaatatttttatcaaagctatgatgtaatttatcatcaaaccgctttcttgccctgctcat
ataggcgttaaaagtattcgcggttattcttttctgcaatcctttcccacggaactttcttttatcattaaataacgcgttcttataccgtgcggctgcggctttaatt
ctcttcgttttgtcgccttgtgggcggtctgaggcatcaattgcctctacctcattcacaagcgtgttgatcaactcaccgattttaccttgctcatcccggttcccc
cgtttctaagtctctggatatcgtaacacaatcaaagaacaagtgacgattaattatgctgctttcgtcacactataagagaacataatagcagaataacaaaggaaca
caacaccggaatagatacatattgttctgcgcttatgttctccttatcagcacacaatagtccattatacgc

#### pJazz-V5

See Figure 2E. 37.6 kb fragment (uppercase) excised from humanised BAC via CRISPR/Cas9 digestion and blunt cloned into pJazz-OC. Replacement of landing pad with pure repeat. pJazz backbone in lowercase. Mouse sequence in **green**. Human sequence in **blue**, selection cassette in **magenta**, non-pathogenic repeat in **red**, AarI site mutation indicated by **gcTggtg**.

gcgtataatgggcaattgtgtgctgatatgttccgtaattgagatatgatatagtgcatatgttccgaattattgatacattaatgttgcattcctatgttgtgttctg
ttatggtgttcttagatagtgggggatgtcacatggattgatgttctttgattgtgtttcaatatagagacggaagagatagagatgcctcaaagtgtataaacagcgt
gaaaatcagcatgttcgcttaacttttgatagcaattggtatgatttattatcatttaaaatcattaagttaggttttaaatataccaacgtaagtaataccgttgcgc
ttagatgaagctaaatagcttctgtagcgcatcaaaaaaattgatttgaaccttaactttgcgttttctaagtcactttgctttttttatgttgctttgctttaaaaacg
taaagtatgatgaaaaacacaaagataagggcatgtttgagaggaaacaggatgtcgttaattaatttgctgaaagactgtatcaatcgcggtcaggaaatgacgcgcg
ctattgccatcgcacagttcggcgatgacagcccggaagcgcgtcgtattactcgccgctggggtataacagaagttgcagacctgatcggcgtaacaccgcaggctat
cagagacgcggaaaaagctggtcgtctaccggctcctgattttgagatgaggggccgagtagaacgtcgcgctggctacacaattgaccaaattagccatatgcggagt
gtgtttggtaacccaaaccaaagacctgacgataaaaacccggttgtactttccgttatgtcacacaagggaggggtttataaaacctcatctgcagtacaccaggcgc
aatggttagccttgcaagggcaccgcgtcctactcgttgaagggaacgacccgcaagggactgcatctatgtaccacggttacgttcctgatttgcacattcacgcaga
cgatacactgcttccatttacctgggtaaacgcgacaacgctgaatatgcgataaaaccaacatgctggccgggtctcgacataatccccagctgcctggcccttcac
cgcattgaaacagatctagcagaaagtcaaaagcctccgaccggaggcttttgacttctgtcacctaggttacgccccgccctgccactcatcgcagtactgttgtaat
tcattaagcattctgccgacatggaagccatcacaaacggcatgatgaacctgaatcgccagcggcatcagcaccttgtcgccttgcgtataatatttgcccatggtga
aaacgggggcgaagaagttgtccatattggccacgtttaaatcaaaactggtgaaactcacccagggattggctgagacgaaaaacatattctcaataaaccctttagg
gaaataggccaggttttcaccgtaacacgccacatcttgcgaatatatgtgtagaaactgccggaaatcgtcgtggtattcactccagagcgatgaaaacgtttcagtt
tgctcatggaaaacggtgtaacaagggtgaacactatcccatatcaccagctcaccgtctttcattgccatacgaaattccggatgagcattcatcaggcgggcaagaa
tgtgaataaaggccggataaaacttgtgcttatttttctttacggtctttaaaaaggccgtaatatccagctgaacggtctggttataggtacattgagcaactgactg
aaatgcctcaaaatgttctttacgatgccattgggatatatcaacggtggtatatccagtgatttttttctccattttagcttccttagctcctgaaaatctcgataac
tcaaaaaatacgcccggtagtgatcttatttcattatggtgaaagttggaacctcttacgtgccgatcagattaaaacgaaaggcccagtctttcgactgagcctttcg
ttttatttgaccatgttggtatgatttaaatggtcagttaatcagttctatgtaccagcaaggtccagttgtaagcggccgcttgacttcaagtctaatggcccGCAAG
GACCCTAGACTCAAGAAAATGTAAGGTTACTAAAGTCTGAAGGACACCAAAGTGAAATGACAATTAGCTTTGGTTTGAGGCATATTTGTTTTGTTCGATTTTTCTGTTT
TTTAAAAGCATATAATCAGAAAGAATTATAGAGCTCGATTGACAATTGCAAGATGAACATGTGAAATAGTATTCCTAGGTGTCAGACAAAATAGAAAATGCCAATCATT
CTTAGACAAATGATTTTTATAACACATTAAAATATAATAGGAAAATCTCACAAAATAAGTCTGGGGCAGGAAGATTGCTCCAAGTTTTAGGCCAACCTGGGCTACAGAG
TGATATCCTGTTAGGTGTAGGTGAAGGAAGAGGAATAGGAAAAAAAAAAGACTCAAGGATAACACTTTTTGGTAACTGGAATGACAGTTCAATGAGACATCCGACATATT
CCAATCACAGTCTGCTAAAAAGGAAATGAATCCACAGGTAACCATATGGTCTTGGGTTTTAATTAGTATGTGACAACACTGACAGTCAATTATTCCATACTGCTACTAA
ATACTGAAAACTCTGCTTACTAAATCCCAATAAATGCAATAAAATATCAACATAAACTGCAGAATGGCTGATGGTTACAGGTAGATGGTGACACTTGGCAGAGATGCTC
GGGAGCACATTTGGAGCTGGCCTGAGAAGTGTGTGGTAGGAGAGTGTGCATTACTGCTGTGTCTCCTCCCCATGGGACCAGCGCCATCAGCCTGCACTGCTGGCCGCAT
CTCTGGTCTATCCAAGTATTAGACATGTACAACCACAAATTCTCTCTTCTTCTAAAGAAACACAATTTCATATTCTCTTTATGATTGCACATTAACATCTAAGATTTAA
GATGATGCTTTTGATTTCCTATGTTTTATGAGGTGCATTCCTCCCCCTGAGACTCTATTTGGTTTTTTGTTTGTTTGTTTGTTTGTGTTTGTGTTTGCTTTTGGGTTTT
TTTGTTTTTTTGTTTTGTGGGTTTTTTGGGTTTTTTGTTTGTTTGGTTGGTTGATTGGTTGGTTGTTTTGGTTGTGTCCTTTAACAGCTGAGCCAAGTCTCCAGCCCAG
GCTTCTGTCCTTAGAAATAGTTACATATAAAGGTAAGGTGCACAGCAGCAATGCACAAGAACAACTGGTCAAAGATGACATTCCTCTCACTCACTAAGGACATTGTTCA
TAAAATCACGAGTCCAAAGACAAGCAGAAAAGAAATGTGTATTGTGCCCTAATGACTTTTTTTTTTTGCAGTACTGAAGCAGTATCATGTGCTGTTAGGATTATAAAGT
GTTGGCTCCAGACAGAAACTCAAGGGTTTTCTATATCATTTCATCCTACTGGCTCCACACACTAAGATTCTATTCGTATTTCAAATGAAGGCAGTATATCATAAAATAC
ATATTTATTAAGTCTTCAGTGAATAAGGAAGCAACCACATGAAATTACATTTCCTAAGAGAAATGAAATGGTCCGTGGTTTATATAGCAATATTCTTCATCCATGTCTG
TCATGAAATTGTTCACATGCCAGTCTCAGTCACAACTAATTTCCTGTAATCTTGGTAAGTAGGAGCTAATCAACAAGCCCTTGGTTCCAGCCCTGCAGCAATGCACATC
ACACTTCTGAAACATATTAGCAGAGAAAATGGAGAAAAGCCTTCCTGACAACTGGCATTTTTGATGAAGCAAGAAGGTTCTCCGTGTTCTGACATATTTATTGGAGTTA
TAAATACGTGTAGAGTGTAAGAGGGTTGTAAGTCATTCATTACTGTATTTATTATATCTGAGTAGTTACCAAAAAAAAAAATGAAAGTAAAACCATCACTTCTAGAATG
TACTGATCTAATTTTCTAGCTCGCCTGATTGCACATAAACCTTACAGCAACCAGGTGACAGATTACTAGGCGTGCTATTTTCCTGCTGAGCAACAGTGTCCTGCTAAAA
GACTGCACTGACCAGCTTCCTGTGGGGTCAAGCAAGGAAGCCCCAGAGGAAGGGACTATCAGAGGAGTGCAATGTACCTATGGGAGTCACTTCCACTCCTATTAGAAGG
GCTGCAGCCATTGCTTTCTGTGGGTGTTCTGCCATCCACACACAAGGGGAACAAAATGCCCTTGTCATGGATACACAGTCTTACCTAGGGTCACACTGGCCACTCTCTC
CATTCCCTCCTTTAAGCCCCCACAGAAAGCTAGCACTGTGGACACTGAGTCACCAAGGAGTGAAGAACTGTCCAGTTCCGTGGTTAACCTAAAGCAGAGACGATACTTA
ATATCTGTTGTGGTTACAGAGCCCAACCTCTTATTTTGCTTTAGAAATCCCCCTTATCGAGGAGTGTCTGCATGACTGAACATTAGTGAAACATGATAGCTGACATTTT
TCTATATCGGGTCTTTAAAGGTGTTGTTCTGGTTGGAAATAGGGGATAGAGTTCTCAGTTCCTGTTTGGAGAGATACTCCCTTTCTTTTTCTTTTCTTTCTTTCTTTCT
TTCTTTCTTTCTTTCTTTCTTTCTTTCTTTCTTTCTTTCTTTCTTTTAATGGGATCCCCCCGCCCCCTTGATAGCCTTCAGCATGTAGTGTAGCCTAGGATAATCTTGA
ACTTCTAATCCTCCTGTCTCAGATTATAGGTGTTTGCCATCTCGTCAGTTCGTTTAGTATTGGGGAGTAAACCAGTACGTGGTAGGCAAGCCCTCTGCCAAATGCCCTC
CACTTTCAGCCCTGGAGATATAGTTCTATAGCCAGCATATTTTCACTTAGAGCCTGGAGTTATTTGAGGTAGATTATTTGTAAAGTTTGAGGACAGGGTTAAAAATC
TATCACACCAATATCATGAATCATATTTCTACATACAGTTAGCATCATTTTCTCTTTGGTTCTCTGGAGGAAAGATGTATTCTACCTTACACCTTTCAGGCAATACAAC
AATTATGAACCAGCATACCTACTTATTATTTTCTTCCTGATTAATAAAAGTTAGCTACAATCCAATTAAAATATATAGGCTACTTTACCTATGCCTAAGTAATTGTTCC
ATCTGATGTGAAGAAAAAGGTCTCCACTCATTGATCACAAAATATAAGTTGTATCATGGTTTATTTCATTTCCTGTGGTCATACAATATTTTAGATCGTGATGTTTCAT
TTTTTTCTCCAAAGAGAGAAAAAGAGAGGAATATGGTAGGGACCAAACCCAGGGAGTCACACATGCTAGGCAAGCCCTCTACCCACGAAGCTACATCCCTAGTCCTTTC
TTGGTTTGATTATTTTTTTTAAATAAAACATTTATTCTAGGACTGATATTACACAAAATAAGTTATTTGAGGGACACTGTTTTTGTCTCTAGGTAAAATTTTGAAGGAA
AAAAAAAACACTAAGAAGGTATATTCCTTCAAAGTTCCAGTCTTATTCTGAAGTGTAATGTTATGTTAGTTTGACTCACAGACAGGTTTTAAAGAAGGGCTTACTTCAA
GAGGACACCAAACAAATACCTTCTATTCCTAGTGGGCTCTGGAATCACAGAAAACTGACCCAATCAATTACATTGATAGCTCTGGCTTACTACAGACAAGCAAATTATC
TTAAGTGTGCATGCATGCGCGTGTATGTGTGTTAGTACCTAACACCCACCTGGGAACTTTTCAGCTTTTCAGTGTGGGATATAGTATAAACGTCTATTCCTCGTGTTGT
GGATTAGCTGACTGGCCTCACTCAGCTGCCTTCCTTACCTGCAAACTCACCCACTTTGACTACAGCATCGCACTCTTAACCCTAGCCTTCCAAACATGGTCCTATGCTA
TTTCTGTGTGTCTGGATGTATTTTTAACTCTCAGATGTATACTTCATTTATGAGATATACATCTGAAGACCACGGTACAAAACACTGTAAGAACTTGATAGAATGACAA
CTGCTAGGTAAAAAAAAAAAAAAAAAAAAAAAAAAAAAAAAAAAAAAAAAAGCATACAATACCTGGTGAGAGTTCTATTTTTACCGAAGGTGGTATTGATAGGTATTCTGTTA
TTAATGCCTTTCTTTTCCCTATAAATGATGAAAAGTTGCTGGAAAATAATAAACACTACTCATCTGTAGTGAAAAGCCACAATACAGTTACAAACCAATCAATCAATCA
ATAAATCAGACGTCATGGTGTTCTTTTCCCAAAGGTTAAAAACAAAGTGCACTGTGCTATTTGGCAAAAATGACGTTTAGAAGAAAACACGGTGACTACGCACAGAGG
GTGGGGGAATCATTGTGCTTGTTGCGGAGTGAACACGTACAGTGTGCACGCAGACTTACGGCATTTAACCGTGTCATAGGGACCAAAGGAAATCCACTCACTCACTAAA
TATTTGTTGAGCACCCACTACCTGCCAACTCCCAAACAAAACAAAGCAAAACTACTTACAACCACAAACTACGCTTCGTAACCTAGATAGATAACGCAGGTGACACTAT
CTATCTAGGTTGAGCTCAGCTCTGCCCATGCTTTTCCTGAGCGGCTCTTGGAAGAAAAGCTACAAAGCCCATGACAGCCTCCGCCTGGCCAGCTGCCACTGGCATCTCA
AGGCTGGCAAAGCAAAGTGAAAGCGCCAACCCGGAACTTACGGAGTCCCACGAGGGAACCGCGGCGCGTCAAGCAGAGACGAGTTCCGCCCACGTGAAAGATGGCGTTT
GTAGTGACAGCCATCCCAATTGCCCTTTCCTTCTAGGTGGAAAGTGGTGTCTAGACAGTCCAGGGAGGGTGTGCGAGGGAGGTGCGTTTTGGTTGCCTCAGCTCGCAAC
TTAACTCCACAACGGTGACCAAGGACAAAAGAAGGAAACAAGACTGCAGAGATCCGCACCGGGGAGCCCTGCAGATTCTGGGTGTGTGTGTTTTTGTTTTTCCCACCCTCT
CTCCCCACTACTTGCTCTCACAGTACTCGCTGAGGGTGAACAAGAAAAGACCTGATAAAGATTAACCCTAGAAAGATAGTCTGCGTAAAATTGACGCATGCATTCTTGA
AATATTGCTCTCTCTTTCTAAATAGCGCGAATCCGTCGCTGTGCATTTAGGACATCTCAGTCGCCGCTTGGAGCTCCCGTGAGGCGTGCTTGTCAATGCGGTAAGTGTC
ACTGATTTTGAACTATAACGACCGCGTGAGTCAAAATGACGCATGATTATCTTTTACGTGACTTTTAAGATTTAACTCATACGATAATTATATTGTTATTTCATGTTCT
ACTTACGTGATAACTTATTATATATATATATTTTCTTGTTATAGATATCGCGGCCGCTCTAGCCTCGAGGCTAGAACTAGTGGATCTCGAGCCCCAGCTGGTTCTTTCCGC
CTCAGAAGCCATAGAGCCCACCGCATCCCCAGCATGCCTGCTATTGTCTTCCCAATCCTCCCCCTTGCTGTCCTGCCCCACCCCACCCCCCAGAATAGAATGACACCTA
CTCAGACAATGCGATGCAATTTCCTCATTTTATTAGGAAAGGACAGTGGGAGTGGCACCTTCCAGGGTCAAGGAAGGCACGGGGGAGGGGCAAACAACAGATGGCTGGC
AACTAGAAGGCACAGTCGAGGCTGATCAGCGAGCTCTAGAGCTCAGAAGAACTCGTCAAGAAGGCGATAGAAGGCGATGCGCTGCGAATCGGGAGCGGCGATACCGTAA
AGCACGAGGAAGCGGTCAGCCCATTCGCCGCCAAGCTCTTCAGCAATATCACGGGTAGCCAACGCTATGTCCTGATAGCGGTCCGCCACACCCAGCCGGCCACAGTCGA
TGAATCCAGAAAAGCGGCCATTTTCCACCATGATATTCGGCAAGCAGGCATCGCCATGGGTCACGACGAGATCCTCGCCGTCGGGCATGCGCGCCTTGAGCCTGGCGAA
CAGTTCGGCTGGCGCGAGCCCCTGATGCTCTTCGTCCAGATCATCCTGATCGACAAGACCGGCTTCCATCCGAGTACGTGCTCGCTCGATGCGATGTTTCGCTTGGTGG
TCGAATGGGCAGGTAGCCGGATCAAGCGTATGCAGCCGCCGCATTGCATCAGCCATGATGGATACTTTCTCGGCAGGAGCAAGGTGAGATGACAGGAGATCCTGCCCCG
GCACTTCGCCCAATAGCAGCCAGTCCCTTCCCGCTTCAGTGACAACGTCGAGCACAGCTGCGCAAGGAACGCCCGTCGTGGCCAGCCACGATAGCCGCGCTGCCTCGTC
CTGCAGTTCATTCAGGGCACCGGACAGGTCGGTCTTGACAAAAAGAACCGGGCGCCCCTGCGCTGACAGCCGGAACACGGCGGCATCAGAGCAGCCGATCGTCTGTTGT
GCCCAGTCATAGCCGAATAGCCTCTCCACCCAAGCGGCCGGAGAACCTGCGTGCAATCCATCTTGTTCAATGGCCGATCCCATGGTTTAGTTCCTCACCTTGTCGTATT

ATACTATGCCGATATACTATGCCGATGATTAATTGTCAACAGGCTGCAGGTCGAAAGGCCCGGAGATGAGGAAGAGGAGAACAGCGCGGCAGACGTGCGCTTTTGAAGC
GTGCAGAATGCCGGGCCTCCGGAGGACCTTCGGGCGCCCGCCCCGCCCCTGAGCCCGCCCCTGAGCCCGCCCCCGGACCCACCCCTTCCCAGCCTCTGAGCCCAGAAAG
CGAAGGAGCAAAGCTGCTATTGGCCGCTGCCCCAAAGGCCTACCCGCTTCCATTGCTCAGCGGTGCTGTCCATCTGCACGAGACTAGTGAGACGTGCTACTTCCATTTG
TCACGTCCTGCACGACGCGAGCTGCGGGGCGGGGGGAACTTCCTGACTAGGGGAGGAGTAGAAGGTGGCGCGAAGGGGCCACCAAAGAACGGAGCCGGTTGGCGCCTA
CCGGTGGATGTGGAATGTGTGCGAGGCCAGAGGCCACTTGTGTAGCGCCAAGTGCCCAGCGGGGCTGCTAAAGCGCATGCTCCAGACTGCCTTGGGAAAAGCGCCTCCC
CTACCCGGTAGAATTAATTCGATATCAAGCTTGCTCGGATGATAGGGCCCTAACGTGTTTTTGCTTTGTTACTTTATAGAAGAAATTTTGAGTTTTTGTTTTTTTTTAA
TAAATAAATAAACATAAATAAATTGTTTGTTGAATTTATTATTAGTATGTAAGTGTAAATATAATAAAACTTAATATCTATTCAAATTAATAAATAAACCTCGATATAC
AGACCGATAAAACACATGCGTCAATTTTACGCATGATTATCTTTAACGTACGTCACAATATGATTATCTTTCTAGGGTTAACCAGAAGAAAACAAGGAGGGAAACAACC
GCAGCCTGTAGCAAGCTCTGGAACTCAGGAGTCGCGCGCTAGGGGCCGGGGCCGGGGCCGGGGCGGGGCTGCGGTTGCGGTGCCTGCGCCCGCGGCGGCGGAGGCGCAG
GCGGTGGCGAGTGGGTGAGTGAGGAGGCGGCATCCTGGCGGGTGGCTGTTTGGGGTTCGGCTGCCGGGAAGAGGCGCGGGTAGAAGCGGGGGCTCTCCTCAGAGCTCGA
CGCATTTTTACTTTCCCTCTCATTTCTCTGACCGAAGCTGGGTGTCGGGCTTTCGCCTCTAGCGACTGGTGGAATTGCCTGCATCCGGGCCCCGGGCTTCCCGGCGGCG
GCGGCGGCGGCGGCGGCGCAGGGACAAGGGATGGGGATCTGGCCTCTTCCTTGCTTTCCCGCCCTCAGTACCCGAGCTGTCTCCTTCCCGGGGACCCGCTGGGAGCGCT
GCCGCTGCGGGCTCGAGAAAAGGGAGCCTCGGGTACTGAGAGGCCTCGCCTGGGGGAAGGCCGGAGGGTGGGCGGCGCGCGGCTTCTGCGGACCAAGTCGGGGTTCGCT
AGGAACCCGAGACGGTCCCTGCCGGCGAGGAGATCATGCGGGATGAGATGGGGGTGTGGAGACGCCTGCACAATTTCAGCCCAAGCTTCTAGAGAGTGGTGATGACTTG
CATATGAGGGCAGCAATGCAAGTCGGTGTGCTCCCCATTCTGTGGGACATGACCTGGTTGCTTCACAGCTCCGAGATGACACAGACTTGCTTAAAGGAAGTGACTATTG
TGACTTGGGCATCACTTGACTGATGGTAATCAGTTGTCTAAAGAAGTGCACAGATTACATGTCCGTGTGCTCATTGGGTCTATCTGGCCGCGTTGAACACCACCAGGCT
TTGTATTCAGAAACAGGAGGGAGGTCCTGCACTTTCCCAGGAGGGGTGGCCCTTTCAGATGCAATCGAGATTGTTAGGCTCTGGGAGAGTAGTTGCCTGGTTGTGGCAG
TTGGTAAATTTCTATTCAAACAGTTGCCATGCACCAGTTGTTCACAACAAGGGTACGTAATCTGTCTGGCATTACTTCTACTTTTGTACAAAGGATCAAAAAAAAAAAA
GATACTGTTAAGATATGATTTTTCTCAGACTTTGGGAAACTTTTAACATAATCTGTGAATATCACAGAAACAAGACTATCATATAGGGGATATTAATAACCTGGAGTCA
GAATACTTGAAATACGGTGTCATTTGACACGGGCATTGTTGTCACCACCTCTGCCAAGGCCTGCCACTTTAGGAAAACCCTGAATCAGTTGGAAACTGCTACATGCTGA
TAGTACATCTGAAACAAGAACGAGAGTAATTACCACATTCCAGATTGTTCACTAAGCCAGCATTTACCTGCTCCAGGAAAAAATTACAAGCACCTTATGAAGTTGATAA
AATATTTTGTTTGGCTATGTTGGCACTCCACAATTTGCTTTCAGAGAAACAAAGTAAACCAAGGAGGACTTCTGTTTTTCAAGTCTGCCCTCGGGTTCTATTCTACGTT
AATTAGATAGTTCCCAGGAGGACTAGGTTAGCCTACCTATTGTCTGAGAAACTTGGAACTGTGAGAAATGGCCAGATAGTGATATGAACTTCACCTTCCAGTCTTCCCT
GATGTTGAAGATTGAGAAAGTGTTGTGAACTTTCTGGTACTGTAAACAGTTCACTGTCCTTGAAGTGGTCCTGGGCAGCTCCTGTTGTGGAAAGTGGACGGTTTAGGAT
CCTGCTTCTCTTTGGGCTGGGAGAAAATAAACAGCATGGTTACAAGTATTGAGAGCCAGGTTGGAGAAGGTGGCTTACACCTGTAATGCCAGAGCTTTGGGAGGCGGAG
GCAAGAGGATCACTTGAAGCCAGGAGTTCAAGCTCAACCTGGGCAACGTAGACCCTGTCTCTACAAAAAATTAAAAACTTAGCCGGGCGTGGTGATGTGCACCTGTAGT
CCTAGCTACTTGGGAGGCTGAGGCAGGAGGGTCATTTGAGCCCAAGAGTTTGAAGTTACCGAGAGCTATGATCCTGCCAGTGCATTCCAGCCTGGATGACAAAACGAGA
CCCTGTCTCTAAAAAACAAGAAGTGAGGGCTTTATGATTGTAGAATTTTCACTACAATAGCAGTGGACCAACCACCTTTCTAAATACCAATCAGGGAAGAGATGGTTGA
TTTTTTAACAGACGTTTAAAGAAAAAGCAAAACCTCAAACTTAGCACTCTACTAACAGTTTTAGCAGATGTTAATTAATGTAATCATGTCTGCATGTATGGGATTATTT
CCAGAAAGTGTATTGGGAAACCTCTCATGAACCCTGTGAGCAAGCCACCGTCTCACTCAATTTGAATCTTGGCTTCCCTCAAAAGACTGGCTAATGTTTGGTAACTCTC
TGGAGTAGACAGCACTACATGTACGTAAGATAGGTACATAAACAACTATTGGTTTTGAGCTGATTTTTTTCAGCTGCATTTGCATGTATGGATTTTTCTCACCAAAGAC
GATGACTTCAAGTATTAGTAAAATAATTGTACAGCTCTCCTGATTATACTTCTCTGTGACATTTCATTTCCCAGGCTATTTCTTTTGGTAGGATTTAAAACTAAGCAAT
TCAGTATGATCTTTGTCCTTCATTTTCTTTCTTATTCTTTTTGTTTGTTTGTTTGTTTGTTTTTTTCTTGAGGCAGAGTCTCTCTCTGTCGCCCAGGCTGGAGTGCAGT
GGCGCCATCTCAGCTCATTGCAACCTCTGCCACCTCCGGGTTCAAGAGATTCTCCTGCCTCAGCCTCCCGAGTAGCTGGGATTACAGGTGTCCACCACCACACCCGGCT
AATTTTTTGTATTTTTAGTAGAGGTGGGGTTTCACCATGTTGGCCAGGCTGGTCTTGAGCTCCTGACCTCAGGTGATCCACCTGCCTCGGCCTACCAAAGAGCTGGGAT
AACAGGTGTGACCCACCATGCCCGGCCCATTTTTTTTTTCTTATTCTGTTAGGAGTGAGAGTGTAACTAGCAGTATAATAGTTCAATTTTCACAACGTGGTAAAAGTTT
CCCTATAATTCAATCAGATTTTGCTCCAGGGTTCAGTTCTGTTTTAGGAAATACTTTTATTTTCAGTTTAATGATGAAATATTAGAGTTGTAATATTGCCTTTATGATT
ATCCACCTTTTTAACCTAAAAGAATGAAAGAAAAATATGTTTGCAATATAATTTTATGGTTGTATGTTAACTTAATTCATTATGTTGGCCTCCAGTTTGCTGTTGTTAG
TTATGACAGCAGTAGTGTCATTACCATTTCAATTCAGATTACATTCCTATATTTGATCATTGTAAACTGACTGCTTACATTGTATTAAAAACAGTGGATATTTTAAAGA
AGCTGTACGGCTTATATCTAGTGCTGTCTCTTAAGACTATTAAATTGATACAACATATTTAAAAGTAAATATTACCTAAATGAATTTTTGAAATTACAAATACACGTGT
TAAAACTGTCGTTGTGTTCAACCATTTCTGTACATACTTAGAGTTAACTGTTTTGCCAGGCTCTGTATGCCTACTCATAATATGATAAAAGCACTCATCTAATGCTCTG
TAAATAGAAGTCAGTGCTTTCCATCAGACTGAACTCTCTTGACAAGATGTGGATGAAATTCTTTAAGTAAAATTGTTTACTTTGTCATACATTTACAGATCAAATGTTA
GCTCCCAAAGCAATCATATGGCAAAGATAGGTATATCATAGTTTGCCTATTAGCTGCTTTGTATTGCTATTATTATAAATAGACTTCACAGTTTTAGACTTGCTTAGGT
GAAATTGCAATTCTTTTTACTTTCAGTCTTAGATAACAAGTCTTCAATTATAGTACAATCACACATTGCTTAGGAATGCATCATTAGGCGATTTTGTCATTATGCAAAC
ATCATAGAGTGTACTTACACAAACCTAGATAGTATAGCCTTTATGTACCTAGGCCGTATGGTATAGTCTGTTGCTCCTAGGCCACAAACCTGTACAACTGTTACTGTAC
TGAATACTATAGACAGTTGTAACACAGTGGTAAATATTTATCTAAATATATGCAAACAGAGAAAAGGTACAGTAAAAGTATGGTATAAAAGATAATGGTATACCTGTGT
AGGCCACTTACCACGAATGGAGCTTGCAGGACTAGAAGTTGCTCTGGGTGAGTCAGTGAGTGAGTGGTGAATTAATGTGAAGGCCTAGAACACTGTACACCACTGTAGA
CTATAAACACAGTACGCTGAAGCTACACCAAATTTATCTTAACAGTTTTTCTTCAATAAAAAATTATAACTTTTTAACTTTGTAAACTTTTTAATTTTTTAACTTTTAA
AATACTTAGCTTGAAACACAAATACATTGTATAGCTATACAAAAATATTTTTTCTTTGTATCCTTATTCTAGAAGCTTTTTTCTATTTTCTATTTTAAATTTTTTTTTT
TACTTGTTAGTCGTTTTTGTTAAAAACTAAAACACACACACTTTCACCTAGGCATAGACAGGATTAGGATCATCAGTATCACTCCCTTCCACCTCACTGCCTTCCACCT
CCACATCTTGTCCCACTGGAAGGTTTTTAGGGGCAATAACACACATGTAGCTGTCACCTATGATAACAGTGCTTTCTGTTGAATACCTCCTGAAGGACTTGCCTGAGGC
TGTTTTACATTTAACTTAAAAAAAAAAAAAGTAGAAGGAGTGCACTCTAAAATAACAATAAAAGGCATAGTATAGTGAATACATAAACCAGCAATGTAGTAGTTTATTA
TCAAGTGTTGTACACTGTAATAATTGTATGTGCTATACTTTAAATAACTTGCAAAATAGTACTAAGACCTTATGATGGTTACAGTGTCACTAAGGCAATAGCATATTTT
CAGGTCCATTGTAATCTAATGGGACTACCATCATATATGCAGTCTACCATTGACTGAAACGTTACATGGCACATAACTGTATTTGCAAGAATGATTTGTTTTACATTAA
TATCACATAGGATGTACCTTTTTAGAGTGGTATGTTTATGTGGATTAAGATGTACAAGTTGAGCAAGGGGACCAAGAGCCCTGGGTTCTGTCTTGGATGTGAGCGTTTA
TGTTCTTCTCCTCATGTCTGTTTTCTCATTAAATTCAAAGGCTTGAACGGGCCCTATTTAGCCCTTCTGTTTTCTACGTGTTCTAAATAACTAAAGCTTTTAAATTCTA
GCCATTTAGTGTAGAACTCTCTTTGCAGTGATGAAATGCTGTATTGGTTTCTTGGCTAGCATATTAAATATTTTTATCTTTGTCTTGATACTTCAATGTCGTTTTAAAC
ATCAGGATCGGGCTTCAGTATTCTCATAACCAGAGAGTTCACTGAGGATACAGGACTGTTTGCCCATTTTTTGTTATGGCTCCAGACTTGTGGTATTTCCATGTCTTTT
TTTTTTTTTTTTTTTTGACCTTTTAGCGGCTTTAAAGTATTTCTGTTGTTAGGTGTTGTATTACTTTTCTAAGATTACTTAACAAAGCACCACAAACTGAGTGGCTTT
AAACAACAGCAATTTATTCTCTCACAATTCTAGAAGCTAGAAGTCCGAAATCAAAGTGTTGACAGGGGCATGATCTTCAAGAGAGAAGACTCTTTCCTTGCCTCTTCCT
GGCTTCTGGTGGTTACCAGCAATCCTGAGTGTTCCTTTCTTGCCTTGTAGTTTCAACAATCCAGTATCTGCCTTTTGTCTTCACATGGCTGTCTACCATTTGTCTCTGT
GTCTCCAAATCTCTCTCCTTATAAACACAGCAGTTATTGGATTAGGCCCCACTCTAATCCAGTATGACCCCATTTTAACATGATTACACTTATTTCTAGATAAGGTCAC
ATTCACGTACACCAAGGGTTAGGAATTGAACATATCTTTTTGGGGGACACAATTCAACCCACAAGTGTCAGTCTCTAGCTGAGCCTTTCCCTTCCTGTTTTTCTCCTTT
TTAGTTGCTATGGGTTAGGGGCCAAATCTCCAGTCATACTAGAATTGCACATGGACTGGATATTTGGGAATACTGCGGGTCTATTCTATGAGCTTTAGTATGTAACATT
TAATATCAGTGTAAAGAAGCCCTTTTTTAAGTTATTTCTTTGAATTTCTAAATGTATGCCCTGAATATAAGTAACAAGTTACCATGTCTTGTAAAATGATCATATCAAC
AAACATTTAATGTGCACCTACTGTGCTAGTTGAATGTCTTTATCCTGATAGGAGATAACAGGATTCCACATCTTTGACTTAAGAGGACAAACCAAATATGTCTAAATCA
TTTGGGGTTTTGATGGATATCTTTAAATTGCTGAACCTAATCATTGGTTTCATATGTCATTGTTTAGATATCTCCGGAGCATTTGGATAATGTGACAGTTGGAATGCAG
TGATGTCGACTCTTTGCCCACCGCCATCTCCAGCTGTTGCCAAGACAGAGATTGCTTTAAGTGGCAAATCACCTTTATTAGCAGCTACTTTTGCTTACTGGGACAATAT
TCTTGGTCCTAGAGTAAGGCACATTTGGGCTCCAAAGACAGAACAGGTACTTCTCAGTGATGGAGAAATAACTTTTCTTGCCAACCACACTCTAAATGGAGAAATCCTT
CGAAATGCAGAGAGTGGTGCTATAGATGTAAAGTTTTTTGTCTTGTCTGAAAAGGGAGTGATTATTGTTTCATTAATCTTTGATGGAAACTGGAATGGGGATCGCAGCA
CATATGGACTATCAATTATACTTCCACAGACAGAACTTAGTTTCTACCTCCCACTTCATAGAGTGTGTGTTGATAGATTAACACATATAATCCGGAAAGGAAGAATATG
GATGCATAAGGTAAGTGATTTTTCAGCTTATTAATCATGTTAACCTATCTGTTGAAAGCTTATTTTCTGGTACATATAAATCTTATTTTTTTAATTATATGCAGTGAAC
ATCAAACAATAAATGTTATTTATTTTGCATTTACCCTATTAGATACAAATACATCTGGTCTGATACCTGTCATCTTCATATTAACTGTGGAAGGTACGAAATGGTAGCT
CCACATTATAGATGAAAAGCTAAAGCTTAGACAAATAAAGAAACTTTTAGACCCTGGATTCTTCTTGGGAGCCTTTGACTCTAATACCTTTTGTTTCCCTTTCATTGCA
CAATTCTGTCTTTTGCTTACTACTATGTGTAAGTATAACAGTTCAAAGTAATAGTTTCATAAGCTGTTGGTCATGTAGCCTTTGGTCTCTTTAACCTCTTTGCCAAGTT
CCCAGGTTCATAAAATGAGGAGGTTGAATGGAATGGTTCCCAAGAGAATTCCTTTTAATCTTACAGAAATTATTGTTTTCCTAAATCCTGTAGTTGAATATATAATGCT
ATTTACATTTCAGTATAGTTTTGATGTATCTAAAGAACACATTGAATTCTCCTTCCTGTGTTCCAGTTTGATACTAACCTGAAAGTCCATTAAGCATTACCAGTTTTAA
AAGGCTTTTGCCCAATAGTAAGGAAAAATAATATCTTTTAAAAGAATAATTTTTTACTATGTTTGCAGGCTTACTTCCTTTTTTCTCACATTATGAAACTCTTAAAATC

AGGAGAATCTTTTAAACAACATCATAATGTTTAATTTGAAAAGTGCAAGTCATTCTTTTCCTTTTTGAAACTATGCAGATGTTACATTGACTGTTTTCTGTGAAGTTAT
CTTTTTTTCACTGCAGAATAAAGGTTGTTTTGATTTTATTTTGTATTGTTTATGAGAACATGCATTTGTTGGGTTAATTTCCTACCCCTGCCCCCATTTTTTCCCTAAA
GTAGAAAGTATTTTTCTTGTGAACTAAATTACTACACAAGAACATGTCTATTGAAAAATAAGCAAGTATCAAAATGTTGTGGGTTGTTTTTTTAAATAAATTTTCTCTT
GCTCAGGAAAGACAAGAAAATGTCCAGAAGATTATCTTAGAAGGCACAGAGAGAATGGAAGATCAGGTATATGCAAATTGCATACTGTCAAATGTTTTTCTCACAGCAT
GTATCTGTATAAGGTTGATGGCTACATTTGTCAAGGCCTTGGAGACATACGAATAAGCCTTTAATGGAGCTTTTATGGAGGTGTACAGAATAAACTGGAGGAAGATTTC
CATATCTTAAACCCAAAGAGTTAAATCAGTAAACAAAGGAAAATAGTAATTGCATCTACAAATTAATATTTGCTCCCTTTTTTTTTCTGTTTGCCCAGAATAAATTTTG
GATAACTTGTTCATAGTAAAAATAAAAAAAATTGTCTCTGATATGTTCTTTAAGGTACTACTTCTCGAACCTTTCCCTAGAAGTAGCTGTAACAGAAGGAGAGCATATG
TACCCCTGAGGTATCTGTCTGGGGTGTAGGCCCAGGTCCACACAATATTTCTTCTAAGTCTTATGTTGTATCGTTAAGACTCATGCAATTTACATTTTATTCCATAACT
ATTTTAGTATTAAAATTTGTCAGTGATATTTCTTACCCTCTCCTCTAGGAAAATGTGCCATGTTTATCCCTTGGCTTTGAATGCCCCTCAGGAACAGACACTAAGAGTT
TGAGAAGCATGGTTACAAGGGTGTGGCTTCCCCTGCGGAAACTAAGTACAGACTATTTCACTGTAAAGCAGAGAAGTTCTTTTGAAGGAGAATCTCCAGTGAAGAAAGA
GTTCTTCACTTTTACTTCCATTTCCTCTTGTGGGTGACCCTCAATGCTCCTTGTAAAACTCCAATATTTTAAACATGGCTGTTTTGCCTTTCTTTGCTTCTTTTTAGCA
TGAATGAGACAGATGATACTTTAAAAAAGTAATTAAAAAAAAAAAACTTGTGAAAATACATGGCCATAATACAGAACCCAATACAATGATCTCCTTTACCAAATTGTTAT
GTTTGTACTTTTGTAGATAGCTTTCCAATTCAGAGACAGTTATTCTGTGTAAAGGTCTGACTTAACAAGAAAAGATTTCCCTTTACCCAAAGAATCCCAGTCCTTATTT
GCTGGTCAATAAGCAGGGTCCCCAGGAATGGGGTAACTTTCAGCACCCTCTAACCCACTAGTTATTAGTAGACTAATTAAGTAAACTTATCGCAAGTTGAGGAAACTTA
GAACCAACTAAAATTCTGCTTTTACTGGGATTTTGTTTTTTCAAACCAGAAACCTTTACTTAAGTTGACTACTATTAATGAATTTTGGTCTCTCTTTTAAGTGCTCTTC
TTAAAAATGTTATCTTACTGCTGAGAAGTTCAAGTTTGGGAAGTACAAGGAGGAATAGAAACTTAAGAGATTTTCTTTTAGAGCCTCTTCTGTATTTAGCCCTGTAGGA
TTTTTTTTTTTTTTTTTTTTTTGGTGTTGTTGAGCTTCAGTGAGGCTATTCATTCACTTATACTGATAATGTCTGAGATACTGTGAATGAAATACTATGTATGCTTAA
ACCTAAGAGGAAATATTTTCCCAAAATTATTCTTCCCGAAAAGGAGGAGTTGCCTTTTGATTGAGTTCTTGCAAATCTCACAACGACTTTATTTTGAACAATACTGTTT
GGGGATGATGCATTAGTTTGAAACAACTTCAGTTGTAGCTGTCATCTGATAAAATTGCTTCACAGGGAAGGAAATTTAACACGGATCTAGTCATTATTCTTGTTAGATT
GAATGTGTGAATTGTAATTGTAAACAGGCATGATAATTATTACTTTAAAAACTAAAAACAGTGAATAGTTAGTTGTGGAGGTTACTAAAGGATGGTTTTTTTTTAAATA
AAACTTTCAGCATTATGCAAATGGGCATATGGCTTAGGATAAAACTTCCAGAAGTAGCATCACATTTAAATTCTCAAGCAACTTAATAATATGGGGCTCTGAAAAACTG
GTTAAGGTTACTCCAAAAATGGCCCTGGGTCTGACAAAGATTCTAACTTAAAGATGCTTATGAAGACTTTGAGTAAAATCATTTCATAAAATAAGTGAGGAAAAACAAC
TAGTATTAAATTCATCTTAAATAATGTATGATTTAAAAAATATGTTTAGCTAAAAATGCATAGTCATTTGACAATTTCATTTATATCTCAAAAAATTTACTTAACCAAG
TTGGTCACAAAACTGATGAGACTGGTGGTGGTAGTGAATAAATGAGGGACCATCCATATTTGAGACACTTTACATTTGTGATGTGTTATACTGAATTTTCAGTTTGATT
CTATAGACTACAAATTTCAAAATTACAATTTCAAGATGTAATAAGTAGTAATATCTTGAAATAGCTCTAAAGGGAATTTTTCTGTTTTATTGATTCTTAAAATATATGT
GCTGATTTTGATTTGCATTTGGGTAGATTATACTTTTATGAGTATGGAGGTTAGGTATTGATTCAAGTTTTCCTTACCTATTTGGTAAGGATTTCAAAGTCTTTTTGTG
CTTGGTTTTCCTCATTTTTAAATATGAAATATATTGATGACCTTTAACAAATTTTTTTTATCTCAAATTTTAAAGGAGATCTTTTCTAAAAGAGGCATGATGACTTAAT
CATTGCATGTAACAGTAAACGATAAACCAATGATTCCATACTCTCTAAAGAATAAAAGTGAGCTTTAGGGCCGGGCATGGTCAGAAATTTGACACCAACCTGGCCAACA
TGGCGAAACCCCGTCTCTACTAAAAATACAAAAATCAGCCGGGCATGGTGGCGGCACCTATAGTCCCAGCTACTTGGGAGGATGAGACAGGAGAGTCACTTGAACCTGG
GAGGAGAGGTTGCAGTGAGCTGAGATCACGCCATTGCACTCCAGCCTGAGCAATGAAAGCAAAACTCCATCTCAAAAAAAAAAAAAAGAAAAGAAAGAATAAAAGTGAGC
TTTGGATTGCATATAAATCCTTTAGACATGTAGTAGACTTGTTTGATACTGTGTTTGAACAAATTACGAAGTATTTTCATCAAAGAATGTTATTGTTTGATGTTATTTT
TATTTTTTATTGCCCAGCTTCTCTCATATTACGTGATTTTCTTCACTTCATGTCACTTTATTGTGCAGGGTCAGAGTATTATTCCAATGCTTACTGGAGAAGTGATTCC
TGTAATGGAACTGCTTTCATCTATGAAATCACACAGTGTTCCTGAAGAAATAGATGTAAGTTTAAATGAGAGCAATTATACACTTTATGAGTTTTTTGGGGTTATAGTA
TTATTATGTATATTATTAATATTCTAATTTTAATAGTAAGGACTTTGTCATACATACTATTCACATACAGTATTAGCCACTTTAGCAAATAAGCACACACAAAATCCTG
GATTTTATGGCAAAACAGAGGCATTTTTGATCAGTGATGACAAAATTAAATTCATTTTGTTTATTTCATTACTTTTATAATTCCTAAAAGTGGGAGGATCCCAGCTCTT
ATAGGAGCAATTAATATTTAATGTAGTGTCTTTTGAAACAAAACTGTGTGCCAAAGTAGTAACCATTAATGGAAGTTTACTTGTAGTCACAAATTTAGTTTCCTTAATC
ATTTGTTGAGGACGTTTTGAATCACACACTATGAGTGTTAAGAGATACCTTTAGGAAACTATTCTTGTTGTTTTCTGATTTTGTCATTTAGGTTAGTCTCCTGATTCTG
ACAGCTCAGAAGAGGAAGTTGTTCTTGTAAAAAATTGTTTAACCTGCTTGACCAGCTTTCACATTTGTTCTTCTGAAGTTTATGGTAGTGCACAGAGATTGTTTTTTGGG
GAGTCTTGATTCTCGGAAATGAAGGCAGTGTGTTATATTGAATCCAGACTTCCGAAAACTTGTATATTAAAAGTGTTATTTCAACACTATGTTACAGCCAGACTAATTT
TTTTATTTTTTGATGCATTTTAGATAGCTGATACAGTACTCAATGATGATGATATTGGTGACAGCTGTCATGAAGGCTTTCTTCTCAAGTAAGAATTTTTCTTTTCATA
AAAGCTGGATGAAGCAGATACCATCTTATGCTCACCTATGACAAGATTTGGAAGAAAGAAAATAACAGACTGTCTACTTAGATTGTTCTAGGGACATTACGTATTTGAA
CTGTTGCTTAAATTTGTGTTATTTTTCACTCATTATATTTCTATATATATTTGGTGTTATTCCATTTGCTATTTAAAGAAACCGAGTTTCCATCCCAGACAAGAAATCA
TGGCCCCTTGCTTGATTCTGGTTTCTTGTTTTACTTCTCATTAAAGCTAACAGAATCCTTTCATATTAAGTTGTACTGTAGATGAACTTAAGTTATTTAGGCGTAGAAC
AAAATTATTCATATTTATACTGATCTTTTTCCATCCAGCAGTGGAGTTTAGTACTTAAGAGTTTGTGCCCTTAAACCAGACTCCCTGGATTAATGCTGTGTACCCGTGG
GCAAGGTGCCTGAATTCTCTATACACCTATTTCCTCATCTGTAAAATGGCAATAATAGTAATAGTACCTAATGTGTAGGGTTGTTATAAGCATTGAGTAAGATAAATAA
TATAAAGCACTTAGAACAGTGCCTGGAACATAAAAACACTTAATAATAGCTCATAGCTAACATTTCCTATTTACATTTCTTCTAGAAATAGCCAGTATTTGTTGAGTGC
CTACATGTTAGTTCCTTTACTAGTTGCTTTACATGTATTATCTTATATTCTGTTTTAAAGTTTCTTCACAGTTACAGATTTTCATGAAATTTTACTTTTAATAAAAGAG
AAGTAAAAGTATAAAGTATTCACTTTTATGTTCACAGTCTTTTCCTTTAGGCTCATGATGGAGTATCAGAGGCATGAGTGTGTTTAACCTAAGAGCCTTAATGGCTTGA
ATCAGAAGCACTTTAGTCCTGTATCTGTTCAGTGTCAGCCTTTCATACATCATTTTAAATCCCATTTGACTTTAAGTAAGTCACTTAATCTCTCTACATGTCAATTTCT
TCAGCTATAAAATGATGGTATTTCAATAAATAAATACATTAATTAAATGATATTATACTGACTAATTGGGCTGTTTTAAGGCTCAATAAGAAAATTTCTGTGAAAGGTC
TCTAGAAAATGTAGGTTCCTATACAAATAAAAGATAACATTGTGCTTATAGCTTCGGTGTTTATCATATAAAGCTATTCTGAGTTATTTGAAGAGCTCACCTACTTTTT
TTTGTTTTTAGTTTGTTAAATTGTTTTATAGGCAATGTTTTTAATCTGTTTTCTTTAACTTACAGTGCCATCAGCTCACACTTGCAAACCTGTGGCTGTTCCGTTGTAG
TAGGTAGCAGTGCAGAGAAAGTAAATAAGGTAGTTTATTTTATAATCTAGCAAATGATTTGACTCTTTAAGACTGATGATATATCATGGATTGTCATTTAAATGGTAGG
TTGCAATTAAAATGATCTAGTAGTATAAGGAGGCAATGTAATCTCATCAAATTGCTAAGACACCTTGTGGCAACAGTGAGTTTGAAATAAACTGAGTAAGAATCATTTA
TCAGTTTATTTTGATAGCTCGGAAATACCAGTGTCAGTAGTGTATAAATGGTTTTGAGAATATATTAAAATCAGATATATAAAAAAAATTACTCTTCTATTTCCCAATG
TTATCTTTAACAAATCTGAAGATAGTCATGTACTTTTGGTAGTAGTTCCAAAGAAATGTTATTTGTTTATTCATCTTGATTTCATTGTCTTCGCTTTCCTTCTAAATCT
GTCCCTTCTAGGGAGCTATTGGGATTAAGTGGTCATTGATTATTATACTTTATTCAGTAATGTTTCTGACCCTTTCCTTCAGTGCTACTTGAGTTAATTAAGGATTAAT
GAACAGTTACATTTCCAAGCATTAGCTAATAAACTAAAGGATTTTGCACTTTTCTTCACTGACCATTAGTTAGAAAGAGTTCAGAGATAAGTATGTGTATCTTTCAATT
TCAGCAAACCTAATTTTTTAAAAAAAGTTTTACATAGGAAATATGTTGGAAATGATACTTTACAAAGATATTCATAATTTTTTTTTGTAATCAGCTACTTTGTATATTT
ACATGAGCCTTAATTTATATTTCTCATATAACCATTTATGAGAGCTTAGTATACCTGTGTCATTATATTGCATCTACGAACTAGTGACCTTATTCCTTCTGTTACCTCA
AACAGGTGGCTTTCCATCTGTGATCTCCAAAGCCTTAGGTTGCACAGAGTGACTGCCGAGCTGCTTTATGAAGGGAGAAAGGCTCCATAGTTGGAGTGTTTTTTTTTTT
TTTTTTAAACATTTTTCCCATCCTCCATCCTCTTGAGGGAGAATAGCTTACCTTTTATCTTGTTTTAATTTGAGAAAGAAGTTGCCACCACTCTAGGTTGAAAACCACT
CCTTTAACATAATAACTGTGGATATGGTTTGAATTTCAAGATAGTTACATGCCTTTTTATTTTTCCTAATAGAGCTGTAGGTCAAATATTATTAGAATCAGATTTCTAA
ATCCCACCCAATGACCTGCTTATTTTAAATCAAATTCAATAATTAATTCTCTTCTTTTTGGAGGATCTGGACATTCTTTGATATTTCTTACAACGAATTTCATGTGTAG
ACCCACTAAACAGAAGCTATAAAAGTTGCATGGTCAAATAAGTCTGAGAAAGTCTGCAGATGATATAATTCACCTGAAGAGTCACAGTATGTAGCCAAATGTTAAAGGT
TTTGAGATGCCATACAGTAAATTTACCAAGCATTTTCTAAATTTATTTGACCACAGAATCCCTATTTTAAGCAACAACTGTTACATCCCATGGATTCCAGGTGACTAAA
GAATACTTATTTCTTAGGATATGTTTTATTGATAATAACAATTAAAATTTCAGATATCTTTCATAAGCAAATCAGTGGTCTTTTTACTTCATGTTTTAATGCTAAAATA
TTTTCTTTTATAGATAGTCAGAACATTATGCCTTTTTCTGACTCCAGCAGAGAGAAAATGCTCCAGGTTATGTGAAGCAGAATCATCATTTAAATATGAGTCAGGGCTC
TTTGTACAAGGCCTGCTAAAGGTATAGTTTCTAGTTATCACAAGTGAAACCACTTTTCTAAAATCATTTTTGAGACTCTTTATAGACAAATCTTAAATATTAGCATTTA
ATGTATCTCATATTGACATGCCCAGAGACTGACTTCCTTTACACAGTTCTGCACATAGACTATATGTCTTATGGATTTATAGTTAGTATCATCAGTGAAACACCATAGA
ATACCCTTTGTGTTCCAGGTGGGTCCCTGTTCCTACATGTCTAGCCTCAGGACTTTTTTTTTTTTTAACACATGCTTAAATCAGGTTGCACATCAAAAATAAGATCATTT
CTTTTTAACTAAATAGATTTGAATTTTATTGAAAAAAAATTTTAAACATCTTTAAGAAGCTTATAGGATTTAAGCAATTCCTATGTATGTGTACTAAAATATATATATT
TCTATATATAATATATATTAGAAAAAAATTGTATTTTTCTTTTATTTGAGTCTACTGTCAAGGAGCAAAACAGAGAAATGTAAATTAGCAATTATTTATAATACTTAAA
GGGAAGAAAGTTGTTCACCTTGTTGAATCTATTATTGTTATTTCAATTATAGTCCCAAGACGTGAAGAAATAGCTTTCCTAATGGTTATGTGATTGTCTCATAGTGACT
ACTTTCTTGAGGATGTAGCCACGGCAAAATGAAATAAAAAAAATTTAAAAATTGTTGCAAATACAAGTTATATTAGGCTTTTGTGCATTTTCAATAATGTGCTGCTATGA
ACTCAGAATGATAGTATTTAAATATAGAAACTAGTTAAAGGAAACGTAGTTTCTATTTGAGTTATACATATCTGTAAATTAGAACTTCTCCTGTTAAAGGCATAATAAA
GTGCTTAATACTTTTGTTTCCTCAGCACCCTCTCATTTAATTATATAATTTTAGTTCTGAAAGGGACCTATACCAGATGCCTAGAGGAAATTTCAAAACTATGATCTAA
TGAAAAAATATTTAATAGTTCTCCATGCAAATACAAATCATATAGTTTTCCAGAAAATACCTTTGACATTATACAAAGATGATTATCACAGCATTATAATAGTAAAAAA

ATGGAAATAGCCTCTTTCTTCTGTTCTGTTCATAGCACAGTGCCTCATACGCAGTAGGTTATTATTACATGGTAACTGGCTACCCCAACTGATTAGGAAAGAAGTAAAT
TTGTTTTATAAAAATACATACTCATTGAGGTGCATAGAATAATTAAGAAATTAAAAGACACTTGTAATTTTGAATCCAGTGAATACCCACTGTTAATATTTGGTATATC
TCTTTCTAGTCTTTTTTTCCCTTTTGCATGTATTTTCTTTAAGACTCCCACCCCCACTGGATCATCTCTGCATGTTCTAATCTGCTTTTTTCACAGCAGATTCTAAGCC
TCTTTGAATATCAACACAAACTTCAACAACTTCATCTATAGATGCCAAATAATAAATTCATTTTTATTTACTTAACCACTTCCTTTGGATGCTTAGGTCATTCTGATGT
TTTGCTATTGAAACCAATGCTATACTGAACACTTCTGTCACTAAAACTTTGCACACACTCATGAATAGCTTCTTAGGATAAATTTTTAGAGATGGATTTGCTAAATCAG
AGACCATTTTTTAAAATTAAAAAACAATTATTCATATCGTTTGGCATGTAAGACAGTAAATTTTCCTTTTATTTTGACAGGATTCAACTGGAAGCTTTGTGCTGCCTTT
CCGGCAAGTCATGTATGCTCCATATCCCACCACACACATAGATGTGGATGTCAATACTGTGAAGCAGATGCCACCCTGTCATGAACATATTTATAATCAGCGTAGATAC
ATGAGATCCGAGCTGACAGCCTTCTGGAGAGCCACTTCAGAAGAAGACATGGCTCAGGATACGATCATCTACACTGACGAAAGCTTTACTCCTGATTTGTACGTAATGC
TCTGCCTGCTGGTACTGTAGTCAAGCAATATGAAATTGTGTCTTTTACGAATAAAAACAAAACAGAAGTTGCATTTAAAAAGAAAGAAATATTACCAGCAGAATTATGC
TTGAAGAAACATTTAATCAAGCATTTTTTTCTTAAATGTTCTTCTTTTTCCATACAATTGTGTTTACCCTAAAATAGGTAAGATTAACCCTTAAAGTAAATATTTAACT
ATTTGTTTAATAAATATATATTGAGCTCCTAGGCACTGTTCTAGGTACCGGGCTTAATAGTGGCCAACCAGACAGCCCCAGCCCCAGCCCCTACATTGTGTATAGTCTA
TTATGTAACAGTTATTGAATGGACTTATTAACAAAACCAAAGAAGTAATTCTAAGTCTTTTTTTTCTTGACATATGAATATAAAATACAGCAAAACTGTTAAAATATAT
TAATGGAACATTTTTTTACTTTGCATTTTATATTGTTATTCACTTCTTATTTTTTTTTAAAAAAAAAAGCCTGAACAGTAAATTCAAAAGGAAAAGTAATGATAATTAA
TTGTTGAGCATGGACCCAACTTGAAAAAAAAAATGATGATGATAAATCTATAATCCTAAAACCCTAAGTAAACACTTAAAAGATGTTCTGAAATCAGGAAAAGAATTAT
AGTATACTTTTGTGTTTCTCTTTTATCAGTTGAAAAAAGGCACAGTAGCTCATGCCTGTAAGAACAGAGCTTTGGGAGTGCAAGGCAGGCGGATCACTTGAGGCCAGGA
GTTCCAGACCAGCCTGGGCAACATAGTGAAACCCCATCTCTACAAAAAATAAAAAAGAATTATTGGAATGTGTTTCTGTGTGCCTGTAATCCTAGCTATTCCGAAAGCT
GAGGCAGGAGGATCTTTTGAGCCCAGGAGTTTGAGGTTACAGGGAGTTATGATGTGCCAGTGTACTCCAGCCTGGGGAACACCGAGACTCTGTCTTATTTAAAAAAAAA
AAAAAAAAAATGCTTGCAATAATGCCTGGCACATAGAAGGTAACAGTAAGTGTTAACTGTAATAACCCAGGTCTAAGTGTGTAAGGCAATAGAAAAATTGGGGCAAATA
AGCCTGACCTATGTATCTACAGAATCAGTTTGAGCTTAGGTAACAGACCTGTGGAGCACCAGTAATTACACAGTAAGTGTTAACCAAAAGCATAGAATAGGAATATCTT
GTTCAAGGGACCCCCAGCCTTATACATCTCAAGGTGCAGAAAGATGACTTAATATAGGACCCATTTTTTCCTAGTTCTCCAGAGTTTTTATTGGTTCTTGAGAAAGTAG
TAGGGGAATGTTTTAGAAAATGAATTGGTCCAACTGAAATTACATGTCAGTAAGTTTTTATATATTGGTAAATTTTAGTAGACATGTAGAAGTTTTCTAATTAATCTGT
GCCTTGAAACATTTTCTTTTTTCCTAAAGTGCTTAGTATTTTTTCCGTTTTTTGATTGGTTACTTGGGAGCTTTTTTGAGGAAATTTAGTGAACTGCAGAATGGGTTTG
CAACCATTTGGTATTTTTGTTTTGTTTTTTAGAGGATGTATGTGTATTTTAACATTTCTTAATCATTTTTAGCCAGCTATGTTTGTTTTGCTGATTTGACAAACTACAG
TTAGACAGCTATTCTCATTTTGCTGATCATGACAAAATAATATCCTGAATTTTTAAATTTTGCATCCAGCTCTAAATTTTCTAAACATAAAATTGTCCAAAAAAATAGTA
TTTTCAGCCACTAGATTGTGTGTTAAGTCTATTGTCACAGAGTCATTTTACTTTTAAGTATATGTTTTTACATGTTAATTATGTTTGTTATTTTTAATTTTAACTTTTT
AAAATAATTCCAGTCACTGCCAATACATGAAAAATTGGTCACTGGAATTTTTTTTTTGACTTTTATTTTAGGTTCATGTGTACATGTGCAGGTGTGTTATACAGGTAAA
TTGCGTGTCATGAGGGTTTGGTGTACAGGTGATTTCATTACCCAGGTAATAAGCATAGTACCCAATAGGTAGTTTTTTGATCCTCACCCTTCTCCCACCCTCAAGTAGG
CCCTGGTGTTGCTGTTTCCTTCTTTGTGTCCATGTATACTCAGTGTTTAGCTCCCACTTAGAAGTGAGAACATGCGGTAGTTGGTTTTCTGTTCCTGGATTAGTTCACT
TAGGATAATGACCTCTAGCTCCATCTGGTTTTTATGGCTGCATAGTATTCCATGGTGTATATGTATCACATTTTCTTTATCCAGTCTACCATTGATAGGCATTTAGGTT
GATTCCCTGTCTTTGTTATCATGAATAGTGCTGTGATGAACATACACATGCATGTGTCTTTATGGTAGAAAAATTTGTATTCCTTTAGGTACATATAGAATAATGGGGT
TGCTAGGGTGAATGGTAGTTCTATTTTCAGTTATTTGAGAAATCTTCAAACTGCTTTTCATAATAGCTAAACTAATTTACAGTCCCGCCAGCAGTGTATAAGTGTTCCC
TTTTCTCCACAACCTTGCCAACATCTGTGATTTTTTGACTTTTTAATAATAGCCATTCCTAGAGAATTGATTTGCAATTCTCTATTAGTGATATTAAGCATTTTTTCAT
ATGCTTTTTAGCTGTCTGTATATATTCTTCTGAAAAATTTTCATGTCCTTTGCCCAGTTTGTAGTGGGGTGGGTTGTTTTTTGCTTGTTAATTAGTTTTAAGTTCCTTC
CAGATTCTGCATATCCCTTTGTTGGATACATGGTTTGCAGATATTTTTCTCCCATTGTGTAGGTTGTCTTTTACTCTGTTGATAGTTTCTTTTGCCATGCAGGAGCTCG
TTAGGTCCCATTTGTGTTTGTTTTTGTTGCAGTTGCTTTTGGCGTCTTCATCATAAAATCTGTGCCAGGGCCTATGTCCAGAATGGTATTTCCTAGGTTGTCTTCCAGG
GTTTTTACAATTTTAGATTTTACGTTTATGTCTTTAATCCATCTTGAGTTGATTTTTGTATATGGCACAAGGAAGGGGTCCAGTTTCACTCCAATTCCTATGGCTAGCA
ATTATCCCAGCACCATTTATTGAATACGGAGTCCTTTCCCCATTGCTTGTTTTTTGTCAACTTTGTTGAAGATCAGATGGTTGTAAGTGTGTGGCTTTATTTCTTGGCT
CTCTATTCTCCATTGGTCTATGTGTCTGTTTTTATAACAGTACCCTGCTGTTCAGGTTCCTATAGCCTTTTAGTATAAAATCGGCTAATGTGATGCCTCCAGCTTTGTT
CTTTTTGCTTAGGATTGCTTTGGCTATTTGGGCTCCTTTTTGGGTCCATATTAATTTTAAAACAGTTTTTTCTGGTTTTGTGAAGGATATCATTGGTAGTTTATAGGAA
TAGCATTGAATCTGTAGATTGCTTTGGGCAGTATGGCCATTTTAACAATATTAATTCTTCCTATCTATGAATATGGAATGTTTTTCCATGTGTTTGTGTCATCTCTTTA
TACCTGATGTATAAAGAAAAGCTGGTATTATTCCTACTCAATCTGTTCCAAAAAATTGAGGAGGAGGAACTCTTCCCTAATGAGGCCAGCATCATTCTGATACCAAAAC
CTGGCAGAGACACAACAGAAAAAAGAAAACTTCAGGCCAATATCCTTGATGAATATAGATGCAAAAATCCTCAACAAAATACTAGCAAACCAAATCCAGCAGCACATCA
AAAAGCTGATCTACTTTGATCAAGTAGGCTTTATCCCTGGGATGCAAGGTTGGTTCAACATACACAAATCAATAAGTGTGATTCATCACATAAACAGAGCTAAAAACAA
AAACCACAAGATTATCTCAATAGGTAGAGAAAAGGTTGTCAATAAAATTTAACATCCTCCATGTTAAAAACCTTCAGTAGGTCAGGTGTAGTGACTCACACCTGTAATC
CCAGCACTTTGGGAGGCCAAGGCGGGCATATCTCTTAAGCCCAGGAGTTCAAGACGAGCCTAGGCAGCATGGTGAAACCCCATCTCTACAAAAAAAAAAAAAAAAAAAA
ATTAGCTTGGTATGGTGACATGCACCTATAGTCCCAGCTATTCAGGAGGTTGAGGTGGGAGGATTGTTTGAGCCCGGGAGGCAGAGGTTGGCAGCGAGCTGAGATCATG
CCACCGCACTCCAGCCTGGGCAACGGAGTGAGACCCTGTCTCAAAAAAGAAAAATCACAAACAATCCTAAACAAACTAGGCATTGAAGGAACATGCCTCAAAAAAATAA
GAACCATCTATGACAGACCCATAGCCAATATCTTACCAAATGGGCAAAAGCTGGAAGTATTCTCCTTGAGAACCGTAACAAGACAAGGATGTCCACTCTCACCACTCCT
TTTCAGCATAGTTCTGGAAGTCCTAGCCAGAGCAATCAGGAAAGAGAAAGAAAGAAAGACATTCAGATAGGAAGAGAAGAAGTCAAACTATTTCTGTTTGCAGGCAGTA
TAATTCTGTACCTAGAAAATCTCATAGTCTCTGCCCAGAAACTCCTAAATCTGTTAAAAATTTCAGCAAAGTTTTGGCATTCTCTATACTCCAACACCTTCCAAAGTGA
GAGCAAAATCAAGAACACAGTCCCATTCACAATAGCCGCAAAACGAATAAAATACCTAGGAATCCAGCTAACCAGGGAGGTGAAAGATCTCTATGAGAATTACAAAACA
CTGCTGAAAGAAATCAGAGATGACACAAACAAATGGAAATGTTCTTTTTTAACACCTTGCTTTATCTAATTCACTTATGATGAAGATACTCATTCAGTGGAACAGGTAT
AATAAGTCCACTCGATTAAATATAAGCCTTATTCTCTTTCCAGAGCCCAAGAAGGGGCACTATCAGTGCCCAGTCAATAATGACGAAATGCTAATATTTTTCCCCTTTA
CGGTTTCTTTCTTCTGTAGTGTGGTACACTCGTTTCTTAAGATAAGGAAACTTGAACTACCTTCCTGTTTGCTTCTACACATACCCATTCTCTTTTTTTGCCACTCTGG
TCAGGTATAGGATGATCCCTACCACTTTCAGTTAAAAACTCCTCCTCTTACTAAATGTTCTCTTACCCTCTGGCCTGAGTAGAACCTAGGGAAAATGGAAGAGAAAAAG
ATGAAAGGGAGGTGGGGCCTGGGAAGGGAATAAGTAGTCCTGTTTGTTTGTGTGTTTGCTTTAGCACCTGCTATATCCTAGGTGCTGTGTTAGGCACACATTATTTTAA
GTGGCCATTATATTACTACTACTCACTCTGGTCGTTGCCAAGGTAGGTAGTACTTTCTTGGATAGTTGGTTCATGTTACTTACAGATGGTGGGCTTGTTGAGGCAAACC
CAGTGGATAATCATCGGAGTGTGTTCTCTAATCTCACTCAAATTTTTCTTCACATTTTTTGGTTTGTTTTGGTTTTTGATGGTAGTGGCTTATTTTTGTTGCTGGTTTG
TTTTTTGTTTTTTTTTGAGATGGCAAGAATTGGTAGTTTTATTTATTAATTGCCTAAGGGTCTCTACTTTTTTTAAAAGATGAGAGTAGTAAAATAGATTGATAGATAC
ATACATACCCTTACTGGGGACTGCTTATATTCTTTAGAGAAAAAATTACATATTAGCCTGACAAACACCAGTAAAATGTAAATATATCCTTGAGTAAATAAATGAATGT
ATATTTTGTGTCTCCAAATATATATATCTATATTCTTACAAATGTGTTTATATGTAATATCAATTTATAAGAACTTAAAATGTTGGCTCAAGTGAGGGATTGTGGAAGG
TAGCATTATATGGCCATTTCAACATTTGAACTTTTTTCTTTTCTTCATTTTCTTCTTTTCTTCAGGAATATTTTTCAAGATGTCTTACACAGAGACACTCTAGTGAAAG
CCTTCCTGGATCAGGTAAATGTTGAACTTGAGATTGTCAGAGTGAATGATATGACATGTTTTCTTTTTTAATATATCCTACAATGCCTGTTCTATATATTTATATTCCC
CTGGATCATGCCCCAGAGTTCTGCTCAGCAATTGCAGTTAAGTTAGTTACACTACAGTTCTCAGAAGAGTCTGTGAGGGCATGTCAAGTGCATCATTACATTGGTTGCC
TCTTGTCCTAGATTTATGCTTCGGGAATTCAGACCTTTGTTTACAATATAATAAATATTATTGCTATCTTTTAAAGATATAATAATAAGATATAAAGTTGACCACAACT
ACTGTTTTTTGAAACATAGAATTCCTGGTTTACATGTATCAAAGTGAAATCTGACTTAGCTTTTACAGATATAATATATACATATATATATCCTGCAATGCTTGTACTA
TATATGTAGTACAAGTATATATATATATGTTTGTGTGTGTATATATATATAGTACGAGCATATATACATATTACCAGCATTGTAGGATATATATATGTTTATATATTAAAA
AAAAGTTATAAACTTAAAACCCTATTATGTTATGTAGAGTATATGTTATATATGATATGTAAAATATATAACATATACTCTATGATAGAGTGTAATATATTTTTTATAT
ATATTTTAACATTTATAAAATGATAGAATTAAGAATTGAGTCCTAATCTGTTTTATTAGGTGCTTTTTGTAGTGTCTGGTCTTTCTAAAGTGTCTAAATGATTTTTCCT
TTTGACTTATTAATGGGGAAGAGCCTGTATATTAACAATTAAGAGTGCAGCATTCCATACGTCAAACAACAAACATTTTAATTCAAGCATTAACCTATAACAAGTAAGT
TTTTTTTTTTTTTTTGAGAAAGGGAGGTTGTTTATTTGCCTGAAATGACTCAAAAATATTTTTGAAACATAGTGTACTTATTTAAATAACATCTTTATTGTTTCATTCT
TTTAAAAAATATCTACTTAATTACACAGTTGAAGGAAATCGTAGATTATATGGAACTTATTTCTTAATATATTACAGTTTGTTATAATAACATTCTGGGGATCAGGCCA
GGAAACTGTGTCATAGATAAAGCTTTGAAATAATGAGATCCTTATGTTTACTAGAAATTTTGGATTGAGATCTATGAGGTCTGTGACATATTGCGAAGTTCAAGGAAAA
TTCGTAGGCCTGGAATTTCATGCTTCTCAAGCTGACATAAAATCCCTCCCACTCTCCACCTCATCATATGCACACATTCTACTCCTACCCACCCACTCCACCCCCTGCA
AAAGTACAGGTATATGAATGTCTCAAAACCATAGGCTCATCTTCTAGGAGCTTCAATGTTATTTGAAGATTTGGGCAGAAAAAATTAAGTAATACGAAATAACTTATGT
ATGAGTTTTAAAAGTGAAGTAAACATGGATGTATTCTGAAGTAGAATGCAAAATTTGAATGCATTTTTAAAGATAAATTAGAAAACTTCTAAAAACTGTCAGATTGTCT
GGGCCTGGTGGCTTATGCCTGTAATCCCAGCACTTTGGGAGTCCGAGGTGGGTGGATCACAAGGTCAGGAGATCGAGACCATCCTGCCAACATGGTGAAACCCCGTCTC
TACTAAGTATACAAAAATTAGCTGGGCGTGGCAGCGTGTGCCTGTAATCCCAGCTACCTGGGAGGCTGAGGCAGGAGAATCGCTTGAACCCAGGAGGTGTAGGTTGCAG

TGAGTCAAGATCGCGCCACTGCACTTTAGCCTGGTGACAGAGCTAGACTCCGTCTCAAAAAAAAAAAAAAATATCAGATTGTTCCTACACCTAGTGCTTCTATACCACA
CTCCTGTTAGGGGGCATCAGTGGAAATGGTTAAGGAGATGTTTAGTGTGTATTGTCTGCCAAGCACTGTCAACACTGTCATAGAAACTTCTGTACGAGTAGAATGTGAG
CAAATTATGTGTTGAAATGGTTCCTCTCCCTGCAGGTCTTTCAGCTGAAACCTGGCTTATCTCTCAGAAGTACTTTCCTTGCACAGTTTCTACTTGTCCTTCACAGAAA
AGCCTTGACACTAATAAAATATATAGAAGACGATACGTGAGTAAAACTCCTACACGGAAGAAAAACCTTTGTACATTGTTTTTTTGTTTTGTTTCCTTTGTACATTTTC
TATATCATAATTTTTGCGCTTCTTTTTTTTTTTTTTTTTTTTTTTTTTTCCATTATTTTAGGCAGAAGGGAAAAAAGCCCTTTAAATCTCTTCGGAACCTGAAGATAGA
CCTTGATTTAACAGCAGAGGGCGATCTTAACATAATAATGGCTCTGGCTGAGAAAATTAAACCAGGCCTACACTCTTTTATCTTTGGAAGACCTTTCTACACTAGTGTG
CAAGAACGAGATGTTCTAATGACTTTTTAACCGTGTGGTTTGCTGTGTCTGTCTCTTCACAGTCACACCTGCTGTTACAGTGTCTCAGCAGTGTGTGGGCACATCCTTC
CTCCCGAGTCCTGCTGCAGGACAGGGTACACTACACTTGTCAGTAGAAGTCTGTACCTGATGTCAGGTGCATCGTTACAGTGAATGACTCTTCCTAGAATAGATGTACT
CTTTTAGGGCCTTATGTTTACAATTATCCTAAGTACTATTGCTGTCTTTTAAAGATATGAATGATGGAATATACACTTGACCATAACTGCTGATTGGTTTTTTGTTTTG
TTTTGTTTGTTTTCTTGGAAACTTATGATTCCTGGTTTACATGTACCACACTGAAACCCTCGTTAGCTTTACAGATAAAGTGTGAGTTGACTTCCTGCCCCTCTGTGTT
CTGTGGTATGTCCGATTACTTCTGCCACAGCTAAACATTAGAGCATTTAAAGTTTGCAGTTCCTCAGAAAGGAACTTAGTCTGACTACAGATTAGTTCTTGAGAGAAGA
CACTGATAGGGCAGAGCTGTAGGTGAAATCAGTTGTTAGCCCTTCCTTTATAGACGTAGTCCTTCAGATTCGGTCTGTACAGAAATGCCGAGGGGTCATGCATGGGCCC
TGAGTATCGTGACCTGTGACAAGTTTTTTGTTGGTTTATTGTAGTTCTGTCAAAGAAAGTGGCATTTGTTTTTATAATTGTTGCCAACTTTTAAGGTTAATTTTCATTA
TTTTTGAGCCGAATTAAAATGCGCACCTCCTGTGCCTTTCCCAATCTTGGAAAATATAATTTCTTGGCAGAGGGTCAGATTTCAGGGCCCAGTCACTTTCATCTGACCA
CCCTTTGCACGGCTGCCGTGTGCCTGGCTTAGATTAGAAGTCCTTGTTAAGTATGTCAGAGTACATTCGCTGATAAGATCTTTGAAGAGCAGGGAAGCGTCTTGCCTCT
TTCCTTTGGTTTCTGCCTGTACTCTGGTGTTTCCCGTGTCACCTGCATCATAGGAACAGCAGAGAAATCTGACCCAGTGCTATTTTTCTAGGTGCTACTATGGCAAACT
CAAGTGGTCTGTTTCTGTTCCTGTAACGTTCGACTATCTCGCTAGCTGTGAAGTACTGATTAGTGGAGTTCTGTGCAACAGCAGTGTAGGAGTATACACAAACACAAAT
ATGTGTTTCTATTTAAAACTGTGGACTTAGCATAAAAAGGGAGAATATATTTATTTTTTACAAAAGGGATAAAAATGGGCCCCGTTCCTCACCCACCAGATTTAGCGAG
AAAAAGCTTTCTATTCTGAAAGGTCACGGTGGCTTTGGCATTACAAATCAGAACAACACACACTGACCATGATGGCTTGTGAACTAACTGCAAGGCACTCCGTCATGGT
AAGCGAGTAGGTCCCACCTCCTAGTGTGCCGCTCATTGCTTTACACAGTAGAATCTTATTTGAGTGCTAATTGTTGTCTTTGCTGCTTTACTGTGTTGTTATAGAAAAT
GTAAGCTGTACAGTGAATAAGTTATTGAAGCATGTGTAAACACTGTTATATATCTTTTCTCCTAGATGGGGAATTTTGAATAAAATACCTTTGAAATTCTGTGTATGTT
TTAGTTCATTATTTAGGGAAAACGCTGCTGTGAAAGGGGGCGTGATCAGCTTCCTATTCTGCGACAGTCGTGTTGAACGGAACCCATTGGTTTTCATCTTCGCTCCCCC
CCCCTTGGTTTTTCGAGACAGGGTTTCTCTGTATAGCCCTGGCTGTCCTGGACCTCACTCTGTAGACCAGGCTGGCCTCGAACTCAGAAATCTACCTGCCTCTGCCTCC
CAAGTGCTGGGAGGCAGTTGCCCCACCAACTAGTCTTCTTTTTTCAAAGAAGATATTTAAAGCTAACGAATAATGCTAGACTCTTACATCTTAAAAAAAAAAGAAGAGA
AAAGAAAAGAAAAGGTAATCACACTGCCCAGTGTGTAGTGCATGCTTCTACTTCCGGTCCTTGGGAGATGGGGGCAGGATGAGACGCTCCAGACCGGCTTCCAATACAG
AGTTCAAGACCCACTGAGCTACGTGAGGCTACACGAGCCTGCCTTTAAAAACATAAAGCTAAAGCTTTCTTCTTAACTTCCAGTATTGCACCTTGATTCCCCCTTCAAA
TTTCACATACAAAATAATTCTTAAATTCTCTTTTGAAAAATGTTCTACTGAGGCCAGAGAGACAGTTCGCTTGGTAAAGGTGCCTGTTGCCAAACGTGATAACCTGAGT
TAAATCATAGCCCCACATGGGGGAGGAAGAAACCCCCGCAGCTTGCCCTCTGATGCCATGTATGCACTAAAACACGCACGTGTGTGCGCACACATTTTTTAAGTTCCTA
TTACATTGATAGTAATATAATTTAAACTGATTTATTCTCCCCAAGTCATTGATACGGGTGTCCAACGTAAAATCCAGCGGCTGAACAAAGCACTTTTAGGCGCTTTAAG
TTGGAAAGCAAGAAACGGAGATTGACACTGTCACTCCAAGAGAAAACTCTTCGTAGTAGCGAGATCGGCTGTGGAGTGAAGATGCTCAGAGGCTGGGAACGCACACAGC
TCAGGAGTGGATAGCATCCCCCAGCCTCAACTCCTAACACTGGGAAAGCGTAGGGCTCTCAGATGAGGAAACAAAACCATACAAAGCTGCTGCAAGCTAAACAGAAAAA
TAGTGGCATTACACTAACTGTTGTGGAATTGTACAGACCGATTCTCCTCCCAATCTGCCGAGTGTGGGCGGCTTGAGAGAATGAAGAGAGCTACTGGCCTCAGGTAACA
GTGCTTCCCACAGGACTGTCTCAGGCTGCCACCACCATAAATAGCATTTAGACGTGACAGAGCTAAGGCTTGACACACAGCCAAAAGCTACTCACATTCCATTTCATC
CCCAGCTGTTCTGTCATCGCTAAGCACAGAGCATTCAGCACAGCTCTTCCCTGTGGTGGGTACTCAGCACTGTTGAGTTGAAAGGATTGAAAAAACTCAAGACTATGTT
CTCAAACATTTTTTTAAGCTCTTTTTAAAACCACCTTAGAATGAAAGCTTTTGACTTCTTATTAACATGCACTAACTTCATATACACATTTAGTGTTATTGTACAGGCA
CGAAGCATACTCTGGTCAGAACCTGTCTCCTTTGGTCCACCCTCCCCACCGTTTTCAGCTTCTATTCCACCTTCCATACGTCTCAAGATCCACATGTGAGAGGGAACAC
TCAGAGCCTTGTCTTTCTGTATCTGGGATATCTCACTTAACATGATATTCTCCAGTTCTGTTCCATCCATTTCATTGCAAAGAGCAAGATTTCACTCTACAGCCAAATA
ACACATTTGTCCATGTATATCCGTATTTTTCCTTATTCATCTGTTGAATGGCACAAGACTGATATCATGGGTAATATCTATGTGGACTTCTAGTCTCTGAGAACAGAGG
CACAGTAAATATGACAGTGATCTTTTTGGTATAATTTCACTTCCTTTGCTTATATGTCCAGAAATGGGATTGAATGATCATCTGGCAGATCCATTTGTGTGTGTGTGTG
TGTGTGTGTGTGTGTGTGTGTGTGTGTGTGTGTGTGTATGTGTGTGTGTGTTCACATGTGTAGGCATAAATGTAGGAGTATGTATATAGACAAATGGCCCATTCCTGTA
AAGGACAAGTTGACACTGGATATCTTCCTCAGTGACTGACTACTTTATTTACTGAGGCTGGGTTGTTTGCTGAACATGGAGCTTGTAGACTCAGCTAGTCTAGGGAGCT
TATCCCAAGGGTCGTCTGTCTTCAGCTTTCAAACACTGGGATTATGAGGTGCCTGGCCGGCTTTTTTAATGTGGGTGCTAAGAGTCTGCACTACAGTCCTTTTGCTTGT
ATAGCAAGCACTTTACCCAGATCCATTTCCCCATCCATCCCCCATGTTTAGCTTTTTCAGGAACCCCCCTATACTATTCTCCATGATAGCCCTACTGATTTACACTTAA
TTACACTGTAGCAAAACACCTGAGAAAAGTCACTTTACAGAAGGACATACACATTTGGGTCTATGAGGTCCGTGGTCACTTGGTCTGTTGTTTCTGAACCTGTAGTAAA
ACAGAGCACCCTGCTTGGGAGGGTACAGAGGAGCAAAGCTGTTGATTAAGGCCGTCCCTACAAGCTCTATGCTAGCAAAGAAATGAACTCCGTGAAACAGGAAGGGGAC
AGAGCTCAGCGTGAGCCTGACACCAGGCCAGTTTAGTATAGCGGTATTTAAGTACAGTACAATTTTGGAAAAAAAGGTTTTCTGGAAAAAACTATCCCAATCCTATGTA
CACAATAAAGTTTAAGGCATGACTGCATCAAAAGTCTTTTGCAAAGTACATGAGACCTCTACCTTGAAGATGGATCTATAGCCCAGGTTCCTGTGGCCTCGAATGGAAG
TCAGCAAGTTATAAAATACATGTGATATCCTCCTCAGGATGGGTTCCACTGGCTGCAAAGCAAAGATAAAGCTAAAGGTCTAGTTGGGGAGAGTGTGGGATAAACATTC
TGGGGAATTTCAGTGAAGCCCAGCCCTACAGGGCGCAAAGGTCACCAGGTTTTTAACAACATCCACTCGCAGCCTTCTGGGTGGGAAACAGGTACACGGCTCCTTCATT
GAGGAGGTGACAATGCCTGATTAGCATTCCTGGTTTCTCCAGTGCTTATCTGTGAGCATTCGCGCCCACTACTAGTATATCATGTAGCCAGGATGCAGAGATAAAGGAG
CCAAGGACAAACTCCACCTTCGACAACACTCTCTCGAGAACTGCTTCCTCCAGCTAGAACCCACTTCCTAAATCTACAAACTTGGCCTGGTTGGATTTTTGTCAGCTTG
ACACAAGCTAGAGTTACCTGGGAAAGGGGACCTCATTTGAGAAAATGCCTCCATCAGATGGTCCATAGGCCAGTCgggccattgggccctagacaagttgtcgcggccg
cactagtgactccagcgtaactggactggccacagttaggccgcaaatgtaatcacactggctcaccttcgggtgggcctttctgcgtctagaggatcccctgttatgc
ccttccttacccatgcgctgagaggtgaaatcactctttaatttgtacgcggtacggacctctatatcgctttgtcgtagaggtggaatctccccagcttgcagccatt
gatagactgcgccgggtgtcacgcccacgccaaccgctgctttttccacatcgccgaaatggcgcacaagttcttcgggtttcatggggattattataatcaataagtg
aaaattaaagctaggtataatttataaaaattatagccagctataaagagatcatttatgattaagggtatgaaaacacgaggcgaacgactgaaagcacgccgtttag
agctgaaactgacgctgaagcaagtggcagaagctgtaggaatctctcttccgggcgtccaaaacttagaacgtggcgacgtgatgccgtcgctggagatcgggctatc
gctggcaaaatgcctgcgtaagcccgtgcaatggatactgtatggcactgaatctgatccagaccgcgttcctgttattggtacgacagagaccggaccggatagagac
tggcagcctggagaacctgccaacacagagcgattcctgccgttcgtgagtcaacggaacaccgtttacgcactgacggtcgggaaccagattcagcgaaactaccagc
cgggagacgttatcctggtcgattcttcacttacgcttgtaccgggtgaggatgtactggtttgtgataataacggcgagatcacaattcaacgattagcccgttatga
cgagcagcattactaccttgatagtgctaactctcaacgggttatccatgataaaagtgatcttcaatttgtgcaccaagtagtcggtacgatcaaatcgttcatggtt
gagggtagatgatacaataacagggtttattgctgactataattctggtttaatccgatctatactttgtcggggttgaaccagaccgtagcagccgaaaaaagacgaa
aaaaaacccgagtcggcaaactcggcctttttcaggaagtagccacgataacgcagatacgtccttcagaaagatagtgcgtttattgtggctgctcctgcggatttt
ttcaacccgaaaaaatgctaattcgcatggaaaggctaaaaaatgaccttacaagaattctacgcggagcgctttggcagcgatccgttttcactgcttgacgcggcac
gtgatgagctgactgagctggcgaagatggccggtatcaactggcctgcgtgtgctgataagattcagctcaatccgcgcggtggtgtagagcgctataccacctacaa
taattcctccccggaagctctgcagaagagccttaaaggccgtgtggaaatctattcacgccttgagcagagcaaggatggcatcagttatcctttcgttaacttcgtt
gaaaaagcgcatgacgctggctcctggagtggcttttcattcctgttttcggagtatcgccgtgaacagcaacggaatggtgcgaccgtggtcgctcagccggaagaag
aacgtgcacgcatggagcgccaggctgaagcacgccgcctccgagctgaacaacaacgcgttaatgatttaaaaaacaatcagatggagcaggaacgcctgcttggatg
gttggctttccatcgcgcctgggaacattcgccagctgaagatggctcctggccttatgcggtaaaaaagggaattcgtgacgtatttggcgcttgtgatatacgtcgc
gtgaccagtcacgacagtgcgaaatggagccgtggaccaacaacttacatggcgatacctctgtcccacctggacggaagaaaagacgggcggattgttggctggcaac
gtatcgatctgaatggtggcaagttccagaccagcgccatcactaacggtgatttcgtcggtgcatgctttgttattggcgacctgcagggggcgcaaaaaattgcgac
tgcagagggtttcgccaccggggcatccatctggctggcgaccagaaacgacccgaaaaaacgctttgacgcggtagtcattgcggtttccgctaacaacatgattcac
gttgtcgagcagctggtgaacatgtacccggctgcgcaaatcacttgcgctctcgataacgaccgcaaatcctcagctgaaggaaaaggcaacacaggcctccgcaccg
gattcgacatcatggagaagttttccggcgtcaaatgtgtttacccaacttttgaggatgaccctgagctggaatgcagcgatttaacgacctgcacagcctgagagg
gctgaaggaagtcgctcgccagttaacgagaaatcatctgagccgtgcaaccgatctgttgtcgatcacgctgaataagctgcgtactctcccgcgtctgaacagacgc
acttttgccaaagaactgcttcgcgccgttgatattggcatgctgacatgcccggtaccaaacagcccgaaagaacttatgcgccttttcagcagcacgctgcgggata
tgggtatcgcagaaatttataacggtaccgttaaagatcacattacgcgccggttgaatcgcaaatgccgtgccgcgcaaacatcacgttcgttcagtgaacgcatcac
caacccgaaccttcgcccgtcacacatcacttacaaacggtttgaaacctccaggatgaccgatgaggtgatgacatacgccgcacagctgcagggcatcgttattgtc

cgcgccgggatggggtcgggtaaatcgacaggcctcctgcgtccactgatgctgcagtccacgcgtggcgtttccgtcgcgcaccgcgtatccctgataggcggcctgc
atgaaatgatgaccgaagggaaaggcgcgaaagccgacattctgcattatcaggatcccggctatcaggaaatggcgccatatgccaacaagctgactatttgcatcaa
ctccatacttaaaggctgctggcaaccgctgatgcgccagcatgacttcttcggcttcgatgaagcaacacagggcctgcgtgccattctggccgggcgtgcgatggaa
aacccggtaggcgtattcaacacgcttatcgacgcgctggcgcgtactgaagagcatgccattatggtggatgccgatgccaacgatctgcttgttgacctggctgaac
tggcgatgaagcgacgcgaggagctgggcctacctgcctggctgcaaattcacgtgattgaactcccggtcgacgttcgcaaccgcgaaaccaacaagcctatccgcgt
attttataccgagaaaaatcggatcatgaccgaggttatcgctgcagtgcaacgcggcgaacgaatcatgctggccaccgacagttcgacgttcgccgaagacgttacc
atgcagctgagactacagttccctgacaaaaagtttctctgcgttaaccagaaaaacaaacaggagaaggaagtcgacgatttcaccaaccagcctaaagtgatggtga
aaaaatatgacggcctcatctacagcccgtcgatatcttcaggtgtatcgattgaggagaaacacttccaccgccatttcggcatgttctgcggcgaagtggtccccag
cgacgccatccagatgctgcgccgcgaccgtacagctcaggaatacattatcggtttcgacaagcttcgcggtaaacgtgaaaccgatccggaaaaaatcaaacgcgcc
tacgcgcaggccttgctcgaaacggctggccactccgggctgctgacagacgttgtctttgacggtgaccgaatttcactcggcgtggctaactcttcattcatgcagc
tcaaaattaaagccgccgcgctcgaggcatcggccagaaatgattatgccagcaacatgatttgcatcatgcatgatgatggctatcaggtcgcgcctatggccaccga
cgcgcttgcaaacagcatcggtaaggatttacgtaaagaagcccgtgaactggtctttgagcagcttatggagcgccacctgagtgtcgatactcctgaccaggccgaa
cacgatgagttgataaaaaaacgcaccctgtccctggatgagcaggcccagctggtccgctgggacatcgaaaaggagctgcagctggatgtcgacgaggtggcgttaa
aattctatttcgacggcggcctgaaaaaagtgcgcctgtttgaaaccatgcagctggatgagataaccgcccgtcgccttgaccgtgaagaagcgttgattcactttac
ctacgcttatcgcgttgccggaagatggcagcagttcgtcactacggccatgacgcgggaacaggccgacgcagagttccaggctaaattcccggccataaccgattac
cgagtgaaatcgaccctgctgttgagataggcatgcgtggtttctacaccctcaaatcggcgacgctgcagcagtacttccgcgactgcggtatcgatccaaaaaccc
tggaaggtgaagcagacatggacgccctcaaacgcgccagagataacctgctcacgccggaacggcgcgatctgctaaacaacgttttacgcatcggcggcttcaacac
agagaaaggcaagaagaaagcgcctgcagacctctgcataggcatcctggagtctatggggctatccagtaagaccagacgcgccagagacggcgatgcacgccccacg
atgcgttccattgacccggactcggtcgagttcctcatgaatatcgtggagaagcgcccgcgaggccggtttatcaattcacgcacgtaaggtggaaaaaaccaccatcg
aagtggatcgcgatttggatctaaatatagatatacatggtaaccctcgatccaaaacagagcacgttccggacgccccacaatcagtaattatccaggcactggaggc
tatcccggtggcggtaccggaggcgtgggcggagaacgcgttgccagcaaccgaaatggaggcggtacgcctgtggccagtggccagcatcgcgagaacgttcgcctcg
ctgtacatgaccgaatttatggacctgctatcagtacgcgaaataaggctgctgaaagcgtttctaagccagcggcaagccgtggcgatataaccggaggggtaatggg
aaaacaacgggaatgccttgtcatcgtggagggctacgacccttatgtggtcaaggtcagccacgacgcgacaagctatagcgattataaaactggccgggaacttcca
ttcacgatcattcggatgaacgggttctcagatagtgagggagtttattgctgcattgccagcgataagagcatggaagaccacacagacgaagccagggcgcttctaa
aagcctataaggtgaagccactatgacgtcgacaccagagttcctgaagcaactggattacgaacagctgaagtactgtcgcgagctgtgcgataaccgcatccggggct
attgaggccgaagagaaaaagatggcctgggcagttactgatggtggcattaactacggctggttccgtacagaagactacatgaaggcggtagaatgcctggttgcta
cagctgcagaacggtgggaagaatcggataaggctgacccatctggacgtggttggcttgacctttcaatccgtggaagccgcctgccggtgtccgagtatgaagcttt
atttgctgatggccagtggggggtgaatagtggcgcatgatctacaaaaaatgatggatggtgagctggtccgcttacaacttgagattaagaaagaacaggagaggcgt
ttagccgaaaagatgattcctgtttttggtgtgtggcaatcacctattaatcaatatgaatttgctgatccagagcaggcaatttcttgtgccgcagggctattggata
aggttcttcgtgaggtcaaggacgatctttccaaagaaggtggattacagggatggaacggcaatttgctacgcatctacgtacagcatgtgaacgagagcgattttgc
agttctacagccttatttaaaggacaaataatcatgatttatcgcagaggatgggtgcctgttttgtggaggaatgagctggaaaagcggctcaaagaagaagggttcg
aaaactggaagcagataagtaactttctttgtggtgaagaggattgctactcggaagcagcaaaaacgccaggacaatacgcttaccaggtcgtcgacaataccgaatg
gatggaacgtcgtgatgcgacgttctggcagcgtctgaaccgtttgtggttcgttccgctgtacctgctgactattccgttccagtggctcattcgtggccgtatgggg
tttgagacaacatctaaggtggctgcggtcatcaataagattaccggactccaataactgatcaggttctgatcagtcagtaaagcaaactataaaaacacctgtgcac
gtgactggtcacggggtaaaatatccacctaaattattgacgtgcgttctgctttggcggtagagttaccccgctgcagcaaaatctgcagccgggcctctcaaccccg
aatgacaagaagcgcacaacacgcgccagcgtgtttttttgtgtgttaaatctgcgcatacctgaattatggtggctcagatggggccaacttcggttgggccggtttc
ttcttgtaccggtgttgagaaccctgtctgggctaccaccccctatagagattctcaactctggtggtagcaccctatacaagataggaatgcataccatgttcaaattc
aaatttgcagctgtcgtccgcacggacaaaaaatcccatatccaccgcctttccaccattgcatcatccgagcgagaagctcgtcgccagttcgccagccgttttgtcc
tcgttctgtcagcccgtatcccggtcagcgaggtgatcgcatgaaccagattcagttaaacgcccagggcctgctggagacgattgaggaacgcctgctgcaggttgaa
gcgcttgtgggctctgctcaccgcactatctccagttatgaggccacgctgtatttgcaggaagctgcggaacttctgcaggtcgcgcgcgaactcacgcaggaagctc
gcggctgctccctgtccctttctgagcagctgaaatcggggaggcgcaatgaaggctctctccgtcttttcatttcaggaaagccaccctatacgggtggttcttgtc
ggtggtgacccgtggtttgtggcgctggatatctgtgctgcattgaatatagccaacccttctgatgcattgcgtaagctggatcatgatgaaaaattgaccctcggtt
taaccgaggcacaaaaacttgatcggatggccagggaagtaaatgtggtatcggaatctggtctctacaccataatcctccgctgccgcgacgcggtgaaacaggggac
gacagcctggcggttccgcaagtgggtgacgaacgaggttctgccagctatccggaaaaacggcgagtatgcttttgtcgagcctgagcctaagaacgccggtgaacca
ctggactggcggcagaaagaagaacttcggggcctgataaacgacatagcacaaagttttcagtaccgtaacgcatgggttagcggcgtctggatggctctccgccgcg
cctgcaggaacccatcaccgaacccaataaccgtggatgatctcccggcaatcatcgccgagctgcgccggatattaacggctgcagaaacggctctgggcaacatgcg
tgtctacgaacgggagctgctgcgcgatgtagttcgtggcggtcgccggagtatgtcttgtggagagctgccgatcaccgatattgatacggaactggagaaggtgctg
ccagcacattttgagctggccattgaaaagctggaaacgctatccacgaaactaacatcgccagcgattgcagaaaaataggttggccagcccgaatttaagccccgc
tttacgcggggaagtccctgcagtcgggacggcgatgtccgtagttgcagctgcagtttccgcctaatccttccgtcccttgcaggggatgggtagggacaggcctcaa
taaaaaagccaccgattaacaccggtggcttttcttttagctgtagtacgtttcccatgcggatcgcattgcggccatagggctatcggcggggccggaccaggcataa
tgctttccatcgtattcaaactctatcttgtacgttccgtccccgttatttttttgcaggcttgaagacgggttttagagcagttggctgatgttcttctggcccttcct
cttcggttggttcatcgccgccaccctcgtcgagctcaatttcatcttcatccagctcgtcgtcttgggactcatcatccggttcgtcgatggtctcaacggattcttc
atcaggcaggacgattgcaggtgtctctatcttcagctgccactgcccgttctcgccaacgaactgccccaatgcatcagcggcaaactccaggtaccggctaatcatc
gtcggggctaaatttaaaggcccggagagtgctgttggttatttttgctgatgggtcctgctccaccagctgcttaacggtttcatggagacggacgccagcgtcacctc
tggcaaagcctggcatttcatcgtccagtttctgcagagccaccagcctggtgttttcatccccaacttcaggtcgccaggttctggagaagttggccagcttgaactg
cttatagtgcagctgggtgttctcatcgtcgtgtccgagaatctccatgaagaacacatcctcgtcgacgtttttccaccgtggatcgacgcggaagaacatctcataa
gcgatgcgagcgtaaatagcgcggctatctttataaacacgacggtcatcgccgaaaaatgatttaacccaagggttaaatgcttttgctaaaatagcatttatcctgc
cgttctcagaccttgtatcatcctttccatatcctttaacaacctcatcgaaatcagatgcagcagagcaagaacgcaattctgttaataattcaacgaataattttgc
ttcgcataaagtataaatcgttctggttacgcttttatcttcagagcgtttttttagcttgccctgagaaattaaccgtatactttcctgaaacggcaaattcaccctga
aacattatctcaatcattcttcgccctgataccgcagccagagcaaaggccaaaggtgccattccagaacgagtgtttaaactaaataaagtcgcaggattattcaaaa
tatcatagatagactgcatgtatgttgggtagtcataaccacaacattacgcttcttctcgcgcagaacatcggcccatcgttgctgtatagatgtacgctccgcagg
gcttagctgcagatggtacagaacctcatggttgaccttgagctggtgtagttcttctaacaacgcagagccttgttggaataacttataaagatagtcacggcgctcc
ttccaatcatcactgtttaaatcactaagagcaaaactccaatctggatatttttttattagtcttgctatttttgcatcactgcctttagagcctattcttacatttg
ataactcttcggcaagcggcattatttctttcaatttagattgtaacgatgacatgtgctggcgaatattagccgtaggcatagaaagccatgaagataattcttcgct
gtaaagaggatacttttccgataatttattaatattttttatcaaagctatgatgtaatttatcatcaaaccgctttcttgccctgctcatataggcgttaaaagtattc
gcggttattcttttctgcaatcctttcccacggaactttcttttatcattaaataacgcgttcttataccgtgcggctgcggctttaattctcttcgttttgtcgcctt
gtgggcggtctgaggcatcaattgcctctacctcattcacaagcgtgttgatcaactcaccgatttttaccttgctcatcccggttcccccgtttctaagtctctggat
atcgtaacacaatcaaagaacaagtgacgattaattatgctgctttcgtcacactataagagaacataatagcagaataacaaaggaacacaacaccggaatagataca
tattgttctgcgcttatgttctccttatcagcacacaatagtccattatacgc

#### Targeting Construct 1

See Figure 2F. pJazz backbone in lowercase. Mouse sequence in **green**. Human sequence in **blue**, selection cassette in **magenta**, pathogenic repeat in **red**, AarI site mutation indicated by gcTggtg.

gcgtataatgggcaattgtgtgctgatatgttccgtaattgagatatgatatagtgcatatgttccgaattattgatacattaatgttgcattcctatgttgtgttctg
ttatggtgttcttagatagtgggggatgtcacatggattgatgttctttgattgtgtttcaatatagagacggaagagatagagatgcctcaaagtgtataaacagcgt
gaaaatcagcatgttcgcttaacttttgatagcaattggtatgatttattatcatttaaaatcattaagttaggttttaaatataccaacgtaagtaataccgttgcgc
ttagatgaagctaaatagcttctgtagcgcatcaaaaaaattgatttgaaccttaactttgcgttttctaagtcactttgcttttttttatgttgctttgctttaaaaacg
taaagtatgatgaaaaacacaaagataagggcatgtttgagaggaaacaggatgtcgttaattaatttgctgaaagactgtatcaatcgcggtcaggaaatgacgcgcg

ctattgccatcgcacagttcggcgatgacagcccggaagcgcgtcgtattactcgccgctggggtataacagaagttgcagacctgatcggcgtaacaccgcaggctat
cagagacgcggaaaaagctggtcgtctaccggctcctgattttgagatgaggggccgagtagaacgtcgcgctggctacacaattgaccaaattagccatatgcggagt
gtgtttggtaacccaaaccaaagacctgacgataaaaacccggttgtactttccgttatgtcacacaagggaggggtttataaaacctcatctgcagtacaccaggcgc
aatggttagccttgcaagggcaccgcgtcctactcgttgaagggaacgacccgcaagggactgcatctatgtaccacggttacgttcctgatttgcacattcacgcaga
cgatacactgcttccatttacctgggtaaacgcgacaacgctgaatatgcgataaaaccaacatgctggccgggtctcgacataatccccagctgcctggcccttcac
cgcattgaaacagatctagcagaaagtcaaaagcctccgaccggaggcttttgacttctgtcacctaggttacgccccgccctctggaactcaggagtcgcgcgctagg
ggccggatcggcaggtgatacttaagtaagccggcttagctagcgggacaggtttcccgactggaaagcgggcagtgagcgcaacgcaattaatgtgagttagctcact
cattaggcaccccaggctttacactttatgcttccggctcgtatgttgtgtggaattgtgagcggacaacaatttcacacaggaaacagctatgaccatgattacgcca
agctatttaggtgagactatagaatactcaagcttgcatgcgatacgtatcgttaacgatggatccgacgcacgtgcgaattcgccctatagtgagtcgtattacaatt
cactggccgtcgttttacaacgtcgtgactgggaaaaccctggcgtcacccaacttaatcgccttgcagcacatcccccttcgccagctggcgtaatagcgaagaggc
ccgcaccgatcgcccttcccaacagttgcgcagctgaatggcgaatcttaagtaggcctccccacctgcatggccggggccggggccggggcgtggtagtgatcttatt
tcattatggtgaaagttggaacctcttacgtgccgatcagattaaaacgaaaggcccagtctttcgactgagcctttcgttttatttgaccatgttggtatgatttaaa
tggtcagttaatcagttctatgtaccagcaaggtccagttgtaagcggccgcttgacttcaagtctaatggcccGCAAGGACCCTAGACTCAAGAAAATGTAAGGTTAC
TAAAGTCTGAAGGACACCAAAGTGAAATGACAATTAGCTTTGGTTTGAGGCATATTTGTTTTGTTCGATTTTTCTGTTTTTTAAAAGCATATAATCAGAAAGAATTATA
GAGCTCGATTGACAATTGCAAGATGAACATGTGAAATAGTATTCCTAGGTGTCAGACAAAATAGAAAATGCCAATCATTCTTAGACAAATGATTTTTATAACACATTAA
AATATAATAGGAAAATCTCACAAAATAAGTCTGGGGCAGGAAGATTGCTCCAAGTTTTAGGCCAACCTGGGCTACAGAGTGATATCCTGTTAGGTGTAGGTGAAGGAAG
AGGAATAGGAAAAAAAAAAGACTCAAGGATAACACTTTTTGGTAACTGGAATGACAGTTCAATGAGACATCCGACATATTCCAATCACAGTCTGCTAAAAAGGAAATGAA
TCCACAGGTAACCATATGGTCTTGGGTTTTAATTAGTATGTGACAACACTGACAGTCAATTATTCCATACTGCTACTAAATACTGAAAACTCTGCTTACTAAATCCCAA
TAAATGCAATAAAATATCAACATAAACTGCAGAATGGCTGATGGTTACAGGTAGATGGTGACACTTGGCAGAGATGCTCGGGAGCACATTTGGAGCTGGCCTGAGAAGT
GTGTGGTAGGAGAGTGTGCATTACTGCTGTGTCTCCTCCCCATGGGACCAGCGCCATCAGCCTGCACTGCTGGCCGCATCTCTGGTCTATCCAAGTATTAGACATGTAC
AACCACAAATTCTCTCTTCTTCTAAAGAAACACAATTTCATATTCTCTTTATGATTGCACATTAACATCTAAGATTTAAGATGATGCTTTTGATTTCCTATGTTTTATG
AGGTGCATTCCTCCCCCTGAGACTCTATTTGGTTTTTTGTTTGTTTGTTTGTTTGTGTTTGTGTTTGCTTTTGGGTTTTTTTGTTTTTTTGTTTTGTGGGTTTTTTGGG
TTTTTTGTTTGTTTGGTTGGTTGATTGGTTGGTTGTTTTGGTTGTGTCCTTTAACAGCTGAGCCAAGTCTCCAGCCCAGGCTTCTGTCCTTAGAAATAGTTACATATAA
AGGTAAGGTGCACAGCAGCAATGCACAAGAACAACTGGTCAAAGATGACATTCCTCTCACTCACTAAGGACATTGTTCATAAAATCACGAGTCCAAAGACAAGCAGAAA
AGAAATGTGTATTGTGCCCTAATGACTTTTTTTTTTTGCAGTACTGAAGCAGTATCATGTGCTGTTAGGATTATAAAGTGTTGGCTCCAGACAGAAACTCAAGGGTTTT
CTATATCATTTCATCCTACTGGCTCCACACACTAAGATTCTATTCGTATTTCAAATGAAGGCAGTATATCATAAAATACATATTTATTAAGTCTTCAGTGAATAAGGAA
GCAACCACATGAAATTACATTTCCTAAGAGAAATGAAATGGTCCGTGGTTTATATAGCAATATTCTTCATCCATGTCTGTCATGAAATTGTTCACATGCCAGTCTCAGT
CACAACTAATTTCCTGTAATCTTGGTAAGTAGGAGCTAATCAACAAGCCCTTGGTTCCAGCCCTGCAGCAATGCACATCACACTTCTGAAACATATTAGCAGAGAAAAT
GGAGAAAAGCCTTCCTGACAACTGGCATTTTTGATGAAGCAAGAAGGTTCTCCGTGTTCTGACATATTTATTGGAGTTATAAATACGTGTAGAGTGTAAGAGGGTTGTA
AGTCATTCATTACTGTATTTATTATATCTGAGTAGTTACCAAAAAAAAAAAATGAAAGTAAAACCATCACTTCTAGAATGTACTGATCTAATTTTCTAGCTCGCCTGATT
GCACATAAACCTTACAGCAACCAGGTGACAGATTACTAGGCGTGCTATTTTCCTGCTGAGCAACAGTGTCCTGCTAAAAGACTGCACTGACCAGCTTCCTGTGGGGTCA
AGCAAGGAAGCCCCAGAGGAAGGGACTATCAGAGGAGTGCAATGTACCTATGGGAGTCACTTCCACTCCTATTAGAAGGGCTGCAGCCATTGCTTTCTGTGGGTGTTCT
GCCATCCACACACAAGGGGAACAAAATGCCCTTGTCATGGATACACAGTCTTACCTAGGGTCACACTGGCCACTCTCTCCATTCCCTCCTTTAAGCCCCCACAGAAAGC
TAGCACTGTGGACACTGAGTCACCAAGGAGTGAAGAACTGTCCAGTTCCGTGGTTAACCTAAAGCAGAGACGATACTTAATATCTGTTGTGGTTACAGAGCCCAACCTC
TTATTTTGCTTTAGAAATCCCCCTTATCGAGGAGTGTCTGCATGACTGAACATTAGTGAAACATGATAGCTGACATTTTTCTATATCGGGTCTTTAAAGGTGTTGTTCT
GGTTGGAAATAGGGGATAGAGTTCTCAGTTCCTGTTTGGAGAGATACTCCCTTTCTTTTTCTTTTCTTTCTTTCTTTCTTTCTTTCTTTCTTTCTTTCTTTCTTTCTTT
CTTTCTTTCTTTCTTTTAATGGGATCCCCCCGCCCCCTTGATAGCCTTCAGCATGTAGTGTAGCCTAGGATAATCTTGAACTTCTAATCCTCCTGTCTCAGATTATAGG
TGTTTGCCATCTCGTCAGTTCGTTTAGTATTGGGGAGTAAACCAGTACGTGGTAGGCAAGCCCTCTGCCAAATGCCCTCCACTTTCAGCCCTGGAGATATAGTTCTATA
GCCAGCATATTTTCACTTAGAGCCTGGAGTTATTTGAGGTAGATTATTTGTAAAGTTTGAGAGGACAGGGTTAAAAATCTATCACACCAATATCATGAATCATATTTCT
ACATACAGTTAGCATCATTTTCTCTTTGGTTCTCTGGAGGAAAGATGTATTCTACCTTACACCTTTCAGGCAATACAACAATTATGAACCAGCATACCTACTTATTATT
TTCTTCCTGATTAATAAAAGTTAGCTACAATCCAATTAAAATATATAGGCTACTTTACCTATGCCTAAGTAATTGTTCCATCTGATGTGAAGAAAAAGGTCTCCACTCA
TTGATCACAAAATATAAGTTGTATCATGGTTTATTTCATTTCCTGTGGTCATACAATATTTTAGATCGTGATGTTTCATTTTTTTCTCCAAAGAGAGAAAAAGAGAGGA
ATATGGTAGGGACCAAACCCAGGGAGTCACACATGCTAGGCAAGCCCTCTACCCACGAAGCTACATCCCTAGTCCTTTCTTGGTTTGATTATTTTTTTTAAATAAAACA
TTTATTCTAGGACTGATATTACACAAAATAAGTTATTTGAGGGACACTGTTTTTGTCTCTAGGTAAAATTTTGAAGGAAAAAAAAAACACTAAGAAGGTATATTCCTTC
AAAGTTCCAGTCTTATTCTGAAGTGTAATGTTATGTTAGTTTGACTCACAGACAGGTTTTAAAGAAGGGCTTACTTCAAGAGGACACCAAACAAATACCTTCTATTCCT
AGTGGGCTCTGGAATCACAGAAAACTGACCCAATCAATTACATTGATAGCTCTGGCTTACTACAGACAAGCAAATTATCTTAAGTGTGCATGCATGCGCGTGTATGTGT
GTTAGTACCTAACACCCACCTGGGAACTTTTCAGCTTTTCAGTGTGGGATATAGTATAAACGTCTATTCCTCGTGTTGTGGATTAGCTGACTGGCCTCACTCAGCTGCC
TTCCTTACCTGCAAACTCACCCACTTTGACTACAGCATCGCACTCTTAACCCTAGCCTTCCAAACATGGTCCTATGCTATTTCTGTGTGTCTGGATGTATTTTTAACTC
TCAGATGTATACTTCATTTATGAGATATACATCTGAAGACCACGGTACAAAACACTGTAAGAACTTGATAGAATGACAACTGCTAGGTAAAAAAAAAAAAAAAAAAAAA
AAAAAAAAAAAAAAAAAGCATACAATACCTGGTGAGAGTTCTATTTTTACCGAAGGTGGTATTGATAGGTATTCTGTTATTAATGCCTTTCTTTTCCCTATAAATGATG
AAAAGTTGCTGGAAAATAATAAACACTACTCATCTGTAGTGAAAAGCCACAATACAGTTACAAACCAATCAATCAATCAATAAATCAGACGTCATGGTGTTCTTTTCCC
AAAGGTTAAAAAACAAAGTGCACTGTGCTATTTGGCAAAAATGACGTTTAGAAGAAAACACGGTGACTACGCACAGAGGGTGGGGGAATCATTGTGCTTGTTGCGGAGT
GAACACGTACAGTGTGCACGCAGACTTACGGCATTTAACCGTGTCATAGGGACCAAAGGAAATCCACTCACTCACTAAATATTTGTTGAGCACCCACTACCTGCCAACT
CCCAAACAAAACAAAGCAAAACTACTTACAACCACAAACTACGCTTCGTAACCTAGATAGATAACGCAGGTGACACTATCTATCTAGGTTGAGCTCAGCTCTGCCCATG
CTTTTCCTGAGCGGCTCTTGGAAGAAAAGCTACAAAGCCCATGACAGCCTCCGCCTGGCCAGCTGCCACTGGCATCTCAAGGCTGGCAAAGCAAAGTGAAAGCGCCAAC
CCGGAACTTACGGAGTCCCACGAGGGAACCGCGGCGCGTCAAGCAGAGACGAGTTCCGCCCACGTGAAAGATGGCGTTTGTAGTGACAGCCATCCCAATTGCCCTTTCC
TTCTAGGTGGAAAGTGGTGTCTAGACAGTCCAGGGAGGGTGTGCGAGGGAGGTGCGTTTTGGTTGCCTCAGCTCGCAACTTAACTCCACAACGGTGACCAAGGACAAAA
GAAGGAAACAAGACTGCAGAGATCCGCACCGGGGAGCCCTGCAGATTCTGGGTGTGTGTTTTTGTTTTTCCCACCCTCTCTCCCCACTACTTGCTCTCACAGTACTCGC
TGAGGGTGAACAAGAAAAGACCTGATAAAGATTAACCCTAGAAAGATAGTCTGCGTAAAATTGACGCATGCATTCTTGAAATATTGCTCTCTCTTTCTAAATAGCGCGA
ATCCGTCGCTGTGCATTTAGGACATCTCAGTCGCCGCTTGGAGCTCCCGTGAGGCGTGCTTGTCAATGCGGTAAGTGTCACTGATTTTGAACTATAACGACCGCGTGAG
TCAAAATGACGCATGATTATCTTTTACGTGACTTTTAAGATTTAACTCATACGATAATTATATTGTTATTTCATGTTCTACTTACGTGATAACTTATTATATATATATT
TTCTTGTTATAGATATCGCGGCCGCTCTAGCCTCGAGGCTAGAACTAGTGGATCTCGAGCCCCAGCTGGTTCTTTCCGCCTCAGAAGCCATAGAGCCCACCGCATCCCC
AGCATGCCTGCTATTGTCTTCCCAATCCTCCCCCTTGCTGTCCTGCCCCACCCCACCCCCCAGAATAGAATGACACCTACTCAGACAATGCGATGCAATTTCCTCATTT
TATTAGGAAAGGACAGTGGGAGTGGCACCTTCCAGGGTCAAGGAAGGCACGGGGGAGGGGCAAACAACAGATGGCTGGCAACTAGAAGGCACAGTCGAGGCTGATCAGC
GAGCTCTAGAGCTCAGAAGAACTCGTCAAGAAGGCGATAGAAGGCGATGCGCTGCGAATCGGGAGCGGCGATACCGTAAAGCACGAGGAAGCGGTCAGCCCATTCGCCG
CCAAGCTCTTCAGCAATATCACGGGTAGCCAACGCTATGTCCTGATAGCGGTCCGCCACACCCAGCCGGCCACAGTCGATGAATCCAGAAAAGCGGCCATTTTCCACCA
TGATATTCGGCAAGCAGGCATCGCCATGGGTCACGACGAGATCCTCGCCGTCGGGCATGCGCGCCTTGAGCCTGGCGAACAGTTCGGCTGGCGCGAGCCCCTGATGCTC
TTCGTCCAGATCATCCTGATCGACAAGACCGGCTTCCATCCGAGTACGTGCTCGCTCGATGCGATGTTTCGCTTGGTGGTCGAATGGGCAGGTAGCCGGATCAAGCGTA
TGCAGCCGCCGCATTGCATCAGCCATGATGGATACTTTCTCGGCAGGAGCAAGGTGAGATGACAGGAGATCCTGCCCCGGCACTTCGCCCAATAGCAGCCAGTCCCTTC
CCGCTTCAGTGACAACGTCGAGCACAGCTGCGCAAGGAACGCCCGTCGTGGCCAGCCACGATAGCCGCGCTGCCTCGTCCTGCAGTTCATTCAGGGCACCGGACAGGTC
GGTCTTGACAAAAAGAACCGGGCGCCCCTGCGCTGACAGCCGGAACACGGCGGCATCAGAGCAGCCGATCGTCTGTTGTGCCCAGTCATAGCCGAATAGCCTCTCCACC
CAAGCGGCCGGAGAACCTGCGTGCAATCCATCTTGTTCAATGGCCGATCCCATGGTTTAGTTCCTCACCTTGTCGTATTATACTATGCCGATATACTATGCCGATGATT
AATTGTCAACAGGCTGCAGGTCGAAAGGCCCGGAGATGAGGAAGAGGAGAACAGCGCGGCAGACGTGCGCTTTTGAAGCGTGCAGAATGCCGGGCCTCCGGAGGACCTT
CGGGCGCCCGCCCCGCCCCTGAGCCCGCCCCTGAGCCCGCCCCCGGACCCACCCCTTCCCAGCCTCTGAGCCCAGAAAGCGAAGGAGCAAAGCTGCTATTGGCCGCTGC
CCCAAAGGCCTACCCGCTTCCATTGCTCAGCGGTGCTGTCCATCTGCACGAGACTAGTGAGACGTGCTACTTCCATTTGTCACGTCCTGCACGACGCGAGCTGCGGGGC
GGGGGGGAACTTCCTGACTAGGGGAGGAGTAGAAGGTGGCGCGAAGGGGCCACCAAAGAACGGAGCCGGTTGGCGCCTACCGGTGGATGTGGAATGTGTGCGAGGCCAG
AGGCCACTTGTGTAGCGCCAAGTGCCCAGCGGGGCTGCTAAAGCGCATGCTCCAGACTGCCTTGGGAAAAGCGCCTCCCCTACCCGGTAGAATTAATTCGATATCAAGC
TTGCTCGGATGATAGGGCCCTAACGTGTTTTTGCTTTGTTACTTTATAGAAGAAATTTTGAGTTTTTGTTTTTTTTTAATAAATAAATAAACATAAATAAATTGTTTGT

TGAATTTATTATTAGTATGTAAGTGTAAATATAATAAAACTTAATATCTATTCAAATTAATAAATAAACCTCGATATACAGACCGATAAAACACATGCGTCAATTTTAC
GCATGATTATCTTTAACGTACGTCACAATATGATTATCTTTCTAGGGTTAACCAGAAGAAAACAAGGAGGGAAACAACCGCAGCCTGTAGCAAGCTCTGGAACTCAGGA
GTCGCGCGCTAGGGGCCGGGGCCGGGGCCGGGGCCGGGGCCGGGGCCGGGGCCGGGGCCGGGGCCGGGGCCGGGGCCGGGGCCGGGGCCGGGGCCGGGGCCGGGGCCGG
GGCCGGGGCCGGGGCCGGGGCCGGGGCCGGGGCCGGGGCCGGGGCCGGGGCCGGGGCCGGGGCCGGGGCCGGGGCCGGGGCCGGGGCCGGGGCCGGGGCCGGGGCCGGG
GCCGGGGCCGGGGCCGGGGCCGGGGCCGGGGCCGGGGCCGGGGCCGGGGCCGGGGCCGGGGCCGGGGCCGGGGCCGGGGCCGGGGCCGGGGCCGGGGCCGGGGCCGGGG
CCGGGGCCGGGGCCGGGGCCGGGGCCGGGGCCGGGGCCGGGGCCGGGGCCGGGGCCGGGGCCGGGGCCGGGGCCGGGGCCGGGGCCGGGGCCGGGGCCGGGGCCGGGGC
CGGGGCCGGGGCCGGGGCCGGGGCCGGGGCCGGGGCCGGGGCCGGGGCCGGGGCCGGGGCCGGGGCCGGGGCCGGGGCCGGGGCCGGGGCCGGGGCCGGGGCCGGGGCC
GGGGCCGGGGCCGGGGCCGGGGCCGGGGCCGGGGCCGGGGCCGGGGCCGGGGCCGGGGCCGGGGCCGGGGCCGGGGCCGGGGCCGGGGCCGGGGCCGGGGCCGGGGCCG
GGGCCGGGGCCGGGGCCGGGGCCGGGGCCGGGGCCGGGGCCGGGGCCGGGGCCGGGGCCGGGGCCGGGGCCGGGGCCGGGGCCGGGGCCGGGGCCGGGGCCGGGGCCGG
GGCCGGGGCCGGGGCCGGGGCCGGGGCCGGGGCCGGGGCCGGGGCCGGGGCCGGGGCCGGGGCCGGGGCCGGGGCCGGGGCCGGGGCCGGGGCCGGGGCCGGGGCCGGG
GCCGGGGCCGGGGCCGGGGCCGGGGCCGGGGCCGGGGCCGGGGCCGGGGCCGGGGCCGGGGCCGGGGCCGGGGCCGGGGCCGGGGCCGGGGCCGGGGCCGGGGCCGGGG
CCGGGGCCGGGGCCGGGGCCGGGGCCGGGGCCGGGGCCGGGGCCGGGGCCGGGGCCGGGGCCGGGGCCGGGGCCGGGGCCGGGGCCGGGGCCGGGGCCGGGGCCGGGGC
CGGGGCCGGGGCCGGGGCCGGGGCCGGGGCCGGGGCCGGGGCCGGGGCCGGGGCCGGGGCCGGGGCCGGGGCCGGGGCCGGGGCCGGGGCCGGGGCCGGGGCCGGGGCC
GGGGCCGGGGCCGGGGCCGGGGCCGGGGCCGGGGCCGGGGCCGGGGCCGGGGCCGGGGCCGGGGCCGGGGCCGGGGCCGGGGCCGGGGCCGGGGCCGGGGCCGGGGCCG
GGGCCGGGGCCGGGGCCGGGGCCGGGGCCGGGGCCGGGGCCGGGGCCGGGGCCGGGGCCGGGGCCGGGGCCGGGGCCGGGGCCGGGGCCGGGGCCGGGGCCGGGGCCGG
GGCCGGGGCCGGGGCCGGGGCCGGGGCCGGGGCCGGGGCCGGGGCCGGGGCCGGGGCCGGGGCCGGGGCCGGGGCCGGGGCCGGGGCCGGGGCCGGGGCCGGGGCCGGG
GCCGGGGCCGGGGCCGGGGCCGGGGCCGGGGCCGGGGCCGGGGCCGGGGCCGGGGCCGGGGCCGGGGCCGGGGCCGGGGCCGGGGCCGGGGCCGGGGCCGGGGCCGGGG
CCGGGGCCGGGGCCGGGGCCGGGGCCGGGGCCGGGGCCGGGGCCGGGGCCGGGGCCGGGGCCGGGGCCGGGGCCGGGGCCGGGGCCGGGGCCGGGGCCGGGGCCGGGGC
CGGGGCCGGGGCCGGGGCCGGGGCCGGGGCCGGGGCCGGGGCCGGGGCCGGGGCCGGGGCCGGGGCCGGGGCCGGGGCCGGGGCCGGGGCCGGGGCCGGGGCCGGGGCC
GGGGCCGGGGCCGGGGCCGGGGCCGGGGCCGGGGCCGGGGCCGGGGCCGGGGCCGGGGCCGGGGCCGGGGCCGGGGCCGGGGCCGGGGCCGGGGCCGGGGCCGGGGCCG
GGGCCGGGGCCGGGGCCGGGGCCGGGGCCGGGGCCGGGGCCGGGGCCGGGGCCGGGGCCGGGGCCGGGGCCGGGGCCGGGGCCGGGGCCGGGGCCGGGGCCGGGGCCGG
GGCCGGGGCCGGGGCCGGGGCCGGGGCCGGGGCCGGGGCCGGGGCCGGGGCCGGGGCCGGGGCCGGGGCCGGGGCCGGGGCCGGGGCCGGGGCCGGGGCCGGGGCCGGG
GCCGGGGCCGGGGCCGGGGCCGGGGCCGGGGCCGGGGCCGGGGCCGGGGCCGGGGCCGGGGCCGGGGCCGGGGCCGGGGCCGGGGCCGGGGCCGGGGCCGGGGCCGGGG
CCGGGGCCGGGGCCGGGGCCGGGGCCGGGGCCGGGGCCGGGGCCGGGGCCGGGGCCGGGGCCGGGGCCGGGGCCGGGGCCGGGGCCGGGGCCGGGGCCGGGGCCGGGGC
CGGGGCCGGGGCCGGGGCCGGGGCCGGGGCCGGGGCCGGGGCCGGGGCCGGGGCCGGGGCCGGGGCCGGGGCCGGGGCCGGGGCCGGGGCCGGGGCCGGGGCCGGGGCC
GGGGCCGGGGCCGGGGCCGGGGCCGGGGCCGGGGCCGGGGCCGGGGCCGGGGCCGGGGCCGGGGCCGGGGCCGGGGCCGGGGCCGGGGCCGGGGCCGGGGCCGGGGCCG
GGGCCGGGGCCGGGGCCGGGGCCGGGGCCGGGGCCGGGGCCGGGGCCGGGGCCGGGGCCGGGGCCGGGGCCGGGGCCGGGGCCGGGGCCGGGGCCGGGGCCGGGGCCGG
GGCCGGGGCCGGGGCCGGGGCCGGGGCCGGGGCCGGGGCCGGGGCCGGGGCCGGGGCCGGGGCCGGGGCCGGGGCCGGGGCCGGGGCCGGGGCCGGGGCCGGGGCCGGG
GCCGGGGCCGGGGCCGGGGCCGGGGCCGGGGCCGGGGCCGGGGCCGGGGCCGGGGCCGGGGCCGGGGCCGGGGCCGGGGCCGGGGCCGGGGCCGGGGCCGGGGCCGGGG
CCGGGGCCGGGGCCGGGGCCGGGGCCGGGGCCGGGGCCGGGGCCGGGGCCGGGGCCGGGGCCGGGGCCGGGGCCGGGGCCGGGGCCGGGGCCGGGGCCGGGGCCGGGGC
CGGGGCCGGGGCCGGGGCCGGGGCCGGGGCCGGGGCCGGGGCCGGGGCCGGGGCCGGGGCCGGGGCCGGGGCCGGGGCCGGGGCCGGGGCCGGGGCCGGGGCCGGGGCC
GGGGCCGGGGCCGGGGCCGGGGCCGGGGCCGGGGCCGGGGCCGGGGCCGGGGCCGGGGCCGGGGCCGGGGCCGGGGCCGGGGCCGGGGCCGGGGCCGGGGCCGGGGCCG
GGGCCGGGGCCGGGGCCGGGGCCGGGGCCGGGGCCGGGGCCGGGGCCGGGGCCGGGGCCGGGGCCGGGGCCGGGGCCGGGGCCGGGGCCGGGGCCGGGGCCGGGGCCGG
GGCCGGGGCCGGGGCCGGGGCCGGGGCCGGGGCCGGGGCCGGGGCCGGGGCCGGGGCCGGGGCCGGGGCCGGGGCCGGGGCCGGGGCCGGGGCCGGGGCCGGGGCCGGG
GCCGGGGCCGGGGCCGGGGCCGGGGCCGGGGCCGGGGCCGGGGCCGGGGCCGGGGCCGGGGCCGGGGCCGGGGCCGGGGCCGGGGCCGGGGCCGGGGCCGGGGCCGGGG
CCGGGGCCGGGGCCGGGGCCGGGGCCGGGGCCGGGGCCGGGGCCGGGGCCGGGGCCGGGGCCGGGGCCGGGGCCGGGGCCGGGGCCGGGGCCGGGGCCGGGGCCGGGGC
CGGGGCCGGGGCCGGGGCCGGGGCCGGGGCCGGGGCCGGGGCCGGGGCCGGGGCCGGGGCCGGGGCCGGGGCCGGGGCCGGGGCCGGGGCCGGGGCCGGGGCCGGGGCC
GGGGCCGGGGCCGGGGCCGGGGCCGGGGCCGGGGCCGGGGCCGGGGCCGGGGCCGGGGCCGGGGCCGGGGCCGGGGCCGGGGCCGGGGCCGGGGCCGGGGCCGGGGCCG
GGGCCGGGGCCGGGGCCGGGGCCGGGGCCGGGGCCGGGGCCGGGGCCGGGGCCGGGGCCGGGGCCGGGGCCGGGGCCGGGGCCGGGGCCGGGGCCGGGGCCGGGGCCGG
GGCCGGGGCCGGGGCCGGGGCCGGGGCCGGGGCCGGGGCCGGGGCCGGGGCCGGGGCCGGGGCCGGGGCCGGGGCCGGGGCCGGGGCCGGGGCCGGGGCCGGGGCCGGG
GCCGGGGCCGGGGCCGGGGCCGGGGCCGGGGCCGGGGCCGGGGCCGGGGCCGGGGCCGGGGCCGGGGCCGGGGCCGGGGCCGGGGCCGGGGCCGGGGCCGGGGCCGGGG
CCGGGGCCGGGGCCGGGGCCGGGGCCGGGGCCGGGGCCGGGGCCGGGGCCGGGGCCGGGGCCGGGGCCGGGGCCGGGGCCGGGGCCGGGGCCGGGGCCGGGGCCGGGGC
CGGGGCCGGGGCCGGGGCCGGGGCCGGGGCCGGGGCCGGGGCCGGGGCCGGGGCCGGGGCCGGGGCCGGGGCCGGGGCCGGGGCCGGGGCCGGGGCCGGGGCCGGGGCC
GGGGCCGGGGCCGGGGCCGGGGCCGGGGCCGGGGCCGGGGCCGGGGCCGGGGCCGGGGCCGGGGCCGGGGCCGGGGCCGGGGCCGGGGCCGGGGCCGGGGCCGGGGCCG
GGGCCGGGGCCGGGGCCGGGGCCGGGGCCGGGGCCGGGGCCGGGGCCGGGGCCGGGGCCGGGGCCGGGGCCGGGGCCGGGGCCGGGGCCGGGGCCGGGGCCGGGGCCGG
GGCCGGGGCCGGGGCCGGGGCCGGGGCCGGGGCCGGGGCCGGGGCCGGGGCCGGGGCCGGGGCCGGGGCCGGGGCCGGGGCCGGGGCCGGGGCCGGGGCCGGGGCCGGG
GCCGGGGCCGGGGCCGGGGCCGGGGCCGGGGCCGGGGCCGGGGCCGGGGCCGGGGCCGGGGCCGGGGCCGGGGCCGGGGCCGGGGCCGGGGCCGGGGCCGGGGCCGGGG
CCGGGGCCGGGGCCGGGGCCGGGGCCGGGGCCGGGGCCGGGGCCGGGGCCGGGGCCGGGGCCGGGGCCGGGGCCGGGGCCGGGGCCGGGGCCGGGGCCGGGGCCGGGGC
CGGGGCCGGGGCCGGGGCCGGGGCCGGGGCCGGGGCCGGGGCCGGGGCCGGGGCCGGGGCCGGGGCCGGGGCCGGGGCCGGGGCCGGGGCCGGGGCCGGGGCCGGGGCC
GGGGCCGGGGCCGGGGCCGGGGCCGGGGCCGGGGCCGGGGCCGGGGCCGGGGCCGGGGCCGGGGCCGGGGCCGGGGCCGGGGCCGGGGCCGGGGCCGGGGCCGGGGCCG
GGGCCGGGGCCGGGGCCGGGGCCGGGGCCGGGGCCGGGGCCGGGGCCGGGGCCGGGGCCGGGGCCGGGGCCGGGGCCGGGGCCGGGGCCGGGGCCGGGGCCGGGGCCGG
GGCCGGGGCCGGGGCCGGGGCCGGGGCCGGGGCCGGGGCCGGGGCCGGGGCCGGGGCCGGGGCCGGGGCCGGGGCCGGGGCCGGGGCCGGGGCCGGGGCCGGGGCCGGG
GCCGGGGCCGGGGCCGGGGCCGGGGCCGGGGCCGGGGCCGGGGCCGGGGCCGGGGCCGGGGCCGGGGCCGGGGCCGGGGCCGGGGCCGGGGCCGGGGCCGGGGCCGGGG
CCGGGGCCGGGGCCGGGGCCGGGGCCGGGGCCGGGGCCGGGGCCGGGGCCGGGGCCGGGGCCGGGGCCGGGGCCGGGGCCGGGGCCGGGGCCGGGGCCGGGGCCGGGGC
CGGGGCCGGGGCCGGGGCCGGGGCCGGGGCCGGGGCCGGGGCCGGGGCCGGGGCCGGGGCCGGGGCCGGGGCCGGGGCCGGGGCCGGGGCCGGGGCCGGGGCCGGGGCC
GGGGCCGGGGCCGGGGCCGGGGCCGGGGCCGGGGCCGGGGCCGGGGCCGGGGCCGGGGCCGGGGCCGGGGCCGGGGCCGGGGCCGGGGCCGGGGCCGGGGCCGGGGCCG
GGGCCGGGGCCGGGGCCGGGGCCGGGGCCGGGGCCGGGGCCGGGGCCGGGGCCGGGGCCGGGGCCGGGGCCGGGGCCGGGGCCGGGGCCGGGGCCGGGGCCGGGGCCGG
GGCCGGGGCCGGGGCCGGGGCCGGGGCCGGGGCCGGGGCCGGGGCCGGGGCCGGGGCCGGGGCCGGGGCCGGGGCCGGGGCCGGGGCCGGGGCCGGGGCCGGGGCCGGG
GCCGGGGCCGGGGCCGGGGCCGGGGCCGGGGCCGGGGCCGGGGCCGGGGCCGGGGCCGGGGCCGGGGCCGGGGCGGGGCTGCGGTTGCGGTGCCTGCGCCCGCGGCGGC
GGAGGCGCAGGCGGTGGCGAGTGGGTGAGTGAGGAGGCGGCATCCTGGCGGGTGGCTGTTTGGGGTTCGGCTGCCGGGAAGAGGCGCGGGTAGAAGCGGGGGCTCTCCT
CAGAGCTCGACGCATTTTTACTTTCCCTCTCATTTCTCTGACCGAAGCTGGGTGTCGGGCTTTCGCCTCTAGCGACTGGTGGAATTGCCTGCATCCGGGCCCCGGGCTT
CCCGGCGGCGGCGGCGGCGGCGGCGCAGGGACAAGGGATGGGGATCTGGCCTCTTCCTTGCTTTCCCGCCCTCAGTACCCGAGCTGTCTCCTTCCCGGGGACCCGC
TGGGAGCGCTGCCGCTGCGGGCTCGAGAAAAGGGAGCCTCGGGTACTGAGAGGCCTCGCCTGGGGGAAGGCCGGAGGGTGGGCGGCGCGCGGCTTCTGCGGACCAAGTC
GGGGTTCGCTAGGAACCCGAGACGGTCCCTGCCGGCGAGGAGATCATGCGGGATGAGATGGGGGTGTGGAGACGCCTGCACAATTTCAGCCCAAGCTTCTAGAGAGTGG
TGATGACTTGCATATGAGGGCAGCAATGCAAGTCGGTGTGCTCCCCATTCTGTGGGACATGACCTGGTTGCTTCACAGCTCCGAGATGACACAGACTTGCTTAAAGGAA
GTGACTATTGTGACTTGGGCATCACTTGACTGATGGTAATCAGTTGTCTAAAGAAGTGCACAGATTACATGTCCGTGTGCTCATTGGGTCTATCTGGCCGCGTTGAACA
CCACCAGGCTTTGTATTCAGAAACAGGAGGGAGGTCCTGCACTTTCCCAGGAGGGGTGGCCCTTTCAGATGCAATCGAGATTGTTAGGCTCTGGGAGAGTAGTTGCCTG
GTTGTGGCAGTTGGTAAATTTCTATTCAAACAGTTGCCATGCACCAGTTGTTCACAACAAGGGTACGTAATCTGTCTGGCATTACTTCTACTTTTGTACAAAGGATCAA
AAAAAAAAAAGATACTGTTAAGATATGATTTTTCTCAGACTTTGGGAAACTTTTAACATAATCTGTGAATATCACAGAAACAAGACTATCATATAGGGGATATTAATAA
CCTGGAGTCAGAATACTTGAAATACGGTGTCATTTGACACGGGCATTGTTGTCACCACCTCTGCCAAGGCCTGCCACTTTAGGAAAACCCTGAATCAGTTGGAAACTGC
TACATGCTGATAGTACATCTGAAACAAGAACGAGAGTAATTACCACATTCCAGATTGTTCACTAAGCCAGCATTTACCTGCTCCAGGAAAAAATTACAAGCACCTTATG
AAGTTGATAAAATATTTTGTTTGGCTATGTTGGCACTCCACAATTTGCTTTCAGAGAAACAAAGTAAACCAAGGAGGACTTCTGTTTTTCAAGTCTGCCCTCGGGTTCT
ATTCTACGTTAATTAGATAGTTCCCAGGAGGACTAGGTTAGCCTACCTATTGTCTGAGAAACTTGGAACTGTGAGAAATGGCCAGATAGTGATATGAACTTCACCTTCC
AGTCTTCCCTGATGTTGAAGATTGAGAAAGTGTTGTGAACTTTCTGGTACTGTAAACAGTTCACTGTCCTTGAAGTGGTCCTGGGCAGCTCCTGTTGTGGAAAGTGGAC
GGTTTAGGATCCTGCTTCTCTTTGGGCTGGGAGAAAATAAACAGCATGGTTACAAGTATTGAGAGCCAGGTTGGAGAAGGTGGCTTACACCTGTAATGCCAGAGCTTTG
GGAGGCGGAGGCAAGAGGATCACTTGAAGCCAGGAGTTCAAGCTCAACCTGGGCAACGTAGACCCTGTCTCTACAAAAAATTAAAAACTTAGCCGGGCGTGGTGATGTG
CACCTGTAGTCCTAGCTACTTGGGAGGCTGAGGCAGGAGGGTCATTTGAGCCCAAGAGTTTGAAGTTACCGAGAGCTATGATCCTGCCAGTGCATTCCAGCCTGGATGA
CAAAACGAGACCCTGTCTCTAAAAAACAAGAAGTGAGGGCTTTATGATTGTAGAATTTTCACTACAATAGCAGTGGACCAACCACCTTTCTAAATACCAATCAGGGAAG

AGATGGTTGATTTTTTAACAGACGTTTAAAGAAAAAGCAAAACCTCAAACTTAGCACTCTACTAACAGTTTTAGCAGATGTTAATTAATGTAATCATGTCTGCATGTAT
GGGATTATTTCCAGAAAGTGTATTGGGAAACCTCTCATGAACCCTGTGAGCAAGCCACCGTCTCACTCAATTTGAATCTTGGCTTCCCTCAAAAGACTGGCTAATGTTT
GGTAACTCTCTGGAGTAGACAGCACTACATGTACGTAAGATAGGTACATAAACAACTATTGGTTTTGAGCTGATTTTTTTCAGCTGCATTTGCATGTATGGATTTTTCT
CACCAAAGACGATGACTTCAAGTATTAGTAAAATAATTGTACAGCTCTCCTGATTATACTTCTCTGTGACATTTCATTTCCCAGGCTATTTCTTTTGGTAGGATTTAAA
ACTAAGCAATTCAGTATGATCTTTGTCCTTCATTTTCTTTCTTATTCTTTTTGTTTGTTTGTTTGTTTGTTTTTTTCTTGAGGCAGAGTCTCTCTCTGTCGCCCAGGCT
GGAGTGCAGTGGCGCCATCTCAGCTCATTGCAACCTCTGCCACCTCCGGGTTCAAGAGATTCTCCTGCCTCAGCCTCCCGAGTAGCTGGGATTACAGGTGTCCACCACC
ACACCCGGCTAATTTTTTGTATTTTTAGTAGAGGTGGGGTTTCACCATGTTGGCCAGGCTGGTCTTGAGCTCCTGACCTCAGGTGATCCACCTGCCTCGGCCTACCAAA
GAGCTGGGATAACAGGTGTGACCCACCATGCCCGGCCCATTTTTTTTTTCTTATTCTGTTAGGAGTGAGAGTGTAACTAGCAGTATAATAGTTCAATTTTCACAACGTG
GTAAAAGTTTCCCTATAATTCAATCAGATTTTGCTCCAGGGTTCAGTTCTGTTTTAGGAAATACTTTTATTTTCAGTTTAATGATGAAATATTAGAGTTGTAATATTGC
CTTTATGATTATCCACCTTTTTAACCTAAAAGAATGAAAGAAAAATATGTTTGCAATATAATTTTATGGTTGTATGTTAACTTAATTCATTATGTTGGCCTCCAGTTTG
CTGTTGTTAGTTATGACAGCAGTAGTGTCATTACCATTTCAATTCAGATTACATTCCTATATTTGATCATTGTAAACTGACTGCTTACATTGTATTAAAAACAGTGGAT
ATTTTAAAGAAGCTGTACGGCTTATATCTAGTGCTGTCTCTTAAGACTATTAAATTGATACAACATATTTAAAAGTAAATATTACCTAAATGAATTTTTGAAATTACAA
ATACACGTGTTAAAACTGTCGTTGTGTTCAACCATTTCTGTACATACTTAGAGTTAACTGTTTTGCCAGGCTCTGTATGCCTACTCATAATATGATAAAAGCACTCATC
TAATGCTCTGTAAATAGAAGTCAGTGCTTTCCATCAGACTGAACTCTCTTGACAAGATGTGGATGAAATTCTTTAAGTAAAATTGTTTACTTTGTCATACATTTACAGA
TCAAATGTTAGCTCCCAAAGCAATCATATGGCAAAGATAGGTATATCATAGTTTGCCTATTAGCTGCTTTGTATTGCTATTATTATAAATAGACTTCACAGTTTTAGAC
TTGCTTAGGTGAAATTGCAATTCTTTTTACTTTCAGTCTTAGATAACAAGTCTTCAATTATAGTACAATCACACATTGCTTAGGAATGCATCATTAGGCGATTTTGTCA
TTATGCAAACATCATAGAGTGTACTTACACAAACCTAGATAGTATAGCCTTTATGTACCTAGGCCGTATGGTATAGTCTGTTGCTCCTAGGCCACAAACCTGTACAACT
GTTACTGTACTGAATACTATAGACAGTTGTAACACAGTGGTAAATATTTATCTAAATATATGCAAACAGAGAAAAGGTACAGTAAAAGTATGGTATAAAAGATAATGGT
ATACCTGTGTAGGCCACTTACCACGAATGGAGCTTGCAGGACTAGAAGTTGCTCTGGGTGAGTCAGTGAGTGAGTGGTGAATTAATGTGAAGGCCTAGAACACTGTACA
CCACTGTAGACTATAAACACAGTACGCTGAAGCTACACCAAATTTATCTTAACAGTTTTTCTTCAATAAAAAAATTATAACTTTTTAACTTTGTAAACTTTTTAATTTTT
TAACTTTTAAAATACTTAGCTTGAAACACAAATACATTGTATAGCTATACAAAAATATTTTTTCTTTGTATCCTTATTCTAGAAGCTTTTTTCTATTTTCTATTTTAAA
TTTTTTTTTTTACTTGTTAGTCGTTTTTGTTAAAAACTAAAACACACACACTTTCACCTAGGCATAGACAGGATTAGGATCATCAGTATCACTCCCTTCCACCTCACTG
CCTTCCACCTCCACATCTTGTCCCACTGGAAGGTTTTTAGGGGCAATAACACACATGTAGCTGTCACCTATGATAACAGTGCTTTCTGTTGAATACCTCCTGAAGGACT
TGCCTGAGGCTGTTTTACATTTAACTTAAAAAAAAAAAAAAGTAGAAGGAGTGCACTCTAAAATAACAATAAAAGGCATAGTATAGTGAATACATAAACCAGCAATGTAG
TAGTTTATTATCAAGTGTTGTACACTGTAATAATTGTATGTGCTATACTTTAAATAACTTGCAAAATAGTACTAAGACCTTATGATGGTTACAGTGTCACTAAGGCAAT
AGCATATTTTCAGGTCCATTGTAATCTAATGGGACTACCATCATATATGCAGTCTACCATTGACTGAAACGTTACATGGCACATAACTGTATTTGCAAGAATGATTTGT
TTTACATTAATATCACATAGGATGTACCTTTTTAGAGTGGTATGTTTATGTGGATTAAGATGTACAAGTTGAGCAAGGGGACCAAGAGCCCTGGGTTCTGTCTTGGATG
TGAGCGTTTATGTTCTTCTCCTCATGTCTGTTTTCTCATTAAATTCAAAGGCTTGAACGGGCCCTATTTAGCCCTTCTGTTTTCTACGTGTTCTAAATAACTAAAGCTT
TTAAATTCTAGCCATTTAGTGTAGAACTCTCTTTGCAGTGATGAAATGCTGTATTGGTTTCTTGGCTAGCATATTAAATATTTTTATCTTTGTCTTGATACTTCAATGT
CGTTTTAAACATCAGGATCGGGCTTCAGTATTCTCATAACCAGAGAGTTCACTGAGGATACAGGACTGTTTGCCCATTTTTTGTTATGGCTCCAGACTTGTGGTATTTC
CATGTCTTTTTTTTTTTTTTTTTTTTGACCTTTTAGCGGCTTTAAAGTATTTCTGTTGTTAGGTGTTGTATTACTTTTCTAAGATTACTTAACAAAGCACCACAAACT
GAGTGGCTTTAAACAACAGCAATTTATTCTCTCACAATTCTAGAAGCTAGAAGTCCGAAATCAAAGTGTTGACAGGGGCATGATCTTCAAGAGAGAAGACTCTTTCCTT
GCCTCTTCCTGGCTTCTGGTGGTTACCAGCAATCCTGAGTGTTCCTTTCTTGCCTTGTAGTTTCAACAATCCAGTATCTGCCTTTTGTCTTCACATGGCTGTCTACCAT
TTGTCTCTGTGTCTCCAAATCTCTCTCCTTATAAACACAGCAGTTATTGGATTAGGCCCCACTCTAATCCAGTATGACCCCATTTTAACATGATTACACTTATTTCTAG
ATAAGGTCACATTCACGTACACCAAGGGTTAGGAATTGAACATATCTTTTTGGGGGACACAATTCAACCCACAAGTGTCAGTCTCTAGCTGAGCCTTTCCCTTCCTGTT
TTTCTCCTTTTTAGTTGCTATGGGTTAGGGGCCAAATCTCCAGTCATACTAGAATTGCACATGGACTGGATATTTGGGAATACTGCGGGTCTATTCTATGAGCTTTAGT
ATGTAACATTTAATATCAGTGTAAAGAAGCCCTTTTTTAAGTTATTTCTTTGAATTTCTAAATGTATGCCCTGAATATAAGTAACAAGTTACCATGTCTTGTAAAATGA
TCATATCAACAAACATTTAATGTGCACCTACTGTGCTAGTTGAATGTCTTTATCCTGATAGGAGATAACAGGATTCCACATCTTTGACTTAAGAGGACAAACCAAATAT
GTCTAAATCATTTGGGGTTTTGATGGATATCTTTAAATTGCTGAACCTAATCATTGGTTTCATATGTCATTGTTTAGATATCTCCGGAGCATTTGGATAATGTGACAGT
TGGAATGCAGTGATGTCGACTCTTTGCCCACCGCCATCTCCAGCTGTTGCCAAGACAGAGATTGCTTTAAGTGGCAAATCACCTTTATTAGCAGCTACTTTTGCTTACT
GGGACAATATTCTTGGTCCTAGAGTAAGGCACATTTGGGCTCCAAAGACAGAACAGGTACTTCTCAGTGATGGAGAAATAACTTTTCTTGCCAACCACACTCTAAATGG
AGAAATCCTTCGAAATGCAGAGAGTGGTGCTATAGATGTAAAGTTTTTTGTCTTGTCTGAAAAGGGAGTGATTATTGTTTCATTAATCTTTGATGGAAACTGGAATGGG
GATCGCAGCACATATGGACTATCAATTATACTTCCACAGACAGAACTTAGTTTCTACCTCCCACTTCATAGAGTGTGTGTTGATAGATTAACACATATAATCCGGAAAG
GAAGAATATGGATGCATAAGGTAAGTGATTTTTCAGCTTATTAATCATGTTAACCTATCTGTTGAAAGCTTATTTTCTGGTACATATAAATCTTATTTTTTTAATTATA
TGCAGTGAACATCAAACAATAAATGTTATTTATTTTGCATTTACCCTATTAGATACAAATACATCTGGTCTGATACCTGTCATCTTCATATTAACTGTGGAAGGTACGA
AATGGTAGCTCCACATTATAGATGAAAAGCTAAAGCTTAGACAAATAAAGAAACTTTTAGACCCTGGATTCTTCTTGGGAGCCTTTGACTCTAATACCTTTTGTTTCCC
TTTCATTGCACAATTCTGTCTTTTGCTTACTACTATGTGTAAGTATAACAGTTCAAAGTAATAGTTTCATAAGCTGTTGGTCATGTAGCCTTTGGTCTCTTTAACCTCT
TTGCCAAGTTCCCAGGTTCATAAAATGAGGAGGTTGAATGGAATGGTTCCCAAGAGAATTCCTTTTAATCTTACAGAAATTATTGTTTTCCTAAATCCTGTAGTTGAAT
ATATAATGCTATTTACATTTCAGTATAGTTTTGATGTATCTAAAGAACACATTGAATTCTCCTTCCTGTGTTCCAGTTTGATACTAACCTGAAAGTCCATTAAGCATTA
CCAGTTTTAAAAGGCTTTTGCCCAATAGTAAGGAAAAATAATATCTTTTAAAAGAATAATTTTTTACTATGTTTGCAGGCTTACTTCCTTTTTTCTCACATTATGAAAC
TCTTAAAATCAGGAGAATCTTTTAAACAACATCATAATGTTTAATTTGAAAAGTGCAAGTCATTCTTTTCCTTTTTGAAACTATGCAGATGTTACATTGACTGTTTTCT
GTGAAGTTATCTTTTTTTCACTGCAGAATAAAGGTTGTTTTGATTTTATTTTGTATTGTTTATGAGAACATGCATTTGTTGGGTTAATTTCCTACCCCTGCCCCCATTT
TTTCCCTAAAGTAGAAAGTATTTTTCTTGTGAACTAAATTACTACACAAGAACATGTCTATTGAAAAATAAGCAAGTATCAAAATGTTGTGGGTTGTTTTTTTAAATAA
ATTTTCTCTTGCTCAGGAAAGACAAGAAAATGTCCAGAAGATTATCTTAGAAGGCACAGAGAGAATGGAAGATCAGGTATATGCAAATTGCATACTGTCAAATGTTTTT
CTCACAGCATGTATCTGTATAAGGTTGATGGCTACATTTGTCAAGGCCTTGGAGACATACGAATAAGCCTTTAATGGAGCTTTTATGGAGGTGTACAGAATAAACTGGA
GGAAGATTTCCATATCTTAAACCCAAAGAGTTAAATCAGTAAACAAAGGAAAATAGTAATTGCATCTACAAATTAATATTTGCTCCCTTTTTTTTTCTGTTTGCCCAGA
ATAAATTTTGGATAACTTGTTCATAGTAAAAATAAAAAAAATTGTCTCTGATATGTTCTTTAAGGTACTACTTCTCGAACCTTTCCCTAGAAGTAGCTGTAACAGAAGG
AGAGCATATGTACCCCTGAGGTATCTGTCTGGGGTGTAGGCCCAGGTCCACACAATATTTCTTCTAAGTCTTATGTTGTATCGTTAAGACTCATGCAATTTACATTTTA
TTCCATAACTATTTTAGTATTAAAATTTGTCAGTGATATTTCTTACCCTCTCCTCTAGGAAAATGTGCCATGTTTATCCCTTGGCTTTGAATGCCCCTCAGGAACAGAC
ACTAAGAGTTTGAGAAGCATGGTTACAAGGGTGTGGCTTCCCCTGCGGAAACTAAGTACAGACTATTTCACTGTAAAGCAGAGAAGTTCTTTTGAAGGAGAATCTCCAG
TGAAGAAAGAGTTCTTCACTTTTACTTCCATTTCCTCTTGTGGGTGACCCTCAATGCTCCTTGTAAAACTCCAATATTTTAAACATGGCTGTTTTGCCTTTCTTTGCTT
CTTTTTAGCATGAATGAGACAGATGATACTTTAAAAAAGTAATTAAAAAAAAAAAACTTGTGAAAATACATGGCCATAATACAGAACCCAATACAATGATCTCCTTTACC
AAATTGTTATGTTTGTACTTTTGTAGATAGCTTTCCAATTCAGAGACAGTTATTCTGTGTAAAGGTCTGACTTAACAAGAAAAGATTTCCCTTTACCCAAAGAATCCCA
GTCCTTATTTGCTGGTCAATAAGCAGGGTCCCCAGGAATGGGGTAACTTTCAGCACCCTCTAACCCACTAGTTATTAGTAGACTAATTAAGTAAACTTATCGCAAGTTG
AGGAAACTTAGAACCAACTAAAATTCTGCTTTTACTGGGATTTTGTTTTTTCAAACCAGAAACCTTTACTTAAGTTGACTACTATTAATGAATTTTGGTCTCTCTTTTA
AGTGCTCTTCTTAAAAATGTTATCTTACTGCTGAGAAGTTCAAGTTTGGGAAGTACAAGGAGGAATAGAAACTTAAGAGATTTTCTTTTAGAGCCTCTTCTGTATTTAG
CCCTGTAGGATTTTTTTTTTTTTTTTTTTTTTGGTGTTGTTGAGCTTCAGTGAGGCTATTCATTCACTTATACTGATAATGTCTGAGATACTGTGAATGAAATACTAT
GTATGCTTAAACCTAAGAGGAAATATTTTCCCAAAATTATTCTTCCCGAAAAGGAGGAGTTGCCTTTTGATTGAGTTCTTGCAAATCTCACAACGACTTTATTTTGAAC
AATACTGTTTGGGGATGATGCATTAGTTTGAAACAACTTCAGTTGTAGCTGTCATCTGATAAAATTGCTTCACAGGGAAGGAAATTTAACACGGATCTAGTCATTATTC
TTGTTAGATTGAATGTGTGAATTGTAATTGTAAACAGGCATGATAATTATTACTTTAAAAACTAAAAACAGTGAATAGTTAGTTGTGGAGGTTACTAAAGGATGGTTTT
TTTTTAAATAAAACTTTCAGCATTATGCAAATGGGCATATGGCTTAGGATAAAACTTCCAGAAGTAGCATCACATTTAAATTCTCAAGCAACTTAATAATATGGGGCTC
TGAAAAACTGGTTAAGGTTACTCCAAAAATGGCCCTGGGTCTGACAAAGATTCTAACTTAAAGATGCTTATGAAGACTTTGAGTAAAATCATTTCATAAAATAAGTGAG
GAAAAACAACTAGTATTAAATTCATCTTAAATAATGTATGATTTAAAAAATATGTTTAGCTAAAAATGCATAGTCATTTGACAATTTCATTTATATCTCAAAAAAATTTA
CTTAACCAAGTTGGTCACAAAACTGATGAGACTGGTGGTGGTAGTGAATAAATGAGGGACCATCCATATTTGAGACACTTTACATTTGTGATGTGTTATACTGAATTTT
CAGTTTGATTCTATAGACTACAAATTTCAAAATTACAATTTCAAGATGTAATAAGTAGTAATATCTTGAAATAGCTCTAAAGGGAATTTTTCTGTTTTATTGATTCTTA
AAATATATGTGCTGATTTTGATTTGCATTTGGGTAGATTATACTTTTATGAGTATGGAGGTTAGGTATTGATTCAAGTTTTCCTTACCTATTTGGTAAGGATTTCAAAG
TCTTTTTGTGCTTGGTTTTCCTCATTTTTAAATATGAAATATATTGATGACCTTTAACAAATTTTTTTTATCTCAAATTTTAAAGGAGATCTTTTCTAAAAGAGGCATG
ATGACTTAATCATTGCATGTAACAGTAAACGATAAACCAATGATTCCATACTCTCTAAAGAATAAAAGTGAGCTTTAGGGCCGGGCATGGTCAGAAATTTGACACCAAC

CTGGCCAACATGGCGAAACCCCGTCTCTACTAAAAATACAAAAATCAGCCGGGCATGGTGGCGGCACCTATAGTCCCAGCTACTTGGGAGGATGAGACAGGAGAGTCAC
TTGAACCTGGGAGGAGAGGTTGCAGTGAGCTGAGATCACGCCATTGCACTCCAGCCTGAGCAATGAAAGCAAAACTCCATCTCAAAAAAAAAAAAAGAAAAGAAAGAAT
AAAAGTGAGCTTTGGATTGCATATAAATCCTTTAGACATGTAGTAGACTTGTTTGATACTGTGTTTGAACAAATTACGAAGTATTTTCATCAAAGAATGTTATTGTTTG
ATGTTATTTTTATTTTTTATTGCCCAGCTTCTCTCATATTACGTGATTTTCTTCACTTCATGTCACTTTATTGTGCAGGGTCAGAGTATTATTCCAATGCTTACTGGAG
AAGTGATTCCTGTAATGGAACTGCTTTCATCTATGAAATCACACAGTGTTCCTGAAGAAATAGATGTAAGTTTAAATGAGAGCAATTATACACTTTATGAGTTTTTTGG
GGTTATAGTATTATTATGTATATTATTAATATTCTAATTTTAATAGTAAGGACTTTGTCATACATACTATTCACATACAGTATTAGCCACTTTAGCAAATAAGCACACA
CAAAATCCTGGATTTTATGGCAAAACAGAGGCATTTTTGATCAGTGATGACAAAATTAAATTCATTTTGTTTATTTCATTACTTTTATAATTCCTAAAAGTGGGAGGAT
CCCAGCTCTTATAGGAGCAATTAATATTTAATGTAGTGTCTTTTGAAACAAAACTGTGTGCCAAAGTAGTAACCATTAATGGAAGTTTACTTGTAGTCACAAATTTAGT
TTCCTTAATCATTTGTTGAGGACGTTTTGAATCACACACTATGAGTGTTAAGAGATACCTTTAGGAAACTATTCTTGTTGTTTTCTGATTTTGTCATTTAGGTTAGTCT
CCTGATTCTGACAGCTCAGAAGAGGAAGTTGTTCTTGTAAAAATTGTTTAACCTGCTTGACCAGCTTTCACATTTGTTCTTCTGAAGTTTATGGTAGTGCACAGAGATT
GTTTTTTGGGGAGTCTTGATTCTCGGAAATGAAGGCAGTGTGTTATATTGAATCCAGACTTCCGAAAACTTGTATATTAAAAGTGTTATTTCAACACTATGTTACAGCC
AGACTAATTTTTTATTTTTTGATGCATTTTAGATAGCTGATACAGTACTCAATGATGATGATATTGGTGACAGCTGTCATGAAGGCTTTCTTCTCAAGTAAGAATTTT
TCTTTTCATAAAAGCTGGATGAAGCAGATACCATCTTATGCTCACCTATGACAAGATTTGGAAGAAAGAAAATAACAGACTGTCTACTTAGATTGTTCTAGGGACATTA
CGTATTTGAACTGTTGCTTAAATTTGTGTTATTTTTCACTCATTATATTTCTATATATATTTGGTGTTATTCCATTTGCTATTTAAAGAAACCGAGTTTCCATCCCAGA
CAAGAAATCATGGCCCCTTGCTTGATTCTGGTTTCTTGTTTTACTTCTCATTAAAGCTAACAGAATCCTTTCATATTAAGTTGTACTGTAGATGAACTTAAGTTATTTA
GGCGTAGAACAAAATTATTCATATTTATACTGATCTTTTTCCATCCAGCAGTGGAGTTTAGTACTTAAGAGTTTGTGCCCTTAAACCAGACTCCCTGGATTAATGCTGT
GTACCCGTGGGCAAGGTGCCTGAATTCTCTATACACCTATTTCCTCATCTGTAAAATGGCAATAATAGTAATAGTACCTAATGTGTAGGGTTGTTATAAGCATTGAGTA
AGATAAATAATATAAAGCACTTAGAACAGTGCCTGGAACATAAAAACACTTAATAATAGCTCATAGCTAACATTTCCTATTTACATTTCTTCTAGAAATAGCCAGTATT
TGTTGAGTGCCTACATGTTAGTTCCTTTACTAGTTGCTTTACATGTATTATCTTATATTCTGTTTTAAAGTTTCTTCACAGTTACAGATTTTCATGAAATTTTACTTTT
AATAAAAGAGAAGTAAAAGTATAAAGTATTCACTTTTATGTTCACAGTCTTTTCCTTTAGGCTCATGATGGAGTATCAGAGGCATGAGTGTGTTTAACCTAAGAGCCTT
AATGGCTTGAATCAGAAGCACTTTAGTCCTGTATCTGTTCAGTGTCAGCCTTTCATACATCATTTTAAATCCCATTTGACTTTAAGTAAGTCACTTAATCTCTCTACAT
GTCAATTTCTTCAGCTATAAAATGATGGTATTTCAATAAATAAATACATTAATTAAATGATATTATACTGACTAATTGGGCTGTTTTAAGGCTCAATAAGAAAATTTCT
GTGAAAGGTCTCTAGAAAATGTAGGTTCCTATACAAATAAAAGATAACATTGTGCTTATAGCTTCGGTGTTTATCATATAAAGCTATTCTGAGTTATTTGAAGAGCTCA
CCTACTTTTTTTTGTTTTTAGTTTGTTAAATTGTTTTATAGGCAATGTTTTTAATCTGTTTTCTTTAACTTACAGTGCCATCAGCTCACACTTGCAAACCTGTGGCTGT
TCCGTTGTAGTAGGTAGCAGTGCAGAGAAAGTAAATAAGGTAGTTTATTTTATAATCTAGCAAATGATTTGACTCTTTAAGACTGATGATATATCATGGATTGTCATTT
AAATGGTAGGTTGCAATTAAAATGATCTAGTAGTATAAGGAGGCAATGTAATCTCATCAAATTGCTAAGACACCTTGTGGCAACAGTGAGTTTGAAATAAACTGAGTAA
GAATCATTTATCAGTTTATTTTGATAGCTCGGAAATACCAGTGTCAGTAGTGTATAAATGGTTTTGAGAATATATTAAAATCAGATATATAAAAAAAATTACTCTTCTA
TTTCCCAATGTTATCTTTAACAAATCTGAAGATAGTCATGTACTTTTGGTAGTAGTTCCAAAGAAATGTTATTTGTTTATTCATCTTGATTTCATTGTCTTCGCTTTCC
TTCTAAATCTGTCCCTTCTAGGGAGCTATTGGGATTAAGTGGTCATTGATTATTATACTTTATTCAGTAATGTTTCTGACCCTTTCCTTCAGTGCTACTTGAGTTAATT
AAGGATTAATGAACAGTTACATTTCCAAGCATTAGCTAATAAACTAAAGGATTTTGCACTTTTCTTCACTGACCATTAGTTAGAAAGAGTTCAGAGATAAGTATGTGTA
TCTTTCAATTTCAGCAAACCTAATTTTTTAAAAAAAGTTTTACATAGGAAATATGTTGGAAATGATACTTTACAAAGATATTCATAATTTTTTTTGTAATCAGCTACT
TTGTATATTTACATGAGCCTTAATTTATATTTCTCATATAACCATTTATGAGAGCTTAGTATACCTGTGTCATTATATTGCATCTACGAACTAGTGACCTTATTCCTTC
TGTTACCTCAAACAGGTGGCTTTCCATCTGTGATCTCCAAAGCCTTAGGTTGCACAGAGTGACTGCCGAGCTGCTTTATGAAGGGAGAAAGGCTCCATAGTTGGAGTGT
TTTTTTTTTTTTTTTAAACATTTTTCCCATCCTCCATCCTCTTGAGGGAGAATAGCTTACCTTTTATCTTGTTTTAATTTGAGAAAGAAGTTGCCACCACTCTAGGTT
GAAAACCACTCCTTTAACATAATAACTGTGGATATGGTTTGAATTTCAAGATAGTTACATGCCTTTTTATTTTTCCTAATAGAGCTGTAGGTCAAATATTATTAGAATC
AGATTTCTAAATCCCACCCAATGACCTGCTTATTTTAAATCAAATTCAATAATTAATTCTCTTCTTTTTGGAGGATCTGGACATTCTTTGATATTTCTTACAACGAATT
TCATGTGTAGACCCACTAAACAGAAGCTATAAAAGTTGCATGGTCAAATAAGTCTGAGAAAGTCTGCAGATGATATAATTCACCTGAAGAGTCACAGTATGTAGCCAAA
TGTTAAAGGTTTTGAGATGCCATACAGTAAATTTACCAAGCATTTTCTAAATTTATTTGACCACAGAATCCCTATTTTAAGCAACAACTGTTACATCCCATGGATTCCA
GGTGACTAAAGAATACTTATTTCTTAGGATATGTTTTATTGATAATAACAATTAAAATTTCAGATATCTTTCATAAGCAAATCAGTGGTCTTTTTACTTCATGTTTTAA
TGCTAAAATATTTTCTTTTATAGATAGTCAGAACATTATGCCTTTTTCTGACTCCAGCAGAGAGAAAATGCTCCAGGTTATGTGAAGCAGAATCATCATTTAAATATGA
GTCAGGGCTCTTTGTACAAGGCCTGCTAAAGGTATAGTTTCTAGTTATCACAAGTGAAACCACTTTTCTAAAATCATTTTTGAGACTCTTTATAGACAAATCTTAAATA
TTAGCATTTAATGTATCTCATATTGACATGCCCAGAGACTGACTTCCTTTACACAGTTCTGCACATAGACTATATGTCTTATGGATTTATAGTTAGTATCATCAGTGAA
ACACCATAGAATACCCTTTGTGTTCCAGGTGGGTCCCTGTTCCTACATGTCTAGCCTCAGGACTTTTTTTTTTTTAACACATGCTTAAATCAGGTTGCACATCAAAAAT
AAGATCATTTCTTTTTAACTAAATAGATTTGAATTTTATTGAAAAAAAAATTTTAAACATCTTTAAGAAGCTTATAGGATTTAAGCAATTCCTATGTATGTGTACTAAAA
TATATATATTTCTATATATAATATATATTAGAAAAAAATTGTATTTTTCTTTTATTTGAGTCTACTGTCAAGGAGCAAAACAGAGAAATGTAAATTAGCAATTATTTAT
AATACTTAAAGGGAAGAAAGTTGTTCACCTTGTTGAATCTATTATTGTTATTTCAATTATAGTCCCAAGACGTGAAGAAATAGCTTTCCTAATGGTTATGTGATTGTCT
CATAGTGACTACTTTCTTGAGGATGTAGCCACGGCAAAATGAAATAAAAAAATTTAAAAATTGTTGCAAATACAAGTTATATTAGGCTTTTGTGCATTTTCAATAATGT
GCTGCTATGAACTCAGAATGATAGTATTTAAATATAGAAACTAGTTAAAGGAAACGTAGTTTCTATTTGAGTTATACATATCTGTAAATTAGAACTTCTCCTGTTAAAG
GCATAATAAAGTGCTTAATACTTTTGTTTCCTCAGCACCCTCTCATTTAATTATATAATTTTAGTTCTGAAAGGGACCTATACCAGATGCCTAGAGGAAATTTCAAAAC
TATGATCTAATGAAAAAATATTTAATAGTTCTCCATGCAAATACAAATCATATAGTTTTCCAGAAAATACCTTTGACATTATACAAAGATGATTATCACAGCATTATAA
TAGTAAAAAAATGGAAATAGCCTCTTTCTTCTGTTCTGTTCATAGCACAGTGCCTCATACGCAGTAGGTTATTATTACATGGTAACTGGCTACCCCAACTGATTAGGAA
AGAAGTAAATTTGTTTTATAAAAATACATACTCATTGAGGTGCATAGAATAATTAAGAAATTAAAAGACACTTGTAATTTTGAATCCAGTGAATACCCACTGTTAATAT
TTGGTATATCTCTTTCTAGTCTTTTTTTCCCTTTTGCATGTATTTTCTTTAAGACTCCCACCCCCACTGGATCATCTCTGCATGTTCTAATCTGCTTTTTTCACAGCAG
ATTCTAAGCCTCTTTGAATATCAACACAAACTTCAACAACTTCATCTATAGATGCCAAATAATAAATTCATTTTTATTTACTTAACCACTTCCTTTGGATGCTTAGGTC
ATTCTGATGTTTTGCTATTGAAACCAATGCTATACTGAACACTTCTGTCACTAAAACTTTGCACACACTCATGAATAGCTTCTTAGGATAAATTTTTAGAGATGGATTT
GCTAAATCAGAGACCATTTTTTAAAATTAAAAAACAATTATTCATATCGTTTGGCATGTAAGACAGTAAATTTTCCTTTTATTTTGACAGGATTCAACTGGAAGCTTTG
TGCTGCCTTTCCGGCAAGTCATGTATGCTCAGCCATATCCCACCACACACATAGATGTGGATGTCAATACTGTGAAGCAGATGCCACCCTGTCATGAACATATTTATAATCA
GCGTAGATACATGAGATCCGAGCTGACAGCCTTCTGGAGAGCCACTTCAGAAGAAGACATGGCTCAGGATACGATCATCTACACTGACGAAAGCTTTACTCCTGATTTG
TACGTAATGCTCTGCCTGCTGGTACTGTAGTCAAGCAATATGAAATTGTGTCTTTTACGAATAAAAACAAAACAGAAGTTGCATTTAAAAAGAAAGAAATATTACCAGC
AGAATTATGCTTGAAGAAACATTTAATCAAGCATTTTTTTTCTTAAATGTTCTTCTTTTTCCATACAATTGTGTTTACCCTAAAATAGGTAAGATTAACCCTTAAAGTAA
ATATTTAACTATTTGTTTAATAAATATATATTGAGCTCCTAGGCACTGTTCTAGGTACCGGGCTTAATAGTGGCCAACCAGACAGCCCCAGCCCCAGCCCCTACATTGT
GTATAGTCTATTATGTAACAGTTATTGAATGGACTTATTAACAAAACCAAAGAAGTAATTCTAAGTCTTTTTTTTTCTTGACATATGAATATAAAATACAGCAAAACTGT
TAAAATATATTAATGGAACATTTTTTTACTTTGCATTTTATATTGTTATTCACTTCTTATTTTTTTTTAAAAAAAAAAAGCCTGAACAGTAAATTCAAAAGGAAAAGTAA
TGATAATTAATTGTTGAGCATGGACCCAACTTGAAAAAAAAAATGATGATGATAAATCTATAATCCTAAAACCCCTAAGTAAACACTTAAAAGATGTTCTGAAATCAGGA
AAAGAATTATAGTATACTTTTGTGTTTCTCTTTTATCAGTTGAAAAAAGGCACAGTAGCTCATGCCTGTAAGAACAGAGCTTTGGGAGTGCAAGGCAGGCGGATCACTT
GAGGCCAGGAGTTCCAGACCAGCCTGGGCAACATAGTGAAACCCCATCTCTACAAAAAATAAAAAAGAATTATTGGAATGTGTTTCTGTGTGCCTGTAATCCTAGCTAT
TCCGAAAGCTGAGGCAGGAGGATCTTTTGAGCCCAGGAGTTTGAGGTTACAGGGAGTTATGATGTGCCAGTGTACTCCAGCCTGGGGAACACCGAGACTCTGTCTTATT
TAAAAAAAAAAAAAAAAAAAATGCTTGCAATAATGCCTGGCACATAGAAGGTAACAGTAAGTGTTAACTGTAATAACCCAGGTCTAAGTGTGTAAGGCAATAGAAAAATT
GGGGCAAATAAGCCTGACCTATGTATCTACAGAATCAGTTTGAGCTTAGGTAACAGACCTGTGGAGCACCAGTAATTACACAGTAAGTGTTAACCAAAAGCATAGAATA
GGAATATCTTGTTCAAGGGACCCCCAGCCTTATACATCTCAAGGTGCAGAAAGATGACTTAATATAGGACCCATTTTTTCCTAGTTCTCCAGAGTTTTTATTGGTTCTT
GAGAAAGTAGTAGGGGAATGTTTTAGAAAATGAATTGGTCCAACTGAAATTACATGTCAGTAAGTTTTTATATATTGGTAAATTTTAGTAGACATGTAGAAGTTTTCTA
ATTAATCTGTGCCTTGAAACATTTTCTTTTTTCCTAAAGTGCTTAGTATTTTTTCCGTTTTTTGATTGGTTACTTGGGAGCTTTTTTGAGGAAATTTAGTGAACTGCAG
AATGGGTTTGCAACCATTTGGTATTTTTGTTTTGTTTTTTAGAGGATGTATGTGTATTTTAACATTTCTTAATCATTTTTAGCCAGCTATGTTTGTTTTGCTGATTTGA
CAAACTACAGTTAGACAGCTATTCTCATTTTGCTGATCATGACAAAATAATATCCTGAATTTTTAAATTTGCATCCAGCTCTAAATTTTCTAAACATAAAATTGTCCA
AAAAATAGTATTTTCAGCCACTAGATTGTGTGTTAAGTCTATTGTCACAGAGTCATTTTACTTTTAAGTATATGTTTTTACATGTTAATTATGTTTGTTATTTTTAATT
TTAACTTTTTAAAATAATTCCAGTCACTGCCAATACATGAAAAATTGGTCACTGGAATTTTTTTTTTGACTTTTATTTTAGGTTCATGTGTACATGTGCAGGTGTGTTA
TACAGGTAAATTGCGTGTCATGAGGGTTTGGTGTACAGGTGATTTCATTACCCAGGTAATAAGCATAGTACCCAATAGGTAGTTTTTTGATCCTCACCCTTCTCCCACC
CTCAAGTAGGCCCTGGTGTTGCTGTTTCCTTCTTTGTGTCCATGTATACTCAGTGTTTAGCTCCCACTTAGAAGTGAGAACATGCGGTAGTTGGTTTTCTGTTCCTGGA

TTAGTTCACTTAGGATAATGACCTCTAGCTCCATCTGGTTTTTATGGCTGCATAGTATTCCATGGTGTATATGTATCACATTTTCTTTATCCAGTCTACCATTGATAGG
CATTTAGGTTGATTCCCTGTCTTTGTTATCATGAATAGTGCTGTGATGAACATACACATGCATGTGTCTTTATGGTAGAAAAATTTGTATTCCTTTAGGTACATATAGA
ATAATGGGGTTGCTAGGGTGAATGGTAGTTCTATTTTCAGTTATTTGAGAAATCTTCAAACTGCTTTTCATAATAGCTAAACTAATTTACAGTCCCGCCAGCAGTGTAT
AAGTGTTCCCTTTTCTCCACAACCTTGCCAACATCTGTGATTTTTTGACTTTTTAATAATAGCCATTCCTAGAGAATTGATTTGCAATTCTCTATTAGTGATATTAAGC
ATTTTTTCATATGCTTTTTAGCTGTCTGTATATATTCTTCTGAAAAATTTTCATGTCCTTTGCCCAGTTTGTAGTGGGGTGGGTTGTTTTTTGCTTGTTAATTAGTTTT
AAGTTCCTTCCAGATTCTGCATATCCCTTTGTTGGATACATGGTTTGCAGATATTTTTCTCCCATTGTGTAGGTTGTCTTTTACTCTGTTGATAGTTTCTTTTGCCATG
CAGGAGCTCGTTAGGTCCCATTTGTGTTTGTTTTTGTTGCAGTTGCTTTTGGCGTCTTCATCATAAAATCTGTGCCAGGGCCTATGTCCAGAATGGTATTTCCTAGGTT
GTCTTCCAGGGTTTTTACAATTTTAGATTTTACGTTTATGTCTTTAATCCATCTTGAGTTGATTTTTGTATATGGCACAAGGAAGGGGTCCAGTTTCACTCCAATTCCT
ATGGCTAGCAATTATCCCAGCACCATTTATTGAATACGGAGTCCTTTCCCCATTGCTTGTTTTTTGTCAACTTTGTTGAAGATCAGATGGTTGTAAGTGTGTGGCTTTA
TTTCTTGGCTCTCTATTCTCCATTGGTCTATGTGTCTGTTTTTATAACAGTACCCTGCTGTTCAGGTTCCTATAGCCTTTTAGTATAAAATCGGCTAATGTGATGCCTC
CAGCTTTGTTCTTTTTGCTTAGGATTGCTTTGGCTATTTGGGCTCCTTTTTGGGTCCATATTAATTTTAAAACAGTTTTTTCTGGTTTTGTGAAGGATATCATTGGTAG
TTTATAGGAATAGCATTGAATCTGTAGATTGCTTTGGGCAGTATGGCCATTTTAACAATATTAATTCTTCCTATCTATGAATATGGAATGTTTTTCCATGTGTTTGTGT
CATCTCTTTATACCTGATGTATAAAGAAAAGCTGGTATTATTCCTACTCAATCTGTTCCAAAAAATTGAGGAGGAGGAACTCTTCCCTAATGAGGCCAGCATCATTCTG
ATACCAAAACCTGGCAGAGACACAACAGAAAAAAGAAAACTTCAGGCCAATATCCTTGATGAATATAGATGCAAAAATCCTCAACAAAATACTAGCAAACCAAATCCAG
CAGCACATCAAAAAGCTGATCTACTTTGATCAAGTAGGCTTTATCCCTGGGATGCAAGGTTGGTTCAACATACACAAATCAATAAGTGTGATTCATCACATAAACAGAG
CTAAAAACAAAAACCACAAGATTATCTCAATAGGTAGAGAAAAGGTTGTCAATAAAATTTAACATCCTCCATGTTAAAAACCTTCAGTAGGTCAGGTGTAGTGACTCAC
ACCTGTAATCCCAGCACTTTGGGAGGCCAAGGCGGGCATATCTCTTAAGCCCAGGAGTTCAAGACGAGCCTAGGCAGCATGGTGAAACCCCATCTCTACAAAAAAAAAA
AAAAAAAAAAATTAGCTTGGTATGGTGACATGCACCTATAGTCCCAGCTATTCAGGAGGTTGAGGTGGGAGGATTGTTTGAGCCCGGGAGGCAGAGGTTGGCAGCGAGC
TGAGATCATGCCACCGCACTCCAGCCTGGGCAACGGAGTGAGACCCTGTCTCAAAAAAGAAAAATCACAAACAATCCTAAACAAACTAGGCATTGAAGGAACATGCCTC
AAAAAAATAAGAACCATCTATGACAGACCCATAGCCAATATCTTACCAAATGGGCAAAAGCTGGAAGTATTCTCCTTGAGAACCGTAACAAGACAAGGATGTCCACTCT
CACCACTCCTTTTCAGCATAGTTCTGGAAGTCCTAGCCAGAGCAATCAGGAAAGAGAAAGAAAGAAAGACATTCAGATAGGAAGAGAAGAAGTCAAACTATTTCTGTTT
GCAGGCAGTATAATTCTGTACCTAGAAAATCTCATAGTCTCTGCCCAGAAACTCCTAAATCTGTTAAAAATTTCAGCAAAGTTTTGGCATTCTCTATACTCCAACACCT
TCCAAAGTGAGAGCAAAATCAAGAACACAGTCCCATTCACAATAGCCGCAAAACGAATAAAATACCTAGGAATCCAGCTAACCAGGGAGGTGAAAGATCTCTATGAGAA
TTACAAAACACTGCTGAAAGAAATCAGAGATGACACAAACAAATGGAAATGTTCTTTTTTAACACCTTGCTTTATCTAATTCACTTATGATGAAGATACTCATTCAGTG
GAACAGGTATAATAAGTCCACTCGATTAAATATAAGCCTTATTCTCTTTCCAGAGCCCAAGAAGGGGCACTATCAGTGCCCAGTCAATAATGACGAAATGCTAATATTT
TTCCCCTTTACGGTTTCTTTCTTCTGTAGTGTGGTACACTCGTTTCTTAAGATAAGGAAACTTGAACTACCTTCCTGTTTGCTTCTACACATACCCATTCTCTTTTTTT
GCCACTCTGGTCAGGTATAGGATGATCCCTACCACTTTCAGTTAAAAACTCCTCCTCTTACTAAATGTTCTCTTACCCTCTGGCCTGAGTAGAACCTAGGGAAAATGGA
AGAGAAAAAGATGAAAGGGAGGTGGGGCCTGGGAAGGGAATAAGTAGTCCTGTTTGTTTGTGTGTTTGCTTTAGCACCTGCTATATCCTAGGTGCTGTGTTAGGCACAC
ATTATTTTAAGTGGCCATTATATTACTACTACTCACTCTGGTCGTTGCCAAGGTAGGTAGTACTTTCTTGGATAGTTGGTTCATGTTACTTACAGATGGTGGGCTTGTT
GAGGCAAACCCAGTGGATAATCATCGGAGTGTGTTCTCTAATCTCACTCAAATTTTTCTTCACATTTTTTGGTTTGTTTTGGTTTTTGATGGTAGTGGCTTATTTTTGT
TGCTGGTTTGTTTTTTGTTTTTTTTTGAGATGGCAAGAATTGGTAGTTTTATTTATTAATTGCCTAAGGGTCTCTACTTTTTTTAAAAGATGAGAGTAGTAAAATAGAT
TGATAGATACATACATACCCTTACTGGGGACTGCTTATATTCTTTAGAGAAAAAATTACATATTAGCCTGACAAACACCAGTAAAATGTAAATATATCCTTGAGTAAAT
AAATGAATGTATATTTTGTGTCTCCAAATATATATATCTATATTCTTACAAATGTGTTTATATGTAATATCAATTTATAAGAACTTAAAATGTTGGCTCAAGTGAGGGA
TTGTGGAAGGTAGCATTATATGGCCATTTCAACATTTGAACTTTTTTCTTTTCTTCATTTTCTTCTTTTCTTCAGGAATATTTTTCAAGATGTCTTACACAGAGACACT
CTAGTGAAAGCCTTCCTGGATCAGGTAAATGTTGAACTTGAGATTGTCAGAGTGAATGATATGACATGTTTTCTTTTTTAATATATCCTACAATGCCTGTTCTATATAT
TTATATTCCCCTGGATCATGCCCCAGAGTTCTGCTCAGCAATTGCAGTTAAGTTAGTTACACTACAGTTCTCAGAAGAGTCTGTGAGGGCATGTCAAGTGCATCATTAC
ATTGGTTGCCTCTTGTCCTAGATTTATGCTTCGGGAATTCAGACCTTTGTTTACAATATAATAAATATTATTGCTATCTTTTAAAGATATAATAATAAGATATAAAGTT
GACCACAACTACTGTTTTTTGAAACATAGAATTCCTGGTTTACATGTATCAAAGTGAAATCTGACTTAGCTTTTACAGATATAATATATACATATATATATCCTGCAAT
GCTTGTACTATATATGTAGTACAAGTATATATATATATGTTTGTGTGTGTATATATATATAGTACGAGCATATATACATATTACCAGCATTGTAGGATATATATATGTTTA
TATATTAAAAAAAAGTTATAAACTTAAAACCCTATTATGTTATGTAGAGTATATGTTATATATGATATGTAAAATATATAACATATACTCTATGATAGAGTGTAATATA
TTTTTTATATATATTTTAACATTTATAAAATGATAGAATTAAGAATTGAGTCCTAATCTGTTTTATTAGGTGCTTTTTGTAGTGTCTGGTCTTTCTAAAGTGTCTAAAT
GATTTTTCCTTTTGACTTATTAATGGGGAAGAGCCTGTATATTAACAATTAAGAGTGCAGCATTCCATACGTCAAACAACAAACATTTTAATTCAAGCATTAACCTATA
ACAAGTAAGTTTTTTTTTTTTTTTTGAGAAAGGGAGGTTGTTTATTTGCCTGAAATGACTCAAAAATATTTTTGAAACATAGTGTACTTATTTAAATAACATCTTTATT
GTTTCATTCTTTTAAAAAATATCTACTTAATTACACAGTTGAAGGAAATCGTAGATTATATGGAACTTATTTCTTAATATATTACAGTTTGTTATAATAACATTCTGGG
GATCAGGCCAGGAAACTGTGTCATAGATAAAGCTTTGAAATAATGAGATCCTTATGTTTACTAGAAATTTTGGATTGAGATCTATGAGGTCTGTGACATATTGCGAAGT
TCAAGGAAAATTCGTAGGCCTGGAATTTCATGCTTCTCAAGCTGACATAAAATCCCTCCCACTCTCCACCTCATCATATGCACACATTCTACTCCTACCCACCCACTCC
ACCCCCTGCAAAAGTACAGGTATATGAATGTCTCAAAACCATAGGCTCATCTTCTAGGAGCTTCAATGTTATTTGAAGATTTGGGCAGAAAAAATTAAGTAATACGAAA
TAACTTATGTATGAGTTTTAAAAGTGAAGTAAACATGGATGTATTCTGAAGTAGAATGCAAAATTTGAATGCATTTTTAAAGATAAATTAGAAAACTTCTAAAAACTGT
CAGATTGTCTGGGCCTGGTGGCTTATGCCTGTAATCCCAGCACTTTGGGAGTCCGAGGTGGGTGGATCACAAGGTCAGGAGATCGAGACCATCCTGCCAACATGGTGAA
ACCCCGTCTCTACTAAGTATACAAAAATTAGCTGGGCGTGGCAGCGTGTGCCTGTAATCCCAGCTACCTGGGAGGCTGAGGCAGGAGAATCGCTTGAACCCAGGAGGTG
TAGGTTGCAGTGAGTCAAGATCGCGCCACTGCACTTTAGCCTGGTGACAGAGCTAGACTCCGTCTCAAAAAAAAAAAAAAAATATCAGATTGTTCCTACACCTAGTGCTT
CTATACCACACTCCTGTTAGGGGGCATCAGTGGAAATGGTTAAGGAGATGTTTAGTGTGTATTGTCTGCCAAGCACTGTCAACACTGTCATAGAAACTTCTGTACGAGT
AGAATGTGAGCAAATTATGTGTTGAAATGGTTCCTCTCCCTGCAGGTCTTTCAGCTGAAACCTGGCTTATCTCTCAGAAGTACTTTCCTTGCACAGTTTCTACTTGTCC
TTCACAGAAAAGCCTTGACACTAATAAAATATATAGAAGACGATACGTGAGTAAAACTCCTACACGGAAGAAAAACCTTTGTACATTGTTTTTTTGTTTTGTTTCCTTT
GTACATTTTCTATATCATAATTTTTGCGCTTCTTTTTTTTTTTTTTTTTTTTTTTTTTCCATTATTTTTAGGCAGAAGGGAAAAAAGCCCTTTAAATCTCTTCGGAACC
TGAAGATAGACCTTGATTTAACAGCAGAGGGCGATCTTAACATAATAATGGCTCTGGCTGAGAAAATTAAACCAGGCCTACACTCTTTTATCTTTGGAAGACCTTTCTA
CACTAGTGTGCAAGAACGAGATGTTCTAATGACTTTTTAACCGTGTGGTTTGCTGTGTCTGTCTCTTCACAGTCACACCTGCTGTTACAGTGTCTCAGCAGTGTGTGGG
CACATCCTTCCTCCCGAGTCCTGCTGCAGGACAGGGTACACTACACTTGTCAGTAGAAGTCTGTACCTGATGTCAGGTGCATCGTTACAGTGAATGACTCTTCCTAGAA
TAGATGTACTCTTTTAGGGCCTTATGTTTACAATTATCCTAAGTACTATTGCTGTCTTTTAAAGATATGAATGATGGAATATACACTTGACCATAACTGCTGATTGGTT
TTTTGTTTTGTTTTGTTTGTTTTCTTGGAAACTTATGATTCCTGGTTTACATGTACCACACTGAAACCCTCGTTAGCTTTACAGATAAAGTGTGAGTTGACTTCCTGCC
CCTCTGTGTTCTGTGGTATGTCCGATTACTTCTGCCACAGCTAAACATTAGAGCATTTAAAGTTTGCAGTTCCTCAGAAAGGAACTTAGTCTGACTACAGATTAGTTCT
TGAGAGAAGACACTGATAGGGCAGAGCTGTAGGTGAAATCAGTTGTTAGCCCTTCCTTTATAGACGTAGTCCTTCAGATTCGGTCTGTACAGAAATGCCGAGGGGTCAT
GCATGGGCCCTGAGTATCGTGACCTGTGACAAGTTTTTGTTGGTTTATTGTAGTTCTGTCAAAGAAAGTGGCATTTGTTTTTATAATTGTTGCCAACTTTTAAGGTTA
ATTTTCATTATTTTTGAGCCGAATTAAAATGCGCACCTCCTGTGCCTTTCCCAATCTTGGAAAATATAATTTCTTGGCAGAGGGTCAGATTTCAGGGCCCAGTCACTTT
CATCTGACCACCCTTTGCACGGCTGCCGTGTGCCTGGCTTAGATTAGAAGTCCTTGTTAAGTATGTCAGAGTACATTCGCTGATAAGATCTTTGAAGAGCAGGGAAGCG
TCTTGCCTCTTTCCTTTGGTTTCTGCCTGTACTCTGGTGTTTCCCGTGTCACCTGCATCATAGGAACAGCAGAGAAATCTGACCCAGTGCTATTTTTCTAGGTGCTACT
ATGGCAAACTCAAGTGGTCTGTTTCTGTTCCTGTAACGTTCGACTATCTCGCTAGCTGTGAAGTACTGATTAGTGGAGTTCTGTGCAACAGCAGTGTAGGAGTATACAC
AAACACAAATATGTGTTTCTATTTAAAACTGTGGACTTAGCATAAAAAGGGAGAATATATTTATTTTTTACAAAAGGGATAAAAATGGGCCCCGTTCCTCACCCACCAG
ATTTAGCGAGAAAAAGCTTTCTATTCTGAAAGGTCACGGTGGCTTTGGCATTACAAATCAGAACAACACACACTGACCATGATGGCTTGTGAACTAACTGCAAGGCACT
CCGTCATGGTAAGCGAGTAGGTCCCACCTCCTAGTGTGCCGCTCATTGCTTTACACAGTAGAATCTTATTTGAGTGCTAATTGTTGTCTTTGCTGCTTTACTGTGTTGT
TATAGAAAATGTAAGCTGTACAGTGAATAAGTTATTGAAGCATGTGTAAACACTGTTATATATCTTTTCTCCTAGATGGGGAATTTTGAATAAAATACCTTTGAAATTC
TGTGTATGTTTTAGTTCATTATTTAGGGAAAACGCTGCTGTGAAAGGGGGCGTGATCAGCTTCCTATTCTGCGACAGTCGTGTTGAACGGAACCCATTGGTTTTCATCT
TCGCTCCCCCCCCCTTGGTTTTTCGAGACAGGGTTTCTCTGTATAGCCCTGGCTGTCCTGGACCTCACTCTGTAGACCAGGCTGGCCTCGAACTCAGAAATCTACCTGC
CTCTGCCTCCCAAGTGCTGGGAGGCAGTTGCCCCACCAACTAGTCTTCTTTTTTCAAAGAAGATATTTAAAGCTAACGAATAATGCTAGACTCTTACATCTTAAAAAAA
AAAGAAGAGAAAAGAAAAGAAAAGGTAATCACACTGCCCAGTGTGTAGTGCATGCTTCTACTTCCGGTCCTTGGGAGATGGGGGCAGGATGAGACGCTCCAGACCGGCT
TCCAATACAGAGTTCAAGACCCACTGAGCTACGTGAGGCTACACGAGCCTGCCTTTAAAAACATAAAGCTAAAGCTTTCTTCTTAACTTCCAGTATTGCACCTTGATTC
CCCCTTCAAATTTCACATACAAAATAATTCTTAAATTCTCTTTTGAAAAATGTTCTACTGAGGCCAGAGAGACAGTTCGCTTGGTAAAGGTGCCTGTTGCCAAACGTGA
TAACCTGAGTTAAATCATAGCCCCACATGGGGGAGGAAGAAACCCCCGCAGCTTGCCCTCTGATGCCATGTATGCACTAAAACACGCACGTGTGTGCGCACACATTTTT

TAAGTTCCTATTACATTGATAGTAATATAATTTAAACTGATTTATTCTCCCCAAGTCATTGATACGGGTGTCCAACGTAAAATCCAGCGGCTGAACAAAGCACTTTTAG
GCGCTTTAAGTTGGAAAGCAAGAAACGGAGATTGACACTGTCACTCCAAGAGAAAACTCTTCGTAGTAGCGAGATCGGCTGTGGAGTGAAGATGCTCAGAGGCTGGGAA
CGCACACAGCTCAGGAGTGGATAGCATCCCCCAGCCTCAACTCCTAACACTGGGAAAGCGTAGGGCTCTCAGATGAGGAAACAAAACCATACAAAGCTGCTGCAAGCTA
AACAGAAAAATAGTGGCATTACACTAACTGTTGTGGAATTGTACAGACCGATTCTCCTCCCAATCTGCCGAGTGTGGGCGGCTTGAGAGAATGAAGAGAGCTACTGGCC
TCAGGTAACAGTGCTTCCCACAGGACTGTCTCAGGCTGCCACCACCATAAATAGCATTTTAGACGTGACAGAGCTAAGGCTTGACACACAGCCAAAAGCTACTCACATT
CCATTTCATCCCCAGCTGTTCTGTCATCGCTAAGCACAGAGCATTCAGCACAGCTCTTCCCTGTGGTGGGTACTCAGCACTGTTGAGTTGAAAGGATTGAAAAAACTCA
AGACTATGTTCTCAAACATTTTTTTAAGCTCTTTTTAAAACCACCTTAGAATGAAAGCTTTTGACTTCTTATTAACATGCACTAACTTCATATACACATTTAGTGTTAT
TGTACAGGCACGAAGCATACTCTGGTCAGAACCTGTCTCCTTTGGTCCACCCTCCCCACCGTTTTCAGCTTCTATTCCACCTTCCATACGTCTCAAGATCCACATGTGA
GAGGGAACACTCAGAGCCTTGTCTTTCTGTATCTGGGATATCTCACTTAACATGATATTCTCCAGTTCTGTTCCATCCATTTCATTGCAAAGAGCAAGATTTCACTCTA
CAGCCAAATAACACATTTGTCCATGTATATCCGTATTTTTCCTTATTCATCTGTTGAATGGCACAAGACTGATATCATGGGTAATATCTATGTGGACTTCTAGTCTCTG
AGAACAGAGGCACAGTAAATATGACAGTGATCTTTTTGGTATAATTTCACTTCCTTTGCTTATATGTCCAGAAATGGGATTGAATGATCATCTGGCAGATCCATTTGTG
TGTGTGTGTGTGTGTGTGTGTGTGTGTGTGTGTGTGTGTGTGTGTGTGTATGTGTGTGTGTGTTCACATGTGTAGGCATAAATGTAGGAGTATGTATATAGACAAATGGCC
CATTCCTGTAAAGGACAAGTTGACACTGGATATCTTCCTCAGTGACTGACTACTTTATTTACTGAGGCTGGGTTGTTTGCTGAACATGGAGCTTGTAGACTCAGCTAGT
CTAGGGAGCTTATCCCAAGGGTCGTCTGTCTTCAGCTTTCAAACACTGGGATTATGAGGTGCCTGGCCGGCTTTTTTAATGTGGGTGCTAAGAGTCTGCACTACAGTCC
TTTTGCTTGTATAGCAAGCACTTTACCCAGATCCATTTCCCCATCCATCCCCCATGTTAGCTTTTTCAGGAACCCCCCTATACTATTCTCCATGATAGCCCTACTGAT
TTACACTTAATTACACTGTAGCAAAACACCTGAGAAAAGTCACTTTACAGAAGGACATACACATTTGGGTCTATGAGGTCCGTGGTCACTTGGTCTGTTGTTTCTGAAC
CTGTAGTAAAACAGAGCACCCTGCTTGGGAGGGTACAGAGGAGCAAAGCTGTTGATTAAGGCCGTCCCTACAAGCTCTATGCTAGCAAAGAAATGAACTCCGTGAAACA
GGAAGGGGACAGAGCTCAGCGTGAGCCTGACACCAGGCCAGTTTAGTATAGCGGTATTTAAGTACAGTACAATTTTGGAAAAAAAAGGTTTTCTGGAAAAAAACTATCCCA
ATCCTATGTACACAATAAAGTTTAAGGCATGACTGCATCAAAAGTCTTTTGCAAAGTACATGAGACCTCTACCTTGAAGATGGATCTATAGCCCAGGTTCCTGTGGCCT
CGAATGGAAGTCAGCAAGTTATAAAATACATGTGATATCCTCCTCAGGATGGGTTCCACTGGCTGCAAAGCAAAGATAAAGCTAAAGGTCTAGTTGGGGAGAGTGTGGG
ATAAACATTCTGGGGAATTTCAGTGAAGCCCAGCCCTACAGGGCGCAAAGGTCACCAGGTTTTTAACAACATCCACTCGCAGCCTTCTGGGTGGGAAACAGGTACACGG
CTCCTTCATTGAGGAGGTGACAATGCCTGATTAGCATTCCTGGTTTCTCCAGTGCTTATCTGTGAGCATTCGCGCCCACTACTAGTATATCATGTAGCCAGGATGCAGA
GATAAAGGAGCCAAGGACAAACTCCACCTTCGACAACACTCTCTCGAGAACTGCTTCCTCCAGCTAGAACCCACTTCCTAAATCTACAAACTTGGCCTGGTTGGATTTT
TGTCAGCTTGACACAAGCTAGAGTTACCTGGGAAAGGGGACCTCATTTGAGAAAATGCCTCCATCAGATGGTCCATAGGCCAGTCgggccattgggccctagacaagtt
gtcgcggccgcactagtgactccagcgtaactggactggccacagttaggccgcaaatgtaatcacactggctcaccttcgggtgggcctttctgcgtctagaggatcc
cctgttatgcccttccttacccatgcgctgagaggtgaaatcactctttaatttgtacgcggtacggacctctatatcgctttgtcgtagaggtggaatctccccagct
tgcagccattgatagactgcgccgggtgtcacgcccacgccaaccgctgctttttccacatcgccgaaatggcgcacaagttcttcgggtttcatggggattattataa
tcaataagtgaaaattaaagctaggtataatttataaaaattatagccagctataaagagatcatttatgattaagggtatgaaaacacgaggcgaacgactgaaagca
cgccgtttagagctgaaactgacgctgaagcaagtggcagaagctgtaggaatctctcttccgggcgtccaaaacttagaacgtggcgacgtgatgccgtcgctggaga
tcgggctatcgctggcaaaatgcctgcgtaagcccgtgcaatggatactgtatggcactgaatctgatccagaccgcgttcctgttattggtacgacagagaccggacc
ggatagagactggcagcctggagaacctgccaacacagagcgattcctgccgttcgtgagtcaacggaacaccgtttacgcactgacggtcgggaaccagattcagcga
aactaccagccgggagacgttatcctggtcgattcttcacttacgcttgtaccgggtgaggatgtactggtttgtgataataacggcgagatcacaattcaacgattag
cccgttatgacgagcagcattactaccttgatagtgctaactctcaacgggttatccatgataaaagtgatcttcaatttgtgcaccaagtagtcggtacgatcaaatc
gttcatggttgagggtagatgatacaataacagggtttattgctgactataattctggtttaatccgatctatactttgtcggggttgaaccagaccgtagcagcccgaa
aaaagacgaaaaaaaacccgagtcggcaaactcgggcctttttcaggaagtagccacgataacgcagatacgtccttcagaaagatagtgcgtttattgtggctgctcc
tgcggatttttttcaacccgaaaaaatgctaattcgcatggaaaggctaaaaaatgaccttacaagaattctacgcggagcgctttggcagcgatccgttttcactgctt
gacgcggcacgtgatgagctgactgagctggcgaagatggccggtatcaactggcctgcgtgtgctgataagattcagctcaatccgcgcggtggtgtagagcgctata
ccacctacaataattcctccccggaagctctgcagaagagccttaaaggccgtgtggaaatctattcacgccttgagcagagcaaggatggcatcagttatcctttcgt
taacttcgttgaaaaagcgcatgacgctggctcctggagtggcttttcattcctgttttcggagtatcgccgtgaacagcaacggaatggtgcgaccgtggtcgctcag
ccggaagaagaacgtgcacgcatggagcgccaggctgaagcacgccgcctccgagctgaacaacaacgcgttaatgatttaaaaaacaatcagatggagcaggaacgcc
tgcttggatggttggctttccatcgcgcctgggaacattcgccagctgaagatggctcctggccttatgcggtaaaaaagggaattcgtgacgtatttggcgcttgtga
tatacgtcgcgtgaccagtcacgacagtgcgaaatggagccgtggaccaacaacttacatggcgatacctctgtcccacctggacggaagaaaagacgggcggattgtt
ggctggcaacgtatcgatctgaatggtggcaagttccagaccagcgccatcactaacggtgatttcgtcggtgcatgctttgttattggcgacctgcagggggcgcaaa
aaattgcgactgcagagggtttcgccaccggggcatccatctggctggcgaccagaaacgacccgaaaaaacgctttgacgcggtagtcattgcggtttccgctaacaa
catgattcacgttgtcgagcagctggtgaacatgtacccggctgcgcaaatcacttgcgctctcgataacgaccgcaaatcctcagctgaaggaaaaggcaacacaggc
ctccgcaccggattcgacatcatggagaagttttccggcgtcaaatgtgtttacccaacttttgaggatgaccctgagctggaatgcagcgattttaacgacctgcaca
gcctgagagggctgaaggaagtcgctcgccagttaacgagaaatcatctgagccgtgcaaccgatctgttgtcgatcacgctgaataagctgcgtactctcccgcgtct
gaacagacgcactttgccaaagaactgcttcgcgccgttgatattggcatgctgacatgcccggtaccaaacagcccgaaagaacttatgcgccttttcagcagcacg
ctgcgggatatgggtatcgcagaaatttataacggtaccgttaaagatcacattacgcgccggttgaatcgcaaatgccgtgccgcgcaaacatcacgttcgttcagtg
aacgcatcaccaacccgaaccttcgcccgtcacacatcacttacaaacggtttgaaacctccaggatgaccgatgaggtgatgacatacgccgcacagctgcagggcat
cgttattgtccgcgcccgggatggggtcgggtaaatcgacaggcctcctgcgtccactgatgctgcagtccacgcgtggcgtttccgtcgcgcaccgcgtatccctgata
ggcggcctgcatgaaatgatgaccgaagggaaaggcgcgaaagccgacattctgcattatcaggatcccggctatcaggaaatggcgccatatgccaacaagctgacta
tttgcatcaactccatacttaaaggctgctggcaaccgctgatgcgccagcatgacttcttcggcttcgatgaagcaacacagggcctgcgtgccattctggccgggcg
tgcgatggaaaacccggtaggcgtattcaacacgcttatcgacgcgctggcgcgtactgaagagcatgccattatggtggatgccgatgccaacgatctgcttgttgac
ctggctgaactggcgatgaagcgacgcgaggagctgggcctacctgcctggctgcaaattcacgtgattgaactcccggtcgacgttcgcaaccgcgaaaccaacaagc
ctatccgcgtattttataccgagaaaaatcggatcatgaccgaggttatcgctgcagtgcaacgcggcgaacgaatcatgctggccaccgacagttcgacgttcgccga
agacgttaccatgcagctgagactacagttccctgacaaaaagtttctctgcgttaaccagaaaaacaaacaggagaaggaagtcgacgatttcaccaaccagcctaaa
gtgatggtgaaaaaatatgacggcctcatctacagcccgtcgatatcttcaggtgtatcgattgaggagaaacacttccaccgccatttcggcatgttctgcggcgaag
tggtccccagcgacgccatccagatgctgcgccgcgaccgtacagctcaggaatacattatcggtttcgacaagcttcgcggtaaacgtgaaaccgatccggaaaaaat
caaacgcgcctacgcgcaggccttgctcgaaacggctggccactccgggctgctgacagacgttgtctttgacggtgaccgaatttcactcggcgtggctaactcttca
ttcatgcagctcaaaattaaagccgccgcgctcgaggcatcggccagaaatgattatgccagcaacatgatttgcatcatgcatgatgatggctatcaggtcgcgccta
tggccaccgacgcgcttgcaaacagcatcggtaaggatttacgtaaagaagcccgtgaactggtctttgagcagcttatggagcgccacctgagtgtcgatactcctga
ccaggccgaacacgatgagttgataaaaaaaacgcaccctgtccctggatgagcaggcccagctggtccgctgggacatcgaaaaggagctgcagctggatgtcgacgag
gtggcgttaaaattctatttcgacggcggcctgaaaaaagtgcgcctgtttgaaaccatgcagctggatgagataaccgcccgtcgccttgaccgtgaagaagcgttga
ttcactttacctacgcttatcgcgttgccggaagatggcagcagttcgtcactacggccatgacgcgggaacaggccgacgcagagttccaggctaaattcccggccat
aaccgattaccgagtgaaatcgacccctgctgttgagataggcatgcgtggtttctacaccctcaaatcggcgacgctgcagcagtacttccgcgactgcggtatcgat
ccaaaaaccctggaaggtgaagcagacatggacgccctcaaacgcgccagagataacctgctcacgccggaacggcgcgatctgctaaacaacgttttacgcatcggcg
gcttcaacacagagaaaggcaagaagaaagcgcctgcagacctctgcataggcatcctggagtctatggggctatccagtaagaccagacgcgccagagacggcgatgc
acgccccacgatgcgttccattgacccggactcggtcgagttcctcatgaatatcgtggagaagcgccgcgaggccggtttatcaattcacgcacgtaaggtggaaaaa
accaccatcgaagtggatcgcgatttggatctaaatatagatatacatggtaaccctcgatccaaaacagagcacgttccggacgccccacaatcagtaattatccagg
cactggaggctatcccggtggcggtaccggaggcgtgggcggagaacgcgttgccagcaaccgaaatggaggcggtacgcctgtggccagtggccagcatcgcgagaac
gttcgcctcgctgtacatgaccgaatttatggacctgctatcagtacgcgaaataaggctgctgaaagcgtttctaagccagcggcaagccgtggcgatataaccggag
gggtaatgggaaaacaacgggaatgccttgtcatcgtggagggctacgacccttatgtggtcaaggtcagccacgacgcgacaagctatagcgattataaaactggccg
ggaacttccattcacgatcattcggatgaacgggttctcagatagtgagggagtttattgctgcattgccagcgataagagcatggaagaccacacagacgaagccagg
gcgcttctaaaagcctataaggtgaagccactatgacgtcgacaccagagttcctgaagcaactggattacgaacagctgaagtactgtcgcgagctgtgcgataaccg
catccgggctattgaggccgaagagaaaaagatggcctgggcagttactgatggtggcattaactacggctggttccgtacagaagactacatgaaggcggtagaatgc
ctggttgctacagctgcagaacggtgggaagaatcggataaggctgacccatctggacgtggttggcttgacctttcaatccgtggaagccgcctgccggtgtccgagt
atgaagctttatttgctgatggccagtgggggtgaatagtggcgcatgatctacaaaaaatgatggatggtgagctggtccgcttacaacttgagattaagaaagaaca

ggagaggcgtttagccgaaaagatgattcctgtttttggtgtgtggcaatcacctattaatcaatatgaatttgctgatccagagcaggcaatttcttgtgccgcaggg
ctattggataaggttcttcgtgaggtcaaggacgatctttccaaagaaggtggattacagggatggaacggcaatttgctacgcatctacgtacagcatgtgaacgaga
gcgattttgcagttctacagccttatttaaaggacaaataatcatgatttatcgcagaggatgggtgcctgttttgtggaggaatgagctggaaaagcggctcaaagaa
gaagggttcgaaaactggaagcagataagtaactttctttgtggtgaagaggattgctactcggaagcagcaaaaacgccaggacaatacgcttaccaggtcgtcgaca
ataccgaatggatggaacgtcgtgatgcgacgttctggcagcgtctgaaccgtttgtggttcgttccgctgtacctgctgactattccgttccagtggctcattcgtgg
ccgtatggggtttgagacaacatctaaggtggctgcggtcatcaataagattaccggactccaataactgatcaggttctgatcagtcagtaaagcaaactataaaaac
acctgtgcacgtgactggtcacggggtaaaatatccacctaaattattgacgtgcgttctgctttggcggtagagttaccccgctgcagcaaaatctgcagccgggcct
ctcaaccccgaatgacaagaagcgcacaacacgcgccagcgtgtttttttgtgtgttaaatctgcgcatacctgaattatggtggctcagatggggccaacttcggttg
ggccggtttcttcttgtaccggtgttgagaaccctgtctgggctaccacccctatagagattctcaactctggtggtagcaccctatacaagataggaatgcatacc at
gttcaaattcaaatttgcagctgtcgtccgcacggacaaaaaatcccatatccaccgcctttccaccattgcatcatccgagcgagaagctcgtcgccagttcgccagc
cgttttgtcctcgttctgtcagcccgtatcccggtcagcgaggtgatcgcatgaaccagattcagttaaacgcccagggcctgctggagacgattgaggaacgcctgct
gcaggttgaagcgcttgtgggctctgctcaccgcactatctccagttatgaggccacgctgtatttgcaggaagctgcggaacttctgcaggtcgcgcgcgaactcacg
caggaagctcgcggctgctccctgtccctttctgagcagctgaaatccggggaggcgcaatgaaggctctctccgtcttttcatttcaggaaagccaccctatacgggt
ggttcttgtcggtggtgacccgtggtttgtggcgctggatatctgtgctgcattgaatatagccaacccttctgatgcattgcgtaagctggatcatgatgaaaaattg
accctcggtttaaccgaggcacaaaaacttgatcggatggccagggaagtaaatgtggtatcggaatctggtctctacaccataatcctccgctgccgcgacgcggtga
aacaggggacgacagcctggcggttccgcaagtgggtgacgaacgaggttctgccagctatccggaaaaacggcgagtatgcttttgtcgagcctgagcctaagaacgc
cggtgaaccactggactggcggcagaaagaagaacttcggggcctgataaacgacatagcacaaagttttcagtaccgtaacgcatgggttagcggcgtctggatggct
ctccgccgcgcctgcaggaacccatcaccgaacccaataaccgtggatgatctcccggcaatcatcgccgagctgcgccggatattaacggctgcagaaacggctctgg
gcaacatgcgtgtctacgaacgggagctgctgcgcgatgtagttcgtggcggtcgccggagtatgtcttgtggagagctgccgatcaccgatattgatacggaactgga
gaaggtgctgccagcacattttgagctggccattgaaaagctggaaacgctatccacgaaactaacatcgccagcgattgcagaaaaatagggttggccagcccgaatt
taagccccgctttacgcggggaagtccctgcagtcgggacggcgatgtccgtagttgcagctgcagtttccgcctaatccttccgtcccttgcaggggatgggtaggga
caggcctcaataaaaaagccaccgattaacaccggtggcttttcttttagctgtagtacgtttcccatgcggatcgcattgcggccatagggctatcggcggggccgga
ccaggcataatgctttccatcgtattcaaactctatcttgtacgttccgtccccgttatttttgcaggcttgaagacgggttttagagcagttggctgatgttcttct
ggcccttcctcttcggttggttcatcgccgccaccctcgtcgagctcaatttcatcttcatccagctcgtcgtcttgggactcatcatccggttcgtcgatggtctcaa
cggattcttcatcaggcaggacgattgcaggtgtctctatcttcagctgccactgcccgttctcgccaacgaactgccccaatgcatcagcggcaaactccaggtaccg
gctaatcatcgtcgggctaaatttaaaggcccggagagtgctgttggttatttttgctgatgggtcctgctccaccagctgcttaacggtttcatggagacggacgcca
gcgtcacctctggcaaagcctggcatttcatcgtccagtttctgcagagccaccagcctggtgttttcatccccaacttcaggtcgccaggttctggagaagttggcca
gcttgaactgcttatagtgcagctgggtgttctcatcgtcgtgtccgagaatctccatgaagaacacatcctcgtcgacgtttttccaccgtggatcgacgcggaagaa
catctcataagcgatgcgagcgtaaatagcgcggctatctttataaacacgacggtcatcgccgaaaaatgatttaacccaagggttaaatgcttttgctaaaatagca
tttatcctgccgttctcagaccttgtatcatcctttccatatcctttaacaacctcatcgaaatcagatgcagcagagcaagaacgcaattctgttaataattcaacga
ataattttgcttcgcataaagtataaatcgttctggttacgcttttatcttcagagcgtttttttagcttgccctgagaaattaaccgtatactttcctgaaacggcaaa
ttcaccctgaaacattatctcaatcattcttcgccctgataccgcagccagagcaaaggccaaaggtgccattccagaacgagtgtttaaactaaataaagtcgcagga
ttattcaaaatatcatagatagactgcatgtatgttgggtagtcaataaccacaacattacgcttcttctcgcgcagaacatcggcccatcgttgctgtatagatgtac
gctccgcagggcttagctgcagatggtacagaacctcatggttgaccttgagctggtgtagttcttctaacaacgcagagccttgttggaataacttataaagatagtc
acggcgctccttccaatcatcactgtttaaatcactaagagcaaaactccaatctggatatttttttattagtcttgctattttgcatcactgcctttagagcctatt
cttacatttgataactcttcggcaagcggcattatttctttcaatttagattgtaacgatgacatgtgctggcgaatattagccgtaggcatagaaagccatgaagata
attcttcgctgtaaagaggatacttttccgataatttattaatatttttatcaaagctatgatgtaatttatcatcaaaccgctttcttgccctgctcatataggcgtt
aaaagtattcgcggttattcttttctgcaatcctttcccacggaactttcttttatcattaaataacgcgttcttataccgtgcggctgcggctttaattctcttcgtt
ttgtcgccttgtgggcggtctgaggcatcaattgcctctacctcattcacaagcgtgttgatcaactcaccgattttaccttgctcatcccggttccccgtttctaa
gtctctggatatcgtaacacaatcaaagaacaagtgacgattaattatgctgctttcgtcacactataagagaacataatagcagaataacaaaggaacacaacaccgg
aatagatacatattgttctgcgcttatgttctccttatcagcacacaatagtccattatacgc

#### Targeting Construct 2

See Figure 2G. pJazz backbone in lowercase. Mouse sequence in **green**. Human sequence in **blue**, selection cassette in **magenta**, pathogenic repeat in **red**, AarI site mutation indicated by **gcTggtg**.

gcgtataatgggcaattgtgtgctgatatgttccgtaattgagatatgatatagtgcatatgttccgaattattgatacattaatgttgcattcctatgttgtgttctg
ttatggtgttcttagatagtgggggatgtcacatggattgatgttctttgattgtgtttcaatatagagacggaagagatagagatgcctcaaagtgtataaacagcgt
gaaaatcagcatgttcgcttaacttttgatagcaattggtatgatttattatcatttaaaatcattaagttaggttttaaatataccaacgtaagtaataccgttgcgc
ttagatgaagctaaatagcttctgtagcgcatcaaaaaattgatttgaaccttaactttgcgttttctaagtcactttgctttttttatgttgctttgctttaaaaacg
taaagtatgatgaaaaacacaaagataagggcatgtttgagaggaaacaggatgtcgttaattaatttgctgaaagactgtatcaatcgcggtcaggaaatgacgcgcg
ctattgccatcgcacagttcggcgatgacagcccggaagcgcgtcgtattactcgccgctggggtataacagaagttgcagacctgatcggcgtaacaccgcaggctat
cagagacgcggaaaaagctggtcgtctaccggctcctgattttgagatgaggggccgagtagaacgtcgcgctggctacacaattgaccaaattagccatatgcggagt
gtgtttggtaacccaaaccaaagacctgacgataaaaacccggttgtactttccgttatgtcacacaagggaggggtttataaacctcatctgcagtacaccaggcgc
aatggttagccttgcaagggcaccgcgtcctactcgttgaagggaacgacccgcaaggactgcatctatgtaccacggttacgttcctgatttgcacattcacgcaga
cgatacactgcttccatttacctgggtaaacgcgacaacgctgaatatgcgataaaaccaacatgctggccgggtctcgacataatccccagctgcctggcccttcac
cgcattgaaacagatctagcagaaagtcaaaagcctccgaccggaggcttttgacttctgtcacctaggttacgccccgccctctggaactcaggagtcgcgcgctagg
ggccggatcggcaggtgatacttaagtaagccggcttagctagcgggacaggtttcccgactggaaagcgggcagtgagcgcaacgcaattaatgtgagttagctcact
cattaggcaccccaggctttacactttatgcttccggctcgtatgttgtgtggaattgtgagcggacaacaatttcacacaggaaacagctatgaccatgattacgcca
agctatttaggtgagactatagaatactcaagcttgcatgcgatacgtatcgttaacgatggatccgacgcacgtgcgaattcgccctatagtgagtcgtattacaatt
cactggccgtcgttttacaacgtcgtgactgggaaaaccctggcgtcacccaacttaatcgccttgcagcacatcccccttcgccagctggcgtaatagcgaagaggc
ccgcaccgatcgcccttcccaacagttgcgcagctgaatggcgaatcttaagtaggcctccccacctgcatggccggggccggggccggggcgtggtagtgatcttatt
tcattatggtgaaagttggaacctcttacgtgccgatcagattaaaacgaaaggcccagtctttcgactgagcctttcgttttatttgaccatgttggtatgatttaaa
tggtcagttaatcagttctatgtaccagcaaggtccagttgtaagcggccgcttgacttcaagtctaatggccc**GCAAGGACCCTAGACTCAAGAAAATGTAAGGTTAC**
**TAAAGTCTGAAGGACACCAAAGTGAAATGACAATTAGCTTTGGTTTGAGGCATATTTGTTTTGTTCGATTTTTCTGTTTTTTAAAAGCATATAATCAGAAAGAATTATA**
**GAGCTCGATTGACAATTGCAAGATGAACATGTGAAATAGTATTCCTAGGTGTCAGACAAAATAGAAAATGCCAATCATTCTTAGACAAATGATTTTTATAACACATTAA**
**AATATAATAGGAAAATCTCACAAAATAAGTCTGGGGCAGGAAGATTGCTCCAAGTTTTAGGCCAACCTGGGCTACAGAGTGATATCCTGTTAGGTGTAGGTGAAGGAAG**
**AGGAATAGGAAAAAAAAAGACTCAAGGATAACACTTTTTGGTAACTGGAATGACAGTTCAATGAGACATCCGACATATTCCAATCACAGTCTGCTAAAAAGGAAATGAA**
**TCCACAGGTAACCATATGGTCTTGGGTTTTAATTAGTATGTGACAACACTGACAGTCAATTATTCCATACTGCTACTAAATACTGAAAACTCTGCTTACTAAATCCCAA**
**TAAATGCAATAAAATATCAACATAAACTGCAGAATGGCTGATGGTTACAGGTAGATGGTGACACTTGGCAGAGATGCTCGGGAGCACATTTGGAGCTGGCCTGAGAAGT**
**GTGTGGTAGGAGAGTGTGCATTACTGCTGTGTCTCCTCCCCATGGGACCAGCGCCATCAGCCTGCACTGCTGGCCGCATCTCTGGTCTATCCAAGTATTAGACATGTAC**
**AACCACAAATTCTCTCTTCTTCTAAAGAAACACAATTTCATATTCTCTTTATGATTGCACATTAACATCTAAGATTTAAGATGATGCTTTTGATTTCCTATGTTTTATG**
**AGGTGCATTCCTCCCCCTGAGACTCTATTTGGTTTTTTGTTTGTTTGTTTGTTTGTGTTTGTGTTTGCTTTTGGGTTTTTTTGTTTTTTTGTTTTGTGGGTTTTTTGGG**
**TTTTTTGTTTGTTTGGTTGGTTGATTGGTTGGTTGTTTTGGTTGTGTCCTTTAACAGCTGAGCCAAGTCTCCAGCCCAGGCTTCTGTCCTTAGAAATAGTTACATATAA**
**AGGTAAGGTGCACAGCAGCAATGCACAAGAACAACTGGTCAAAGATGACATTCCTCTCACTCACTAAGGACATTGTTCATAAAATCACGAGTCCAAAGACAAGCAGAAA**
**AGAAATGTGTATTGTGCCCTAATGACTTTTTTTTTTTGCAGTACTGAAGCAGTATCATGTGCTGTTAGGATTATAAAGTGTTGGCTCCAGACAGAAACTCAAGGGTTTT**
**CTATATCATTTCATCCTACTGGCTCCACACACTAAGATTCTATTCGTATTTCAAATGAAGGCAGTATATCATAAAATACATATTTATTAAGTCTTCAGTGAATAAGGAA**
**GCAACCACATGAAATTACATTTCCTAAGAGAAATGAAATGGTCCGTGGTTTATATAGCAATATTCTTCATCCATGTCTGTCATGAAATTGTTCACATGCCAGTCTCAGT**
**CACAACTAATTTCCTGTAATCTTGGTAAGTAGGAGCTAATCAACAAGCCCTTGGTTCCAGCCCTGCAGCAATGCACATCACACTTCTGAAACATATTAGCAGAGAAAAT**

GGAGAAAAGCCTTCCTGACAACTGGCATTTTTGATGAAGCAAGAAGGTTCTCCGTGTTCTGACATATTTATTGGAGTTATAAATACGTGTAGAGTGTAAGAGGGTTGTA
AGTCATTCATTACTGTATTTATTATATCTGAGTAGTTACCAAAAAAAAAAATGAAAGTAAAACCATCACTTCTAGAATGTACTGATCTAATTTTCTAGCTCGCCTGATT
GCACATAAACCTTACAGCAACCAGGTGACAGATTACTAGGCGTGCTATTTTCCTGCTGAGCAACAGTGTCCTGCTAAAAGACTGCACTGACCAGCTTCCTGTGGGGTCA
AGCAAGGAAGCCCCAGAGGAAGGGACTATCAGAGGAGTGCAATGTACCTATGGGAGTCACTTCCACTCCTATTAGAAGGGCTGCAGCCATTGCTTTCTGTGGGTGTTCT
GCCATCCACACACAAGGGGAACAAAATGCCCTTGTCATGGATACACAGTCTTACCTAGGGTCACACTGGCCACTCTCTCCATTCCCTCCTTTAAGCCCCCACAGAAAGC
TAGCACTGTGGACACTGAGTCACCAAGGAGTGAAGAACTGTCCAGTTCCGTGGTTAACCTAAAGCAGAGACGATACTTAATATCTGTTGTGGTTACAGAGCCCAACCTC
TTATTTTGCTTTAGAAATCCCCCTTATCGAGGAGTGTCTGCATGACTGAACATTAGTGAAACATGATAGCTGACATTTTTCTATATCGGGTCTTTAAAGGTGTTGTTCT
GGTTGGAAATAGGGGATAGAGTTCTCAGTTCCTGTTTGGAGAGATACTCCCTTTCTTTTTCTTTTCTTTCTTTCTTTCTTTCTTTCTTTCTTTCTTTCTTTCTTTCTTT
CTTTCTTTCTTTCTTTTAATGGATCCCCCCGCCCCCTTGATAGCCTTCAGCATGTAGTGTAGCCTAGGATAATCTTGAACTTCTAATCCTCCTGTCTCAGATTATAGG
TGTTTGCCATCTCGTCAGTTCGTTTAGTATTGGGGAGTAAACCAGTACGTGGTAGGCAAGCCCTCTGCCAAATGCCCTCCACTTTCAGCCCTGGAGATATAGTTCTATA
GCCAGCATATTTTCACTTAGAGCCTGGAGTTATTTGAGGTAGATTATTTGTAAAGTTTGAGAGGACAGGGTTAAAAATCTATCACACCAATATCATGAATCATATTTCT
ACATACAGTTAGCATCATTTTCTCTTTGGTTCTCTGGAGGAAAGATGTATTCTACCTTACACCTTTCAGGCAATACAACAATTATGAACCAGCATACCTACTTATTATT
TTCTTCCTGATTAATAAAAGTTAGCTACAATCCAATTAAAATATATAGGCTACTTTACCTATGCCTAAGTAATTGTTCCATCTGATGTGAAGAAAAAGGTCTCCACTCA
TTGATCACAAAATATAAGTTGTATCATGGTTTATTTCATTTCCTGTGGTCATACAATATTTTAGATCGTGATGTTTCATTTTTTTCTCCAAAGAGAGAAAAAGAGAGGA
ATATGGTAGGGACCAAACCCAGGGAGTCACACATGCTAGGCAAGCCCTCTACCCACGAAGCTACATCCCTAGTCCTTTCTTGGTTTGATTATTTTTTTTAAATAAAACA
TTTATTCTAGGACTGATATTACACAAAATAAGTTATTTGAGGGACACTGTTTTTGTCTCTAGGTAAAATTTTGAAGGAAAAAAAAAACACTAAGAAGGTATATTCCTTC
AAAGTTCCAGTCTTATTCTGAAGTGTAATGTTATGTTAGTTTGACTCACAGACAGGTTTTAAAGAAGGGCTTACTTCAAGAGGACACCAAACAAATACCTTCTATTCCT
AGTGGGCTCTGGAATCACAGAAAACTGACCCAATCAATTACATTGATAGCTCTGGCTTACTACAGACAAGCAAATTATCTTAAGTGTGCATGCATGCGCGTGTATGTGT
GTTAGTACCTAACACCCACCTGGGAACTTTTCAGCTTTTCAGTGTGGGATATAGTATAAACGTCTATTCCTCGTGTTGTGGATTAGCTGACTGGCCTCACTCAGCTGCC
TTCCTTACCTGCAAACTCACCCACTTTGACTACAGCATCGCACTCTTAACCCTAGCCTTCCAAACATGGTCCTATGCTATTTCTGTGTGTCTGGATGTATTTTTAACTC
TCAGATGTATACTTCATTTATGAGATATACATCTGAAGACCACGGTACAAAACACTGTAAGAACTTGATAGAATGACAACTGCTAGGTAAAAAAAAAAAAAAAAAAAAA
AAAAAAAAAAAAAAAAAGCATACAATACCTGGTGAGAGTTCTATTTTTACCGAAGGTGGTATTGATAGGTATTCTGTTATTAATGCCTTTCTTTTCCCTATAAATGATG
AAAAGTTGCTGGAAAATAATAAACACTACTCATCTGTAGTGAAAAGCCACAATACAGTTACAAACCAATCAATCAATCAATAAATCAGACGTCATGGTGTTCTTTTCCC
AAAGGTTAAAAACAAAGTGCACTGTGCTATTTGGCAAAAATGACGTTTAGAAGAAAACACGGTGACTACGCACAGAGGGTGGGGGAATCATTGTGCTTGTTGCGGAGT
GAACACGTACAGTGTGCACGCAGACTTACGGCATTTAACCGTGTCATAGGGACCAAAGGAAATCCACTCACTCACTAAATATTTGTTGAGCACCCACTACCTGCCAACT
CCCAAACAAAACAAAGCAAAACTACTTACAACCACAAACTACGCTTCGTAACCTAGATAGATAACGCAGGTGACACTATCTATCTAGGTTGAGCTCAGCTCTGCCCATG
CTTTTCCTGAGCGGCTCTTGGAAGAAAAGCTACAAAGCCCATGACAGCCTCCGCCTGGCCAGCTGCCACTGGCATCTCAAGGCTGGCAAAGCAAAGTGAAAGCGCCAAC
CCGGAACTTACGGAGTCCCACGAGGGAACCGCGGCGCGTCAAGCAGAGACGAGTTCCGCCCACGTGAAAGATGGCGTTTGTAGTGACAGCCATCCCAATTGCCCTTTCC
TTCTAGGTGGAAAGTGGTGTCTAGACAGTCCAGGGAGGGTGTGCGAGGGAGGTGCGTTTTGGTTGCCTCAGCTCGCAACTTAACTCCACAACGGTGACCAAGGACAAAA
GAAGGAAACAAGACTGCAGAGATCCGCACCGGGGAGCCCTGCAGATTCTGGGTGTGTGTTTTTGTTTTTCCCACCCTCTCTCCCCACTACTTGCTCTCACAGTACTCGC
TGAGGGTGAACAAGAAAAGACCTGATAAAGATTAACCCTAGAAAGATAGTCTGCGTAAAATTGACGCATGCATTCTTGAAATATTGCTCTCTCTTTCTAAATAGCGCGA
ATCCGTCGCTGTGCATTTAGGACATCTCAGTCGCCGCTTGGAGCTCCCGTGAGGCGTGCTTGTCAATGCGGTAAGTGTCACTGATTTTGAACTATAACGACCGCGTGAG
TCAAAATGACGCATGATTATCTTTTACGTGACTTTTAAGATTTAACTCATACGATAATTATATTGTTATTTCATGTTCTACTTACGTGATAACTTATTATATATATATT
TTCTTGTTATAGATATCGCGGCCGCTCTAGCCTCGAGGCTAGAACTAGTGGATCTCGAGCCCCAGCTGGTTCTTTCCGCCTCAGAAGCCATAGAGCCCACCGCATCCCC
AGCATGCCTGCTATTGTCTTCCCAATCCTCCCCCTTGCTGTCCTGCCCCACCCCACCCCCCAGAATAGAATGACACCTACTCAGACAATGCGATGCAATTTCCTCATTT
TATTAGGAAAGGACAGTGGGAGTGGCACCTTCCAGGGTCAAGGAAGGCACGGGGGAGGGGCAAACAACAGATGGCTGGCAACTAGAAGGCACAGTCGAGGCTGATCAGC
GAGCTCTAGAGCTCAGAAGAACTCGTCAAGAAGGCGATAGAAGGCGATGCGCTGCGAATCGGGAGCGGCGATACCGTAAAGCACGAGGAAGCGGTCAGCCCATTCGCCG
CCAAGCTCTTCAGCAATATCACGGGTAGCCAACGCTATGTCCTGATAGCGGTCCGCCACACCCAGCCGGCCACAGTCGATGAATCCAGAAAAGCGGCCATTTTCCACCA
TGATATTCGGCAAGCAGGCATCGCCATGGGTCACGACGAGATCCTCGCCGTCGGGCATGCGCGCCTTGAGCCTGGCGAACAGTTCGGCTGGCGCGAGCCCCTGATGCTC
TTCGTCCAGATCATCCTGATCGACAAGACCGGCTTCCATCCGAGTACGTGCTCGCTCGATGCGATGTTTCGCTTGGTGGTCGAATGGGCAGGTAGCCGGATCAAGCGTA
TGCAGCCGCCGCATTGCATCAGCCATGATGGATACTTTCTCGGCAGGAGCAAGGTGAGATGACAGGAGATCCTGCCCCGGCACTTCGCCCAATAGCAGCCAGTCCCTTC
CCGCTTCAGTGACAACGTCGAGCACAGCTGCGCAAGGAACGCCCGTCGTGGCCAGCCACGATAGCCGCGCTGCCTCGTCCTGCAGTTCATTCAGGGCACCGGACAGGTC
GGTCTTGACAAAAAGAACCGGGCGCCCCTGCGCTGACAGCCGGAACACGGCGGCATCAGAGCAGCCGATCGTCTGTTGTGCCCAGTCATAGCCGAATAGCCTCTCCACC
CAAGCGGCCGGAGAACCTGCGTGCAATCCATCTTGTTCAATGGCCGATCCCATGGTTTAGTTCCTCACCTTGTCGTATTATACTATGCCGATATACTATGCCGATGATT
AATTGTCAACAGGCTGCAGGTCGAAAGGCCCGGAGATGAGGAAGAGGAGAACAGCGCGGCAGACGTGCGCTTTTGAAGCGTGCAGAATGCCGGGCCTCCGGAGGACCTT
CGGGCGCCCGCCCCGCCCCTGAGCCCGCCCCTGAGCCCGCCCCCGGACCCACCCCTTCCCAGCCTCTGAGCCCAGAAAGCGAAGGAGCAAAGCTGCTATTGGCCGCTGC
CCCAAAGGCCTACCCGCTTCCATTGCTCAGCGGTGCTGTCCATCTGCACGAGACTAGTGAGACGTGCTACTTCCATTTGTCACGTCCTGCACGACGCGAGCTGCGGGGC
GGGGGGGAACTTCCTGACTAGGGGAGGAGTAGAAGGTGGCGCGAAGGGGCCACCAAAGAACGGAGCCGGTTGGCGCCTACCGGTGGATGTGGAATGTGTGCGAGGCCAG
AGGCCACTTGTGTAGCGCCAAGTGCCCAGCGGGGCTGCTAAAGCGCATGCTCCAGACTGCCTTGGGAAAAGCGCCTCCCCTACCCGGTAGAATTAATTCGATATCAAGC
TTGCTCGGATGATAGGGCCCTAACGTGTTTTTGCTTTGTTACTTTATAGAAGAAATTTTGAGTTTTTGTTTTTTTTTAATAAATAAATAAACATAAATAAATTGTTTGT
TGAATTTATTATTAGTATGTAAGTGTAAATATAATAAAACTTAATATCTATTCAAATTAATAAATAAACCTCGATATACAGACCGATAAAACACATGCGTCAATTTTAC
GCATGATTATCTTTAACGTACGTCACAATATGATTATCTTTCTAGGGTTAACCAGAAGAAAACAAGGAGGGAAACAACCGCAGCCTGTAGCAAGCTCTGGAACTCAGGA
GTCGCGCGCTAGGGGCCGGGGCCGGGGCCGGGGCCGGGGCCGGGGCCGGGGCCGGGGCCGGGGCCGGGGCCGGGGCCGGGGCCGGGGCCGGGGCCGGGGCCGGGGCCGG
GGCCGGGGCCGGGGCCGGGGCCGGGGCCGGGGCCGGGGCCGGGGCCGGGGCCGGGGCCGGGGCCGGGGCCGGGGCCGGGGCCGGGGCCGGGGCCGGGGCCGGGGCCGGG
GCCGGGGCCGGGGCCGGGGCCGGGGCCGGGGCCGGGGCCGGGGCCGGGGCCGGGGCCGGGGCCGGGGCCGGGGCCGGGGCCGGGGCCGGGGCCGGGGCCGGGGCCGGGG
CCGGGGCCGGGGCCGGGGCCGGGGCCGGGGCCGGGGCCGGGGCCGGGGCCGGGGCCGGGGCCGGGGCCGGGGCCGGGGCCGGGGCCGGGGCCGGGGCCGGGGCCGGGGC
CGGGGCCGGGGCCGGGGCCGGGGCCGGGGCCGGGGCCGGGGCCGGGGCCGGGGCCGGGGCCGGGGCCGGGGCCGGGGCCGGGGCCGGGGCCGGGGCCGGGGCCGGGGCC
GGGGCCGGGGCCGGGGCCGGGGCCGGGGCCGGGGCCGGGGCCGGGGCCGGGGCCGGGGCCGGGGCCGGGGCCGGGGCCGGGGCCGGGGCCGGGGCCGGGGCCGGGGCCG
GGGCCGGGGCCGGGGCCGGGGCCGGGGCCGGGGCCGGGGCCGGGGCCGGGGCCGGGGCCGGGGCCGGGGCCGGGGCCGGGGCCGGGGCCGGGGCCGGGGCCGGGGCCGG
GGCCGGGGCCGGGGCCGGGGCCGGGGCCGGGGCCGGGGCCGGGGCCGGGGCCGGGGCCGGGGCCGGGGCCGGGGCCGGGGCCGGGGCCGGGGCCGGGGCCGGGGCCGGG
GCCGGGGCCGGGGCCGGGGCCGGGGCCGGGGCCGGGGCCGGGGCCGGGGCCGGGGCCGGGGCCGGGGCCGGGGCCGGGGCCGGGGCCGGGGCCGGGGCCGGGGCCGGGG
CCGGGGCCGGGGCCGGGGCCGGGGCCGGGGCCGGGGCCGGGGCCGGGGCCGGGGCCGGGGCCGGGGCCGGGGCCGGGGCCGGGGCCGGGGCCGGGGCCGGGGCCGGGGC
CGGGGCCGGGGCCGGGGCCGGGGCCGGGGCCGGGGCCGGGGCCGGGGCCGGGGCCGGGGCCGGGGCCGGGGCCGGGGCCGGGGCCGGGGCCGGGGCCGGGGCCGGGGCC
GGGGCCGGGGCCGGGGCCGGGGCCGGGGCCGGGGCCGGGGCCGGGGCCGGGGCCGGGGCCGGGGCCGGGGCCGGGGCCGGGGCCGGGGCCGGGGCCGGGGCCGGGGCCG
GGGCCGGGGCCGGGGCCGGGGCCGGGGCCGGGGCCGGGGCCGGGGCCGGGGCCGGGGCCGGGGCCGGGGCCGGGGCCGGGGCCGGGGCCGGGGCCGGGGCCGGGGCCGG
GGCCGGGGCCGGGGCCGGGGCCGGGGCCGGGGCCGGGGCCGGGGCCGGGGCCGGGGCCGGGGCCGGGGCCGGGGCCGGGGCCGGGGCCGGGGCCGGGGCCGGGGCCGGG
GCCGGGGCCGGGGCCGGGGCCGGGGCCGGGGCCGGGGCCGGGGCCGGGGCCGGGGCCGGGGCCGGGGCCGGGGCCGGGGCCGGGGCCGGGGCCGGGGCCGGGGCCGGGG
CCGGGGCCGGGGCCGGGGCCGGGGCCGGGGCCGGGGCCGGGGCCGGGGCCGGGGCCGGGGCCGGGGCCGGGGCCGGGGCCGGGGCCGGGGCCGGGGCCGGGGCCGGGGC
CGGGGCCGGGGCCGGGGCCGGGGCCGGGGCCGGGGCCGGGGCCGGGGCCGGGGCCGGGGCCGGGGCCGGGGCCGGGGCCGGGGCCGGGGCCGGGGCCGGGGCCGGGGCC
GGGGCCGGGGCCGGGGCCGGGGCCGGGGCCGGGGCCGGGGCCGGGGCCGGGGCCGGGGCCGGGGCCGGGGCCGGGGCCGGGGCCGGGGCCGGGGCCGGGGCCGGGGCCG
GGGCCGGGGCCGGGGCCGGGGCCGGGGCCGGGGCCGGGGCCGGGGCCGGGGCCGGGGCCGGGGCCGGGGCCGGGGCCGGGGCCGGGGCCGGGGCCGGGGCCGGGGCCGG
GGCCGGGGCCGGGGCCGGGGCCGGGGCCGGGGCCGGGGCCGGGGCCGGGGCCGGGGCCGGGGCCGGGGCCGGGGCCGGGGCCGGGGCCGGGGCCGGGGCCGGGGCCGGG
GCCGGGGCCGGGGCCGGGGCCGGGGCCGGGGCCGGGGCCGGGGCCGGGGCCGGGGCCGGGGCCGGGGCCGGGGCCGGGGCCGGGGCCGGGGCCGGGGCCGGGGCCGGGG
CCGGGGCCGGGGCCGGGGCCGGGGCCGGGGCCGGGGCCGGGGCCGGGGCCGGGGCCGGGGCCGGGGCCGGGGCCGGGGCCGGGGCCGGGGCCGGGGCCGGGGCCGGGGC
CGGGGCCGGGGCCGGGGCCGGGGCCGGGGCCGGGGCCGGGGCCGGGGCCGGGGCCGGGGCCGGGGCCGGGGCCGGGGCCGGGGCCGGGGCCGGGGCCGGGGCCGGGGCC
GGGGCCGGGGCCGGGGCCGGGGCCGGGGCCGGGGCCGGGGCCGGGGCCGGGGCCGGGGCCGGGGCCGGGGCCGGGGCCGGGGCCGGGGCCGGGGCCGGGGCCGGGGCCG
GGGCCGGGGCCGGGGCCGGGGCCGGGGCCGGGGCCGGGGCCGGGGCCGGGGCCGGGGCCGGGGCCGGGGCCGGGGCCGGGGCCGGGGCCGGGGCCGGGGCCGGGGCCGG
GGCCGGGGCCGGGGCCGGGGCCGGGGCCGGGGCCGGGGCCGGGGCCGGGGCCGGGGCCGGGGCCGGGGCCGGGGCCGGGGCCGGGGCCGGGGCCGGGGCCGGGGCCGGG

GCCGGGGCCGGGGCCGGGGCCGGGGCCGGGGCCGGGGCCGGGGCCGGGGCCGGGGCCGGGGCCGGGGCCGGGGCCGGGGCCGGGGCCGGGGCCGGGGCCGGGGCCGGGG
CCGGGGCCGGGGCCGGGGCCGGGGCCGGGGCCGGGGCCGGGGCCGGGGCCGGGGCCGGGGCCGGGGCCGGGGCCGGGGCCGGGGCCGGGGCCGGGGCCGGGGCCGGGGC
CGGGGCCGGGGCCGGGGCCGGGGCCGGGGCCGGGGCCGGGGCCGGGGCCGGGGCCGGGGCCGGGGCCGGGGCCGGGGCCGGGGCCGGGGCCGGGGCCGGGGCCGGGGCC
GGGGCCGGGGCCGGGGCCGGGGCCGGGGCCGGGGCCGGGGCCGGGGCCGGGGCCGGGGCCGGGGCCGGGGCCGGGGCCGGGGCCGGGGCCGGGGCCGGGGCCGGGGCCG
GGGCCGGGGCCGGGGCCGGGGCCGGGGCCGGGGCCGGGGCCGGGGCCGGGGCCGGGGCCGGGGCCGGGGCCGGGGCCGGGGCCGGGGCCGGGGCCGGGGCCGGGGCCGG
GGCCGGGGCCGGGGCCGGGGCCGGGGCCGGGGCCGGGGCCGGGGCCGGGGCCGGGGCCGGGGCCGGGGCCGGGGCCGGGGCCGGGGCCGGGGCCGGGGCCGGGGCCGGG
GCCGGGGCCGGGGCCGGGGCCGGGGCCGGGGCCGGGGCCGGGGCCGGGGCCGGGGCCGGGGCCGGGGCCGGGGCCGGGGCCGGGGCCGGGGCCGGGGCCGGGGCCGGGG
CCGGGGCCGGGGCCGGGGCCGGGGCCGGGGCCGGGGCCGGGGCCGGGGCCGGGGCCGGGGCCGGGGCCGGGGCCGGGGCCGGGGCCGGGGCCGGGGCCGGGGCCGGGGC
CGGGGCCGGGGCCGGGGCCGGGGCCGGGGCCGGGGCCGGGGCCGGGGCCGGGGCCGGGGCCGGGGCCGGGGCCGGGGCCGGGGCCGGGGCCGGGGCCGGGGCCGGGGCC
GGGGCCGGGGCCGGGGCCGGGGCCGGGGCCGGGGCCGGGGCCGGGGCCGGGGCCGGGGCCGGGGCCGGGGCCGGGGCCGGGGCCGGGGCCGGGGCCGGGGCCGGGGCCG
GGGCCGGGGCCGGGGCCGGGGCCGGGGCCGGGGCCGGGGCCGGGGCCGGGGCCGGGGCCGGGGCCGGGGCCGGGGCCGGGGCCGGGGCCGGGGCCGGGGCCGGGGCCGG
GGCCGGGGCCGGGGCCGGGGCCGGGGCCGGGGCCGGGGCCGGGGCCGGGGCCGGGGCCGGGGCCGGGGCCGGGGCCGGGGCCGGGGCCGGGGCCGGGGCCGGGGCCGGG
GCCGGGGCCGGGGCCGGGGCCGGGGCCGGGGCCGGGGCCGGGGCCGGGGCCGGGGCCGGGGCCGGGGCCGGGGCCGGGGCCGGGGCCGGGGCCGGGGCCGGGGCCGGGG
CCGGGGCCGGGGCCGGGGCCGGGGCCGGGGCCGGGGCCGGGGCCGGGGCCGGGGCCGGGGCCGGGGCCGGGGCCGGGGCCGGGGCCGGGGCCGGGGCCGGGGCCGGGGC
CGGGGCCGGGGCCGGGGCCGGGGCCGGGGCCGGGGCCGGGGCCGGGGCCGGGGCCGGGGCCGGGGCCGGGGCCGGGGCCGGGGCCGGGGCCGGGGCCGGGGCCGGGGCC
GGGGCCGGGGCCGGGGCCGGGGCCGGGGCCGGGGCCGGGGCCGGGGCCGGGGCCGGGGCCGGGGCCGGGGCCGGGGCCGGGGCCGGGGCCGGGGCCGGGGCCGGGGCCG
GGGCCGGGGCCGGGGCCGGGGCCGGGGCCGGGGCCGGGGCCGGGGCCGGGGCCGGGGCCGGGGCCGGGGCCGGGGCCGGGGCCGGGGCCGGGGCCGGGGCCGGGGCCGG
GGCCGGGGCCGGGGCCGGGGCCGGGGCCGGGGCCGGGGCCGGGGCCGGGGCCGGGGCCGGGGCCGGGGCCGGGGCCGGGGCCGGGGCCGGGGCCGGGGCCGGGGCCGGG
GCCGGGGCCGGGGCCGGGGCCGGGGCCGGGGCCGGGGCCGGGGCCGGGGCCGGGGCCGGGGCCGGGGCCGGGGCCGGGGCCGGGGCCGGGGCCGGGGCCGGGGCCGGGG
CCGGGGCCGGGGCCGGGGCCGGGGCCGGGGCCGGGGCCGGGGCCGGGGCCGGGGCCGGGGCCGGGGCCGGGGCCGGGGCCGGGGCCGGGGCCGGGGCCGGGGCCGGGGC
CGGGGCCGGGGCCGGGGCCGGGGCCGGGGCCGGGGCCGGGGCCGGGGCCGGGGCCGGGGCCGGGGCCGGGGCCGGGGCCGGGGCCGGGGCCGGGGCCGGGGCCGGGGCC
GGGGCCGGGGCCGGGGCCGGGGCCGGGGCCGGGGCCGGGGCCGGGGCCGGGGCCGGGGCCGGGGCCGGGGCCGGGGCCGGGGCCGGGGCCGGGGCCGGGGCCGGGGCCG
GGGCCGGGGCCGGGGCCGGGGCCGGGGCCGGGGCCGGGGCCGGGGCCGGGGCCGGGGCCGGGGCCGGGGCCGGGGCCGGGGCCGGGGCCGGGGCCGGGGCCGGGGCCGG
GGCCGGGGCCGGGGCCGGGGCCGGGGCCGGGGCCGGGGCCGGGGCCGGGGCCGGGGCCGGGGCCGGGGCCGGGGCCGGGGCCGGGGCCGGGGCCGGGGCCGGGGCCGGG
GCCGGGGCCGGGGCCGGGGCCGGGGCCGGGGCCGGGGCCGGGGCCGGGGCCGGGGCCGGGGCCGGGGCCGGGGCCGGGGCCGGGGCCGGGGCCGGGGCCGGGGCCGGGG
CCGGGGCCGGGGCCGGGGCCGGGGCCGGGGCCGGGGCCGGGGCCGGGGCCGGGGCCGGGGCCGGGGCCGGGGCCGGGGCCGGGGCCGGGGCCGGGGCCGGGGCCGGGGC
CGGGGCCGGGGCCGGGGCCGGGGCCGGGGCCGGGGCCGGGGCCGGGGCCGGGGCCGGGGCCGGGGCCGGGGCCGGGGCCGGGGCCGGGGCCGGGGCCGGGGCCGGGGCC
GGGGCCGGGGCCGGGGCCGGGGCCGGGGCCGGGGCCGGGGCCGGGGCCGGGGCCGGGGCCGGGGCCGGGGCCGGGGCCGGGGCCGGGGCCGGGGCCGGGGCCGGGGCCG
GGGCCGGGGCCGGGGCCGGGGCCGGGGCCGGGGCCGGGGCCGGGGCCGGGGCCGGGGCCGGGGCCGGGGCCGGGGCCGGGGCCGGGGCCGGGGCCGGGGCCGGGGCCGG
GGCCGGGGCCGGGGCCGGGGCCGGGGCCGGGGCCGGGGCCGGGGCCGGGGCCGGGGCCGGGGCCGGGGCCGGGGCCGGGGCCGGGGCCGGGGCCGGGGCCGGGGCCGGG
GCCGGGGCCGGGGCCGGGGCCGGGGCCGGGGCCGGGGCCGGGGCCGGGGCCGGGGCCGGGGCCGGGGCCGGGGCGGGGCTGCGGTTGCGGTGCCTGCGCCCGCGGCGGC
GGAGGCGCAGGCGGTGGCGAGTGGGTGAGTGAGGAGGCGGCATCCTGGCGGGTGGCTGTTTGGGGTTCGGCTGCCGGGAAGAGGCGCGGGTAGAAGCGGGGGCTCTCCT
CAGAGCTCGACGCATTTTTACTTTCCCTCTCATTTCTCTGACCGAAGCTGGGTGTCGGGCTTTCGCCTCTAGCGACTGGTGGAATTGCCTGCATCCGGGCCCCGGGCTT
CCCGGCGGCGGCGGCGGCGGCGGCGCAGGGACAAGGGATGGGGATCTGGCCTCTTCCTTGCTTTCCCGCCCTCAGTACCCGAGCTGTCTCCTTCCCGGGGACCCGC
TGGGAGCGCTGCCGCTGCGGGCTCGAGAAAAGGGAGCCTCGGGTACTGAGAGGCCTCGCCTGGGGGAAGGCCGGAGGGTGGGCGGCGCGCGGCTTCTGCGGACCAAGTC
GGGGTTCGCTAGGAACCCGAGACGGTCCCTGCCGGCGAGGAGATCATGCGGGATGAGATGGGGGTGTGGAGACGCCTGCACAATTTCAGCCCAAGCTTCTAGAGAGTGG
TGATGACTTGCATATGAGGGCAGCAATGCAAGTCGGTGTGCTCCCCATTCTGTGGGACATGACCTGGTTGCTTCACAGCTCCGAGATGACACAGACTTGCTTAAAGGAA
GTGACTATTGTGACTTGGGCATCACTTGACTGATGGTAATCAGTTGTCTAAAGAAGTGCACAGATTACATGTCCGTGTGCTCATTGGGTCTATCTGGCCGCGTTGAACA
CCACCAGGCTTTGTATTCAGAAACAGGAGGGAGGTCCTGCACTTTCCCAGGAGGGGTGGCCCTTTCAGATGCAATCGAGATTGTTAGGCTCTGGGAGAGTAGTTGCCTG
GTTGTGGCAGTTGGTAAATTTCTATTCAAACAGTTGCCATGCACCAGTTGTTCACAACAAGGGTACGTAATCTGTCTGGCATTACTTCTACTTTTGTACAAAGGATCAA
AAAAAAAAAAGATACTGTTAAGATATGATTTTTCTCAGACTTTGGGAAACTTTTAACATAATCTGTGAATATCACAGAAACAAGACTATCATATAGGGGATATTAATAA
CCTGGAGTCAGAATACTTGAAATACGGTGTCATTTGACACGGGCATTGTTGTCACCACCTCTGCCAAGGCCTGCCACTTTAGGAAAACCCTGAATCAGTTGGAAACTGC
TACATGCTGATAGTACATCTGAAACAAGAACGAGAGTAATTACCACATTCCAGATTGTTCACTAAGCCAGCATTTACCTGCTCCAGGAAAAAATTACAAGCACCTTATG
AAGTTGATAAAATATTTTGTTTGGCTATGTTGGCACTCCACAATTTGCTTTCAGAGAAACAAAGTAAACCAAGGAGGACTTCTGTTTTTCAAGTCTGCCCTCGGGTTCT
ATTCTACGTTAATTAGATAGTTCCCAGGAGGACTAGGTTAGCCTACCTATTGTCTGAGAAACTTGGAACTGTGAGAAATGGCCAGATAGTGATATGAACTTCACCTTCC
AGTCTTCCCTGATGTTGAAGATTGAGAAAGTGTTGTGAACTTTCTGGTACTGTAAACAGTTCACTGTCCTTGAAGTGGTCCTGGGCAGCTCCTGTTGTGGAAAGTGGAC
GGTTTAGGATCCTGCTTCTCTTTGGGCTGGGAGAAAATAAACAGCATGGTTACAAGTATTGAGAGCCAGGTTGGAGAAGGTGGCTTACACCTGTAATGCCAGAGCTTTG
GGAGGCGGAGGCAAGAGGATCACTTGAAGCCAGGAGTTCAAGCTCAACCTGGGCAACGTAGACCCTGTCTCTACAAAAAATTAAAAACTTAGCCGGGCGTGGTGATGTG
CACCTGTAGTCCTAGCTACTTGGGAGGCTGAGGCAGGAGGGTCATTTGAGCCCAAGAGTTTGAAGTTACCGAGAGCTATGATCCTGCCAGTGCATTCCAGCCTGGATGA
CAAAACGAGACCCTGTCTCTAAAAAACAAGAAGTGAGGGCTTTATGATTGTAGAATTTTCACTACAATAGCAGTGGACCAACCACCTTTCTAAATACCAATCAGGGAAG
AGATGGTTGATTTTTTAACAGACGTTTAAAGAAAAAGCAAAACCTCAAACTTAGCACTCTACTAACAGTTTTAGCAGATGTTAATTAATGTAATCATGTCTGCATGTAT
GGGATTATTTCCAGAAAGTGTATTGGGAAACCTCTCATGAACCCTGTGAGCAAGCCACCGTCTCACTCAATTTGAATCTTGGCTTCCCTCAAAAGACTGGCTAATGTTT
GGTAACTCTCTGGAGTAGACAGCACTACATGTACGTAAGATAGGTACATAAACAACTATTGGTTTTGAGCTGATTTTTTTCAGCTGCATTTGCATGTATGGATTTTTCT
CACCAAAGACGATGACTTCAAGTATTAGTAAAATAATTGTACAGCTCTCCTGATTATACTTCTCTGTGACATTTCATTTCCCAGGCTATTTCTTTTGGTAGGATTTAAA
ACTAAGCAATTCAGTATGATCTTTGTCCTTCATTTTCTTTCTTATTCTTTTTGTTTGTTTGTTTGTTTGTTTTTTTCTTGAGGCAGAGTCTCTCTCTGTCGCCCAGGCT
GGAGTGCAGTGGCGCCATCTCAGCTCATTGCAACCTCTGCCACCTCCGGGTTCAAGAGATTCTCCTGCCTCAGCCTCCCGAGTAGCTGGGATTACAGGTGTCCACCACC
ACACCCGGCTAATTTTTTGTATTTTTAGTAGAGGTGGGGTTTCACCATGTTGGCCAGGCTGGTCTTGAGCTCCTGACCTCAGGTGATCCACCTGCCTCGGCCTACCAAA
GAGCTGGGATAACAGGTGTGACCCACCATGCCCGGCCCATTTTTTTTTTCTTATTCTGTTAGGAGTGAGAGTGTAACTAGCAGTATAATAGTTCAATTTTCACAACGTG
GTAAAAGTTTCCCTATAATTCAATCAGATTTTGCTCCAGGGTTCAGTTCTGTTTTAGGAAATACTTTTATTTTCAGTTTAATGATGAAATATTAGAGTTGTAATATTGC
CTTTATGATTATCCACCTTTTTAACCTAAAAGAATGAAAGAAAAATATGTTTGCAATATAATTTTATGGTTGTATGTTAACTTAATTCATTATGTTGGCCTCCAGTTTG
CTGTTGTTAGTTATGACAGCAGTAGTGTCATTACCATTTCAATTCAGATTACATTCCTATATTTGATCATTGTAAACTGACTGCTTACATTGTATTAAAAACAGTGGAT
ATTTTAAAGAAGCTGTACGGCTTATATCTAGTGCTGTCTCTTAAGACTATTAAATTGATACAACATATTTAAAAGTAAATATTACCTAAATGAATTTTTGAAATTACAA
ATACACGTGTTAAAACTGTCGTTGTGTTCAACCATTTCTGTACATACTTAGAGTTAACTGTTTTGCCAGGCTCTGTATGCCTACTCATAATATGATAAAAGCACTCATC
TAATGCTCTGTAAATAGAAGTCAGTGCTTTCCATCAGACTGAACTCTCTTGACAAGATGTGGATGAAATTCTTTAAGTAAAATTGTTTACTTTGTCATACATTTACAGA
TCAAATGTTAGCTCCCAAAGCAATCATATGGCAAAGATAGGTATATCATAGTTTGCCTATTAGCTGCTTTGTATTGCTATTATTATAAATAGACTTCACAGTTTTAGAC
TTGCTTAGGTGAAATTGCAATTCTTTTTACTTTCAGTCTTAGATAACAAGTCTTCAATTATAGTACAATCACACATTGCTTAGGAATGCATCATTAGGCGATTTTGTCA
TTATGCAAACATCATAGAGTGTACTTACACAAACCTAGATAGTATAGCCTTTATGTACCTAGGCCGTATGGTATAGTCTGTTGCTCCTAGGCCACAAACCTGTACAACT
GTTACTGTACTGAATACTATAGACAGTTGTAACACAGTGGTAAATATTTATCTAAATATATGCAAACAGAGAAAAGGTACAGTAAAAGTATGGTATAAAAGATAATGGT
ATACCTGTGTAGGCCACTTACCACGAATGGAGCTTGCAGGACTAGAAGTTGCTCTGGGTGAGTCAGTGAGTGAGTGGTGAATTAATGTGAAGGCCTAGAACACTGTACA
CCACTGTAGACTATAAACACAGTACGCTGAAGCTACACCAAATTTATCTTAACAGTTTTTCTTCAATAAAAAATTATAACTTTTTAACTTTGTAAACTTTTTAATTTTT
TAACTTTTAAAATACTTAGCTTGAAACACAAATACATTGTATAGCTATACAAAAATATTTTTTCTTTGTATCCTTATTCTAGAAGCTTTTTTCTATTTTCTATTTTAAA
TTTTTTTTTTTACTTGTTAGTCGTTTTTGTTAAAAACTAAAACACACACACTTTCACCTAGGCATAGACAGGATTAGGATCATCAGTATCACTCCCTTCCACCTCACTG
CCTTCCACCTCCACATCTTGTCCCACTGGAAGGTTTTTAGGGGCAATAACACACATGTAGCTGTCACCTATGATAACAGTGCTTTCTGTTGAATACCTCCTGAAGGACT
TGCCTGAGGCTGTTTTACATTTAACTTAAAAAAAAAAAAAGTAGAAGGAGTGCACTCTAAAATAACAATAAAAGGCATAGTATAGTGAATACATAAACCAGCAATGTAG
TAGTTTATTATCAAGTGTTGTACACTGTAATAATTGTATGTGCTATACTTTAAATAACTTGCAAAATAGTACTAAGACCTTATGATGGTTACAGTGTCACTAAGGCAAT
AGCATATTTTCAGGTCCATTGTAATCTAATGGGACTACCATCATATATGCAGTCTACCATTGACTGAAACGTTACATGGCACATAACTGTATTTGCAAGAATGATTTGT
TTTACATTAATATCACATAGGATGTACCTTTTTAGAGTGGTATGTTTATGTGGATTAAGATGTACAAGTTGAGCAAGGGGACCAAGAGCCCTGGGTTCTGTCTTGGATG
TGAGCGTTTATGTTCTTCTCCTCATGTCTGTTTTCTCATTAAATTCAAAGGCTTGAACGGGCCCTATTTAGCCCTTCTGTTTTCTACGTGTTCTAAATAACTAAAGCTT

TTAAATTCTAGCCATTTAGTGTAGAACTCTCTTTGCAGTGATGAAATGCTGTATTGGTTTCTTGGCTAGCATATTAAATATTTTTATCTTTGTCTTGATACTTCAATGT
CGTTTTAAACATCAGGATCGGGCTTCAGTATTCTCATAACCAGAGAGTTCACTGAGGATACAGGACTGTTTGCCCATTTTTTGTTATGGCTCCAGACTTGTGGTATTTC
CATGTCTTTTTTTTTTTTTTTTTTTTTGACCTTTTAGCGGCTTTAAAGTATTTCTGTTGTTAGGTGTTGTATTACTTTTCTAAGATTACTTAACAAAGCACCACAAACT
GAGTGGCTTTAAACAACAGCAATTTATTCTCTCACAATTCTAGAAGCTAGAAGTCCGAAATCAAAGTGTTGACAGGGGCATGATCTTCAAGAGAGAAGACTCTTTCCTT
GCCTCTTCCTGGCTTCTGGTGGTTACCAGCAATCCTGAGTGTTCCTTTCTTGCCTTGTAGTTTCAACAATCCAGTATCTGCCTTTTGTCTTCACATGGCTGTCTACCAT
TTGTCTCTGTGTCTCCAAATCTCTCTCCTTATAAACACAGCAGTTATTGGATTAGGCCCCACTCTAATCCAGTATGACCCCATTTTAACATGATTACACTTATTTCTAG
ATAAGGTCACATTCACGTACACCAAGGGTTAGGAATTGAACATATCTTTTTGGGGGACACAATTCAACCCACAAGTGTCAGTCTCTAGCTGAGCCTTTCCCTTCCTGTT
TTTCTCCTTTTTAGTTGCTATGGGTTAGGGGCCAAATCTCCAGTCATACTAGAATTGCACATGGACTGGATATTTGGGAATACTGCGGGTCTATTCTATGAGCTTTAGT
ATGTAACATTTAATATCAGTGTAAAGAAGCCCTTTTTTAAGTTATTTCTTTGAATTTCTAAATGTATGCCCTGAATATAAGTAACAAGTTACCATGTCTTGTAAAATGA
TCATATCAACAAACATTTAATGTGCACCTACTGTGCTAGTTGAATGTCTTTATCCTGATAGGAGATAACAGGATTCCACATCTTTGACTTAAGAGGACAAACCAAATAT
GTCTAAATCATTTGGGGTTTTGATGGATATCTTTAAATTGCTGAACCTAATCATTGGTTTCATATGTCATTGTTTAGATATCTCCGGAGCATTTGGATAATGTGACAGT
TGGAATGCAGTGATGTCGACTATCTGCCCCCCACCATCTCCTGCTGTTGCCAAGACAGAGATTGCTTTAAGTGGTGAATCACCCTTGTTGGCGGCTACCTTTGCTTACT
GGGATAATATTCTTGGTCCTAGAGTAAGGCATATTTGGGCTCCAAAGACAGACCAAGTGCTTCTCAGTGATGGAGAAATAACTTTTCTTGCCAACCACACTCTAAATGG
AGAAATTCTTCGAAATGCAGAGAGTGGGGCTATAGATGTAAAATTTTTTGTCTTATCTGAAAAAGGGGTAATTATTGTTTCATTAATCTTCGACGGAAACTGGAATGGA
GATCGGAGCACTTATGGACTATCAATTATACTGCCGCAGACAGAGCTGAGCTTCTACCTCCCACTTCACAGAGTGTGTGTTGACAGGCTAACACACATTATTCGAAAAG
GAAGAATATGGATGCATAAGGTAAGGGGCTTTTGAGCTTGATCATGGTAGCCTGGCCAATGAAAGTTTTTTTCTGGTACAGTTACACTTAAGTTTTGGAAATTATATGC
TGCTAACACCAGACAGCTGTTATGTTGTGTCTCCTGGGCACAGAAAGCCCTGCTCTCATGCCTGGGGTCTTCACAGTCCTAATGGAAAGTAAGATCTTATAAACATTGT
GTCTGAGTTTGTTCTGGAAGCTGTGACTCTACCTTCTTGTTTTCCTTTCCCTGTGTGACTTTGTCCTTTGCTTACAACAGTGCAAAAGTATAAATATTCTCAGATTTTG
ATAAGCTGTCAGCCACACAGCCTTAGTAACTAAGCTGCTGTCCCACGCTCCCAGTTCTGTATAACGAGGATGGACCAATTAGATTCTAAGGAGTTATTCCTTTCAATTT
GCAAATTTAGCTAAAGGAAATATTGTTTTCTCCTGATATTTACATTGCTTTTCATTTTCAGCATATCTAAAGAACAAACCTAATTCTCCTTCCTACTTTCTAGTTTAAT
ATAATCCTAAAAATCCATTAAAACATGACTAATTCTATAAGGCCTCTAACCTACAAAGGGAAGTAGCATTTTGAAAAGAATAGTTTTCTCTATTATACCTATTCATGCA
GACTTCCTTCCTTATTTCTGACATACTTAACAAAAATCATTTAGATTCAAACAGTTTAGCTGCAGGTGATATTACAGACAAGTAATCCCAGTGCTCTATCTAGTCTGAG
GCAAAAGGATTTGAGCTCAGTGCCAGCCTGTTCTATCTACCTGGTGAGTTCCAGTCCCATAAATAAACAAACTAAAACAACCGTTCCTCTGTTCCTCAGATGCGAGTCG
ATCTTGTTTGATTTAAATAGTGTGTAATTATTTTCTTTTGAAGCTGCAGGTGTTATGTGGGCTGTTTTAGACTAAATTCTCTCTTTACTGTGGAGTAAAGGGTGCTGTG
ATTGTATTTCATGTTCTCTGCGAGAGCTTGAACTTGTTGGGCTAATCGCTTGTCTCCATCCTGTCTCCCCACCTGCGTAAAAAGTATTTTCCTGTGAGCTGTACATGAT
AGAGCATATCTACATTGAAAAATGAACGAGCATCAAAATGGATTTGTTAAAGTAAATTTTCTTTTTCTTAGGAAAGACAAGAAAATGTCCAGAAAATTGTCTTGGAAGG
CACAGAGAGGATGGAAGATCAGGTACAGTGCATATCACATGCTGCCTGTGGCAGGTCCTCTTTGCTTATGTCGGTATAAAGTTGGTGGGTACTTCTGGTAAGGACCTGA
GGATACATTCATTTGACGGAAGGAGCCTGAAAATGAGTATTCTTGTTAAGCTGTATAGAATGAACTGAATAAAAATTTCTGCAGCCTAAGTTTGAATTTTAAAAAAATT
TAATTACATCTACAAATTAGTATTTGGCCACCCTTTTTCAATCAGCAAGAATATGTTTGAGGTCATTTATTTGTAGTAAAATTGCATGCAGTTTATTTATTTTATTGAA
AATAGGTTTTTTAAACTATATTTTCTGATTATGGTTTTCCCTCCTCTGAATCCTCCTAGAACCTCCACCTACCCAAATCTATATCTGTTCTTTCTCTCTCTCATTAGGA
TACAATCAGGCATGTAAAATAATAGTAGTAGTAGTAGTAATAATAATGTAAAATAAGTTAAAGTAAAAACAAACCAGAGTAGGACAACATAAATAGAAGTAGAAAAGAG
CCAAATAAGAAATTCAAGAAACACATATAGACACAGACACAATATTTGCATACACAGAAATTGCATAAAACCGCAAGACTGGAAACCATAATATGTATGTAAGGTGGAG
TGGGAAGCCCTGACAGCACAGTGAGTAAAGCACTTTCAAAAACACCACTGACTTTGTGTTGTGTTGCCTGTCTGCTGGGCATGAGGCCTGGCCTTAGAGAGTGGTGTGT
ATACCCAGGAAGACTTACATAAACACTTAGCTTTTCATTTGTGACCTGATAGCAATTGGAAATAGTGTCTGGGCTAGGCATTCCGGCTTATTGCCACTTCCCCTCAGCA
CTGAGGCCCCATCTGAATCGGATCCGTGCAACCCTTGTGCATATGCAGTTTTAAAAGTTATCCCTTCTGCAACTATGCTCACAGGAGTTGCCGTCTTAAGGGAGTGAGC
ACACCCCTGAGGCATGGCTCCAGGGGTGCAGAGCCAGCCATAGGCACAGTTTTTTTTAAAAGGTTTATGTTGTAGTTTTGAAACTCAAATTTATGTGTATTTGTGGCAG
ATTGTTTGAATGTTGAAATTTGCCAGTAACATCTTTTATCTTCTTCCCTTTAGCCTGGCATGCCACCCACCCTCATTTGTCCTTGTCAAACTCCAGTAATTAAACATGG
CTATGTGGCCTTTTCTCTCATTTTCCTTAGCATGGCTAAGGAGAATGGGACTTAAAAAATAATAATCATCATTTTAAGTATGTCTGAGGGTTTGAGGATATAGTGGTAG
AATATCTGCCTAGCTTCCATAGCTTGATCCTACATTTGATCCCTGGCAAAACACACACACACACACATATACACACACATAAAATGACTTTTATAAAGTTAGTGTGCTG
TGCTGTGATGAACAGTGCCATAGGAAATATTCTTGGAAAAGACCTGAAACTAAATGCTCTAAAAGGTCTAATCTTTACTTGCTTGCTGATCGTTAAGCAGAGTCTCCAA
GTATAAAGTCACTTTCACCAACCTCTGCACTGGATTTCTGGAGTAATTAGGGAGAGTCATTTCAATATAAGAAAATTTAGTACCAAATAAAATTTTCATTCAGTGAAAT
TTTGTTTTTGAAAGTAAGAGCCCACTGTGGTGGTTTGAATATGCTTGGCCCAGGGAGTGTCCTGTAAGATTTTTGTTGTTGTTGAACTCCATTGAGACTTATGTTGACA
ATAAATGCCTGAGAGTCCATGTCTAAAATGCTGTACCTGTCTGAACCCAACGGAGATAAAACTTACCATTTCTGAAAAGGATGAGGTGTTTTATTTACATAGCTGATGT
AATGTGCTTGCAACAGCTCTATTATGAATCTTAATACTACTTCAGTATATCACAGCACTTCAGGAAATTTAACATACATTGTTTAATTCCATGTCTTAATTGTATTTGT
AAACAGACATTTCAGCAGTTACTCTAAAAAGTAGAAATAATGAGTGGTTGCTTCTGGTCATTAGGATGAAATATTGAAATGATAAAATTTTCTGGGCTGGAGAGATGGC
TCAGAGGTTAAGAGCACTGACTGCTCTTCCAGAGATCCTGAGTTCAATTCCCAGCAACCACATGGTAGCTCACAACCATCTGTAATGGGGATCTGATGCCCTCTTCTGG
TGTGTCTGAAGACAACTACAGTGAACTCATACAAATAAAAATAAATAAATCTTTTTTTAAAAATCTATATCTGCATAGGCATTTCTAGATTAGGATAAATTTTCCAAAG
GAAATAAGCACCTCCATGATAAGGGCATTGGAAATGAAGCCCCCGCCCCCACCCCCGGTCTGCACGTGTGTTGAGGATGAGATCTAGGGCCTCCTTATACATGCCAGGC
AGCTGTTCTGTCACCAAGTGGAATATAATCCTCAACCCTTAATTTGAGGTTCTAACTTTAAAATAGATGTGAGGGGTTTAAATAATCATTTCATGAAACTTAAATGAGC
AAGTTTATTACTGAGGTGAGTATAAGTAATTGATAATTTTAAATATATTTAGCTGAGATTGATAGACACTTGGCAATGTCAGCATCTTATTTAGGTGATCATAAACTGA
TGGGAGAAATGGTAAATGTTAGGGGGTGTCGCTCATGTCACACACCGCAGTTATGCTGCAAACAAGATGCCGGGAAATAGAAATTCAAGGTCTTGTTTTGCGGGTGCAG
ACTCTTCTGTCTCACTGATTCTATGTGGTAACTTCAGTATGCATTTGGATAGATTATGTCCCATTTTGAATGTGGAAGCTGGCTGTTGAGAGGAGACTTCCTGGTGAAT
TCCTTTTTCTAAGCATTACCATCTGTCTTAGTCAGGGTTTCTATTCCTGCACAAACATTATGACCAAGAAGCACTTGGGGAGGAAAGGGTTTATTCAGCTTACACTTCC
ACACTGCTGTTCATCACCAAGGAAGTCAGGACTGGAACTTAAGCAGGTCAGGAAGCAGGAGCTGATGCAGAGGCCACGGAGGGATGTTCTTTACTGGCTTGCTTCCCTG
GCTTGCTCAGCCTGCTGTCTTATAGAACCCAAGACTACTAGCCTAGGGATGGCACCACCCACAATGGGCCCTCCCCCCTTGATCACTAATTGAGAAAATGCCCCACAGC
TGGATCTCATGGAGGCATTTCCTCAACTGAAACTCCTTTCTCTGTGATAACTCCAGCCTGTGTCAAGTTGACACACAAAACCAGCCAGTACAACATCTTTTCACATTTA
ATTTTTCTCACTTTAAACGTGGCCTTTAACAAGCGCTTATAAAAATGCTTAAGCTTAAATGTTATTTAAGCTTAATATACTTAATATACAGCACTGTAGCTTAAATGTT
GCATGTGAGAGTATATGATAAGCCATGCTCACCAAGGAAAAGAAGCTTAAAGAGCATAAAAACCCTGACAGCGGTTTCTGAGTGGGAGGCTCGGGGACTGTGCTGAGCA
ATTCCAACCAAGGGTGTTTTACTCTCTGCCTCCATTTGAAATGTTTTTCCTGCACAACCTACCCACCCTGTGATTTCGTTCACTCGATTATGTTTGATCTAGGGTCAGA
GTATCATTCCCATGCTTACTGGGGAAGTCATTCCTGTAATGGAGCTGCTTGCATCTATGAAATCCCACAGTGTTCCTGAAGACATTGATGTAAGTGTCATGTATCTTTT
ATGGGTTCCCTTGAGTGGTGAGTGGGTGGATGTGTGGTGCATGTGCGTGTGTGTGCTTGCATACTGGGAATTGAACCCAAGTCCTCAGGAAGAGCAGCCGGTGCTCTTA
AGCACTGAGCCATCTCTTCAGAACCTCTTCCACCAGTTTCTTTGACCATTTGTTGAGAATATTCCAGTCACACATTTTCCGTGAGTAAATCTCTCTAATGCTGATTTGT
CATTAAGCTCAGTCTCCTAATTCTGATAGCTAAGAAGGGTAAATTATTAAAAAGTGCCCTTTACTCTTCCTGGCCAATTCCCCTTTGTTCTTCTGAAAAGTGCATAGAC
AGCATCACTTTATAGATCACCTTGATGCTCGTGAGAGGGCTGGCTCGTGCTGGCTCTAGACTTCGGCACACTTATTAAGAGTTCTCCCAACACTGTAAACAGACTAATT
TTTATATTGTGCATTTTAGATAGCTGATACAGTGCTCAATGATGATGACATTGGTGACAGCTGTCACGAAGGCTTTCTTCTCAAGTAAGAATTTTACTTCTTTTTCTGA
ATGCTAAGTAAAGCAGATTAAAAATCTTAATGCTCACCCATGACAAGATTTACAGGGAAAAGATGGTAGAAAACCTACTTCCTCCAATTATTTAGGGTCAACATGGCAC
ATTTGAGCTTACACGTGTTGTTCTCACCCATACAACAGTGGCATATCTGACATTACTCTTCCCACAGTCTAAAAAGGCAGAGTTTCCGTAGTACCCAGGGAAGTTCTGG
TCTGTGTTTGGGTCTGGTTTCTTCTTTCAATTCTCACTAAGTATAACCCTTAGGAATCTATCAAGTTGAGTTGCATTTTAAATTCCTGTGAATTCTTCAGGTCTAGAAA
TGGAAATCATTCATATTTTAGACTGACATTTTTCATCTTCTTGTGTAATTTAACATTTAAGAACTTGAGCTCTAATATCAGACTGTCTAGGTTACAACTGGGAAAACTT
GGTGAAGCTACCCAAAGCTGAACCTCCATTTTCTTACCTGTGAAATGTGAACAGTGATAACAGCTAGTTTCTTGGGTCCTTGTAGGCACCAAATGACAGGATAATATAA
AGCACCTAGGACAGTGGAGCCAATGAGCCAGGAGCCAGTGTGCCCTATTATATCTGCTCTAAGAAAGACAGTAAGTGGAATAGCCAATACTGACTGTCTTAGTCAGGCT
TTCTATTCCTGAACAAAAAACATCATGACCAAGAAGCAAGCTGGGGAGGAAAGGGTTTATTCAGCTTACACTTCCACGTTGCTGTTCCTCACCAAAGGAAGTCAGGACT
GGAACTCAGATCAGGAAACAGGAGCAGATGCAGAGGCCATGGAGGAATGTTACTTACTAGCTTGCTTTCTTATAGACCCCAAGACTACCAGCCCAGAGATGGTCCCACC
CACAAGGGACCCTGCCCCCTTGATCACTAATTGAGAAAATGCCCCACAGCTGGATCTCATGGAGGCATTTCCCCAACTGAAACTCCTTTCTCTATGATAACTCCAGCCT
GTTTCAAGTTGACACAAAACCAGCCAGTACGCTGACCGAGCAGCTGTGTGTTCCTCTGCAGGGCTGTGTTCTCTGTTTGTCCCTCATCTCCTGTTGTAGTCTCCTTTAC
AGTTACAGACTGTCATCAGTAACGAGAGAGAAGTGAATAGGATTTTGTTAAAGTGTTTACTTCTATGTCACATTCCCTTCCTATAATAAGCTCACAGTGAAATACCAGG
TGACCGTGCTTAACGGCATCTATTACCTAACTGGGGTATCTTTTTCCTTAAAATGGATTTAATTTTATGTGTGTTTGAATACCTGCATATGTGTATGTACACCATATTT
ATGTATGCCTGGTACCTGAAAAAGGGAAAAGAGGGCTTTGGCTTTCTTGAACTAGATGGTTGTGAGTCTCCATGTGGGTTCTGGATTGTCTCTGCAAGAGCGGCAGGC

ACACTTTAGCAGTGAGCCGCTCCTGTCCCGAGTTGTCTTAAGACCTGTGAAAGGTCCCTAAAAAATGCAGGGTTTTACCCGAATAAAAGATGACATCATGCAGATGGCT
TTGGTGTTCATCAAGCTCTTGTGTGTTGTCCTAACCTTGCTGGGCTTTGTCGTTGTGAAGCTGTAACTCCGTCAATGTTTTCCTTACCTACAGTGCCATCAGCTCACAC
CTGCAGACCTGTGGCTGTTCCGTTGTAGTTGGCAGCAGTGCAGAGAAAGTAAATAAGGTAATTCGTTCTACAGTTGAACATGATCTGACTTTTATCATCACTAGCATAT
CATACATTATCATCTAAACAGTAGGCTGCAATTGAAATAACCCCATAGTATAAGGAAGCAATGTAATTTTACCAAATTTCTCTGACACCCTCTAGCAGAACTGACTCTA
ATAGAATGAGTAAGAATTCAATTACCAAATTAATTTTGATACTCTTTTTTATTTTTGTTATTACTTTTTTATTTTATTTTAATTAGGTATTTTCTTCATTTACATTTCC
AATGCTATCCCAAAAGTTTCCCATACCCTCCCACCCACTCCCACTCCCCTATCCACCCACTCCCCTTTGGCCTTGGCGTTCACCTGTACTGAGACATATAAAATTTGCA
AGACCAATGGGCCTCTCTTTCCAATGATGGCCAACTAGACCATCTTCTGATACATATGCAGCTAGAGACACGAGCTCCAGGGGGTACTGGTTAGTTCATATTGTTGTTC
CACCTAAAGGGTTGCAGACCCCTTTAGCTCCTTAGGTACTTTCTCTAGCTCCTCCATTGGGGGCCCTGTGATCCATCCAATAGCTGACTGTGAGCATCCACTTCTCTGT
TTGCTAGGCCCCAGCATAGCCTCACAAGAGACAGCTATATCAGGGTCCTTTTAGCAAAATCTTGCTAGTGTGTGCAATGGTGTCAGCGTTTGGAAGCTGATTATGAGAT
GGATCCCCAGGATGGCAGTATCTAGATCGTCCATCCTTTCGTCTCAGTTCCAAACTTTGTCTCTGTAACTCCTTCCATGGGTGTTTTGTTCCCAATTCTAAGAAGGGAC
AAAGTGTCCACACTTTGGTTTTCATTCTTCTTGAATTTCATGTGTTTTGCAAATTGTATCTTATATCTTGGGTATCCTAAGTTTCTGGGCTAATATCCACTTATCAGTG
AGTACATATTGTGTGAGTTCCTTTGTGATTGGGTTACCTCACTCAGGATGATGCCCTCCAAGTCCATCCATTTGCCTAGGAATTTCATAAATTCATTCTTTTTAATAGC
TGAGTAGTACTCCATTGTATAAATGTACCACATTTTCTGTATCCATTCCTCTGTTGAAGGACATCTGGGTTCTTTCCAGCTTCTGGCTATTATAAATAAGGCTGCTATG
AACATAGTGGAGCATGTGACCTTCTTACCGGgccattgggccctagacaagttgtcgcggccgcactagtgactccagcgtaactggactggccacagttaggccgcaa
atgtaatcacactggctcaccttcgggtgggcctttctgcgtctagaggatcccctgttatgcccttccttacccatgcgctgagaggtgaaatcactctttaatttgt
acgcggtacggacctctatatcgctttgtcgtagaggtggaatctccccagcttgcagccattgatagactgcgccgggtgtcacgcccacgccaaccgctgcttttc
cacatcgccgaaatggcgcacaagttcttcgggtttcatggggattattataatcaataagtgaaaattaaagctaggtataatttataaaaattatagccagctataa
agagatcatttatgattaagggtatgaaaacacgaggcgaacgactgaaagcacgccgtttagagctgaaactgacgctgaagcaagtggcagaagctgtaggaatctc
tcttccgggcgtccaaaacttagaacgtggcgacgtgatgccgtcgctggagatcgggctatcgctggcaaaatgcctgcgtaagcccgtgcaatggatactgtatggc
actgaatctgatccagaccgcgttcctgttattggtacgacagagaccggaccggatagagactggcagcctggagaacctgccaacacagagcgattcctgccgttcg
tgagtcaacggaacaccgtttacgcactgacggtcgggaaccagattcagcgaaactaccagccgggagacgttatcctggtcgattcttcacttacgcttgtaccggg
tgaggatgtactggtttgtgataataacggcgagatcacaattcaacgattagcccgttatgacgagcagcattactaccttgatagtgctaactctcaacgggttatc
catgataaaagtgatcttcaatttgtgcaccaagtagtcggtacgatcaaatcgttcatggttgagggtagatgatacaataacagggtttattgctgactataattct
ggtttaatccgatctatactttgtcggggttgaaccagaccgtagcagccgaaaaaagacgaaaaaaaacccgagtcggcaaactcgggcctttttcaggaagtagcca
cgataacgcagatacgtccttcagaaagatagtgcgtttattgtggctgctcctgcggatttttcaacccgaaaaaatgctaattcgcatggaaaggctaaaaaatga
ccttacaagaattctacgcggagcgctttggcagcgatccgttttcactgcttgacgcggcacgtgatgagctgactgagctggcgaagatggccggtatcaactggcc
tgcgtgtgctgataagattcagctcaatccgcgcggtggtgtagagcgctataccacctacaataattcctccccggaagctctgcagaagagccttaaaggccgtgtg
gaaatctattcacgccttgagcagagcaaggatggcatcagttatcctttcgttaacttcgttgaaaaagcgcatgacgctggctcctggagtggcttttcattcctgt
tttcggagtatcgccgtgaacagcaacggaatggtgcgaccgtggtcgctcagccggaagaagaacgtgcacgcatggagcgccaggctgaagcacgccgcctccgagc
tgaacaacaacgcgttaatgatttaaaaaacaatcagatggagcaggaacgcctgcttggatggttggctttccatcgcgcctgggaacattcgccagctgaagatggc
tcctggccttatgcggtaaaaaagggaattcgtgacgtatttggcgcttgtgatatacgtcgcgtgaccagtcacgacagtgcgaaatggagccgtggaccaacaactt
acatggcgatacctctgtcccacctggacggaagaaaagacgggcggattgttggctggcaacgtatcgatctgaatggtggcaagttccagaccagcgccatcactaa
cggtgatttcgtcggtgcatgctttgttattggcgacctgcaggggcgcaaaaaattgcgactgcagagggtttcgccaccggggcatccatctggctggcgaccaga
aacgacccgaaaaaacgctttgacgcggtagtcattgcggtttccgctaacaacatgattcacgttgtcgagcagctggtgaacatgtacccggctgcgcaaatcactt
gcgctctcgataacgaccgcaaatcctcagctgaaggaaaaggcaacacaggcctccgcaccggattcgacatcatggagaagttttccggcgtcaaatgtgtttaccc
aacttttgaggatgaccctgagctggaatgcagcgattttaacgacctgcacagcctgagagggctgaaggaagtcgctcgccagttaacgagaaatcatctgagccgt
gcaaccgatctgttgtcgatcacgctgaataagctgcgtactctcccgcgtctgaacagacgcactttgccaaagaactgcttcgcgccgttgatattggcatgctga
catgcccggtaccaaacagcccgaaagaacttatgcgccttttcagcagcacgctgcgggatatgggtatcgcagaaatttataacggtaccgttaaagatcacattac
gcgccggttgaatcgcaaatgccgtgccgcgcaaacatcacgttcgttcagtgaacgcatcaccaacccgaaccttcgcccgtcacacatcacttacaaacggtttgaa
acctccaggatgaccgatgaggtgatgacatacgccgcacagctgcagggcatcgttattgtccgcgccgggatggggtcggtaaatcgacaggcctcctgcgtccac
tgatgctgcagtccacgcgtggcgtttccgtcgcgcaccgcgtatccctgataggcggcctgcatgaaatgatgaccgaagggaaaggcgcgaaagccgacattctgca
ttatcaggatcccggctatcaggaaatggcgccatatgccaacaagctgactatttgcatcaactccatacttaaaggctgctggcaaccgctgatgcgccagcatgac
ttcttcggcttcgatgaagcaacacagggcctgcgtgccattctggccgggcgtgcgatggaaaacccggtaggcgtattcaacacgcttatcgacgcgctggcgcgta
ctgaagagcatgccattatggtggatgccgatgccaacgatctgcttgttgacctggctgaactggcgatgaagcgacgcgaggagctgggcctacctgcctggctgca
aattcacgtgattgaactcccggtcgacgttcgcaaccgcgaaaccaacaagcctatccgcgtattttataccgagaaaaatcggatcatgaccgaggttatcgctgca
gtgcaacgcggcgaacgaatcatgctggccaccgacagttcgacgttcgccgaagacgttaccatgcagctgagactacagttccctgacaaaaagtttctctgcgtta
accagaaaaacaaacaggagaaggaagtcgacgatttcaccaaccagcctaaagtgatggtgaaaaaatatgacggcctcatctacagcccgtcgatatcttcaggtgt
atcgattgaggagaaacacttccaccgccatttcggcatgttctgcggcgaagtggtccccagcgacgccatccagatgctgcgccgcgaccgtacagctcaggaatac
attatcggtttcgacaagcttcgcggtaaacgtgaaaccgatccggaaaaatcaaacgcgcctacgcgcaggccttgctcgaaacggctggccactccgggctgctga
cagacgttgtctttgacggtgaccgaatttcactcggcgtggctaactcttcattcatgcagctcaaaattaaagccgccgcgctcgaggcatcggccagaaatgatta
tgccagcaacatgatttgcatcatgcatgatgatggctatcaggtcgcgcctatggccaccgacgcgcttgcaaacagcatcggtaaggatttacgtaaagaagcccgt
gaactggtctttgagcagcttatggagcgccacctgagtgtcgatactcctgaccaggccgaacacgatgagttgataaaaaaacgcaccctgtccctggatgagcagg
cccagctggtccgctgggacatcgaaaaggagctgcagctggatgtcgacgaggtggcgttaaaattctatttcgacggcggcctgaaaaaagtgcgcctgtttgaaac
catgcagctggatgagataaccgcccgtcgccttgaccgtgaagaagcgttgattcactttacctacgcttatcgcgttgccggaagatggcagcagttcgtcactacg
gccatgacgcgggaacaggccgacgcagagttccaggctaaattcccggccataaccgattaccgagtgaaatcgacccctgctgttgagataggcatgcgtggtttct
acaccctcaaatcggcgacgctgcagcagtacttccgcgactgcggtatcgatccaaaaaccctggaaggtgaagcagacatggacgccctcaaacgcgccagagataa
cctgctcacgccggaacggcgcgatctgctaaacaacgttttacgcatcggcggcttcaacacagagaaaggcaagaagaaagcgcctgcagacctctgcataggcatc
ctggagtctatggggctatccagtaagaccagacgcgccagagacggcgatgcacgccccacgatgcgttccattgacccggactcggtcgagttcctcatgaatatcg
tggagaagcgccgcgaggccggtttatcaattcacgcacgtaaggtggaaaaaaccaccatcgaagtggatcgcgatttggatctaaatatagatatacatggtaaccc
tcgatccaaaacagagcacgttccggacgccccacaatcagtaattatccaggcactggaggctatcccggtggcggtaccggaggcgtggcggagaacgcgttgcca
gcaaccgaaatggaggcggtacgcctgtggccagtggccagcatcgcgagaacgttcgcctcgctgtacatgaccgaatttatggacctgctatcagtacgcgaaataa
ggctgctgaaagcgtttctaagccagcggcaagccgtggcgatataaccggaggggtaatgggaaaacaacggaatgccttgtcatcgtggaggctacgacccttat
gtggtcaaggtcagccacgacgcgacaagctatagcgattataaaactggccgggaacttccattcacgatcattcggatgaacgggttctcagatagtgagggagttt
attgctgcattgccagcgataagagcatggaagaccacacagacgaagccagggcgcttctaaaagcctataaggtgaagccactatgacgtcgacaccagagttcctg
aagcaactggattacgaacagctgaagtactgtcgcgagctgtgcgataaccgcatccgggctattgaggccgaagagaaaaagatggcctgggcagttactgatggtg
gcattaactacggctggttccgtacagaagactacatgaaggcggtagaatgcctggttgctacagctgcagaacggtgggaagaatcggataaggctgacccatctgg
acgtggttggcttgacctttcaatccgtggaagccgcctgccggtgtccgagtatgaagctttatttgctgatggccagtggggggtgaatagtggcgcatgatctacaa
aaaatgatggatggtgagctggtccgcttacaacttgagattaagaaagaacaggagaggcgtttagccgaaaagatgattcctgtttttggtgtgtggcaatcaccta
ttaatcaatatgaatttgctgatccagagcaggcaatttcttgtgccgcagggctattggataaggttcttcgtgaggtcaaggacgatctttccaaagaaggtggatt
acagggatggaacggcaatttgctacgcatctacgtacagcatgtgaacgagagcgattttgcagttctacagccttatttaaaggacaaataatcatgatttatcgca
gaggatgggtgcctgttttgtggaggaatgagctggaaaagcggctcaaagaagaagggttcgaaaactggaagcagataagtaactttctttgtggtgaagaggattg
ctactcggaagcagcaaaaacgccaggacaatacgcttaccaggtcgtcgacaataccgaatggatggaacgtcgtgatgcgacgttctggcagcgtctgaaccgtttg
tggttcgttccgctgtacctgctgactattccgttccagtggctcattcgtggccgtatggggtttgagacaacatctaaggtggctgcggtcatcaataagattaccg
gactccaataactgatcaggttctgatcagtcagtaaagcaaactataaaaacacctgtgcacgtgactggtcacggggtaaaatatccacctaaattattgacgtgcg
ttctgctttggcggtagagttaccccgctgcagcaaaatctgcagccgggcctctcaaccccgaatgacaagaagcgcacaacacgcgccagcgtgtttttttgtgtgt
taaatctgcgcatacctgaattatggtggctcagatggggccaacttcggttgggccggtttcttcttgtaccggtgttgagaaccctgtctgggctaccacccctata
gagattctcaactctggtggtagcaccctatacaagataggaatgcataccatgttcaaattcaaatttgcagctgtcgtccgcacggacaaaaaatcccatatccacc
gcctttccaccattgcatcatccgagcgagaagctcgtcgccagttcgccagccgttttgtcctcgttctgtcagcccgtatcccggtcagcgaggtgatcgcatgaac

cagattcagttaaacgcccagggcctgctggagacgattgaggaacgcctgctgcaggttgaagcgcttgtgggctctgctcaccgcactatctccagttatgaggcca
cgctgtatttgcaggaagctgcggaacttctgcaggtcgcgcgcgaactcacgcaggaagctcgcggctgctccctgtccctttctgagcagctgaaatccggggaggc
gcaatgaaggctctctccgtcttttcatttcaggaaagccaccctatacgggtggttcttgtcggtggtgacccgtggtttgtggcgctggatatctgtgctgcattga
atatagccaacccttctgatgcattgcgtaagctggatcatgatgaaaaattgaccctcggtttaaccgaggcacaaaaacttgatcggatggccagggaagtaaatgt
ggtatcggaatctggtctctacaccataatcctccgctgccgcgacgcggtgaaacaggggacgacagcctggcggttccgcaagtgggtgacgaacgaggttctgcca
gctatccggaaaaacggcgagtatgcttttgtcgagcctgagcctaagaacgccggtgaaccactggactggcggcagaaagaagaacttcggggcctgataaacgaca
tagcacaaagttttcagtaccgtaacgcatgggttagcggcgtctggatggctctccgccgcgcctgcaggaacccatcaccgaacccaataaccgtggatgatctccc
ggcaatcatcgccgagctgcgccggatattaacggctgcagaaacggctctgggcaacatgcgtgtctacgaacgggagctgctgcgcgatgtagttcgtggcggtcgc
cggagtatgtcttgtggagagctgccgatcaccgatattgatacggaactggagaaggtgctgccagcacattttgagctggccattgaaaagctggaaacgctatcca
cgaaactaacatcgccagcgattgcagaaaaatagggttggccagcccgaatttaagccccgctttacgcggggaagtccctgcagtcgggacggcgatgtccgtagtt
gcagctgcagtttccgcctaatccttccgtcccttgcaggggatgggtagggacaggcctcaataaaaaagccaccgattaacaccggtggcttttcttttagctgtag
tacgtttcccatgcggatcgcattgcggccatagggctatcggcggggccggaccaggcataatgctttccatcgtattcaaactctatcttgtacgttccgtccccgt
tattttttgcaggcttgaagacgggttttagagcagttggctgatgttcttctggcccttcctcttcggttggttcatcgccgccaccctcgtcgagctcaatttcatc
ttcatccagctcgtcgtcttgggactcatcatccggttcgtcgatggtctcaacggattcttcatcaggcaggacgattgcaggtgtctctatcttcagctgccactgc
ccgttctcgccaacgaactgccccaatgcatcagcggcaaactccaggtaccggctaatcatcgtcgggctaaatttaaaggcccggagagtgctgttggttattttttg
ctgatgggtcctgctccaccagctgcttaacggtttcatggagacggacgccagcgtcacctctggcaaagcctggcatttcatcgtccagtttctgcagagccaccag
cctggtgttttcatccccaacttcaggtcgccaggttctggagaagttggccagcttgaactgcttatagtgcagctgggtgttctcatcgtcgtgtccgagaatctcc
atgaagaacacatcctcgtcgacgtttttccaccgtggatcgacgcggaagaacatctcataagcgatgcgagcgtaaatagcgcggctatctttataaacacgacggt
catcgccgaaaaatgatttaacccaagggttaaatgcttttgctaaaatagcatttatcctgccgttctcagaccttgtatcatcctttccatatcctttaacaacctc
atcgaaatcagatgcagcagagcaagaacgcaattctgttaataattcaacgaataattttgcttcgcataaagtataaatcgttctggttacgcttttatcttcagag
cgtttttttagcttgccctgagaaattaaccgtatactttcctgaaacggcaaattcaccctgaaacattatctcaatcattcttcgccctgataccgcagccagagcaa
aggccaaaggtgccattccagaacgagtgtttaaactaaataaagtcgcaggattattcaaaatatcatagatagactgcatgtatgttgggtagtcaataaccacaac
attacgcttcttctcgcgcagaacatcggcccatcgttgctgtatagatgtacgctccgcagggcttagctgcagatggtacagaacctcatggttgaccttgagctgg
tgtagttcttctaacaacgcagagccttgttggaataacttataaagatagtcacggcgctccttccaatcatcactgtttaatcactaagagcaaaactccaatctg
gatatttttttattagtcttgctatttttgcatcactgcctttagagcctattcttacatttgataactcttcggcaagcggcattatttctttcaatttagattgtaa
cgatgacatgtgctggcgaatattagccgtaggcatagaaagccatgaagataattcttcgctgtaaagaggatacttttccgataatttattaatatttttatcaaag
ctatgatgtaatttatcatcaaaccgctttcttgccctgctcatataggcgttaaaagtattcgcggttattcttttctgcaatcctttcccacggaactttcttttat
cattaaataacgcgttcttataccgtgcggctgcggctttaattctcttcgttttgtcgccttgtgggcggtctgaggcatcaattgcctctacctcattcacaagcgt
gttgatcaactcaccgattttaccttgctcatcccggttcccccgtttctaagtctctggatatcgtaacacaatcaaagaacaagtgacgattaattatgctgcttt
cgtcacactataagagaacataatagcagaataacaaaggaacacaacaccggaatagatacatattgttctgcgcttatgttctccttatcagcacacaatagtccat
tatacgc

**10x GGC repeat building cassette (Supplementary Figure 1A)**
(annealed pair of DNA oligos; blunt cloned into SmaI digested pJAZZ-OK-delta-AarI)
5’-CACCTGCATATCGGCGGCGGCGGCGGCGGCGGCGGCGGCGGCGGCATGTCTTC-5’
3’-GTGGACGTATAGCCGCCGCCGCCGCCGCCGCCGCCGCCGCCGCCGTACAGAAG-3’

### Supplementary File 3

#### Recombineering and cloning steps to create the humanised C9orf72 mouse BAC

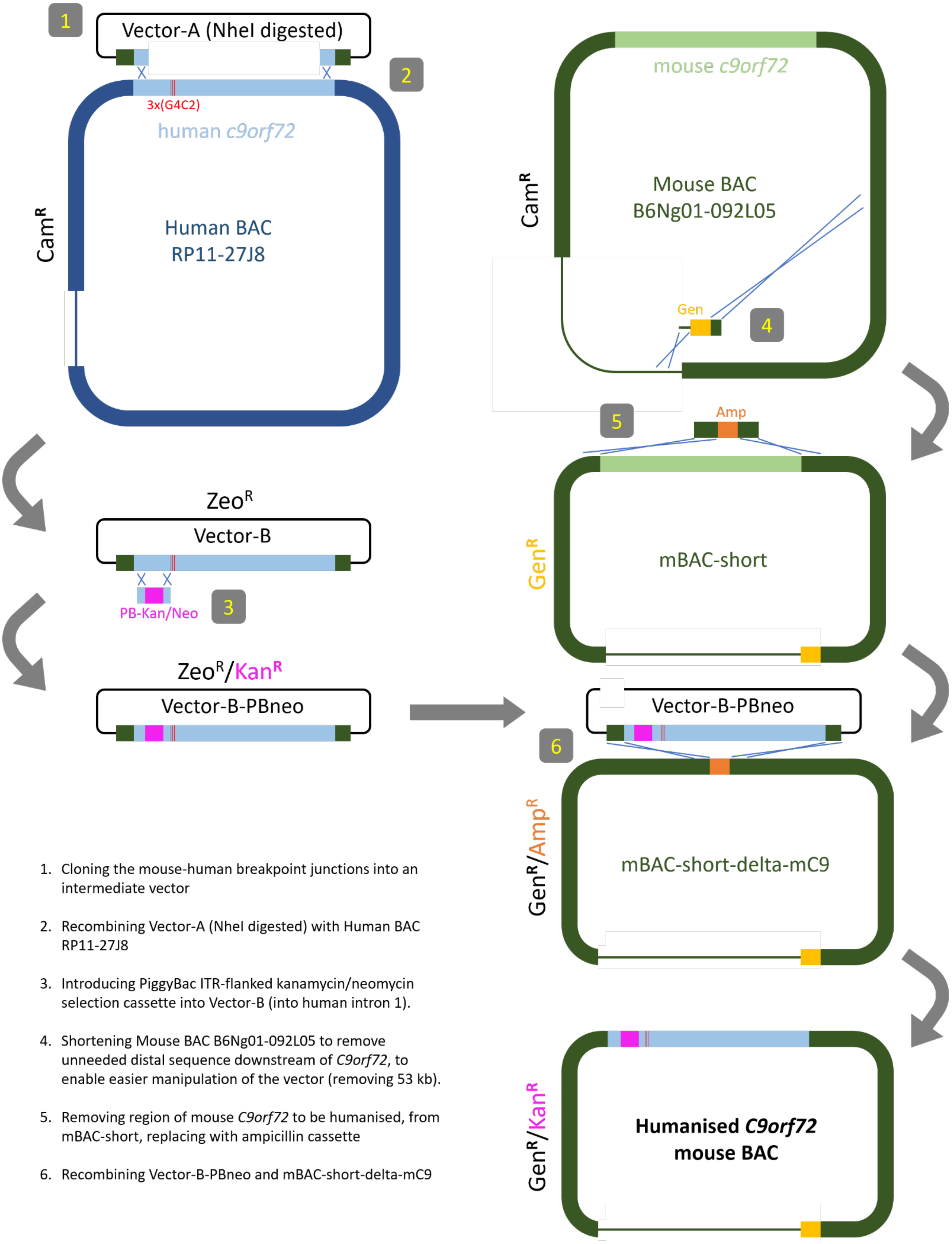

1. Cloning the mouse-human breakpoint junctions into an intermediate vector
2. Recombining Vector-A (NheI digested) with Human BAC RP11-27J8
3. Introducing PiggyBac ITR-flanked kanamycin/neomycin selection cassette into Vector-B (into human intron 1).
4. Shortening Mouse BAC B6Ng01-092L05 to remove unneeded distal sequence downstream of *C9orf72*, to enable easier manipulation of the vector (removing 53 kb).
5. Removing region of mouse *C9orf72* to be humanised, from mBAC-short, replacing with ampicillin cassette
6. Recombining Vector-B-PBneo and mBAC-short-delta-mC9

#### 1. Cloning the mouse-human breakpoint junctions into an intermediate vector

**Modified vector name:** Vector-A (zeocin$^R$)

**Method:** commercial purchase

**Vector:** empty BAC backbone (pSMART BAC; Lucigen)

**Insert sequence**:

**Mouse in green**; **human in blue**: 5′ and 3′ distal parts for humanisation; NheI site needed for next step highlighted; Zeocin antibiotic selection cassette underlined.

ggatccatcgat**CTACCTGCCAACTCCCAAACAAAACAAAGCAAAACTACTTACAACCACAAACTACGCTTCGTAACCTAGATAGATAACGCAGGTGACACTATCTATC**
**TAGGTTGAGCTCAGCTCTGCCCATGCTTTTCCTGAGCGGCTCTTGGAAGAAAAGCTACAAAGCCCATGACAGCCTCCGCCTGGCCAGCTGCCACTGGCATCTCAAGGCT**
**GGCAAAGCAAAGTGAAAGCGCCAACCCGGAACTTACGGAGTCCCACGAGGGAACCGCGGCGTCAAGCAGAGACGAGTTCCGCCCACGTGAAAGATGGCGTTTGTAGT**
**GACAGCCATCCCAATTGCCCTTTCCTTCTAGGTGGAAAGTGGTGTCTAGACAGTCCAGGGAGGGTGTGCGAGGGAGGTGCGTTTTGGTTGCCTCAGCTCGCAACTTAAC**
**TCCACAACGGTGACCAAGGACAAAAGAAGGAAACAAGACTGCAGAGATCCGCACCGGGGAGCCCTGCAGATTCTGGGTGTGTGTTTTTGTTTTTCCCACCCTCTCTCCC**
**CACTACTTGCTCTCACAGTACTCGCTGAGGGTGAACAAGAAAAGACCTGATAAAGATTAACCAGAAGAAAACAAGGAGGGAAACAACCGCAGCCTGTAGCAAGCTCTGG**
**AACTCAGGAGTCGCGCGCTAGCTAGCTGTCATAGAAACTTCTGTACGAGTAGAATGTGAGCAAATTATGTGTTGAAATGGTTCCTCTCCCTGCAGGTCTTTCAGCTGAA**
**ACCTGGCTTATCTCTCAGAAGTACTTTCCTTGCACAGTTTCTACTTGTCCTTCACAGAAAAGCCTTGACACTAATAAAATATATAGAAGACGATACGTGAGTAAAACTC**
**CTACACGGAAGAAAAACCTTTGTACATTGTTTTTTTGTTTTGTTTCCTTTGTACATTTTCTATATCATAATTTTTGCGCTTCTTTTTTTTTTTTTTTTTTTTTTTTTTC**
**CATTATTTTTAGGCAGAAGGGAAAAAAGCCCTTTAAATCTCTTCGGAACCTGAAGATAGACCTTGATTTAACAGCAGAGGGCGATCTTAACATAATAATGGCTCTGGCT**
**GAGAAAATTAAACCAGGCCTACACTCTTTTATCTTTGGAAGACCTTTCTACACTAGTGTGCAAGAACGAGATGTTCTAATGACTTTTTAACCGTGTGGTTTGCTGTGTC**
**TGTCTCTTCACAGTCACACCTGCTGTTACAGTGTCTCAGCAGTGTGTGGGCACATCCTTCCTCCCGAGTCCTGCTGCAGGACAGGGTACACTACACTTGTCAGTAGAAG**
**TCTGTACCTGATGTCAGGTGCATCGTTACAGTGAATGACTCTTCCTAGAATAGATGTACTCTTTTAGGGCCTTATGTTTACAATTATCCTAAGTACTATTGCTGTCTTT**
**TAAAGATATGAATGATGGAATATACACTTGACCATAACTGCTGATTGGTTTTTTGTTTTGTTTTGTTTGTTTTCTTGGAAACTTATGATTCCTGGTTTACATGTACCAC**
**ACTGAAACCCTCGTTAGCTTTACAGATAAAGTGTGAGTTGACTTCCTGCCCCTCTGTGTTCTGTGGTATGTCCGATTACTTCTGCCACAGCTAAACATTAGAGCATTTA**
**AAGTTTGCAGTTCCTCAGAAAGGAACTTAGTCTGACTACAGATTA**gtttaaacatacaattctaccacctgcagcagcacgtgttgacaattaatcatcggcatagtat
atcggcatagtataatacgacaaggtgaggaactaaaccatggccaagttgaccagtgccgttccggtgctcaccgcgcgcgacgtcgccggagcggtcgagttctgga
ccgaccggctcgggttctcccgggacttcgtggaggacgacttcgccggtgtggtccgggacgacgtgaccctgttcatcagcgcggtccaggaccaggtggtgccgga
caacaccctggcctgggtgtgggtgcgcggcctggacgagctgtacgccgagtggtcggaggtcgtgtccacgaacttccgggacgcctccgggccggccatgaccgag
atcggcgagcagccgtgggggcgggagttcgccctgcgcgacccggccggcaactgcgtgcacttcgtggccgaggagcaggactgaggatcc

#### 2. Recombining Vector-A (NheI digested) with Human BAC RP11-27J8

**Modified vector name:** Vector-B (zeocin$^R$)

**Method**: Recombineering

**Vector**: Vector-A (NheI linearised)

**Donating vector:** Human BAC RP11-27J8

**Result**: Full region to be humanised (human intron 1 to ATG start codon) recombineered within Vector-A (at NheI site) via human homology sequences indicated above, and thus flanked by correct mouse sequence breakpoints

#### 3. Introducing PiggyBac ITR-flanked kanamycin/neomycin selection cassette into Vector-B (into human intron 1).

**Modified vector name:** Vector-B-PBneo (zeocin$^R$ and kanamycin$^R$)

**Method:** Recombineering

**Vector:** Vector B

**Donating fragment:**

Human sequence homology arms in **uppercase**. PiggyBac ITR-flanked Kanamycin/Neomycin cassette in lowercase. Endogenous human **TTAA** insertion site underlined.

**TTGCTCTCACAGTACTCGCTGAGGGTGAACAAGAAAAGACCTGATAAAGA**ttaaccctagaaagatagtctgcgtaaaattgacgcatgcattcttgaaatattgctct
ctctttctaaatagcgcgaatccgtcgctgtgcatttaggacatctcagtcgccgcttggagctcccgtgaggcgtgcttgtcaatgcggtaagtgtcactgattttga
actataacgaccgcgtgagtcaaaatgacgcatgattatcttttacgtgacttttaagatttaactcatacgataattatattgttatttcatgttctacttacgtgat
aacttattatatatatatttcttgttatagatatcgcggccgctctagcctcgaggctagaactagtggatctcgagccccagctggttctttccgcctcagaagcca
tagagcccaccgcatcCCagcatgcctgctattgtcttcccaatcctcccccttgctgtcctgccccaccccaccccccagaatagaatgacacctactcagacaatg
cgatgcaatttcctcattttattaggaaaggacagtgggagtggcaccttccagggtcaaggaaggcacgggggaggggcaaacaacagatggctggcaactagaaggc
acagtcgaggctgatcagcgagctctagagctcagaagaactcgtcaagaaggcgatagaaggcgatgcgctgcgaatcgggagcggcgataccgtaaagcacgaggaa
gcggtcagcccattcgccgccaagctcttcagcaatatcacgggtagccaacgctatgtcctgatagcggtccgccacacccagccggccacagtcgatgaatccagaa
aagcggccattttccaccatgatattcggcaagcaggcatcgccatgggtcacgacgagatcctcgccgtcgggcatgcgcgccttgagcctggcgaacagttcggctg
gcgcgagcccctgatgctcttcgtccagatcatcctgatcgacaagaccggcttccatccgagtacgtgctcgctcgatgcgatgtttcgcttggtggtcgaatgggca
ggtagccggatcaagcgtatgcagccgccgcattgcatcagccatgatggatactttctcggcaggagcaaggtgagatgacaggagatcctgccccggcacttcgccc
aatagcagccagtcccttcccgcttcagtgacaacgtcgagcacagctgcgcaaggaacgcccgtcgtggccagccacgatagccgcgctgcctcgtcctgcagttcat
tcagggcaccggacaggtcggtcttgacaaaaagaaccgggcgcccctgcgctgacagccggaacacggcggcatcagagcagccgatcgtctgttgtgcccagtcata
gccgaatagcctctccacccaagcggccggagaacctgcgtgcaatccatcttgttcaatggccgatcccatggtttagttcctcaccttgtcgtattatactatgccg
atatactatgccgatgattaattgtcaacaggctgcaggtcgaaaggcccggagatgaggaagaggagaacagcgcggcagacgtgcgcttttgaagcgtgcagaatgc
cgggcctccggaggaccttcgggcgcccgccccgcccctgagcccgcccctgagcccgcccccggacccaccccttcccagcctctgagcccagaaagcgaaggagcaa

agctgctattggccgctgccccaaaggcctacccgcttccattgctcagcggtgctgtccatctgcacgagactagtgagacgtgctacttccatttgtcacgtcctgc
acgacgcgagctgcggggcggggggaacttcctgactaggggaggagtagaaggtggcgcgaaggggccaccaaagaacggagccggttggcgcctaccggtggatgt
ggaatgtgtgcgaggccagaggccacttgtgtagcgccaagtgcccagcggggctgctaaagcgcatgctccagactgccttgggaaaagcgcctcccctacccggtag
aattaattcgatatcaagcttgctcggatgatagggccctaacgtgtttttgctttgttactttatagaagaaattttgagtttttgtttttttttaataaataaataa
acataaataaattgtttgttgaatttattattagtatgtaagtgtaaatataataaaacttaatatctattcaaattaataaataaacctcgatatacagaccgataaa
acacatgcgtcaattttacgcatgattatctttaacgtacgtcacaatatgattatctttctaggg**TTAACCAGAAGAAAACAAGGAGGGAAACAACCGCAGCCTGTAG
CAAGCTCTGGA**

#### 4. Shortening mouse BAC to remove unneeded distal sequence downstream of *C9orf72*, to enable easier manipulation of the vector (removing 53 kb).

**Modified vector name:** mBAC-short (gentamycin$^R$)

**Method**: Recombineering

**Vector**: Mouse BAC B6Ng01-092L05

**Donating fragment**:

Vector homology arms in **uppercase**. 5' homology arm matches mouse BAC sequence (green). 3' homology arm matches part of the BAC vector backbone. Gentamycin cassette underlined, replacing Chloramphenicol in parent vector backbone.

**TTTAGCTTCCTTAGCTCCTGAAAATCTCGATAACTCAAAAAATACGCCCG**atgttacgcagcagcaacgatgttacgcagcagggcagtcgccctaaaacaaagttagg
tggctcaagtatgggcatcattcgcacatgtaggctcggccctgaccaagtcaaatccatgcgggctgctcttgatcttttcggtcgtgagttcggagacgtagccacc
tactcccaacatcagccggactccgattacctcgggaacttgctccgtagtaagacattcatcgcgcttgctgccttcgaccaagaagcggttgttggcgctctcgcgg
cttacgttctgcccaggtttgagcagccgcgtagtgagatctatatctatgatctcgcagtctccggcgagcaccggaggcagggcattgccaccgcgctcatcaatct
cctcaagcatgaggccaacgcgcttggtgcttatgtgatctacgtgcaagcagattacggtgacgatcccgcagtggctctctatacaaagttgggcatacgggaagaa
gtgatgcactttgatatcgacccaagtaccgccacctaaggaggacaatcactatgtcgggtgcggagaaagaggtaatgaaatgtgctctgtcacctaggGCAACAGC
TTCTGGGACAGGAGATAAAGAGTAAGCTCCTCCAGCCGGAGC

#### 5. Removing region of mouse *C9orf72* to be humanised, from mBAC-short, replacing with ampicillin cassette

**Modified vector name:** mBAC-short-delta-mC9 (gentamycin$^R$ and ampicillin$^R$)

**Method**: Recombineering

**Vector**: mBAC-short

**Fragment to recombine**:

Mouse sequence homology arms in **uppercase green.** Ampicillin cassette in lowercase.

**AAGGAAACAAGACTGCAGAGATCCGCACCGGGGAGCCCTGCAGATTCTGG**aacaaaggcctggtgatgatggcgggatcgttgtatatttcttgacacctttcggcat
cgccctaaaattcggcgtcctcatattgtgtgaggacgttttattacgtgtttacgaagcaaaagctaaaaccaggagctatttaatggcaacagttaaccagctggta
cgcaaaccacgtgctcgcaaagttgcgaaaagcaacgtgcctgcgctggaagcatgcccgcaaaaacgtggcgtatgtactcgtgtatatactaccactcctaaaaaac
cgaactccgcgctgcgtaaagtatgccgtgttcgtctgactaacggtttcgaagtgacttcctacatcggtggtgaaggtcacaacctgcaggagcactccgtgatcct
gatccgtggcggtcgtgttaaagacctcccgggtgttcgttaccacaccgtacgtggtgcgcttgactgctccggcgttaaagaccgtaagcaggctcgttccaagtat
ggcgtgaagcgtcctaaggcttaaggaggacaatcttcaaatatgtatccgctcatgagacaataaccctgataaatgcttcaataatattgaaaaaggaagagtatga
gtattcaacatttccgtgtcgcccttattcccttttttgcggcattttgccttcctgtttttgctcacccagaaacgctggtgaaagtaaaagatgctgaagatcagtt
gggtgcacgagtgggttacatcgaactggatctcaacagcggtaagatccttgagagttttcgccccgaagaacgttttccaatgatgagcacttttaaagttctgcta
tgtggcgcggtattatcccgtattgacgccgggcaagagcaactcggtcgccgcatacactattctcagaatgacttggttgagtactcaccagtcacagaaaagcatc
ttacggatggcatgacagtaagagaattatgcagtgctgccataaccatgagtgataacactgcggccaacttacttctgacaacgatcggaggaccgaaggagctaac
cgcttttttgcacaacatgggggatcatgtaactcgccttgatcgttgggaaccggagctgaatgaagccataccaaacgacgagcgtgacaccacgatgcctgtagca
atggcaacaacgttgcgcaaactattaactggcgaactacttactctagcttcccggcaacaattaatagactggatggaggcggataaagttgcaggaccacttctgc
gctcggcccttccggctggctggtttattgctgataaatctggagccggtgagcgtgggtctcgcggtatcattgcagcactggggccagatggtaagccctcccgtat
cgtagttatctacacgacggggagtcaggcaactatggatgaacgaaatagacagatcgctgagataggtgcctcactgattaagcattggtaa**CCGTGTGGTTTGCTG
TGTCTGTCTCTTCACAGTCACACCTGCTGTTACAG**

#### 6. Recombining Vector-B-PBneo and mBAC-short-delta-mC9

**Modified vector name:** Humanised C9orf72 mouse BAC (gentamycin$^R$ and kanamycin$^R$)

**Method**: Recombineering

**Vector**: mBAC-short-delta-mC9

**Donating vector**: Vector-B-PBneo

**Result**: Humanised *C9orf72* mouse BAC

### Supplementary File 4

(All sequences are orientated 5’-3’)

### Mouse *C9orf72* CRISPR/Cas9 target sites for ES cell gene targeting

5’ CRISPR target site ATTCTGGGTCTGCTGTGGAC
3’ CRISPR target site AGTCGACATCCCTGCATCCC

The full humanisation strategy used only the 5’ CRISPR target site. The intron 1 humanisation strategy used the 5’ and 3’ target sites together.

Oligos for cloning into modified px330:
5’ target site cloning oligos CACCgATTCTGGGTCTGCTGTGGAC & AAACGTCCACAGCAGACCCAGAATc
3’ target site cloning oligos CACCgAGTCGACATCCCTGCATCCC & AAACGGGATGCAGGGATGTCGACTc

### On-locus PCR to detect correct targeting events via gel electrophoresis and Sanger sequencing:

**Two independent long range PCR assays spanning the 5’ homology arm**

Applicable for screening a correct targeting event from both Targeting Constructs
A positive result indicates targeted (not random) locus integration

5’-span-F1 TCGGGATGAAGAGGCAGGTA (mouse sequence outside of targeting construct)
5’-Span-R1 CTTGTTCACCCTCAGCGAGT (human intron 1)
**Amplicon: 5127 bp**

5’-span-F2 GCCAGAGCTGTCACACAGAG (mouse sequence outside of targeting construct)
5’-Span-R2 GGAGAGAGGGTGGGAAAAAC (human intron 1)
**Amplicon: 5108 bp**

**Internal allele assay to detect human sequence, 3’ to the repeat**

hC9internal-F TCAGTACCCGAGCTGTCTCC (human intron 1)
hC9internal-R AAGCTTGGGCTGAAATTGTG (human intron 1)
**Amplicon: 242 bp**

**Internal assay spanning the 3’ mouse-human junction**

(intron 1 humanisation allele only)
hC9orf72-F1 GGGTTAGGGGCCAAATCTCC (human intron 1)
mC9orf72-R1 TGGGAGGTAGAAGCTCAGCT (mouse 3’ homology arm)
**Amplicon: 805 bp**

**Internal assay to detect successful scarless removal of selection cassette**

(forward primer used for Sanger sequencing)
SCR-F GAACTTACGGAGTCCCACGA
SCR-R CAGGCTGCGGTTGTTTCC
**Amplicon: 395 bp**

### Copycount qPCR probe assays:

**<u>Assay to detect humanised *C9orf72* allele</u>**

Spans 5' mouse-human breakpoint

| | |
|---|---|
| hC9orf72-F | AACGGTGACCAAGGACAAA |
| hC9orf72-probe | AGGAAACAAGACTGCAGAGATCCGC |
| hC9orf72-R | GAGTACTGTGAGAGCAAGTAGTG |

**<u>Assay to detect wild-type *mC9orf72* allele</u>**

Spans from 3' end of mouse intron 1 to 5' end of mouse exon 2

Assay will drop if humanised allele present

| | |
|---|---|
| mC9orf72-F | CTATTGCAAGCGTTCGGATAATG |
| mC9orf72-probe | TGGAATGCAGTGAGACCTGGGATG |
| mC9orf72-R | CTTGGCAACAGCAGGAGAT |

**<u>Assay to detect presence of selection cassette with the humanised allele</u>**

Spans from 5' end of human intron 1 to 5' end of selection cassette

Assay will drop if selection cassette is removed.

| | |
|---|---|
| SC-F | CGCTGAGGGTGAACAAGAA |
| SC-probe | TGACGCATGCATTCTTGAAATATTGCTCTC |
| SC-R | GACGGATTCGCGCTATTTAGA |

**<u>Assay to detect PiggyBac transposase transgenic allele at the Rosa26 locus</u>**

| | |
|---|---|
| Rosa26-PB-F | CGGTAAACCGCAAATGGTTATG |
| Rosa26-PB-probe | TCAAACTAAAGGCGGAGTGGACACG |
| Rosa26-PB-R | CATAGGCCACCTATTCGTCTTC |

**<u>Assay to detect wild type Rosa26 locus</u>**

Assay will drop if PiggyBac transposase transgene is present

| | |
|---|---|
| Rosa26-WT-F | TCCCTCGTGATCTGCAAC |
| Rosa26-WT-probe | CAGTCTTTCTAGAAGATGGGCGGGA |
| Rosa26-WT-R | AACGCCCACACACCAGGTTAG |

#### qRT-PCR assays:

**Assay to detect total *C9orf72* transcripts (exon 5/6) in wild type versus *hC9orf72*$^{370}$ mice**

C9 ex5 Fwd CTGTCACGAAGGCTTTCTTCT
C9 ex6 Rev GCTGCCAACTACAACGGAAC

**Assay to detect human *C9orf72* intron 1 transcripts**

C9 intron 1a Fwd CCCCACTACTTGCTCTCACA
C9 intron 1a Rev CGGTTGTTTCCCTCCTTGTT

**GAPDH control assay (IDT)**

GAPDH mouse Fwd/Rev mix Predesigned catalog (Mm.PT.39a.1)
